# Automated assessments can support regional Red Lists but need to be applied with caution: A case study from central Germany

**DOI:** 10.1101/2023.10.31.564918

**Authors:** Alexander Zizka, Indra Starke-Ottich, David Eichenberg, Dirk Bönsel, Georg Zizka

## Abstract

**Aim:** Expert-based regional Red Lists (RL) carry conservation legislation globally. Yet, they are often difficult to reproduce and their regular compilation by a dwindling number of expert assessors burdens many regional conservation authorities. Here we batch-estimate RL indicators and extinction risk for >1,100 plant species and test the potential of this automated approach to support the expert-based RL process.

**Location:** State of Hesse, Central Germany

**Methods:** First, we estimated current population status, short-term population trend, long-term population trend, and extinction risk by binning existing occurrence probabilities modelled at three time slices with cut-off values derived from RL methodology. Subsequently we compared the results with the latest version of the Hessian expert-based RL using summary statistics and selected example species.

**Results:** We find the assessments of extinction risk to agree in c. 60% of the cases, mostly for species not threatened with extinction. Existing mismatch was by one category in most cases, but up to 6 categories in some cases (mean: 1.6 categories). Furthermore, agreement was highest for extreme categories and very abundant species.

**Main conclusions:** Automated assessments were simplistic for many rare and taxonomically challenging species, but we considered them more accurate than the expert assessments for species with intermediate population size and for species of anthropogenic habitats. Furthermore, the automated assessments are particularly informative for the estimation of long-term and short-term population trends, for which experts are often left to guesstimate based on little data.

## Introduction

Regional Red Lists (RL) quantify species’ risk of local extinction and drive conservation action on the state and national level in many countries (Azam et al., 2016; Bennun et al., 2018; Betts et al., 2020; Jiang et al., 2020; Rodrigues et al., 2006). For instance, in Germany, the National Strategy for Biodiversity, a primary policy to counteract an accelerating national biodiversity loss (Habel et al., 2016; Jandt et al., 2022; Kamp et al., 2021), is based partly on RL.

RL follow a simple procedure: Specialist(s) with expertise in the biota of a region assess the extinction risk of species following fixed criteria, and, as a result, deliver a documented extinction risk category for each species. The criteria differ among RL, but are virtually always based on an assessment of habitat availability, range size, population size, and population trends through time or combinations thereof (IUCN Standards and Petitions Subcommittee, 2017; Schippmann, 2020). For instance, the RL of Germany classifies species into one of nine extinction risk categories based on three indicators: current populations status, long-term population trend (percent change in the last 50-150 years) and short-term population trend (10-25 years; Ludwig et al., 2006; Starke-Ottich et al., 2019).

The laborious process to estimate indicators and derive extinction risk assessment based on expert-opinion are the root for multiple challenges for RL (IUCN Standards and Petitions Subcommittee, 2017): First, taxonomic bias. Available expertise for animal and plant groups is proportional to their appeal to scientists and the general public. Therefore, charismatic mammals, amphibians, birds, and, to a lesser extent, flowering plants are better assessed than more elusive groups, such as soil biota and most invertebrates (Cowie et al., 2022). Likewise, even within individual plant and animal groups, assessments are more elaborate for well-studied genera and species. Second, low reproducibility. While expert’s opinion can provide a high-quality interpretation of all available information, individual experts may judge population trends and extinction risk differently and therefore deliver different assessments based on identical data. Third, uncertainty for species with intermediate population size. Estimating population dynamics through time requires detailed data and understanding of a species’ biology. Experts usually have a good understanding for extremely rare and very rare species, because those species are of particular interest to the specialist community and therefore most scrutinized. Likewise, common and very common species can be assumed to not to be threatened with extinction. In contrast, species with intermediate population size challenge expert assessments: on the one hand they are too common to be of particular interest to specialists, but on the other hand they are rare enough, so that unobserved population declines may put them under risk of extinction.

The challenges to RL are exacerbated by the need for regular re-assessment. RL assessments need to be repeated every 10 years to account for changes in species status in a changing world: new threats and the effectiveness of conservation measures need to be quantified to ensure effective species conservation (Rondinini et al., 2014). Since RL are limited by expert availability and often rely on volunteer contributions, many RL are (at least partly) outdated (Cowie et al., 2022; Rondinini et al., 2014). For instance, the last available RL for plants in several German states date to the late 1990ies and early 2000s. This effect is likely to intensify in the future, because the number of available experts is dwindling (Frobel & Schlumprecht, 2013; Starke-Ottich et al., 2019).

A possible remedy to the lack of person-power and data on species’ population status for RL assessments can be to use species distribution models (Cazalis et al., 2022). SDM can estimate species geographic ranges from few geo-referenced occurrence records and environmental variables in the absence of complete sampling for many species at once (Franklin, 2010). SDM-based range estimates can then be used to approximate current population status, and its change over time, which is encouraged by some RL guidelines (Starke-Ottich et al., 2019). SDM encompass a multitude of algorithms from simple climate envelopes to ensemble approaches combining multiple machine learning techniques. In a simple form (Area of Habitat) SDM based analyses are widely used in global RL assessments (IUCN Standards and Petitions Subcommittee, 2017). Despite their flexibility many SDM may be unsuitable to estimate temporal trends in species’ distribution or population for RL due to the nature of the underlying data: If the effort to observe species differs among time slices, as it most often does, inferred trends would be distorted. The FRESCALO (an SDM in the broad sense) estimates the probability of a species occurrence based on environmental similarity and species co-occurrence across neighbouring locations (Hill, 2012), and efficiently corrects for this observation bias by harnessing common species as benchmark to estimate sampling effort and correct for its distortion (Isaac et al., 2014); an approach that has proven suitable for to estimate trends in plant species distribution across Germany (Eichenberg et al., 2021).

The application of modelled species occurrence probabilities by the FRESCALO algorithm or comparable approaches (short as SDM hereafter) for RL is particularly promising in combination with a second class of automated data analysis techniques: automated assessments (AA). AA standardize the interpretation of species occurrence dynamics to batch estimate the extinction risk of many species. Similar to SDM, AA exist in varying complexity (Cazalis et al., 2022; Henry et al., 2023), from simple approaches to estimate range size indicators from species occurrence records (Bachman et al., 2011, 2020; Dauby et al., 2017) to artificial intelligence using a broad range of data (Walker et al., 2022; Zizka et al., 2021, 2022). SDM and AA may be applied individually or in combination to assist Red Listing, and, as data-driven baseline, have the potential to guide assessment effort (Cazalis et al., 2023), increase reproducibility (Cazalis et al., 2022) and reduce the time needed for assessments by an order of magnitude (Silva et al., 2022).

Despite their potential, SDM and AA have only been sporadically used for RL. Likely, the reason is that both methods add complexity to the assessment process and pose methodological hurdles for assessors (which are mostly taxonomic specialists). Furthermore, AA are under active development, so that no standard for their application exists, and their performance for RL is untested. For instance, AA have so far predominantly been developed for the global IUCN Red List (with few academic exceptions, e.g. Ralimanana et al., 2022; Schmidt et al., 2017) and their reliability on the regional scale remains questionable.

Here, we ask the question *“How well can the combination of SDM and AA reproduce expert-based estimates of current population status, long-term and short-term population trends and extinction risk on the regional scale?”*. We mimic the expert-based Red Listing process by batch-applying RL thresholds to approximations of species current population status, long-term and short-term population trend derived from modelled distributions ranges of >1,100 plant species from the state of Hesse in central Germany.

## Methods

### Study area

Hesse is a state in central Germany covering c. 21,100 km²; mid-elevation mountains, and the floodplains of the Rhine and Main rivers are the major habitat types. We chose Hesse as model region for three reasons:

1. A long-lasting tradition of scientific floristic exploration (since the 1600s) and of formal Red Listing of flowering plants (five version of the state’s RL have been published since 1976: Buttler et al., 1996; Hemm et al., 2008; Hessische Landesanstalt für Umwelt, 1976; Kalheber et al., 1980; Starke-Ottich et al., 2019) and therefore many data are available on species populations trends and threat status.
2. The RL of Hesse is compiled for four subregions separately, increasing data availability.
3. Up-to-date distribution models of the Hessian flora exist in high resolution (25 km^2^) and at three time slices (1960-1987, 1988-1996, 1997-2017; Eichenberg et al., 2021).

### Automated regional Red List (ARL)

We generated an automated Red List for the Flora of Hessen (ARL) by estimating current population situation, short-term population trend and long-term population trend (“RL indicators” hereafter) and extinction risk for the four subregions and the entire state by thresholding modelled species distributions.

First, we obtained modelled occurrence probabilities of individual plant species in a 5×5 km^2^ grid across Germany at three time slices: 1960-1987, 1988-1996, 1997-2017 from Eichenberg et al. (2021). The modelled occurrence probabilities were based on 29 million occurrence records (from the whole of Germany) from a variety of sources and equally distributed among the three time-slices. Species occurrence probabilities were then modelled based on ecological similarity from 76 environmental variables and species co-occurrences across grid-cells using the FRESCALO algorithm, accounting for sampling bias. The underlying method may be applied to any geographic area, taxon and time slice, given enough data are available (See Eichenberg et al., 2021 for details of the methodology). For our study, we extracted the state-wide, gridded occurrence probabilities of all species that occupied any Hessian grid cell (Fig. 1A).

**Figure 1.**
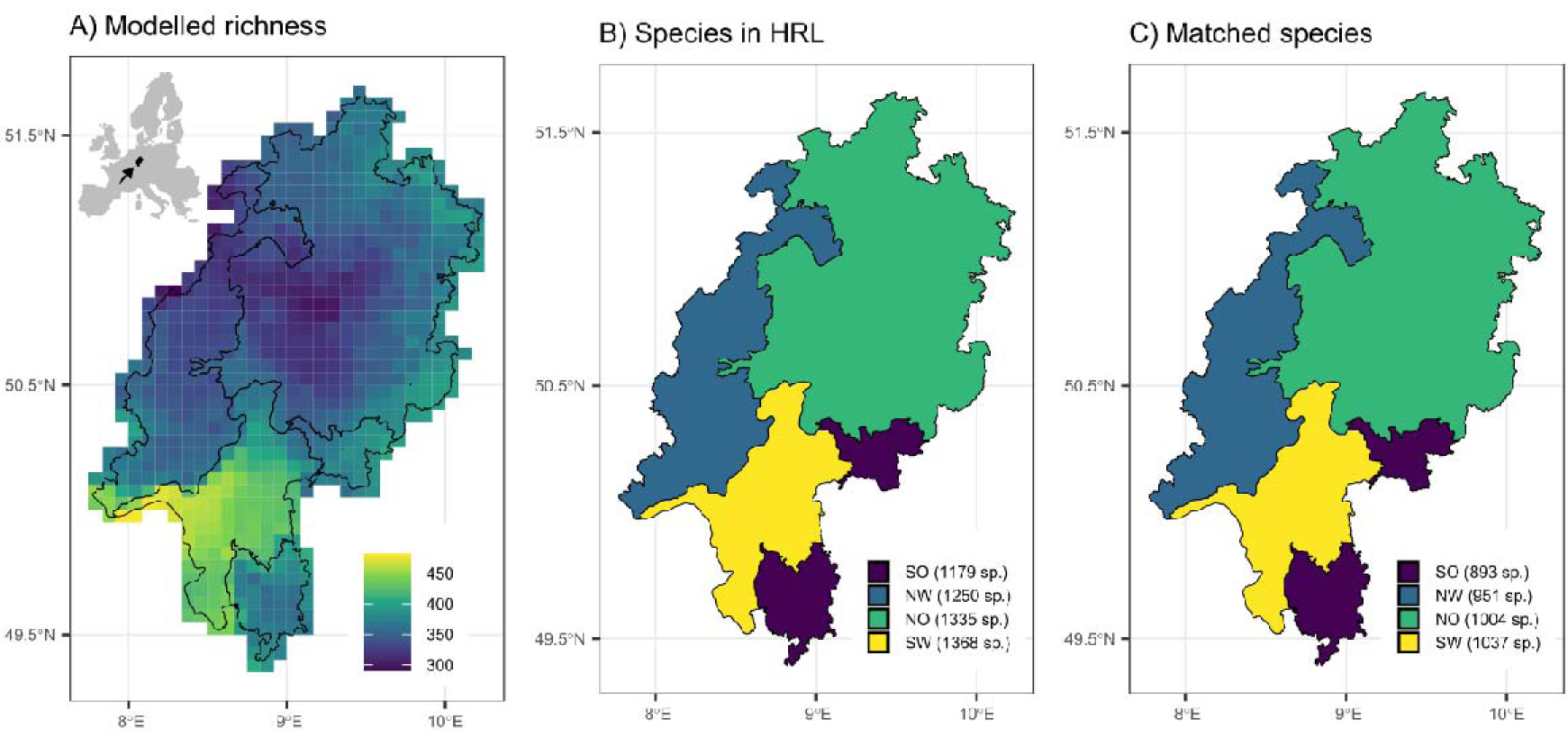
The study area, the state of Hesse in central Germany. **A)** Modelled richness of all species with distribution models available, as the cumulative occurrence probability in 25 km^2^ grid cells. **B)** The number species in each subregion assessed in the 5^th^ version of the Hessian Red List of vascular plants (HRL). **C)** The number of species included in the comparison between automated assessment and HRL. The inlay in A shows the position of the study area in Europe.

Next, we estimated species’ ranges in Hesse at each time slice based on the modelled occurrence probabilities. To do so, we sampled 200 replicates from the occurrence probability of each species in each grid cell at each time-slice with an independent binomial draw and then summarized the mean number of grid cells occupied by in each time-slice per species.

Then, we used the modelled species ranges per time slice to approximate RL indicators: We estimated species current population status by the proportion of grid cells occupied by the species in 1997-2017; the short-term population trend by the percentage of grid-cells lost or gained from 1988-1996 to 1997-2017 (15 years); and the long-term population trend by the percentage of grid-cells lost or gained from 1960-1987 to 1997-2017 (34 years). We then used the cut-off values proposed by the German Federal Ministry for Nature Conservation (Ludwig et al., 2006) and the methodology of the 5^th^ version of the Hessian Red List for vascular plants („HRL“; Starke-Ottich et al., 2019) to convert the numeric values into RL categories for each indicator (Table 1). For instance, a current population status of 12% grid cells occupied translated into the current population status category “rare” (S), and a decrease in cells occupied of 30% from 1988-1996 to 1997-2017 translated into the short-term population change category “strong decrease” (↓↓). Notably, the approximation of population trends from modelled species distributions is a simplification: It may lead to erroneous estimates, if species occur over a relatively large range but in low densities, or in limited areas with large populations. Similarly, a constant geographic range through time may conceal a decline of number of individuals within this range. However, the official Hessian Red List (HRL) explicitly encourages the use modelled species distributions in this way (Starke-Ottich et al., 2019). Additionally, ARL has the advantage of being explicitly defined, documented and hence reproducible, in contrast to expert-opinion based estimates of population trends which often originate from a similar, “range-size-for-population-size” proxy.

**Table 1.** Categories of current population status, long-term trend and short-term trend used in the 5^th^ version of the Hessian Red List for vascular plants (HRL) in the automated assessment in this study (ARL), and the cut-off values used to categorize modelled RL indicators. Risk factors were not used.

| **Current Population status** |  |  |
| --- | --- | --- |
| Symbol | Category name | Percent grid-cells occupied |
| ex | Extinct | 0 |
| es | Extremely rare | 0-1% |
| ss | Very rare | 1-5% |
| s | Rare | 5-15% |
| mh | Moderately common | 15-35% |
| h | Common | 35-70% |
| sh | Very common | 75-100% |
| ? | Unknown | - |
| **Short-term population trend** |  |  |
| Symbol | Category name | Change in cells occupied between 1988-1996 and 1997-2017 |
| ↓↓↓ | Very strong decrease | -100 - -35% |
| ↓↓ | Strong decrease | -35 - -25% |
| (↓) | Moderate decrease, or of unknown magnitude | -25 - -10% |
| = | Stagnant | -10 - +10% |
| ↑ | Distinct increase | > +10% |
| ? | Data deficient | - |
| **Long-term population trend** |  |  |
| Symbol | Category name | Change in cells occupied between 1960-1987 and 1997-2017 |
| <<< | Very strong decrease | -100 - -80% |
| << | Strong decrease | -80 - -55% |
| < | Moderate decrease | -55 - -30% |
| (<) | Decrease of unknown magnitude | - |
| = | Stagnant | -30 - +30% |
| > | Distinct increase | > +30% |
| ? | Data deficient | - |
| **Extinction risk** |  |  |
| Symbol | Category name | Corresponding IUCN category |
| 0 | Extinct or lost | RE |
| 1 | Threatened with extinction | CR |
| 2 | Strongly threatened | EN |
| 3 | Threatened | VU |
| G | Threat of unknown magnitude | [I] |
| R | Extremely rare | [R] |
| V | Near threatened | NT |
| D | Data deficient | DD |
| * | Not threatened | LC |

Finally, we combined the categorical estimates of RL indicators into the seven most relevant extinction risk categories used for the 5^th^ version of expert-based Hessian Red List (0, 1, 2, 3, R, V, *) following the same methodology used in HRL (Starke-Ottich et al., 2019). Each combination of RL indicators translated into a specific extinction risk category. For instance, the combination of current population status “very rare” (ss), long-term population trend “stagnant” (=), and short-term population trend “moderate decrease” (↓), ss=(↓), resulted in the extinction risk category “threatened” (3). Similarly, the combination of “very common” (sh), “moderate decrease” (<) and “distinct increase” (↑), sh<↑, resulted in the extinction risk category “not threatened” (*).

### Expert-based regional Red List (HRL)

HRL is published by Hesse’s Agency for Nature Conservation, Environment and Geology every 10 years and includes assessments of all native species in the state (but neither neophytes nor established archeophytes). In contrast to earlier versions, the 5^th^ version of HRL is based on methodology provided by the German Federal Ministry for Nature Conservation for Red Listing (Ludwig et al., 2006) to increase comparability with other German states. The next paragraphs give a short overview on the methodology of HRL, see Ludwig (2006) and Starke-Ottich et al. (2019) for details.

HRL treated four subregions within Hesse separately, including Northwest (NW), Northeast (NO), Southwest (SW), Southeast (SO) (Fig. 1; Buttler et al., 1996). The regions represented dominant landscape types in the state (Klausing, 1988), and therefore differed in size: NO covered 52% of Hesse, SO covered only 9%. While regions NW, NO, and SO principally comprised mid-elevation mountain habitats, region SW mostly comprised the lowlands of the Rhine and Main river floodplains.

For each subregion and species, HRL provided estimates of current population status (extinct to very common), short-term population trend (very strong decrease to strong increase), long-term population trend (very strong decrease to strong increase) (Table 1) based on the evaluation of a total of 12 experts. The relevant time-span for the long-term population trend was 50-150 years. Assessments for this period were based on information from herbarium collections and literature. For the short-term population trend the relevant time span was 10-25 years, and assessment was based on changes compared to the 4^th^ version of the HRL (Hemm et al., 2008). HRL only provided RL indicators for the subregions, but not for the entire state.

Based on the three RL indicators HRL classified each species in each subregion into one of nine extinction risk categories (Ludwig et al., 2006): extinct or lost (0), threatened with extinction (1), highly threatened (2), threatened (3), threat of unknown extent (G), extremely rare (R), near threatened (V), not threatened (*) and data deficient (D). The same extinction risk categories are provided for the entire state based on the most positive evaluation in a region. While the methodology (Ludwig et al., 2006) in principle allowed the inclusion of specific risk factors, these were not used for HRL.

### Comparing HRL and ARL

To compare ARL and HRL, we first harmonized the taxonomic reference between both lists since the data underlying the ARL used a different taxonomic backbone than HRL (HRL: Jäger, 2017; ARL: GermanSL; Jansen & Dengler, 2008 version 1.4, https://germansl.infinitenature.org/). Next, we removed species which we found incomparable between HRL and ARL including: (1) subspecies and varieties; (2) hybrids; (3) species which names we could not harmonize between HRL and ARL because one name in one list corresponded to multiple accepted names in the other list; (4) taxonomically difficult groups, for instance apomictic species in *Rubus* and *Taraxacum*; (5) species in HRL with no modelled distribution available from Eichenberg et al. (2021); (6) species predicted as present in Hesse by ARL, but that were never considered present in Hesse by the HRL (species formerly present and now considered extinct in HRL are included in the analyses); (7) species that were considered present in Hesse by HRL but never by ARL.

To test the agreement between ARL und HRL, we lumped cases from all subregions and compared the accuracy and magnitude of error for RL indicators and extinction risk. The magnitude of error was the mismatch in number of ordinal classes, e. g. HRL: * and ARL: V amounted to an error of 1, HRL: * and ARL: 1 amounted to an error of 4 (Table 1). To calculate accuracy, we excluded any instances (an instance is the estimate of an RL indicator or extinction risk in a specific region) classified as “?” in HRL or ARL. Likewise, for calculating the accuracy of the current population status we excluded species classified as “common” (h) in ARL, since this category was not used by HRL (which only uses “moderately common”, mh). Lumping the “h” category with the categories “sh” or “mh” was not advisable, because records in “h” could either be close to “sh” or “mh”. Additionally, to calculate the magnitude of error, we excluded instances classified in category “R”, because this category was not easily set in ordinal relation with the other categories. Finally, we compared the modelled percentage of grid-cells covered for current population status, and the percentage change in coverage for short-term population trend and long-term population trend with the categorized estimates from HRL. We also compared the match of the extinction risk assessment between ARL and HRL for species on the state level (RL indicators are not provided on the state level by HRL).

To identify systematic reasons for ARL-HRL mismatch, we first compared the accuracy of estimation and magnitude of error among the current population status classes from HRL (expected higher uncertainty for species with intermediate population size). Furthermore, we identified instances for which ARL and HRL disagreed most to discuss possible improvements of ARL and the integration of ARL into the next version of HRL. We defined instances as “disagreeing most”, if: they were considered as “not threatened” (*) in one list and as one of “extinct” (0), “threatened with extinction” (1), or “extremely rare” (R) in the other list (160 instances); or their summed magnitude of error across populations indicators was more than 5 categories (82 instances); or ARL predicted a positive short-term or long-term trend, since these are rare in HRL and only one expert used them at all (395 instances).

We consider identifying mismatches as the major contribution of automated assessments to RL in the future (besides increased reproducibility), since it will allow a more efficient use of effort by pointing experts to species which need increased scrutiny. Unfortunately, identifying mismatches is also unsatisfying as it leaves the central question “Which one is right: HRL or ARL?”, unanswered. To give a rough hint on the answer to this question, we mimic what we anticipate as future standard procedure, and use our expertise of the local flora (including one contributing expert and the lead editor of HRL) to loosely score either HRL or ARL as “likely more accurate” for cases in which both approaches disagreed most. To do so, we explicitly consider new data which has emerged since 2019 (including on www.inaturalist.org) when the last version of HRL was published.

### Software used

We did all analyses in R (R Core Team, 2022) and used the caret (Kuhn, 2022), cowplot (Wilke, 2020), plotly (Sievert, 2020), tidyverse v1.3.0 (Wickham et al., 2019), readxl v1.3.1 (Wickham & Bryan, 2022), viridis v0.5.1 (Garnier et al., 2021), and writexl (Ooms, 2021) packages.

## Results

We obtained modelled occurrence probabilities for 1,255 species in Hesse (Fig. 1A) and information for 1,933 taxa from HRL (Fig. 1B). During matching we removed a total of 1,040 taxa with 4,111 instances. Of these, 147 taxa (730 instances) were subspecies or varieties; 39 species (195 instances) were hybrids; 38 species (190 instances) couldn’t be harmonized between HRL and ARL; 236 species (1,180 instances) belonged to taxonomically difficult groups; 259 species (1,295 instances) had no modelled distribution; 348 species (519 instances) were not considered present in Hesse by HRL. In the end we retained 1,139 species with 5,023 instances (Fig. 1C) for the comparison of HRL and ARL, corresponding to 40.8% of the 2,790 taxa included in HRL (see Supplementary material S1 and S2 for a full list of matched species and Supplementary material S3 for a list of removed taxa).

For the current population status our analyses included 3,848 instances of which ARL and HRL agreed for 1,420 instances (36.9%, Fig. 2A). When excluding the 729 instances with assessment “h” in ARL (See Methods), a total of 3,119 instances remained, with 1,420 instances of identical assessment (44,3%). In a striking 1,539 instances ARL suggested a higher current population status than HRL (49,3%; magnitude of error of 1 category in 1,226 instances/39,3%, 2 categories: 285/9,1%, 3 categories: 14/0,4%, 4: 4/0,1%, Fig. 2A). In contrast, ARL suggested a lower current populations status in 170 instances (5,5%; magnitude of error of 1 category in 150 instances/4,8%, 2 categories: 16/0,5% and 3 categories: 4/0,1%).

**Figure 2.**
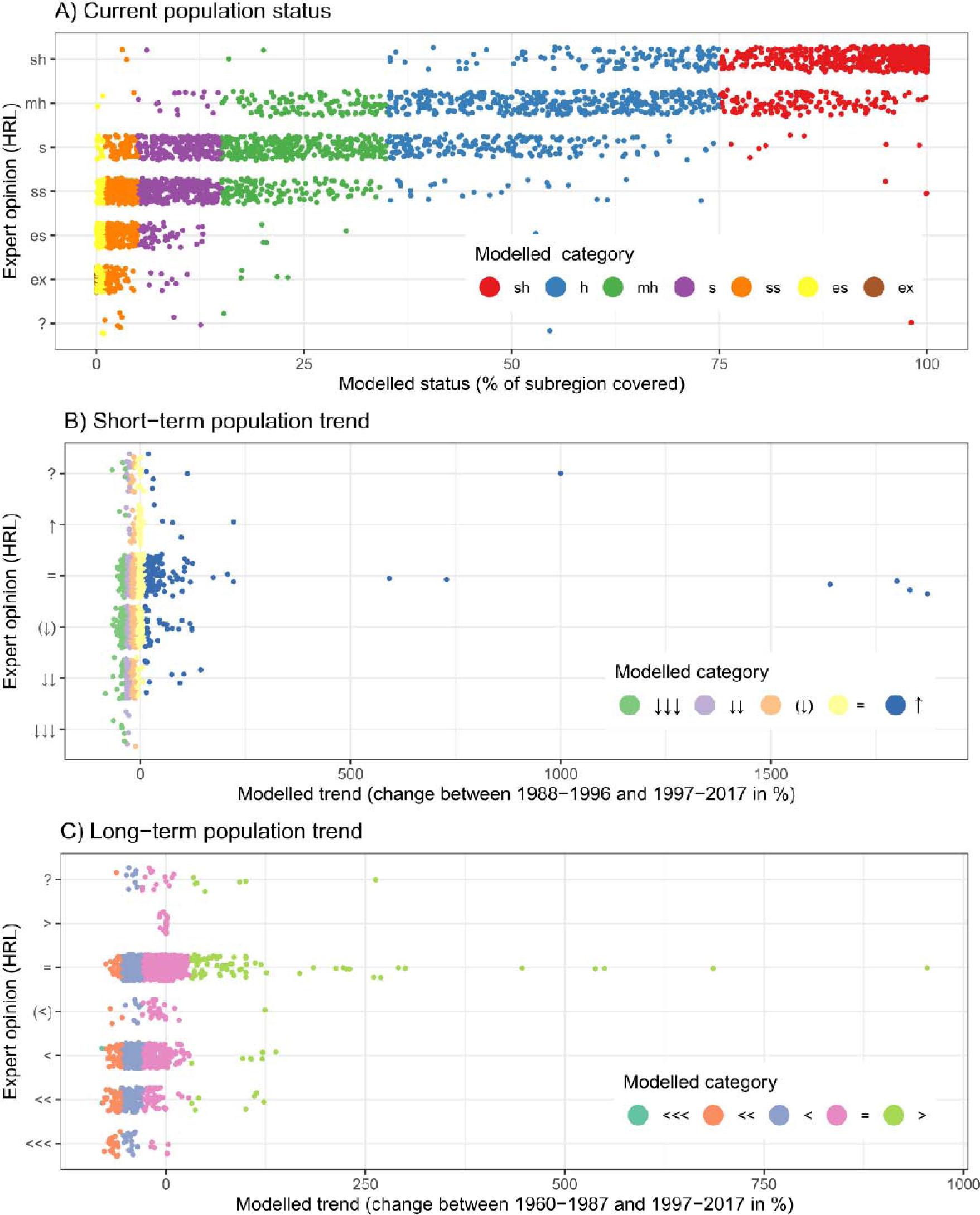
Comparison of expert-based (HRL) and modelled (ARL) RL indicators. **A)** Current population status, **B)** Short-term population trend, C) Long-term population trend. Each datapoint represents an instance (species per subregion). Supplementary material S6-S8 are interactive versions of the subfigures indicating the species names and modelled distribution for each datapoint.

For the short-term population trend our analyses included 3,463 instances, of which HRL and ARL agreed in 1,723 instances (nearly 50%, Fig. 2B). In 471 instances ARL suggested a more positive short-term population trend than HRL (magnitude of error of 1 category in 404 instances/11,7%, 2 categories: 57/1,65%, 3 categories: 9/0,3%). In contrast, in 1,253 instances ARL suggested a more negative trend than HRL (magnitude of error of 1 category in 865 instances/25,1%, 2 categories: 286/8,3%, 3 categories: 100/2,9%, 4 categories: 2/0,05%, Fig. 3). The mismatch was particularly severe in the HRL categories (V) and =, for which modelled assessment span from ↑ to ↓↓↓ (Fig. 2B).

**Figure 3.**
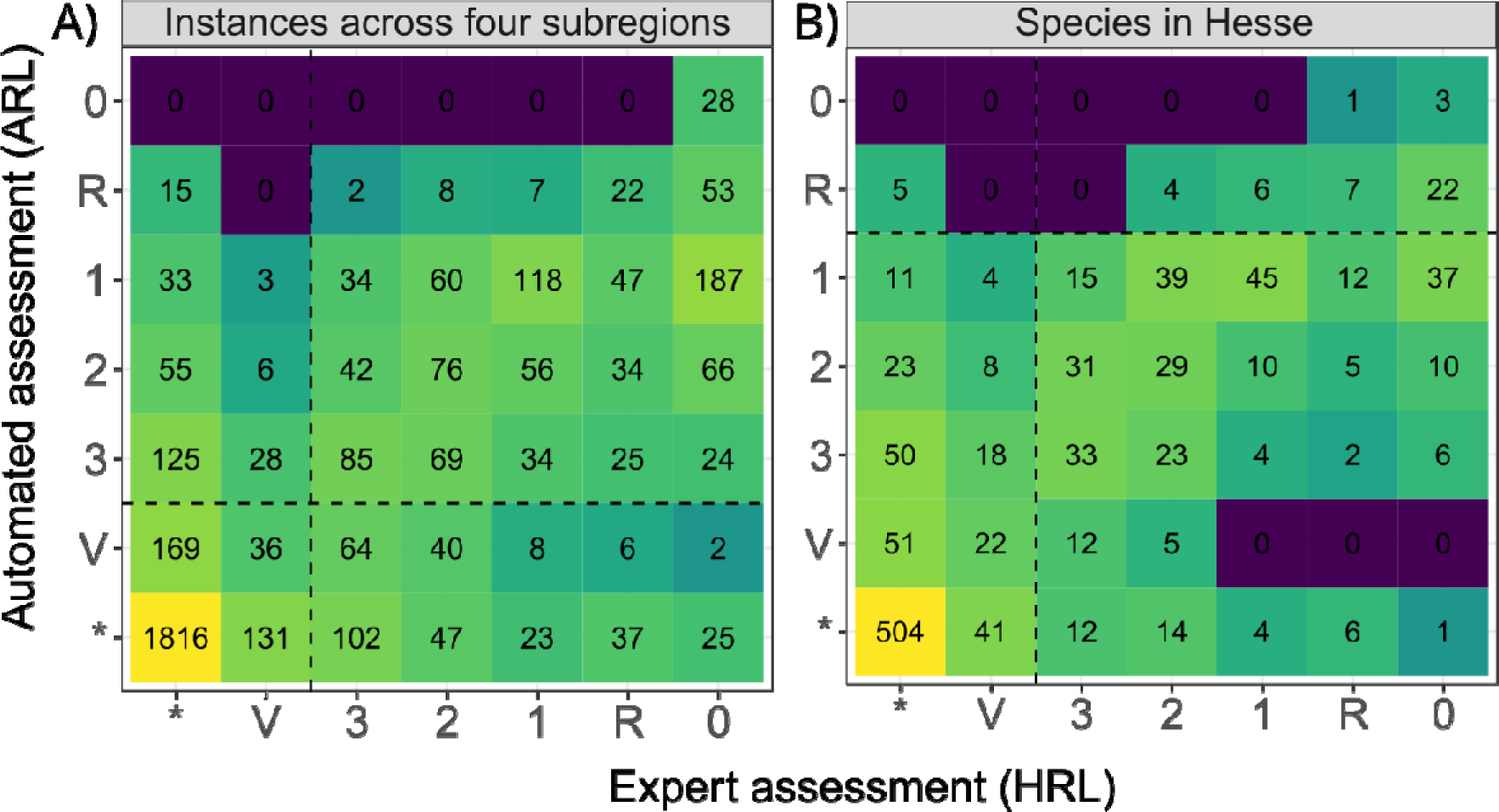
Error matrix comparing the extinction risk estimates from our automated assessment (ARL) with the 5^th^ version of the Hessian Red List of vascular plants (HRL). A) Extinction risk assessments for species per subregion (instance). B) State-wide extinction risk for 1,135 species. Numbers and colours indicate the number of species in each category. See Supplementary material S11 for an interactive versions of subfigure B indicating the species names.

For the long-term population trend our analyses included 3,463 instances, of which HRL and ARL agreed in 2,237 instances (65%), 1,902 of them in category “=” (Fig. 2C). ARL suggested a more positive long-term population trend than HRL in 550 instances (17,7%; magnitude of error of 1 category in 462 instances/13,4%, 2: 75/2,2%, 3: 13/0,4%). In contrast, ARL suggested a more negative trend in 655 instances (19%; magnitude of error of 1 category in 609 instances/17,7%, 2 categories: 43/1,2%, 3 categories: 3/0,1%).

For extinction risk assessments our analysis included 3,452 instances, of which ARL and HRL agreed in 2,741 instances on the binary threatened/not threatened level (79%, Fig. 3A) and in 2,147 instances (62%) on the detailed level of extinction risk categories, 1,816 of them in the category not threatened (*). ARL suggested a lower extinction risk than HRL in 567 instances (16.5%). In contrast, ARL suggested a higher extinction risk than HRL in 555 instances (16%); 6% of instances were considered in extinction risk category (R). In cases of mismatch between ARL and HRL, the mean magnitude of error was 1.65 categories. The agreement in extinction risk assessments between ARL and HRL was similar in all regions, with 60% in NO and NW, 63% in SW and 66% in SO.

For the extinction risk across the entire state, our analysis included 1,135 species, of which ARL and HRL agreed for 847 species on the binary threatened/non-threatened level (229 threatened and 618 not threatened; accuracy of ARL: 0.84, Fig. 3B). HRL considered 288 of species as threatened (3: 103, 2:114, 1: 69) and 737 not threatened (*: 644, V: 93), 33 species as rare (R), 2 species as data deficient (?), and 79 species as extinct (ex). In contrast, ARL considered 415 species threatened (3: 136, 2: 116, 1: 163), 676 species not threatened (*: 585, V: 91), 44 species as rare, and 4 species as extinct for Hesse. Concerning the detailed extinction risk categories, the ARL and HRL agreed on 643 species (56,7%). For 266 species (23,4%) ARL suggested a higher extinction risk than HRL (magnitude of error of 1 category: 146 species/12,9%, 2 categories: 77/6,8%, 3 categories: 27/2,4%, 4 categories: 11/0,9%, and 5 categories: 5/0,4%). In contrast, for 226 species (20%), ARL suggested a lower extinction risk than HRL (magnitude of error of 1 category: 120 species/10,6%, 2 categories: 63/5,6%, 3 categories: 26/2,3%, 4 categories: 10/0,9%, 5 categories: 6/0,5%, and 6 categories: 1/0,09%). Notably, ARL considered more than double the number of species in the highest threat category than HRL. Per class sensitivity varied between 0.26 and 0.79 and was much higher for the extreme classes (* = 0.79, 1 = 0.71) as compared to the intermediate classes (V: 0.24, 3: 0.32, 2: 0.26).

For the relation between current population status and ARL-HRL agreement of extinction risk estimates, we found the overall agreement across regions to be the lower, the rarer a species was (Fig. 4). The agreement between ARL and HRL was high for very common species (775 out of 778 instances, accuracy: 0.99) and common species (550 out of 653 instances, accuracy: 0.84) but decreased to 91 out of 332 species for extremely rare species (accuracy: 0.34). Similarly, the mean magnitude of error was the higher, the rarer a species was, and increased from 1.3 for very common species to 1.8 for extremely rare species (Fig. 4).

**Figure 4.**
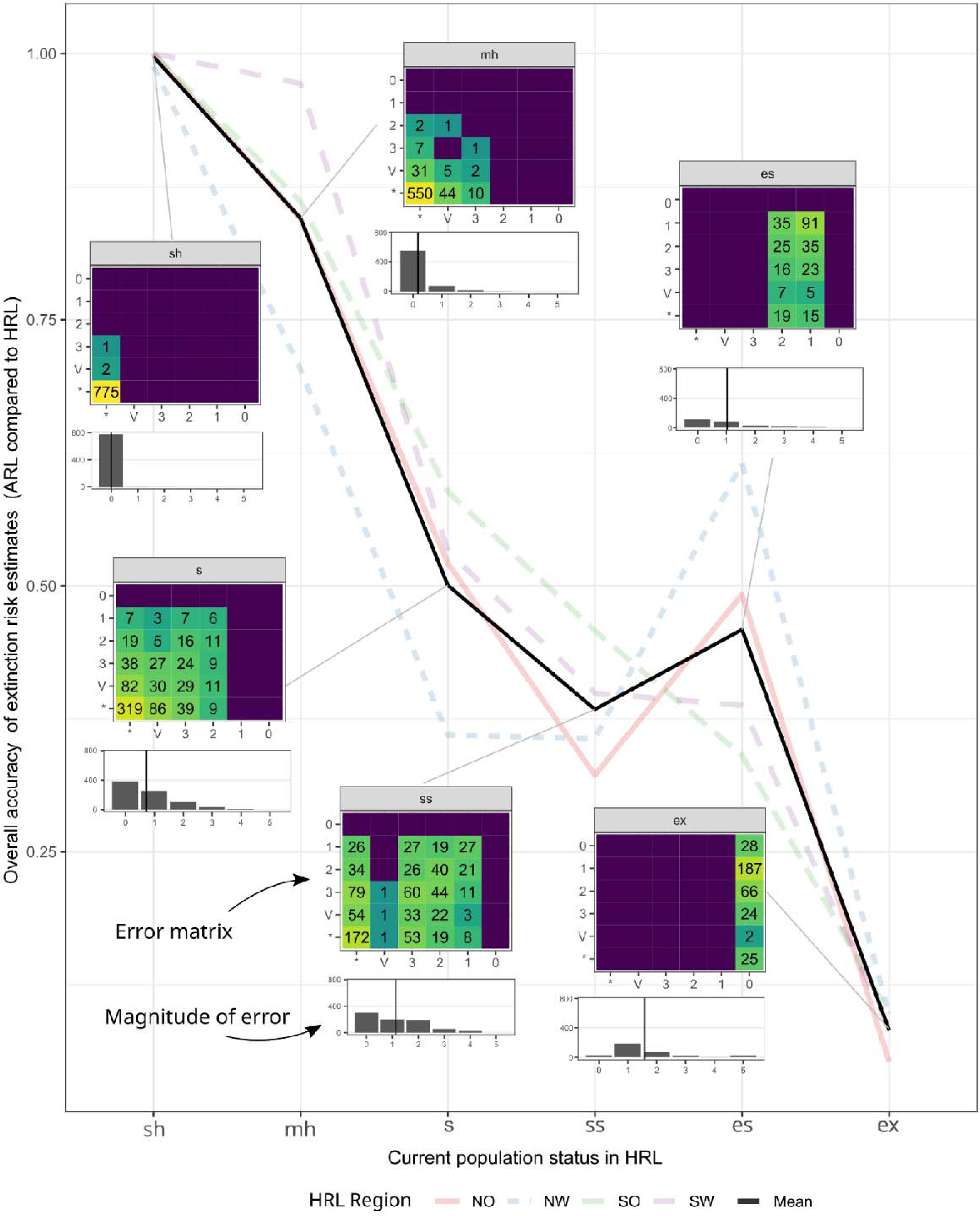
The accuracy of ARL compared to HRL depending on the species current population status. Inlets show the error matrices for each current population status category (HRL on the x-axis and ARL on the y-axis) and the distribution of the magnitude of error (by how many categories the assessments disagree in how many cases). Data from all subregions are shown together.

For the species which disagreed most between we judged ARL to estimate the current population situation as more accurate in 246 cases and HRL to be more accurate in 536 cases (Supplementary material S4).

## Discussion

Here we compared estimates of RL indicators and extinction risk from an Automated Red List (ARL) based on species distribution models and automated assessments with a traditional expert-based Red List (HRL) in the state of Hesse, a well-documented study area in central Europe. We found both approaches to agree for extinction risk in between 57%-62% of cases and the estimates for individual RL indicators consistent in around half of the cases (current population status: 44,3%, short-term trend: 50%, long-term trend: 65%, Fig. 2). In many cases the mismatch was small (difference of 1 category in 23,5% of the instances) but could be considerable in some cases (up to 6 categories, Fig 3). We found the discrepancies between ARL and HRL to increase with species rarity (Fig. 4).

In the following paragraphs, we use our results to discuss possible improvements of ARL and the potential to integrate ARL in the next version of HRL (due 2029) as well as its potential for expert-based RL in general, along six themes.

### Very Rare species (“es” and “ss”)

Examples from this category often occured in forests or dry lawns, such as *Vicia sylvatica* (HRL: *; ARL: 2), *Iris sibirica* (HRL: 2; ARL: *), *Moenchia erecta* (HRL: 2; ARL: *), and *Hepatica nobilis* (HRL: *, ARL: 2). In addition to a potential mis-approximation of population trends by range size (see Methods) a main reason for mismatch of this category is that documented occurrence records are often scarce for rare species (which by definition are found less frequently or may be particularly protected and hence not be documented). The performance of SDM, including the FRESCALO algorithm, depends to the amount of data available for model training (Wisz et al., 2008). Hence, a lower performance of ARL is expected for rare species. Additionally, the modelled occurrence probabilities used for ARL were based on distribution data from across Germany, so that edge effects may cause the model to predict species that only occur in neighbouring states as present in Hessen.

In the future, the limitation from data availability will be remedied by rapidly increasing data from ongoing efforts to compile existing data (e.g. https://www.gbif.org, https://land.gbif.de/) and to harness verified data from citizen science platforms (e.g., https://www.inaturalist.org/, https://floraincognita.com/) which will continuously improve ARL performance for rare species by increasing the data basis for SDM. Additionally, ARL performance can increase by using SDMs tailored to the study area. Such models may be implemented as part of an integrated pipeline for automated regional red listing (comparable to what sRedList does for the IUCN Red List, https://sredlist.eu), and additionally be combined with predictive automated assessment methods which can directly process incomplete information on population size and trends together with species functional traits and evolutionary relationship (Cazalis et al., 2022).

### Species with intermediate population size (s, mh, h)

Examples for this category for which ARL and HRL differed in the estimation of current population status are *Salvia pratensis, Populus alba* (HRL: ss, ARL: h), *Arum maculatum, Dipsacus fullonum, Echium vulgare, Primula veris* (HRL: s, ARL: h), *Digitaria ischaemum, Saponaria officinalis, Sedum album* (HRL: ss, ARL: mh). Likely, a reason for the mismatch is limited data availability to experts and a relatively strong performance of ARL. Experts can usually assign very rare and very common species confidently to extinction risk categories and approximate trends in their population status. In contrast, recognizing trends in rare to common species is more challenging, since less information is available to experts, because they usually receive less attention (Starke-Ottich et al., 2019) but often are particularly threatened (Jansen et al., 2020). Hence, these species may increase or decrease substantially before changes are picked up by experts, losing valuable time for conservation. In the future, ARL estimates of RL indicators have the potential to serve as data-driven, reproducible baseline to inform experts during the assessment process in particular for long-term and short-term population trend.

### Species of azonal (small-scale) habitats

Examples in this category are species from wet meadows, bogs, flooding meadows, dry lawns (following www.floraweb.de) which ARL often considered less threatened than HRL, such as *Equisetum pratense* (HRL: 3, ARL: *), *Carex flava, Iris sibirica, Moenchia erecta* (HRL: 2, ARL: *), *Crepis pulchra* (HRL: R, ARL: *). There may be multiple non-mutually exclusive explanations for this mismatch. First, these habitats have received considerable conservation and research attention by specialists in the last years making HRL particularly precise and accurate. Second, these habitats are often protected and, in some cases, hard to access, so that particularly few collection and observation records (but see Sperle & Bruelheide, 2021) as basis for the modelled distributions exist. Third, modelling the ranges for species of azonal, small-scale habitats is challenging. Our SDM approach is comprehensive, as it includes 76 variables to measure ecological similarity and includes species co-occurrences to model occurrence probabilities among grid cells (Eichenberg et al., 2021), but due to the 25 km^2^ resolution it may fail to capture microhabitat conditions and therefore systematic underestimate the occurrence probability of species of azonal habitats. In the future higher resolution SDM may improve the match between ARL and HRL, as could the relaxation of the neighbourhood criterion in the modelling algorithm, yet ARL may remain of limited use for these species.

### Species in taxonomically difficult groups and aggregate species

Examples in this category are *Alchemilla vulgaris* (NW: ARL: sh, HRL: s)*, Bistorta officinalis* (SW: ARL: h, HRL: ss)*, Carex flava* (SO: ARL: h, HRL: ss; SW: ARL: mh, HRL: es), and *Galium mollugo* (SW: ARL: sh, HRL: ss). The reasons are likely an unsatisfactory taxonomic matching between HRL and ARL and the high identification uncertainty for these species. Although we have taken great care to resolve the taxonomies between the two datasets and have excluded particularly complex groups from the analyses, the correct assessment remains a challenge. Another interesting example is *Nymphaea alba*, where a high percentage of recorded occurrences was of hybrid origin or even misidentified, which is correctly reflected in the HRL but not ARL (Starke-Ottich et al., 2019). As a route forward for ARL, further effort for taxonomic harmonisation before analyses may remedy this problem, yet it will likely persist, since a correct identification of species in taxonomic difficult groups cannot be guaranteed when using data from multiple sources including for instance citizen science.

### Species related to cultivation or anthropogenic habitats

Examples in this category for which ARL overestimates extinction risk (as compared to HRL and our post-hoc expert judgement) are *Juglans regia*, *Setaria viridis*, *Setaria pumila*, and *Solanum nigrum*. One reason for this systematic error was the low number of occurrence records included as basis for the modelling in ARL for these species, likely caused by the disregard of data collectors due to an ascribed low value or relevance for RL. In contrast, examples of species occurring naturally in agricultural fields or ruderal or urban habitats for which HRL underestimated current distribution range, were *Anchusa officinalis, Barbarea stricta, Bryonia dioica, Filago germanica, Bupleurum rotundifolium, Galeopsis ladanum, Geranium molle, Odontites vernus*. An explanation could be, that especially urban habitats and their flora receive less attention from experts in the course of Red list assessments. These results suggest, that currently ARL estimates can supplement expert-based assessments for species native in anthropogenic habitats, but are of limited value for cultivated species. In the future, information on cultivation or habitat could be used to exclude species a priori from ARL or more sophisticated ARL algorithms could incorporated this information in the prediction.

### Quantifying uncertainty and methodological considerations

All Red Listing approaches rely on arbitrarily chosen thresholds, either to convert continuous modelled distributions (in the case of ARL) or abstract expert knowledge (in the case of HRL) into the distinct categories required for Red Lists. While we have carefully chosen the thresholds for ARL following the HRL methodology as much as possible, the guidelines are necessarily vague (since they ought to apply to many taxa) and different thresholds could change the results of ARL. Compared to HRL, ARL has the advantage, that the thresholds are clearly documented and therefore the assessments are reproducible. Furthermore, it is conceivable to statistically quantify this uncertainty in the future, for instance by integrating assessments over multiple thresholds.

The definition of the reference time points for estimating short term and long-term population trends add uncertainty. There are several reasons for this, which affect HRL and ARL equally. First, less data is available further back in time. The recommended baseline for the long-term trend is 50-150 years (Ludwig et al., 2006). This is conceptually challenging for ARL, where theoretically some, but considerably less data exist from this time as basis for modelling, and environmental variables used for modelling may have changed, but likely even more severe for HRL, where experts may have some experience on the upper bound of this interval and general information from specimen or (often grey) literature may be accessible (e.g., Händler, 2019). In practice, for ARL we had to rely the first time slice with modelled species distributions available, which was from 1960-1987 as baseline for the long-term trend (on average 50 years ago at the time of the analyses).

## Conclusion

While we consider expert assessments more accurate in many cases, particularly for very rare species and taxonomic difficult groups, our results indicate that the combination of SDM and automated assessments can provide valuable, reproducible input to the expert-assessment process, particularly for estimating short-term and long-term population trend and for species with intermediate abundance or from anthropogenic habitats. The value of automated assessment will potentially increase in the future, as more distribution data become available and if quantification of uncertainty and predictive approaches were to be included in an analyses pipeline specifically tailored for regional Red Lists. In Supplementary material S5-S11 we provide interactive graphs to identify differences between the automated assessment and expert-based regional RL to facilitate the integration of automated assessments into the regional Red List.

## Supporting information

Supplemental Table 1 - Full dataset

Supplemental Table 2 - Metadata for Table S1

Supplemental Table 3 - Instances removed from either dataset before analysis

Supplemental Table 4 - Expert evaluation of assessmetn quality

Supplemental Figure 1 - Interactive version of figure 2A

Supplemental Figure 2 - Interactive version of figure 2B

Supplemental Figure 3 - Interactive version of figure 2C

Supplemental Figure 4 - Interactive confusion matrix current state

Supplemental Figure 5 - Interactive confusion matrix short-term trend

Supplemental Figure 5 - Interactive confusion matrix long-term trend

Supplemental Figure 5 - Interactive Figure 3B

## Data accessibility statement

The data are available in Supplementary material S1 and S2. All modelled occurrences and analysis scripts will be available from a zenodo repository upon publication.

