## Supplemental Figure 1 - Interactive version of figure 2A for "Automated assessments can support regional Red Lists but need to be applied with caution: A case study from central Germany": plotly.js

```
HTMLWidgets.widget({
  name: "plotly",
  type: "output",

  initialize: function(el, width, height) {
    return {};
  },

  resize: function(el, width, height, instance) {
    if (instance.autosize) {
      var width = instance.width || width;
      var height = instance.height || height;
      Plotly.relayout(el.id, {width: width, height: height});
    }
  },  
  
  renderValue: function(el, x, instance) {
    
    // Plotly.relayout() mutates the plot input object, so make sure to 
    // keep a reference to the user-supplied width/height *before*
    // we call Plotly.plot();
    var lay = x.layout || {};
    instance.width = lay.width;
    instance.height = lay.height;
    instance.autosize = lay.autosize || true;
    
    /* 
    / 'inform the world' about highlighting options this is so other
    / crosstalk libraries have a chance to respond to special settings 
    / such as persistent selection. 
    / AFAIK, leaflet is the only library with such intergration
    / https://github.com/rstudio/leaflet/pull/346/files#diff-ad0c2d51ce5fdf8c90c7395b102f4265R154
    */
    var ctConfig = crosstalk.var('plotlyCrosstalkOpts').set(x.highlight);
      
    if (typeof(window) !== "undefined") {
      // make sure plots don't get created outside the network (for on-prem)
      window.PLOTLYENV = window.PLOTLYENV || {};
      window.PLOTLYENV.BASE_URL = x.base_url;
      
      // Enable persistent selection when shift key is down
      // https://stackoverflow.com/questions/1828613/check-if-a-key-is-down
      var persistOnShift = function(e) {
        if (!e) window.event;
        if (e.shiftKey) { 
          x.highlight.persistent = true; 
          x.highlight.persistentShift = true;
        } else {
          x.highlight.persistent = false; 
          x.highlight.persistentShift = false;
        }
      };
      
      // Only relevant if we haven't forced persistent mode at command line
      if (!x.highlight.persistent) {
        window.onmousemove = persistOnShift;
      }
    }

    var graphDiv = document.getElementById(el.id);
    
    // TODO: move the control panel injection strategy inside here...
    HTMLWidgets.addPostRenderHandler(function() {
      
      // lower the z-index of the modebar to prevent it from highjacking hover
      // (TODO: do this via CSS?)
      // https://github.com/ropensci/plotly/issues/956
      // https://www.w3schools.com/jsref/prop_style_zindex.asp
      var modebars = document.querySelectorAll(".js-plotly-plot .plotly .modebar");
      for (var i = 0; i < modebars.length; i++) {
        modebars[i].style.zIndex = 1;
      }
    });
      
      // inject a "control panel" holding selectize/dynamic color widget(s)
    if ((x.selectize || x.highlight.dynamic) && !instance.plotly) {
      var flex = document.createElement("div");
      flex.class = "plotly-crosstalk-control-panel";
      flex.style = "display: flex; flex-wrap: wrap";
      
      // inject the colourpicker HTML container into the flexbox
      if (x.highlight.dynamic) {
        var pickerDiv = document.createElement("div");
        
        var pickerInput = document.createElement("input");
        pickerInput.id = el.id + "-colourpicker";
        pickerInput.placeholder = "asdasd";
        
        var pickerLabel = document.createElement("label");
        pickerLabel.for = pickerInput.id;
        pickerLabel.innerHTML = "Brush color&nbsp;&nbsp;";
        
        pickerDiv.appendChild(pickerLabel);
        pickerDiv.appendChild(pickerInput);
        flex.appendChild(pickerDiv);
      }
      
      // inject selectize HTML containers (one for every crosstalk group)
      if (x.selectize) {
        var ids = Object.keys(x.selectize);
        
        for (var i = 0; i < ids.length; i++) {
          var container = document.createElement("div");
          container.id = ids[i];
          container.style = "width: 80%; height: 10%";
          container.class = "form-group crosstalk-input-plotly-highlight";
          
          var label = document.createElement("label");
          label.for = ids[i];
          label.innerHTML = x.selectize[ids[i]].group;
          label.class = "control-label";
          
          var selectDiv = document.createElement("div");
          var select = document.createElement("select");
          select.multiple = true;
          
          selectDiv.appendChild(select);
          container.appendChild(label);
          container.appendChild(selectDiv);
          flex.appendChild(container);
        }
      }
      
      // finally, insert the flexbox inside the htmlwidget container,
      // but before the plotly graph div
      graphDiv.parentElement.insertBefore(flex, graphDiv);
      
      if (x.highlight.dynamic) {
        var picker = $("#" + pickerInput.id);
        var colors = x.highlight.color || [];
        // TODO: let users specify options?
        var opts = {
          value: colors[0],
          showColour: "both",
          palette: "limited",
          allowedCols: colors.join(" "),
          width: "20%",
          height: "10%"
        };
        picker.colourpicker({changeDelay: 0});
        picker.colourpicker("settings", opts);
        picker.colourpicker("value", opts.value);
        // inform crosstalk about a change in the current selection colour
        var grps = x.highlight.ctGroups || [];
        for (var i = 0; i < grps.length; i++) {
          crosstalk.group(grps[i]).var('plotlySelectionColour')
            .set(picker.colourpicker('value'));
        }
        picker.on("change", function() {
          for (var i = 0; i < grps.length; i++) {
            crosstalk.group(grps[i]).var('plotlySelectionColour')
              .set(picker.colourpicker('value'));
          }
        });
      }
    }
    
    // if no plot exists yet, create one with a particular configuration
    if (!instance.plotly) {
      
      var plot = Plotly.newPlot(graphDiv, x);
      instance.plotly = true;
      
    } else if (x.layout.transition) {
      
      var plot = Plotly.react(graphDiv, x);
    
    } else {
      
      // this is essentially equivalent to Plotly.newPlot(), but avoids creating 
      // a new webgl context
      // https://github.com/plotly/plotly.js/blob/2b24f9def901831e61282076cf3f835598d56f0e/src/plot_api/plot_api.js#L531-L532

      // TODO: restore crosstalk selections?
      Plotly.purge(graphDiv);
      // TODO: why is this necessary to get crosstalk working?
      graphDiv.data = undefined;
      graphDiv.layout = undefined;
      var plot = Plotly.newPlot(graphDiv, x);
    }
    
    // Trigger plotly.js calls defined via `plotlyProxy()`
    plot.then(function() {
      if (HTMLWidgets.shinyMode) {
        Shiny.addCustomMessageHandler("plotly-calls", function(msg) {
          var gd = document.getElementById(msg.id);
          if (!gd) {
            throw new Error("Couldn't find plotly graph with id: " + msg.id);
          }
          // This isn't an official plotly.js method, but it's the only current way to 
          // change just the configuration of a plot 
          // https://community.plot.ly/t/update-config-function/9057
          if (msg.method == "reconfig") {
            Plotly.react(gd, gd.data, gd.layout, msg.args);
            return;
          }
          if (!Plotly[msg.method]) {
            throw new Error("Unknown method " + msg.method);
          }
          var args = [gd].concat(msg.args);
          Plotly[msg.method].apply(null, args);
        });
      }
      
      // plotly's mapbox API doesn't currently support setting bounding boxes
      // https://www.mapbox.com/mapbox-gl-js/example/fitbounds/
      // so we do this manually...
      // TODO: make sure this triggers on a redraw and relayout as well as on initial draw
      var mapboxIDs = graphDiv._fullLayout._subplots.mapbox || [];
      for (var i = 0; i < mapboxIDs.length; i++) {
        var id = mapboxIDs[i];
        var mapOpts = x.layout[id] || {};
        var args = mapOpts._fitBounds || {};
        if (!args) {
          continue;
        }
        var mapObj = graphDiv._fullLayout[id]._subplot.map;
        mapObj.fitBounds(args.bounds, args.options);
      }
      
    });
    
    // Attach attributes (e.g., "key", "z") to plotly event data
    function eventDataWithKey(eventData) {
      if (eventData === undefined || !eventData.hasOwnProperty("points")) {
        return null;
      }
      return eventData.points.map(function(pt) {
        var obj = {
          curveNumber: pt.curveNumber, 
          pointNumber: pt.pointNumber, 
          x: pt.x,
          y: pt.y
        };
        
        // If 'z' is reported with the event data, then use it!
        if (pt.hasOwnProperty("z")) {
          obj.z = pt.z;
        }
        
        if (pt.hasOwnProperty("customdata")) {
          obj.customdata = pt.customdata;
        }
        
        /* 
          TL;DR: (I think) we have to select the graph div (again) to attach keys...
          
          Why? Remember that crosstalk will dynamically add/delete traces 
          (see traceManager.prototype.updateSelection() below)
          For this reason, we can't simply grab keys from x.data (like we did previously)
          Moreover, we can't use _fullData, since that doesn't include 
          unofficial attributes. It's true that click/hover events fire with 
          pt.data, but drag events don't...
        */
        var gd = document.getElementById(el.id);
        var trace = gd.data[pt.curveNumber];
        
        if (!trace._isSimpleKey) {
          var attrsToAttach = ["key"];
        } else {
          // simple keys fire the whole key
          obj.key = trace.key;
          var attrsToAttach = [];
        }
        
        for (var i = 0; i < attrsToAttach.length; i++) {
          var attr = trace[attrsToAttach[i]];
          if (Array.isArray(attr)) {
            if (typeof pt.pointNumber === "number") {
              obj[attrsToAttach[i]] = attr[pt.pointNumber];
            } else if (Array.isArray(pt.pointNumber)) {
              obj[attrsToAttach[i]] = attr[pt.pointNumber[0]][pt.pointNumber[1]];
            } else if (Array.isArray(pt.pointNumbers)) {
              obj[attrsToAttach[i]] = pt.pointNumbers.map(function(idx) { return attr[idx]; });
            }
          }
        }
        return obj;
      });
    }
    
    
    var legendEventData = function(d) {
      // if legendgroup is not relevant just return the trace
      var trace = d.data[d.curveNumber];
      if (!trace.legendgroup) return trace;
      
      // if legendgroup was specified, return all traces that match the group
      var legendgrps = d.data.map(function(trace){ return trace.legendgroup; });
      var traces = [];
      for (i = 0; i < legendgrps.length; i++) {
        if (legendgrps[i] == trace.legendgroup) {
          traces.push(d.data[i]);
        }
      }
      
      return traces;
    };

    
    // send user input event data to shiny
    if (HTMLWidgets.shinyMode && Shiny.setInputValue) {
      
      // Some events clear other input values
      // TODO: always register these?
      var eventClearMap = {
        plotly_deselect: ["plotly_selected", "plotly_selecting", "plotly_brushed", "plotly_brushing", "plotly_click"],
        plotly_unhover: ["plotly_hover"],
        plotly_doubleclick: ["plotly_click"]
      };
    
      Object.keys(eventClearMap).map(function(evt) {
        graphDiv.on(evt, function() {
          var inputsToClear = eventClearMap[evt];
          inputsToClear.map(function(input) {
            Shiny.setInputValue(input + "-" + x.source, null, {priority: "event"});
          });
        });
      });
      
      var eventDataFunctionMap = {
        plotly_click: eventDataWithKey,
        plotly_sunburstclick: eventDataWithKey,
        plotly_hover: eventDataWithKey,
        plotly_unhover: eventDataWithKey,
        // If 'plotly_selected' has already been fired, and you click
        // on the plot afterwards, this event fires `undefined`?!?
        // That might be considered a plotly.js bug, but it doesn't make 
        // sense for this input change to occur if `d` is falsy because,
        // even in the empty selection case, `d` is truthy (an object),
        // and the 'plotly_deselect' event will reset this input
        plotly_selected: function(d) { if (d) { return eventDataWithKey(d); } },
        plotly_selecting: function(d) { if (d) { return eventDataWithKey(d); } },
        plotly_brushed: function(d) {
          if (d) { return d.range ? d.range : d.lassoPoints; }
        },
        plotly_brushing: function(d) {
          if (d) { return d.range ? d.range : d.lassoPoints; }
        },
        plotly_legendclick: legendEventData,
        plotly_legenddoubleclick: legendEventData,
        plotly_clickannotation: function(d) { return d.fullAnnotation }
      };
      
      var registerShinyValue = function(event) {
        var eventDataPreProcessor = eventDataFunctionMap[event] || function(d) { return d ? d : el.id };
        // some events are unique to the R package
        var plotlyJSevent = (event == "plotly_brushed") ? "plotly_selected" : (event == "plotly_brushing") ? "plotly_selecting" : event;
        // register the event
        graphDiv.on(plotlyJSevent, function(d) {
          Shiny.setInputValue(
            event + "-" + x.source,
            JSON.stringify(eventDataPreProcessor(d)),
            {priority: "event"}
          );
        });
      }
    
      var shinyEvents = x.shinyEvents || [];
      shinyEvents.map(registerShinyValue);
    }
    
    // Given an array of {curveNumber: x, pointNumber: y} objects,
    // return a hash of {
    //   set1: {value: [key1, key2, ...], _isSimpleKey: false}, 
    //   set2: {value: [key3, key4, ...], _isSimpleKey: false}
    // }
    function pointsToKeys(points) {
      var keysBySet = {};
      for (var i = 0; i < points.length; i++) {
        
        var trace = graphDiv.data[points[i].curveNumber];
        if (!trace.key || !trace.set) {
          continue;
        }
        
        // set defaults for this keySet
        // note that we don't track the nested property (yet) since we always 
        // emit the union -- http://cpsievert.github.io/talks/20161212b/#21
        keysBySet[trace.set] = keysBySet[trace.set] || {
          value: [],
          _isSimpleKey: trace._isSimpleKey
        };
        
        // Use pointNumber by default, but aggregated traces should emit pointNumbers
        var ptNum = points[i].pointNumber;
        var hasPtNum = typeof ptNum === "number";
        var ptNum = hasPtNum ? ptNum : points[i].pointNumbers;
        
        // selecting a point of a "simple" trace means: select the 
        // entire key attached to this trace, which is useful for,
        // say clicking on a fitted line to select corresponding observations 
        var key = trace._isSimpleKey ? trace.key : Array.isArray(ptNum) ? ptNum.map(function(idx) { return trace.key[idx]; }) : trace.key[ptNum];
        // http://stackoverflow.com/questions/10865025/merge-flatten-an-array-of-arrays-in-javascript
        var keyFlat = trace._isNestedKey ? [].concat.apply([], key) : key;
        
        // TODO: better to only add new values?
        keysBySet[trace.set].value = keysBySet[trace.set].value.concat(keyFlat);
      }
      
      return keysBySet;
    }
    
    
    x.highlight.color = x.highlight.color || [];
    // make sure highlight color is an array
    if (!Array.isArray(x.highlight.color)) {
      x.highlight.color = [x.highlight.color];
    }

    var traceManager = new TraceManager(graphDiv, x.highlight);

    // Gather all *unique* sets.
    var allSets = [];
    for (var curveIdx = 0; curveIdx < x.data.length; curveIdx++) {
      var newSet = x.data[curveIdx].set;
      if (newSet) {
        if (allSets.indexOf(newSet) === -1) {
          allSets.push(newSet);
        }
      }
    }

    // register event listeners for all sets
    for (var i = 0; i < allSets.length; i++) {
      
      var set = allSets[i];
      var selection = new crosstalk.SelectionHandle(set);
      var filter = new crosstalk.FilterHandle(set);
      
      var filterChange = function(e) {
        removeBrush(el);
        traceManager.updateFilter(set, e.value);
      };
      filter.on("change", filterChange);
      
      
      var selectionChange = function(e) {
        
        // Workaround for 'plotly_selected' now firing previously selected
        // points (in addition to new ones) when holding shift key. In our case,
        // we just want the new keys 
        if (x.highlight.on === "plotly_selected" && x.highlight.persistentShift) {
          // https://stackoverflow.com/questions/1187518/how-to-get-the-difference-between-two-arrays-in-javascript
          Array.prototype.diff = function(a) {
              return this.filter(function(i) {return a.indexOf(i) < 0;});
          };
          e.value = e.value.diff(e.oldValue);
        }
        
        // array of "event objects" tracking the selection history
        // this is used to avoid adding redundant selections
        var selectionHistory = crosstalk.var("plotlySelectionHistory").get() || [];
        
        // Construct an event object "defining" the current event. 
        var event = {
          receiverID: traceManager.gd.id,
          plotlySelectionColour: crosstalk.group(set).var("plotlySelectionColour").get()
        };
        event[set] = e.value;
        // TODO: is there a smarter way to check object equality?
        if (selectionHistory.length > 0) {
          var ev = JSON.stringify(event);
          for (var i = 0; i < selectionHistory.length; i++) {
            var sel = JSON.stringify(selectionHistory[i]);
            if (sel == ev) {
              return;
            }
          }
        }
        
        // accumulate history for persistent selection
        if (!x.highlight.persistent) {
          selectionHistory = [event];
        } else {
          selectionHistory.push(event);
        }
        crosstalk.var("plotlySelectionHistory").set(selectionHistory);
        
        // do the actual updating of traces, frames, and the selectize widget
        traceManager.updateSelection(set, e.value);
        // https://github.com/selectize/selectize.js/blob/master/docs/api.md#methods_items
        if (x.selectize) {
          if (!x.highlight.persistent || e.value === null) {
            selectize.clear(true);
          }
          selectize.addItems(e.value, true);
          selectize.close();
        }
      }
      selection.on("change", selectionChange);
      
      // Set a crosstalk variable selection value, triggering an update
      var turnOn = function(e) {
        if (e) {
          var selectedKeys = pointsToKeys(e.points);
          // Keys are group names, values are array of selected keys from group.
          for (var set in selectedKeys) {
            if (selectedKeys.hasOwnProperty(set)) {
              selection.set(selectedKeys[set].value, {sender: el});
            }
          }
        }
      };
      if (x.highlight.debounce > 0) {
        turnOn = debounce(turnOn, x.highlight.debounce);
      }
      graphDiv.on(x.highlight.on, turnOn);
      
      graphDiv.on(x.highlight.off, function turnOff(e) {
        // remove any visual clues
        removeBrush(el);
        // remove any selection history
        crosstalk.var("plotlySelectionHistory").set(null);
        // trigger the actual removal of selection traces
        selection.set(null, {sender: el});
      });
          
      // register a callback for selectize so that there is bi-directional
      // communication between the widget and direct manipulation events
      if (x.selectize) {
        var selectizeID = Object.keys(x.selectize)[i];
        var items = x.selectize[selectizeID].items;
        var first = [{value: "", label: "(All)"}];
        var opts = {
          options: first.concat(items),
          searchField: "label",
          valueField: "value",
          labelField: "label",
          maxItems: 50
        };
        var select = $("#" + selectizeID).find("select")[0];
        var selectize = $(select).selectize(opts)[0].selectize;
        // NOTE: this callback is triggered when *directly* altering 
        // dropdown items
        selectize.on("change", function() {
          var currentItems = traceManager.groupSelections[set] || [];
          if (!x.highlight.persistent) {
            removeBrush(el);
            for (var i = 0; i < currentItems.length; i++) {
              selectize.removeItem(currentItems[i], true);
            }
          }
          var newItems = selectize.items.filter(function(idx) { 
            return currentItems.indexOf(idx) < 0;
          });
          if (newItems.length > 0) {
            traceManager.updateSelection(set, newItems);
          } else {
            // Item has been removed...
            // TODO: this logic won't work for dynamically changing palette 
            traceManager.updateSelection(set, null);
            traceManager.updateSelection(set, selectize.items);
          }
        });
      }
    } // end of selectionChange
    
  } // end of renderValue
}); // end of widget definition

/**
 * @param graphDiv The Plotly graph div
 * @param highlight An object with options for updating selection(s)
 */
function TraceManager(graphDiv, highlight) {
  // The Plotly graph div
  this.gd = graphDiv;

  // Preserve the original data.
  // TODO: try using Lib.extendFlat() as done in  
  // https://github.com/plotly/plotly.js/pull/1136 
  this.origData = JSON.parse(JSON.stringify(graphDiv.data));
  
  // avoid doing this over and over
  this.origOpacity = [];
  for (var i = 0; i < this.origData.length; i++) {
    this.origOpacity[i] = this.origData[i].opacity === 0 ? 0 : (this.origData[i].opacity || 1);
  }

  // key: group name, value: null or array of keys representing the
  // most recently received selection for that group.
  this.groupSelections = {};
  
  // selection parameters (e.g., transient versus persistent selection)
  this.highlight = highlight;
}

TraceManager.prototype.close = function() {
  // TODO: Unhook all event handlers
};

TraceManager.prototype.updateFilter = function(group, keys) {

  if (typeof(keys) === "undefined" || keys === null) {
    
    this.gd.data = JSON.parse(JSON.stringify(this.origData));
    
  } else {
  
    var traces = [];
    for (var i = 0; i < this.origData.length; i++) {
      var trace = this.origData[i];
      if (!trace.key || trace.set !== group) {
        continue;
      }
      var matchFunc = getMatchFunc(trace);
      var matches = matchFunc(trace.key, keys);
      
      if (matches.length > 0) {
        if (!trace._isSimpleKey) {
          // subsetArrayAttrs doesn't mutate trace (it makes a modified clone)
          trace = subsetArrayAttrs(trace, matches);
        }
        traces.push(trace);
      }
    }
    this.gd.data = traces;
  }
  
  Plotly.redraw(this.gd);
  
  // NOTE: we purposely do _not_ restore selection(s), since on filter,
  // axis likely will update, changing the pixel -> data mapping, leading 
  // to a likely mismatch in the brush outline and highlighted marks
  
};

TraceManager.prototype.updateSelection = function(group, keys) {
  
  if (keys !== null && !Array.isArray(keys)) {
    throw new Error("Invalid keys argument; null or array expected");
  }
  
  // if selection has been cleared, or if this is transient
  // selection, delete the "selection traces"
  var nNewTraces = this.gd.data.length - this.origData.length;
  if (keys === null || !this.highlight.persistent && nNewTraces > 0) {
    var tracesToRemove = [];
    for (var i = 0; i < this.gd.data.length; i++) {
      if (this.gd.data[i]._isCrosstalkTrace) tracesToRemove.push(i);
    }
    Plotly.deleteTraces(this.gd, tracesToRemove);
    this.groupSelections[group] = keys;
  } else {
    // add to the groupSelection, rather than overwriting it
    // TODO: can this be removed?
    this.groupSelections[group] = this.groupSelections[group] || [];
    for (var i = 0; i < keys.length; i++) {
      var k = keys[i];
      if (this.groupSelections[group].indexOf(k) < 0) {
        this.groupSelections[group].push(k);
      }
    }
  }
  
  if (keys === null) {
    
    Plotly.restyle(this.gd, {"opacity": this.origOpacity});
    
  } else if (keys.length >= 1) {
    
    // placeholder for new "selection traces"
    var traces = [];
    // this variable is set in R/highlight.R
    var selectionColour = crosstalk.group(group).var("plotlySelectionColour").get() || 
      this.highlight.color[0];

    for (var i = 0; i < this.origData.length; i++) {
      // TODO: try using Lib.extendFlat() as done in  
      // https://github.com/plotly/plotly.js/pull/1136 
      var trace = JSON.parse(JSON.stringify(this.gd.data[i]));
      if (!trace.key || trace.set !== group) {
        continue;
      }
      // Get sorted array of matching indices in trace.key
      var matchFunc = getMatchFunc(trace);
      var matches = matchFunc(trace.key, keys);
      
      if (matches.length > 0) {
        // If this is a "simple" key, that means select the entire trace
        if (!trace._isSimpleKey) {
          trace = subsetArrayAttrs(trace, matches);
        }
        // reach into the full trace object so we can properly reflect the 
        // selection attributes in every view
        var d = this.gd._fullData[i];
        
        /* 
        / Recursively inherit selection attributes from various sources, 
        / in order of preference:
        /  (1) official plotly.js selected attribute
        /  (2) highlight(selected = attrs_selected(...))
        */
        // TODO: it would be neat to have a dropdown to dynamically specify these!
        $.extend(true, trace, this.highlight.selected);
        
        // if it is defined, override color with the "dynamic brush color""
        if (d.marker) {
          trace.marker = trace.marker || {};
          trace.marker.color =  selectionColour || trace.marker.color || d.marker.color;
        }
        if (d.line) {
          trace.line = trace.line || {};
          trace.line.color =  selectionColour || trace.line.color || d.line.color;
        }
        if (d.textfont) {
          trace.textfont = trace.textfont || {};
          trace.textfont.color =  selectionColour || trace.textfont.color || d.textfont.color;
        }
        if (d.fillcolor) {
          // TODO: should selectionColour inherit alpha from the existing fillcolor?
          trace.fillcolor = selectionColour || trace.fillcolor || d.fillcolor;
        }
        // attach a sensible name/legendgroup
        trace.name = trace.name || keys.join("<br />");
        trace.legendgroup = trace.legendgroup || keys.join("<br />");
        
        // keep track of mapping between this new trace and the trace it targets
        // (necessary for updating frames to reflect the selection traces)
        trace._originalIndex = i;
        trace._newIndex = this.gd._fullData.length + traces.length;
        trace._isCrosstalkTrace = true;
        traces.push(trace);
      }
    }
    
    if (traces.length > 0) {
      
      Plotly.addTraces(this.gd, traces).then(function(gd) {
        // incrementally add selection traces to frames
        // (this is heavily inspired by Plotly.Plots.modifyFrames() 
        // in src/plots/plots.js)
        var _hash = gd._transitionData._frameHash;
        var _frames = gd._transitionData._frames || [];
        
        for (var i = 0; i < _frames.length; i++) {
          
          // add to _frames[i].traces *if* this frame references selected trace(s)
          var newIndices = [];
          for (var j = 0; j < traces.length; j++) {
            var tr = traces[j];
            if (_frames[i].traces.indexOf(tr._originalIndex) > -1) {
              newIndices.push(tr._newIndex);
              _frames[i].traces.push(tr._newIndex);
            }
          }
          
          // nothing to do...
          if (newIndices.length === 0) {
            continue;
          }
          
          var ctr = 0;
          var nFrameTraces = _frames[i].data.length;
          
          for (var j = 0; j < nFrameTraces; j++) {
            var frameTrace = _frames[i].data[j];
            if (!frameTrace.key || frameTrace.set !== group) {
              continue;
            }
            
            var matchFunc = getMatchFunc(frameTrace);
            var matches = matchFunc(frameTrace.key, keys);
            
            if (matches.length > 0) {
              if (!trace._isSimpleKey) {
                frameTrace = subsetArrayAttrs(frameTrace, matches);
              }
              var d = gd._fullData[newIndices[ctr]];
              if (d.marker) {
                frameTrace.marker = d.marker;
              }
              if (d.line) {
                frameTrace.line = d.line;
              }
              if (d.textfont) {
                frameTrace.textfont = d.textfont;
              }
              ctr = ctr + 1;
              _frames[i].data.push(frameTrace);
            }
          }
          
          // update gd._transitionData._frameHash
          _hash[_frames[i].name] = _frames[i];
        }
      
      });
      
      // dim traces that have a set matching the set of selection sets
      var tracesToDim = [],
          opacities = [],
          sets = Object.keys(this.groupSelections),
          n = this.origData.length;
          
      for (var i = 0; i < n; i++) {
        var opacity = this.origOpacity[i] || 1;
        // have we already dimmed this trace? Or is this even worth doing?
        if (opacity !== this.gd._fullData[i].opacity || this.highlight.opacityDim === 1) {
          continue;
        }
        // is this set an element of the set of selection sets?
        var matches = findMatches(sets, [this.gd.data[i].set]);
        if (matches.length) {
          tracesToDim.push(i);
          opacities.push(opacity * this.highlight.opacityDim);
        }
      }
      
      if (tracesToDim.length > 0) {
        Plotly.restyle(this.gd, {"opacity": opacities}, tracesToDim);
        // turn off the selected/unselected API
        Plotly.restyle(this.gd, {"selectedpoints": null});
      }
      
    }
    
  }
};

/* 
Note: in all of these match functions, we assume needleSet (i.e. the selected keys)
is a 1D (or flat) array. The real difference is the meaning of haystack.
findMatches() does the usual thing you'd expect for 
linked brushing on a scatterplot matrix. findSimpleMatches() returns a match iff 
haystack is a subset of the needleSet. findNestedMatches() returns 
*/

function getMatchFunc(trace) {
  return (trace._isNestedKey) ? findNestedMatches : 
    (trace._isSimpleKey) ? findSimpleMatches : findMatches;
}

// find matches for "flat" keys
function findMatches(haystack, needleSet) {
  var matches = [];
  haystack.forEach(function(obj, i) {
    if (obj === null || needleSet.indexOf(obj) >= 0) {
      matches.push(i);
    }
  });
  return matches;
}

// find matches for "simple" keys
function findSimpleMatches(haystack, needleSet) {
  var match = haystack.every(function(val) {
    return val === null || needleSet.indexOf(val) >= 0;
  });
  // yes, this doesn't make much sense other than conforming 
  // to the output type of the other match functions
  return (match) ? [0] : []
}

// find matches for a "nested" haystack (2D arrays)
function findNestedMatches(haystack, needleSet) {
  var matches = [];
  for (var i = 0; i < haystack.length; i++) {
    var hay = haystack[i];
    var match = hay.every(function(val) { 
      return val === null || needleSet.indexOf(val) >= 0; 
    });
    if (match) {
      matches.push(i);
    }
  }
  return matches;
}

function isPlainObject(obj) {
  return (
    Object.prototype.toString.call(obj) === '[object Object]' &&
    Object.getPrototypeOf(obj) === Object.prototype
  );
}

function subsetArrayAttrs(obj, indices) {
  var newObj = {};
  Object.keys(obj).forEach(function(k) {
    var val = obj[k];

    if (k.charAt(0) === "_") {
      newObj[k] = val;
    } else if (k === "transforms" && Array.isArray(val)) {
      newObj[k] = val.map(function(transform) {
        return subsetArrayAttrs(transform, indices);
      });
    } else if (k === "colorscale" && Array.isArray(val)) {
      newObj[k] = val;
    } else if (isPlainObject(val)) {
      newObj[k] = subsetArrayAttrs(val, indices);
    } else if (Array.isArray(val)) {
      newObj[k] = subsetArray(val, indices);
    } else {
      newObj[k] = val;
    }
  });
  return newObj;
}

function subsetArray(arr, indices) {
  var result = [];
  for (var i = 0; i < indices.length; i++) {
    result.push(arr[indices[i]]);
  }
  return result;
}

// Convenience function for removing plotly's brush 
function removeBrush(el) {
  var outlines = el.querySelectorAll(".select-outline");
  for (var i = 0; i < outlines.length; i++) {
    outlines[i].remove();
  }
}


// https://davidwalsh.name/javascript-debounce-function

// Returns a function, that, as long as it continues to be invoked, will not
// be triggered. The function will be called after it stops being called for
// N milliseconds. If `immediate` is passed, trigger the function on the
// leading edge, instead of the trailing.
function debounce(func, wait, immediate) {
	var timeout;
	return function() {
		var context = this, args = arguments;
		var later = function() {
			timeout = null;
			if (!immediate) func.apply(context, args);
		};
		var callNow = immediate && !timeout;
		clearTimeout(timeout);
		timeout = setTimeout(later, wait);
		if (callNow) func.apply(context, args);
	};
};
```
