## Supplemental Figure 1 - Interactive version of figure 2A for "Automated assessments can support regional Red Lists but need to be applied with caution: A case study from central Germany": plotly-latest.min.js

```
/**
* plotly.js v2.11.1
* Copyright 2012-2022, Plotly, Inc.
* All rights reserved.
* Licensed under the MIT license
*/
!function(t){if("object"==typeof exports&&"undefined"!=typeof module)module.exports=t();else if("function"==typeof define&&define.amd)define([],t);else{("undefined"!=typeof window?window:"undefined"!=typeof global?global:"undefined"!=typeof self?self:this).Plotly=t()}}((function(){return function t(e,r,n){function i(o,s){if(!r[o]){if(!e[o]){var l="function"==typeof require&&require;if(!s&&l)return l(o,!0);if(a)return a(o,!0);var c=new Error("Cannot find module '"+o+"'");throw c.code="MODULE_NOT_FOUND",c}var u=r[o]={exports:{}};e[o][0].call(u.exports,(function(t){return i(e[o][1][t]||t)}),u,u.exports,t,e,r,n)}return r[o].exports}for(var a="function"==typeof require&&require,o=0;o<n.length;o++)i(n[o]);return i}({1:[function(t,e,r){"use strict";var n=t("../src/lib"),i={"X,X div":'direction:ltr;font-family:"Open Sans",verdana,arial,sans-serif;margin:0;padding:0;',"X input,X button":'font-family:"Open Sans",verdana,arial,sans-serif;',"X input:focus,X button:focus":"outline:none;","X a":"text-decoration:none;","X a:hover":"text-decoration:none;","X .crisp":"shape-rendering:crispEdges;","X .user-select-none":"-webkit-user-select:none;-moz-user-select:none;-ms-user-select:none;-o-user-select:none;user-select:none;","X svg":"overflow:hidden;","X svg a":"fill:#447adb;","X svg a:hover":"fill:#3c6dc5;","X .main-svg":"position:absolute;top:0;left:0;pointer-events:none;","X .main-svg .draglayer":"pointer-events:all;","X .cursor-default":"cursor:default;","X .cursor-pointer":"cursor:pointer;","X .cursor-crosshair":"cursor:crosshair;","X .cursor-move":"cursor:move;","X .cursor-col-resize":"cursor:col-resize;","X .cursor-row-resize":"cursor:row-resize;","X .cursor-ns-resize":"cursor:ns-resize;","X .cursor-ew-resize":"cursor:ew-resize;","X .cursor-sw-resize":"cursor:sw-resize;","X .cursor-s-resize":"cursor:s-resize;","X .cursor-se-resize":"cursor:se-resize;","X .cursor-w-resize":"cursor:w-resize;","X .cursor-e-resize":"cursor:e-resize;","X .cursor-nw-resize":"cursor:nw-resize;","X .cursor-n-resize":"cursor:n-resize;","X .cursor-ne-resize":"cursor:ne-resize;","X .cursor-grab":"cursor:-webkit-grab;cursor:grab;","X .modebar":"position:absolute;top:2px;right:2px;","X .ease-bg":"-webkit-transition:background-color .3s ease 0s;-moz-transition:background-color .3s ease 0s;-ms-transition:background-color .3s ease 0s;-o-transition:background-color .3s ease 0s;transition:background-color .3s ease 0s;","X .modebar--hover>:not(.watermark)":"opacity:0;-webkit-transition:opacity .3s ease 0s;-moz-transition:opacity .3s ease 0s;-ms-transition:opacity .3s ease 0s;-o-transition:opacity .3s ease 0s;transition:opacity .3s ease 0s;","X:hover .modebar--hover .modebar-group":"opacity:1;","X .modebar-group":"float:left;display:inline-block;box-sizing:border-box;padding-left:8px;position:relative;vertical-align:middle;white-space:nowrap;","X .modebar-btn":"position:relative;font-size:16px;padding:3px 4px;height:22px;cursor:pointer;line-height:normal;box-sizing:border-box;","X .modebar-btn svg":"position:relative;top:2px;","X .modebar.vertical":"display:flex;flex-direction:column;flex-wrap:wrap;align-content:flex-end;max-height:100%;","X .modebar.vertical svg":"top:-1px;","X .modebar.vertical .modebar-group":"display:block;float:none;padding-left:0px;padding-bottom:8px;","X .modebar.vertical .modebar-group .modebar-btn":"display:block;text-align:center;","X [data-title]:before,X [data-title]:after":"position:absolute;-webkit-transform:translate3d(0, 0, 0);-moz-transform:translate3d(0, 0, 0);-ms-transform:translate3d(0, 0, 0);-o-transform:translate3d(0, 0, 0);transform:translate3d(0, 0, 0);display:none;opacity:0;z-index:1001;pointer-events:none;top:110%;right:50%;","X [data-title]:hover:before,X [data-title]:hover:after":"display:block;opacity:1;","X [data-title]:before":'content:"";position:absolute;background:transparent;border:6px solid transparent;z-index:1002;margin-top:-12px;border-bottom-color:#69738a;margin-right:-6px;',"X [data-title]:after":"content:attr(data-title);background:#69738a;color:#fff;padding:8px 10px;font-size:12px;line-height:12px;white-space:nowrap;margin-right:-18px;border-radius:2px;","X .vertical [data-title]:before,X .vertical [data-title]:after":"top:0%;right:200%;","X .vertical [data-title]:before":"border:6px solid transparent;border-left-color:#69738a;margin-top:8px;margin-right:-30px;","X .select-outline":"fill:none;stroke-width:1;shape-rendering:crispEdges;","X .select-outline-1":"stroke:#fff;","X .select-outline-2":"stroke:#000;stroke-dasharray:2px 2px;",Y:'font-family:"Open Sans",verdana,arial,sans-serif;position:fixed;top:50px;right:20px;z-index:10000;font-size:10pt;max-width:180px;',"Y p":"margin:0;","Y .notifier-note":"min-width:180px;max-width:250px;border:1px solid #fff;z-index:3000;margin:0;background-color:#8c97af;background-color:rgba(140,151,175,.9);color:#fff;padding:10px;overflow-wrap:break-word;word-wrap:break-word;-ms-hyphens:auto;-webkit-hyphens:auto;hyphens:auto;","Y .notifier-close":"color:#fff;opacity:.8;float:right;padding:0 5px;background:none;border:none;font-size:20px;font-weight:bold;line-height:20px;","Y .notifier-close:hover":"color:#444;text-decoration:none;cursor:pointer;"};for(var a in i){var o=a.replace(/^,/," ,").replace(/X/g,".js-plotly-plot .plotly").replace(/Y/g,".plotly-notifier");n.addStyleRule(o,i[a])}},{"../src/lib":498}],2:[function(t,e,r){"use strict";e.exports=t("../src/transforms/aggregate")},{"../src/transforms/aggregate":1113}],3:[function(t,e,r){"use strict";e.exports=t("../src/traces/bar")},{"../src/traces/bar":651}],4:[function(t,e,r){"use strict";e.exports=t("../src/traces/barpolar")},{"../src/traces/barpolar":664}],5:[function(t,e,r){"use strict";e.exports=t("../src/traces/box")},{"../src/traces/box":674}],6:[function(t,e,r){"use strict";e.exports=t("../src/components/calendars")},{"../src/components/calendars":359}],7:[function(t,e,r){"use strict";e.exports=t("../src/traces/candlestick")},{"../src/traces/candlestick":683}],8:[function(t,e,r){"use strict";e.exports=t("../src/traces/carpet")},{"../src/traces/carpet":702}],9:[function(t,e,r){"use strict";e.exports=t("../src/traces/choropleth")},{"../src/traces/choropleth":716}],10:[function(t,e,r){"use strict";e.exports=t("../src/traces/choroplethmapbox")},{"../src/traces/choroplethmapbox":723}],11:[function(t,e,r){"use strict";e.exports=t("../src/traces/cone")},{"../src/traces/cone":729}],12:[function(t,e,r){"use strict";e.exports=t("../src/traces/contour")},{"../src/traces/contour":744}],13:[function(t,e,r){"use strict";e.exports=t("../src/traces/contourcarpet")},{"../src/traces/contourcarpet":755}],14:[function(t,e,r){"use strict";e.exports=t("../src/core")},{"../src/core":476}],15:[function(t,e,r){"use strict";e.exports=t("../src/traces/densitymapbox")},{"../src/traces/densitymapbox":763}],16:[function(t,e,r){"use strict";e.exports=t("../src/transforms/filter")},{"../src/transforms/filter":1114}],17:[function(t,e,r){"use strict";e.exports=t("../src/traces/funnel")},{"../src/traces/funnel":773}],18:[function(t,e,r){"use strict";e.exports=t("../src/traces/funnelarea")},{"../src/traces/funnelarea":782}],19:[function(t,e,r){"use strict";e.exports=t("../src/transforms/groupby")},{"../src/transforms/groupby":1115}],20:[function(t,e,r){"use strict";e.exports=t("../src/traces/heatmap")},{"../src/traces/heatmap":795}],21:[function(t,e,r){"use strict";e.exports=t("../src/traces/heatmapgl")},{"../src/traces/heatmapgl":806}],22:[function(t,e,r){"use strict";e.exports=t("../src/traces/histogram")},{"../src/traces/histogram":818}],23:[function(t,e,r){"use strict";e.exports=t("../src/traces/histogram2d")},{"../src/traces/histogram2d":824}],24:[function(t,e,r){"use strict";e.exports=t("../src/traces/histogram2dcontour")},{"../src/traces/histogram2dcontour":828}],25:[function(t,e,r){"use strict";e.exports=t("../src/traces/icicle")},{"../src/traces/icicle":834}],26:[function(t,e,r){"use strict";e.exports=t("../src/traces/image")},{"../src/traces/image":847}],27:[function(t,e,r){"use strict";var n=t("./core");n.register([t("./bar"),t("./box"),t("./heatmap"),t("./histogram"),t("./histogram2d"),t("./histogram2dcontour"),t("./contour"),t("./scatterternary"),t("./violin"),t("./funnel"),t("./waterfall"),t("./image"),t("./pie"),t("./sunburst"),t("./treemap"),t("./icicle"),t("./funnelarea"),t("./scatter3d"),t("./surface"),t("./isosurface"),t("./volume"),t("./mesh3d"),t("./cone"),t("./streamtube"),t("./scattergeo"),t("./choropleth"),t("./scattergl"),t("./splom"),t("./pointcloud"),t("./heatmapgl"),t("./parcoords"),t("./parcats"),t("./scattermapbox"),t("./choroplethmapbox"),t("./densitymapbox"),t("./sankey"),t("./indicator"),t("./table"),t("./carpet"),t("./scattercarpet"),t("./contourcarpet"),t("./ohlc"),t("./candlestick"),t("./scatterpolar"),t("./scatterpolargl"),t("./barpolar"),t("./scattersmith"),t("./aggregate"),t("./filter"),t("./groupby"),t("./sort"),t("./calendars")]),e.exports=n},{"./aggregate":2,"./bar":3,"./barpolar":4,"./box":5,"./calendars":6,"./candlestick":7,"./carpet":8,"./choropleth":9,"./choroplethmapbox":10,"./cone":11,"./contour":12,"./contourcarpet":13,"./core":14,"./densitymapbox":15,"./filter":16,"./funnel":17,"./funnelarea":18,"./groupby":19,"./heatmap":20,"./heatmapgl":21,"./histogram":22,"./histogram2d":23,"./histogram2dcontour":24,"./icicle":25,"./image":26,"./indicator":28,"./isosurface":29,"./mesh3d":30,"./ohlc":31,"./parcats":32,"./parcoords":33,"./pie":34,"./pointcloud":35,"./sankey":36,"./scatter3d":37,"./scattercarpet":38,"./scattergeo":39,"./scattergl":40,"./scattermapbox":41,"./scatterpolar":42,"./scatterpolargl":43,"./scattersmith":44,"./scatterternary":45,"./sort":46,"./splom":47,"./streamtube":48,"./sunburst":49,"./surface":50,"./table":51,"./treemap":52,"./violin":53,"./volume":54,"./waterfall":55}],28:[function(t,e,r){"use strict";e.exports=t("../src/traces/indicator")},{"../src/traces/indicator":855}],29:[function(t,e,r){"use strict";e.exports=t("../src/traces/isosurface")},{"../src/traces/isosurface":861}],30:[function(t,e,r){"use strict";e.exports=t("../src/traces/mesh3d")},{"../src/traces/mesh3d":866}],31:[function(t,e,r){"use strict";e.exports=t("../src/traces/ohlc")},{"../src/traces/ohlc":871}],32:[function(t,e,r){"use strict";e.exports=t("../src/traces/parcats")},{"../src/traces/parcats":880}],33:[function(t,e,r){"use strict";e.exports=t("../src/traces/parcoords")},{"../src/traces/parcoords":891}],34:[function(t,e,r){"use strict";e.exports=t("../src/traces/pie")},{"../src/traces/pie":902}],35:[function(t,e,r){"use strict";e.exports=t("../src/traces/pointcloud")},{"../src/traces/pointcloud":911}],36:[function(t,e,r){"use strict";e.exports=t("../src/traces/sankey")},{"../src/traces/sankey":917}],37:[function(t,e,r){"use strict";e.exports=t("../src/traces/scatter3d")},{"../src/traces/scatter3d":955}],38:[function(t,e,r){"use strict";e.exports=t("../src/traces/scattercarpet")},{"../src/traces/scattercarpet":962}],39:[function(t,e,r){"use strict";e.exports=t("../src/traces/scattergeo")},{"../src/traces/scattergeo":970}],40:[function(t,e,r){"use strict";e.exports=t("../src/traces/scattergl")},{"../src/traces/scattergl":984}],41:[function(t,e,r){"use strict";e.exports=t("../src/traces/scattermapbox")},{"../src/traces/scattermapbox":994}],42:[function(t,e,r){"use strict";e.exports=t("../src/traces/scatterpolar")},{"../src/traces/scatterpolar":1002}],43:[function(t,e,r){"use strict";e.exports=t("../src/traces/scatterpolargl")},{"../src/traces/scatterpolargl":1010}],44:[function(t,e,r){"use strict";e.exports=t("../src/traces/scattersmith")},{"../src/traces/scattersmith":1017}],45:[function(t,e,r){"use strict";e.exports=t("../src/traces/scatterternary")},{"../src/traces/scatterternary":1025}],46:[function(t,e,r){"use strict";e.exports=t("../src/transforms/sort")},{"../src/transforms/sort":1117}],47:[function(t,e,r){"use strict";e.exports=t("../src/traces/splom")},{"../src/traces/splom":1035}],48:[function(t,e,r){"use strict";e.exports=t("../src/traces/streamtube")},{"../src/traces/streamtube":1043}],49:[function(t,e,r){"use strict";e.exports=t("../src/traces/sunburst")},{"../src/traces/sunburst":1051}],50:[function(t,e,r){"use strict";e.exports=t("../src/traces/surface")},{"../src/traces/surface":1060}],51:[function(t,e,r){"use strict";e.exports=t("../src/traces/table")},{"../src/traces/table":1068}],52:[function(t,e,r){"use strict";e.exports=t("../src/traces/treemap")},{"../src/traces/treemap":1079}],53:[function(t,e,r){"use strict";e.exports=t("../src/traces/violin")},{"../src/traces/violin":1092}],54:[function(t,e,r){"use strict";e.exports=t("../src/traces/volume")},{"../src/traces/volume":1100}],55:[function(t,e,r){"use strict";e.exports=t("../src/traces/waterfall")},{"../src/traces/waterfall":1108}],56:[function(t,e,r){!function(n,i){"object"==typeof r&&void 0!==e?i(r,t("d3-array"),t("d3-collection"),t("d3-shape"),t("elementary-circuits-directed-graph")):i(n.d3=n.d3||{},n.d3,n.d3,n.d3,null)}(this,(function(t,e,r,n,i){"use strict";function a(t){return t.target.depth}function o(t,e){return t.sourceLinks.length?t.depth:e-1}function s(t){return function(){return t}}i=i&&i.hasOwnProperty("default")?i.default:i;var l="function"==typeof Symbol&&"symbol"==typeof Symbol.iterator?function(t){return typeof t}:function(t){return t&&"function"==typeof Symbol&&t.constructor===Symbol&&t!==Symbol.prototype?"symbol":typeof t};function c(t,e){return f(t.source,e.source)||t.index-e.index}function u(t,e){return f(t.target,e.target)||t.index-e.index}function f(t,e){return t.partOfCycle===e.partOfCycle?t.y0-e.y0:"top"===t.circularLinkType||"bottom"===e.circularLinkType?-1:1}function h(t){return t.value}function p(t){return(t.y0+t.y1)/2}function d(t){return p(t.source)}function m(t){return p(t.target)}function g(t){return t.index}function v(t){return t.nodes}function y(t){return t.links}function x(t,e){var r=t.get(e);if(!r)throw new Error("missing: "+e);return r}function b(t,e){return e(t)}function _(t,e,r){var n=0;if(null===r){for(var a=[],o=0;o<t.links.length;o++){var s=t.links[o],l=s.source.index,c=s.target.index;a[l]||(a[l]=[]),a[c]||(a[c]=[]),-1===a[l].indexOf(c)&&a[l].push(c)}var u=i(a);u.sort((function(t,e){return t.length-e.length}));var f={};for(o=0;o<u.length;o++){var h=u[o].slice(-2);f[h[0]]||(f[h[0]]={}),f[h[0]][h[1]]=!0}t.links.forEach((function(t){var e=t.target.index,r=t.source.index;e===r||f[r]&&f[r][e]?(t.circular=!0,t.circularLinkID=n,n+=1):t.circular=!1}))}else t.links.forEach((function(t){t.source[r]<t.target[r]?t.circular=!1:(t.circular=!0,t.circularLinkID=n,n+=1)}))}function w(t,e){var r=0,n=0;t.links.forEach((function(i){i.circular&&(i.source.circularLinkType||i.target.circularLinkType?i.circularLinkType=i.source.circularLinkType?i.source.circularLinkType:i.target.circularLinkType:i.circularLinkType=r<n?"top":"bottom","top"==i.circularLinkType?r+=1:n+=1,t.nodes.forEach((function(t){b(t,e)!=b(i.source,e)&&b(t,e)!=b(i.target,e)||(t.circularLinkType=i.circularLinkType)})))})),t.links.forEach((function(t){t.circular&&(t.source.circularLinkType==t.target.circularLinkType&&(t.circularLinkType=t.source.circularLinkType),q(t,e)&&(t.circularLinkType=t.source.circularLinkType))}))}function T(t){var e=Math.abs(t.y1-t.y0),r=Math.abs(t.target.x0-t.source.x1);return Math.atan(r/e)}function k(t,e){var r=0;t.sourceLinks.forEach((function(t){r=t.circular&&!q(t,e)?r+1:r}));var n=0;return t.targetLinks.forEach((function(t){n=t.circular&&!q(t,e)?n+1:n})),r+n}function A(t){var e=t.source.sourceLinks,r=0;e.forEach((function(t){r=t.circular?r+1:r}));var n=t.target.targetLinks,i=0;return n.forEach((function(t){i=t.circular?i+1:i})),!(r>1||i>1)}function M(t,e,r){return t.sort(E),t.forEach((function(n,i){var a,o,s=0;if(q(n,r)&&A(n))n.circularPathData.verticalBuffer=s+n.width/2;else{for(var l=0;l<i;l++)if(a=t[i],o=t[l],!(a.source.column<o.target.column||a.target.column>o.source.column)){var c=t[l].circularPathData.verticalBuffer+t[l].width/2+e;s=c>s?c:s}n.circularPathData.verticalBuffer=s+n.width/2}})),t}function S(t,r,i,a){var o=e.min(t.links,(function(t){return t.source.y0}));t.links.forEach((function(t){t.circular&&(t.circularPathData={})})),M(t.links.filter((function(t){return"top"==t.circularLinkType})),r,a),M(t.links.filter((function(t){return"bottom"==t.circularLinkType})),r,a),t.links.forEach((function(e){if(e.circular){if(e.circularPathData.arcRadius=e.width+10,e.circularPathData.leftNodeBuffer=5,e.circularPathData.rightNodeBuffer=5,e.circularPathData.sourceWidth=e.source.x1-e.source.x0,e.circularPathData.sourceX=e.source.x0+e.circularPathData.sourceWidth,e.circularPathData.targetX=e.target.x0,e.circularPathData.sourceY=e.y0,e.circularPathData.targetY=e.y1,q(e,a)&&A(e))e.circularPathData.leftSmallArcRadius=10+e.width/2,e.circularPathData.leftLargeArcRadius=10+e.width/2,e.circularPathData.rightSmallArcRadius=10+e.width/2,e.circularPathData.rightLargeArcRadius=10+e.width/2,"bottom"==e.circularLinkType?(e.circularPathData.verticalFullExtent=e.source.y1+25+e.circularPathData.verticalBuffer,e.circularPathData.verticalLeftInnerExtent=e.circularPathData.verticalFullExtent-e.circularPathData.leftLargeArcRadius,e.circularPathData.verticalRightInnerExtent=e.circularPathData.verticalFullExtent-e.circularPathData.rightLargeArcRadius):(e.circularPathData.verticalFullExtent=e.source.y0-25-e.circularPathData.verticalBuffer,e.circularPathData.verticalLeftInnerExtent=e.circularPathData.verticalFullExtent+e.circularPathData.leftLargeArcRadius,e.circularPathData.verticalRightInnerExtent=e.circularPathData.verticalFullExtent+e.circularPathData.rightLargeArcRadius);else{var s=e.source.column,l=e.circularLinkType,c=t.links.filter((function(t){return t.source.column==s&&t.circularLinkType==l}));"bottom"==e.circularLinkType?c.sort(C):c.sort(L);var u=0;c.forEach((function(t,n){t.circularLinkID==e.circularLinkID&&(e.circularPathData.leftSmallArcRadius=10+e.width/2+u,e.circularPathData.leftLargeArcRadius=10+e.width/2+n*r+u),u+=t.width})),s=e.target.column,c=t.links.filter((function(t){return t.target.column==s&&t.circularLinkType==l})),"bottom"==e.circularLinkType?c.sort(I):c.sort(P),u=0,c.forEach((function(t,n){t.circularLinkID==e.circularLinkID&&(e.circularPathData.rightSmallArcRadius=10+e.width/2+u,e.circularPathData.rightLargeArcRadius=10+e.width/2+n*r+u),u+=t.width})),"bottom"==e.circularLinkType?(e.circularPathData.verticalFullExtent=Math.max(i,e.source.y1,e.target.y1)+25+e.circularPathData.verticalBuffer,e.circularPathData.verticalLeftInnerExtent=e.circularPathData.verticalFullExtent-e.circularPathData.leftLargeArcRadius,e.circularPathData.verticalRightInnerExtent=e.circularPathData.verticalFullExtent-e.circularPathData.rightLargeArcRadius):(e.circularPathData.verticalFullExtent=o-25-e.circularPathData.verticalBuffer,e.circularPathData.verticalLeftInnerExtent=e.circularPathData.verticalFullExtent+e.circularPathData.leftLargeArcRadius,e.circularPathData.verticalRightInnerExtent=e.circularPathData.verticalFullExtent+e.circularPathData.rightLargeArcRadius)}e.circularPathData.leftInnerExtent=e.circularPathData.sourceX+e.circularPathData.leftNodeBuffer,e.circularPathData.rightInnerExtent=e.circularPathData.targetX-e.circularPathData.rightNodeBuffer,e.circularPathData.leftFullExtent=e.circularPathData.sourceX+e.circularPathData.leftLargeArcRadius+e.circularPathData.leftNodeBuffer,e.circularPathData.rightFullExtent=e.circularPathData.targetX-e.circularPathData.rightLargeArcRadius-e.circularPathData.rightNodeBuffer}if(e.circular)e.path=function(t){var e="";e="top"==t.circularLinkType?"M"+t.circularPathData.sourceX+" "+t.circularPathData.sourceY+" L"+t.circularPathData.leftInnerExtent+" "+t.circularPathData.sourceY+" A"+t.circularPathData.leftLargeArcRadius+" "+t.circularPathData.leftSmallArcRadius+" 0 0 0 "+t.circularPathData.leftFullExtent+" "+(t.circularPathData.sourceY-t.circularPathData.leftSmallArcRadius)+" L"+t.circularPathData.leftFullExtent+" "+t.circularPathData.verticalLeftInnerExtent+" A"+t.circularPathData.leftLargeArcRadius+" "+t.circularPathData.leftLargeArcRadius+" 0 0 0 "+t.circularPathData.leftInnerExtent+" "+t.circularPathData.verticalFullExtent+" L"+t.circularPathData.rightInnerExtent+" "+t.circularPathData.verticalFullExtent+" A"+t.circularPathData.rightLargeArcRadius+" "+t.circularPathData.rightLargeArcRadius+" 0 0 0 "+t.circularPathData.rightFullExtent+" "+t.circularPathData.verticalRightInnerExtent+" L"+t.circularPathData.rightFullExtent+" "+(t.circularPathData.targetY-t.circularPathData.rightSmallArcRadius)+" A"+t.circularPathData.rightLargeArcRadius+" "+t.circularPathData.rightSmallArcRadius+" 0 0 0 "+t.circularPathData.rightInnerExtent+" "+t.circularPathData.targetY+" L"+t.circularPathData.targetX+" "+t.circularPathData.targetY:"M"+t.circularPathData.sourceX+" "+t.circularPathData.sourceY+" L"+t.circularPathData.leftInnerExtent+" "+t.circularPathData.sourceY+" A"+t.circularPathData.leftLargeArcRadius+" "+t.circularPathData.leftSmallArcRadius+" 0 0 1 "+t.circularPathData.leftFullExtent+" "+(t.circularPathData.sourceY+t.circularPathData.leftSmallArcRadius)+" L"+t.circularPathData.leftFullExtent+" "+t.circularPathData.verticalLeftInnerExtent+" A"+t.circularPathData.leftLargeArcRadius+" "+t.circularPathData.leftLargeArcRadius+" 0 0 1 "+t.circularPathData.leftInnerExtent+" "+t.circularPathData.verticalFullExtent+" L"+t.circularPathData.rightInnerExtent+" "+t.circularPathData.verticalFullExtent+" A"+t.circularPathData.rightLargeArcRadius+" "+t.circularPathData.rightLargeArcRadius+" 0 0 1 "+t.circularPathData.rightFullExtent+" "+t.circularPathData.verticalRightInnerExtent+" L"+t.circularPathData.rightFullExtent+" "+(t.circularPathData.targetY+t.circularPathData.rightSmallArcRadius)+" A"+t.circularPathData.rightLargeArcRadius+" "+t.circularPathData.rightSmallArcRadius+" 0 0 1 "+t.circularPathData.rightInnerExtent+" "+t.circularPathData.targetY+" L"+t.circularPathData.targetX+" "+t.circularPathData.targetY;return e}(e);else{var f=n.linkHorizontal().source((function(t){return[t.source.x0+(t.source.x1-t.source.x0),t.y0]})).target((function(t){return[t.target.x0,t.y1]}));e.path=f(e)}}))}function E(t,e){return O(t)==O(e)?"bottom"==t.circularLinkType?C(t,e):L(t,e):O(e)-O(t)}function L(t,e){return t.y0-e.y0}function C(t,e){return e.y0-t.y0}function P(t,e){return t.y1-e.y1}function I(t,e){return e.y1-t.y1}function O(t){return t.target.column-t.source.column}function z(t){return t.target.x0-t.source.x1}function D(t,e){var r=T(t),n=z(e)/Math.tan(r);return"up"==H(t)?t.y1+n:t.y1-n}function R(t,e){var r=T(t),n=z(e)/Math.tan(r);return"up"==H(t)?t.y1-n:t.y1+n}function F(t,e,r,n){t.links.forEach((function(i){if(!i.circular&&i.target.column-i.source.column>1){var a=i.source.column+1,o=i.target.column-1,s=1,l=o-a+1;for(s=1;a<=o;a++,s++)t.nodes.forEach((function(o){if(o.column==a){var c,u=s/(l+1),f=Math.pow(1-u,3),h=3*u*Math.pow(1-u,2),p=3*Math.pow(u,2)*(1-u),d=Math.pow(u,3),m=f*i.y0+h*i.y0+p*i.y1+d*i.y1,g=m-i.width/2,v=m+i.width/2;g>o.y0&&g<o.y1?(c=o.y1-g+10,c="bottom"==o.circularLinkType?c:-c,o=N(o,c,e,r),t.nodes.forEach((function(t){b(t,n)!=b(o,n)&&t.column==o.column&&B(o,t)&&N(t,c,e,r)}))):(v>o.y0&&v<o.y1||g<o.y0&&v>o.y1)&&(c=v-o.y0+10,o=N(o,c,e,r),t.nodes.forEach((function(t){b(t,n)!=b(o,n)&&t.column==o.column&&t.y0<o.y1&&t.y1>o.y1&&N(t,c,e,r)})))}}))}}))}function B(t,e){return t.y0>e.y0&&t.y0<e.y1||(t.y1>e.y0&&t.y1<e.y1||t.y0<e.y0&&t.y1>e.y1)}function N(t,e,r,n){return t.y0+e>=r&&t.y1+e<=n&&(t.y0=t.y0+e,t.y1=t.y1+e,t.targetLinks.forEach((function(t){t.y1=t.y1+e})),t.sourceLinks.forEach((function(t){t.y0=t.y0+e}))),t}function j(t,e,r,n){t.nodes.forEach((function(i){n&&i.y+(i.y1-i.y0)>e&&(i.y=i.y-(i.y+(i.y1-i.y0)-e));var a=t.links.filter((function(t){return b(t.source,r)==b(i,r)})),o=a.length;o>1&&a.sort((function(t,e){if(!t.circular&&!e.circular){if(t.target.column==e.target.column)return t.y1-e.y1;if(!V(t,e))return t.y1-e.y1;if(t.target.column>e.target.column){var r=R(e,t);return t.y1-r}if(e.target.column>t.target.column)return R(t,e)-e.y1}return t.circular&&!e.circular?"top"==t.circularLinkType?-1:1:e.circular&&!t.circular?"top"==e.circularLinkType?1:-1:t.circular&&e.circular?t.circularLinkType===e.circularLinkType&&"top"==t.circularLinkType?t.target.column===e.target.column?t.target.y1-e.target.y1:e.target.column-t.target.column:t.circularLinkType===e.circularLinkType&&"bottom"==t.circularLinkType?t.target.column===e.target.column?e.target.y1-t.target.y1:t.target.column-e.target.column:"top"==t.circularLinkType?-1:1:void 0}));var s=i.y0;a.forEach((function(t){t.y0=s+t.width/2,s+=t.width})),a.forEach((function(t,e){if("bottom"==t.circularLinkType){for(var r=e+1,n=0;r<o;r++)n+=a[r].width;t.y0=i.y1-n-t.width/2}}))}))}function U(t,e,r){t.nodes.forEach((function(e){var n=t.links.filter((function(t){return b(t.target,r)==b(e,r)})),i=n.length;i>1&&n.sort((function(t,e){if(!t.circular&&!e.circular){if(t.source.column==e.source.column)return t.y0-e.y0;if(!V(t,e))return t.y0-e.y0;if(e.source.column<t.source.column){var r=D(e,t);return t.y0-r}if(t.source.column<e.source.column)return D(t,e)-e.y0}return t.circular&&!e.circular?"top"==t.circularLinkType?-1:1:e.circular&&!t.circular?"top"==e.circularLinkType?1:-1:t.circular&&e.circular?t.circularLinkType===e.circularLinkType&&"top"==t.circularLinkType?t.source.column===e.source.column?t.source.y1-e.source.y1:t.source.column-e.source.column:t.circularLinkType===e.circularLinkType&&"bottom"==t.circularLinkType?t.source.column===e.source.column?t.source.y1-e.source.y1:e.source.column-t.source.column:"top"==t.circularLinkType?-1:1:void 0}));var a=e.y0;n.forEach((function(t){t.y1=a+t.width/2,a+=t.width})),n.forEach((function(t,r){if("bottom"==t.circularLinkType){for(var a=r+1,o=0;a<i;a++)o+=n[a].width;t.y1=e.y1-o-t.width/2}}))}))}function V(t,e){return H(t)==H(e)}function H(t){return t.y0-t.y1>0?"up":"down"}function q(t,e){return b(t.source,e)==b(t.target,e)}function G(t,r,n){var i=t.nodes,a=t.links,o=!1,s=!1;if(a.forEach((function(t){"top"==t.circularLinkType?o=!0:"bottom"==t.circularLinkType&&(s=!0)})),0==o||0==s){var l=e.min(i,(function(t){return t.y0})),c=(n-r)/(e.max(i,(function(t){return t.y1}))-l);i.forEach((function(t){var e=(t.y1-t.y0)*c;t.y0=(t.y0-l)*c,t.y1=t.y0+e})),a.forEach((function(t){t.y0=(t.y0-l)*c,t.y1=(t.y1-l)*c,t.width=t.width*c}))}}t.sankeyCircular=function(){var t,n,i=0,a=0,b=1,T=1,A=24,M=g,E=o,L=v,C=y,P=32,I=2,O=null;function z(){var t={nodes:L.apply(null,arguments),links:C.apply(null,arguments)};D(t),_(t,M,O),R(t),B(t),w(t,M),N(t,P,M),V(t);for(var e=4,r=0;r<e;r++)j(t,T,M),U(t,T,M),F(t,a,T,M),j(t,T,M),U(t,T,M);return G(t,a,T),S(t,I,T,M),t}function D(t){t.nodes.forEach((function(t,e){t.index=e,t.sourceLinks=[],t.targetLinks=[]}));var e=r.map(t.nodes,M);return t.links.forEach((function(t,r){t.index=r;var n=t.source,i=t.target;"object"!==(void 0===n?"undefined":l(n))&&(n=t.source=x(e,n)),"object"!==(void 0===i?"undefined":l(i))&&(i=t.target=x(e,i)),n.sourceLinks.push(t),i.targetLinks.push(t)})),t}function R(t){t.nodes.forEach((function(t){t.partOfCycle=!1,t.value=Math.max(e.sum(t.sourceLinks,h),e.sum(t.targetLinks,h)),t.sourceLinks.forEach((function(e){e.circular&&(t.partOfCycle=!0,t.circularLinkType=e.circularLinkType)})),t.targetLinks.forEach((function(e){e.circular&&(t.partOfCycle=!0,t.circularLinkType=e.circularLinkType)}))}))}function B(t){var e,r,n;for(e=t.nodes,r=[],n=0;e.length;++n,e=r,r=[])e.forEach((function(t){t.depth=n,t.sourceLinks.forEach((function(t){r.indexOf(t.target)<0&&!t.circular&&r.push(t.target)}))}));for(e=t.nodes,r=[],n=0;e.length;++n,e=r,r=[])e.forEach((function(t){t.height=n,t.targetLinks.forEach((function(t){r.indexOf(t.source)<0&&!t.circular&&r.push(t.source)}))}));t.nodes.forEach((function(t){t.column=Math.floor(E.call(null,t,n))}))}function N(o,s,l){var c=r.nest().key((function(t){return t.column})).sortKeys(e.ascending).entries(o.nodes).map((function(t){return t.values}));!function(r){if(n){var s=1/0;c.forEach((function(t){var e=T*n/(t.length+1);s=e<s?e:s})),t=s}var l=e.min(c,(function(r){return(T-a-(r.length-1)*t)/e.sum(r,h)}));l*=.3,o.links.forEach((function(t){t.width=t.value*l}));var u=function(t){var r=0,n=0,i=0,a=0,o=e.max(t.nodes,(function(t){return t.column}));return t.links.forEach((function(t){t.circular&&("top"==t.circularLinkType?r+=t.width:n+=t.width,0==t.target.column&&(a+=t.width),t.source.column==o&&(i+=t.width))})),{top:r=r>0?r+25+10:r,bottom:n=n>0?n+25+10:n,left:a=a>0?a+25+10:a,right:i=i>0?i+25+10:i}}(o),f=function(t,r){var n=e.max(t.nodes,(function(t){return t.column})),o=b-i,s=T-a,l=o/(o+r.right+r.left),c=s/(s+r.top+r.bottom);return i=i*l+r.left,b=0==r.right?b:b*l,a=a*c+r.top,T*=c,t.nodes.forEach((function(t){t.x0=i+t.column*((b-i-A)/n),t.x1=t.x0+A})),c}(o,u);l*=f,o.links.forEach((function(t){t.width=t.value*l})),c.forEach((function(t){var e=t.length;t.forEach((function(t,n){t.depth==c.length-1&&1==e||0==t.depth&&1==e?(t.y0=T/2-t.value*l,t.y1=t.y0+t.value*l):t.partOfCycle?0==k(t,r)?(t.y0=T/2+n,t.y1=t.y0+t.value*l):"top"==t.circularLinkType?(t.y0=a+n,t.y1=t.y0+t.value*l):(t.y0=T-t.value*l-n,t.y1=t.y0+t.value*l):0==u.top||0==u.bottom?(t.y0=(T-a)/e*n,t.y1=t.y0+t.value*l):(t.y0=(T-a)/2-e/2+n,t.y1=t.y0+t.value*l)}))}))}(l),y();for(var u=1,g=s;g>0;--g)v(u*=.99,l),y();function v(t,r){var n=c.length;c.forEach((function(i){var a=i.length,o=i[0].depth;i.forEach((function(i){var s;if(i.sourceLinks.length||i.targetLinks.length)if(i.partOfCycle&&k(i,r)>0);else if(0==o&&1==a)s=i.y1-i.y0,i.y0=T/2-s/2,i.y1=T/2+s/2;else if(o==n-1&&1==a)s=i.y1-i.y0,i.y0=T/2-s/2,i.y1=T/2+s/2;else{var l=e.mean(i.sourceLinks,m),c=e.mean(i.targetLinks,d),u=((l&&c?(l+c)/2:l||c)-p(i))*t;i.y0+=u,i.y1+=u}}))}))}function y(){c.forEach((function(e){var r,n,i,o=a,s=e.length;for(e.sort(f),i=0;i<s;++i)(n=o-(r=e[i]).y0)>0&&(r.y0+=n,r.y1+=n),o=r.y1+t;if((n=o-t-T)>0)for(o=r.y0-=n,r.y1-=n,i=s-2;i>=0;--i)(n=(r=e[i]).y1+t-o)>0&&(r.y0-=n,r.y1-=n),o=r.y0}))}}function V(t){t.nodes.forEach((function(t){t.sourceLinks.sort(u),t.targetLinks.sort(c)})),t.nodes.forEach((function(t){var e=t.y0,r=e,n=t.y1,i=n;t.sourceLinks.forEach((function(t){t.circular?(t.y0=n-t.width/2,n-=t.width):(t.y0=e+t.width/2,e+=t.width)})),t.targetLinks.forEach((function(t){t.circular?(t.y1=i-t.width/2,i-=t.width):(t.y1=r+t.width/2,r+=t.width)}))}))}return z.nodeId=function(t){return arguments.length?(M="function"==typeof t?t:s(t),z):M},z.nodeAlign=function(t){return arguments.length?(E="function"==typeof t?t:s(t),z):E},z.nodeWidth=function(t){return arguments.length?(A=+t,z):A},z.nodePadding=function(e){return arguments.length?(t=+e,z):t},z.nodes=function(t){return arguments.length?(L="function"==typeof t?t:s(t),z):L},z.links=function(t){return arguments.length?(C="function"==typeof t?t:s(t),z):C},z.size=function(t){return arguments.length?(i=a=0,b=+t[0],T=+t[1],z):[b-i,T-a]},z.extent=function(t){return arguments.length?(i=+t[0][0],b=+t[1][0],a=+t[0][1],T=+t[1][1],z):[[i,a],[b,T]]},z.iterations=function(t){return arguments.length?(P=+t,z):P},z.circularLinkGap=function(t){return arguments.length?(I=+t,z):I},z.nodePaddingRatio=function(t){return arguments.length?(n=+t,z):n},z.sortNodes=function(t){return arguments.length?(O=t,z):O},z.update=function(t){return w(t,M),V(t),t.links.forEach((function(t){t.circular&&(t.circularLinkType=t.y0+t.y1<T?"top":"bottom",t.source.circularLinkType=t.circularLinkType,t.target.circularLinkType=t.circularLinkType)})),j(t,T,M,!1),U(t,T,M),S(t,I,T,M),t},z},t.sankeyCenter=function(t){return t.targetLinks.length?t.depth:t.sourceLinks.length?e.min(t.sourceLinks,a)-1:0},t.sankeyLeft=function(t){return t.depth},t.sankeyRight=function(t,e){return e-1-t.height},t.sankeyJustify=o,Object.defineProperty(t,"__esModule",{value:!0})}))},{"d3-array":102,"d3-collection":103,"d3-shape":114,"elementary-circuits-directed-graph":124}],57:[function(t,e,r){!function(n,i){"object"==typeof r&&void 0!==e?i(r,t("d3-array"),t("d3-collection"),t("d3-shape")):i(n.d3=n.d3||{},n.d3,n.d3,n.d3)}(this,(function(t,e,r,n){"use strict";function i(t){return t.target.depth}function a(t,e){return t.sourceLinks.length?t.depth:e-1}function o(t){return function(){return t}}function s(t,e){return c(t.source,e.source)||t.index-e.index}function l(t,e){return c(t.target,e.target)||t.index-e.index}function c(t,e){return t.y0-e.y0}function u(t){return t.value}function f(t){return(t.y0+t.y1)/2}function h(t){return f(t.source)*t.value}function p(t){return f(t.target)*t.value}function d(t){return t.index}function m(t){return t.nodes}function g(t){return t.links}function v(t,e){var r=t.get(e);if(!r)throw new Error("missing: "+e);return r}function y(t){return[t.source.x1,t.y0]}function x(t){return[t.target.x0,t.y1]}t.sankey=function(){var t=0,n=0,i=1,y=1,x=24,b=8,_=d,w=a,T=m,k=g,A=32;function M(){var t={nodes:T.apply(null,arguments),links:k.apply(null,arguments)};return S(t),E(t),L(t),C(t),P(t),t}function S(t){t.nodes.forEach((function(t,e){t.index=e,t.sourceLinks=[],t.targetLinks=[]}));var e=r.map(t.nodes,_);t.links.forEach((function(t,r){t.index=r;var n=t.source,i=t.target;"object"!=typeof n&&(n=t.source=v(e,n)),"object"!=typeof i&&(i=t.target=v(e,i)),n.sourceLinks.push(t),i.targetLinks.push(t)}))}function E(t){t.nodes.forEach((function(t){t.value=Math.max(e.sum(t.sourceLinks,u),e.sum(t.targetLinks,u))}))}function L(e){var r,n,a;for(r=e.nodes,n=[],a=0;r.length;++a,r=n,n=[])r.forEach((function(t){t.depth=a,t.sourceLinks.forEach((function(t){n.indexOf(t.target)<0&&n.push(t.target)}))}));for(r=e.nodes,n=[],a=0;r.length;++a,r=n,n=[])r.forEach((function(t){t.height=a,t.targetLinks.forEach((function(t){n.indexOf(t.source)<0&&n.push(t.source)}))}));var o=(i-t-x)/(a-1);e.nodes.forEach((function(e){e.x1=(e.x0=t+Math.max(0,Math.min(a-1,Math.floor(w.call(null,e,a))))*o)+x}))}function C(t){var i=r.nest().key((function(t){return t.x0})).sortKeys(e.ascending).entries(t.nodes).map((function(t){return t.values}));!function(){var r=e.max(i,(function(t){return t.length})),a=2/3*(y-n)/(r-1);b>a&&(b=a);var o=e.min(i,(function(t){return(y-n-(t.length-1)*b)/e.sum(t,u)}));i.forEach((function(t){t.forEach((function(t,e){t.y1=(t.y0=e)+t.value*o}))})),t.links.forEach((function(t){t.width=t.value*o}))}(),d();for(var a=1,o=A;o>0;--o)l(a*=.99),d(),s(a),d();function s(t){i.forEach((function(r){r.forEach((function(r){if(r.targetLinks.length){var n=(e.sum(r.targetLinks,h)/e.sum(r.targetLinks,u)-f(r))*t;r.y0+=n,r.y1+=n}}))}))}function l(t){i.slice().reverse().forEach((function(r){r.forEach((function(r){if(r.sourceLinks.length){var n=(e.sum(r.sourceLinks,p)/e.sum(r.sourceLinks,u)-f(r))*t;r.y0+=n,r.y1+=n}}))}))}function d(){i.forEach((function(t){var e,r,i,a=n,o=t.length;for(t.sort(c),i=0;i<o;++i)(r=a-(e=t[i]).y0)>0&&(e.y0+=r,e.y1+=r),a=e.y1+b;if((r=a-b-y)>0)for(a=e.y0-=r,e.y1-=r,i=o-2;i>=0;--i)(r=(e=t[i]).y1+b-a)>0&&(e.y0-=r,e.y1-=r),a=e.y0}))}}function P(t){t.nodes.forEach((function(t){t.sourceLinks.sort(l),t.targetLinks.sort(s)})),t.nodes.forEach((function(t){var e=t.y0,r=e;t.sourceLinks.forEach((function(t){t.y0=e+t.width/2,e+=t.width})),t.targetLinks.forEach((function(t){t.y1=r+t.width/2,r+=t.width}))}))}return M.update=function(t){return P(t),t},M.nodeId=function(t){return arguments.length?(_="function"==typeof t?t:o(t),M):_},M.nodeAlign=function(t){return arguments.length?(w="function"==typeof t?t:o(t),M):w},M.nodeWidth=function(t){return arguments.length?(x=+t,M):x},M.nodePadding=function(t){return arguments.length?(b=+t,M):b},M.nodes=function(t){return arguments.length?(T="function"==typeof t?t:o(t),M):T},M.links=function(t){return arguments.length?(k="function"==typeof t?t:o(t),M):k},M.size=function(e){return arguments.length?(t=n=0,i=+e[0],y=+e[1],M):[i-t,y-n]},M.extent=function(e){return arguments.length?(t=+e[0][0],i=+e[1][0],n=+e[0][1],y=+e[1][1],M):[[t,n],[i,y]]},M.iterations=function(t){return arguments.length?(A=+t,M):A},M},t.sankeyCenter=function(t){return t.targetLinks.length?t.depth:t.sourceLinks.length?e.min(t.sourceLinks,i)-1:0},t.sankeyLeft=function(t){return t.depth},t.sankeyRight=function(t,e){return e-1-t.height},t.sankeyJustify=a,t.sankeyLinkHorizontal=function(){return n.linkHorizontal().source(y).target(x)},Object.defineProperty(t,"__esModule",{value:!0})}))},{"d3-array":102,"d3-collection":103,"d3-shape":114}],58:[function(t,e,r){(function(){var t={version:"3.8.0"},r=[].slice,n=function(t){return r.call(t)},i=self.document;function a(t){return t&&(t.ownerDocument||t.document||t).documentElement}function o(t){return t&&(t.ownerDocument&&t.ownerDocument.defaultView||t.document&&t||t.defaultView)}if(i)try{n(i.documentElement.childNodes)[0].nodeType}catch(t){n=function(t){for(var e=t.length,r=new Array(e);e--;)r[e]=t[e];return r}}if(Date.now||(Date.now=function(){return+new Date}),i)try{i.createElement("DIV").style.setProperty("opacity",0,"")}catch(t){var s=this.Element.prototype,l=s.setAttribute,c=s.setAttributeNS,u=this.CSSStyleDeclaration.prototype,f=u.setProperty;s.setAttribute=function(t,e){l.call(this,t,e+"")},s.setAttributeNS=function(t,e,r){c.call(this,t,e,r+"")},u.setProperty=function(t,e,r){f.call(this,t,e+"",r)}}function h(t,e){return t<e?-1:t>e?1:t>=e?0:NaN}function p(t){return null===t?NaN:+t}function d(t){return!isNaN(t)}function m(t){return{left:function(e,r,n,i){for(arguments.length<3&&(n=0),arguments.length<4&&(i=e.length);n<i;){var a=n+i>>>1;t(e[a],r)<0?n=a+1:i=a}return n},right:function(e,r,n,i){for(arguments.length<3&&(n=0),arguments.length<4&&(i=e.length);n<i;){var a=n+i>>>1;t(e[a],r)>0?i=a:n=a+1}return n}}}t.ascending=h,t.descending=function(t,e){return e<t?-1:e>t?1:e>=t?0:NaN},t.min=function(t,e){var r,n,i=-1,a=t.length;if(1===arguments.length){for(;++i<a;)if(null!=(n=t[i])&&n>=n){r=n;break}for(;++i<a;)null!=(n=t[i])&&r>n&&(r=n)}else{for(;++i<a;)if(null!=(n=e.call(t,t[i],i))&&n>=n){r=n;break}for(;++i<a;)null!=(n=e.call(t,t[i],i))&&r>n&&(r=n)}return r},t.max=function(t,e){var r,n,i=-1,a=t.length;if(1===arguments.length){for(;++i<a;)if(null!=(n=t[i])&&n>=n){r=n;break}for(;++i<a;)null!=(n=t[i])&&n>r&&(r=n)}else{for(;++i<a;)if(null!=(n=e.call(t,t[i],i))&&n>=n){r=n;break}for(;++i<a;)null!=(n=e.call(t,t[i],i))&&n>r&&(r=n)}return r},t.extent=function(t,e){var r,n,i,a=-1,o=t.length;if(1===arguments.length){for(;++a<o;)if(null!=(n=t[a])&&n>=n){r=i=n;break}for(;++a<o;)null!=(n=t[a])&&(r>n&&(r=n),i<n&&(i=n))}else{for(;++a<o;)if(null!=(n=e.call(t,t[a],a))&&n>=n){r=i=n;break}for(;++a<o;)null!=(n=e.call(t,t[a],a))&&(r>n&&(r=n),i<n&&(i=n))}return[r,i]},t.sum=function(t,e){var r,n=0,i=t.length,a=-1;if(1===arguments.length)for(;++a<i;)d(r=+t[a])&&(n+=r);else for(;++a<i;)d(r=+e.call(t,t[a],a))&&(n+=r);return n},t.mean=function(t,e){var r,n=0,i=t.length,a=-1,o=i;if(1===arguments.length)for(;++a<i;)d(r=p(t[a]))?n+=r:--o;else for(;++a<i;)d(r=p(e.call(t,t[a],a)))?n+=r:--o;if(o)return n/o},t.quantile=function(t,e){var r=(t.length-1)*e+1,n=Math.floor(r),i=+t[n-1],a=r-n;return a?i+a*(t[n]-i):i},t.median=function(e,r){var n,i=[],a=e.length,o=-1;if(1===arguments.length)for(;++o<a;)d(n=p(e[o]))&&i.push(n);else for(;++o<a;)d(n=p(r.call(e,e[o],o)))&&i.push(n);if(i.length)return t.quantile(i.sort(h),.5)},t.variance=function(t,e){var r,n,i=t.length,a=0,o=0,s=-1,l=0;if(1===arguments.length)for(;++s<i;)d(r=p(t[s]))&&(o+=(n=r-a)*(r-(a+=n/++l)));else for(;++s<i;)d(r=p(e.call(t,t[s],s)))&&(o+=(n=r-a)*(r-(a+=n/++l)));if(l>1)return o/(l-1)},t.deviation=function(){var e=t.variance.apply(this,arguments);return e?Math.sqrt(e):e};var g=m(h);function v(t){return t.length}t.bisectLeft=g.left,t.bisect=t.bisectRight=g.right,t.bisector=function(t){return m(1===t.length?function(e,r){return h(t(e),r)}:t)},t.shuffle=function(t,e,r){(a=arguments.length)<3&&(r=t.length,a<2&&(e=0));for(var n,i,a=r-e;a;)i=Math.random()*a--|0,n=t[a+e],t[a+e]=t[i+e],t[i+e]=n;return t},t.permute=function(t,e){for(var r=e.length,n=new Array(r);r--;)n[r]=t[e[r]];return n},t.pairs=function(t){for(var e=0,r=t.length-1,n=t[0],i=new Array(r<0?0:r);e<r;)i[e]=[n,n=t[++e]];return i},t.transpose=function(e){if(!(a=e.length))return[];for(var r=-1,n=t.min(e,v),i=new Array(n);++r<n;)for(var a,o=-1,s=i[r]=new Array(a);++o<a;)s[o]=e[o][r];return i},t.zip=function(){return t.transpose(arguments)},t.keys=function(t){var e=[];for(var r in t)e.push(r);return e},t.values=function(t){var e=[];for(var r in t)e.push(t[r]);return e},t.entries=function(t){var e=[];for(var r in t)e.push({key:r,value:t[r]});return e},t.merge=function(t){for(var e,r,n,i=t.length,a=-1,o=0;++a<i;)o+=t[a].length;for(r=new Array(o);--i>=0;)for(e=(n=t[i]).length;--e>=0;)r[--o]=n[e];return r};var y=Math.abs;function x(t){for(var e=1;t*e%1;)e*=10;return e}function b(t,e){for(var r in e)Object.defineProperty(t.prototype,r,{value:e[r],enumerable:!1})}function _(){this._=Object.create(null)}t.range=function(t,e,r){if(arguments.length<3&&(r=1,arguments.length<2&&(e=t,t=0)),(e-t)/r==1/0)throw new Error("infinite range");var n,i=[],a=x(y(r)),o=-1;if(t*=a,e*=a,(r*=a)<0)for(;(n=t+r*++o)>e;)i.push(n/a);else for(;(n=t+r*++o)<e;)i.push(n/a);return i},t.map=function(t,e){var r=new _;if(t instanceof _)t.forEach((function(t,e){r.set(t,e)}));else if(Array.isArray(t)){var n,i=-1,a=t.length;if(1===arguments.length)for(;++i<a;)r.set(i,t[i]);else for(;++i<a;)r.set(e.call(t,n=t[i],i),n)}else for(var o in t)r.set(o,t[o]);return r};function w(t){return"__proto__"==(t+="")||"\0"===t[0]?"\0"+t:t}function T(t){return"\0"===(t+="")[0]?t.slice(1):t}function k(t){return w(t)in this._}function A(t){return(t=w(t))in this._&&delete this._[t]}function M(){var t=[];for(var e in this._)t.push(T(e));return t}function S(){var t=0;for(var e in this._)++t;return t}function E(){for(var t in this._)return!1;return!0}function L(){this._=Object.create(null)}function C(t){return t}function P(t,e,r){return function(){var n=r.apply(e,arguments);return n===e?t:n}}function I(t,e){if(e in t)return e;e=e.charAt(0).toUpperCase()+e.slice(1);for(var r=0,n=O.length;r<n;++r){var i=O[r]+e;if(i in t)return i}}b(_,{has:k,get:function(t){return this._[w(t)]},set:function(t,e){return this._[w(t)]=e},remove:A,keys:M,values:function(){var t=[];for(var e in this._)t.push(this._[e]);return t},entries:function(){var t=[];for(var e in this._)t.push({key:T(e),value:this._[e]});return t},size:S,empty:E,forEach:function(t){for(var e in this._)t.call(this,T(e),this._[e])}}),t.nest=function(){var e,r,n={},i=[],a=[];function o(t,a,s){if(s>=i.length)return r?r.call(n,a):e?a.sort(e):a;for(var l,c,u,f,h=-1,p=a.length,d=i[s++],m=new _;++h<p;)(f=m.get(l=d(c=a[h])))?f.push(c):m.set(l,[c]);return t?(c=t(),u=function(e,r){c.set(e,o(t,r,s))}):(c={},u=function(e,r){c[e]=o(t,r,s)}),m.forEach(u),c}return n.map=function(t,e){return o(e,t,0)},n.entries=function(e){return function t(e,r){if(r>=i.length)return e;var n=[],o=a[r++];return e.forEach((function(e,i){n.push({key:e,values:t(i,r)})})),o?n.sort((function(t,e){return o(t.key,e.key)})):n}(o(t.map,e,0),0)},n.key=function(t){return i.push(t),n},n.sortKeys=function(t){return a[i.length-1]=t,n},n.sortValues=function(t){return e=t,n},n.rollup=function(t){return r=t,n},n},t.set=function(t){var e=new L;if(t)for(var r=0,n=t.length;r<n;++r)e.add(t[r]);return e},b(L,{has:k,add:function(t){return this._[w(t+="")]=!0,t},remove:A,values:M,size:S,empty:E,forEach:function(t){for(var e in this._)t.call(this,T(e))}}),t.behavior={},t.rebind=function(t,e){for(var r,n=1,i=arguments.length;++n<i;)t[r=arguments[n]]=P(t,e,e[r]);return t};var O=["webkit","ms","moz","Moz","o","O"];function z(){}function D(){}function R(t){var e=[],r=new _;function n(){for(var r,n=e,i=-1,a=n.length;++i<a;)(r=n[i].on)&&r.apply(this,arguments);return t}return n.on=function(n,i){var a,o=r.get(n);return arguments.length<2?o&&o.on:(o&&(o.on=null,e=e.slice(0,a=e.indexOf(o)).concat(e.slice(a+1)),r.remove(n)),i&&e.push(r.set(n,{on:i})),t)},n}function F(){t.event.preventDefault()}function B(){for(var e,r=t.event;e=r.sourceEvent;)r=e;return r}function N(e){for(var r=new D,n=0,i=arguments.length;++n<i;)r[arguments[n]]=R(r);return r.of=function(n,i){return function(a){try{var o=a.sourceEvent=t.event;a.target=e,t.event=a,r[a.type].apply(n,i)}finally{t.event=o}}},r}t.dispatch=function(){for(var t=new D,e=-1,r=arguments.length;++e<r;)t[arguments[e]]=R(t);return t},D.prototype.on=function(t,e){var r=t.indexOf("."),n="";if(r>=0&&(n=t.slice(r+1),t=t.slice(0,r)),t)return arguments.length<2?this[t].on(n):this[t].on(n,e);if(2===arguments.length){if(null==e)for(t in this)this.hasOwnProperty(t)&&this[t].on(n,null);return this}},t.event=null,t.requote=function(t){return t.replace(j,"\\$&")};var j=/[\\\^\$\*\+\?\|\[\]\(\)\.\{\}]/g,U={}.__proto__?function(t,e){t.__proto__=e}:function(t,e){for(var r in e)t[r]=e[r]};function V(t){return U(t,Y),t}var H=function(t,e){return e.querySelector(t)},q=function(t,e){return e.querySelectorAll(t)},G=function(t,e){var r=t.matches||t[I(t,"matchesSelector")];return(G=function(t,e){return r.call(t,e)})(t,e)};"function"==typeof Sizzle&&(H=function(t,e){return Sizzle(t,e)[0]||null},q=Sizzle,G=Sizzle.matchesSelector),t.selection=function(){return t.select(i.documentElement)};var Y=t.selection.prototype=[];function W(t){return"function"==typeof t?t:function(){return H(t,this)}}function X(t){return"function"==typeof t?t:function(){return q(t,this)}}Y.select=function(t){var e,r,n,i,a=[];t=W(t);for(var o=-1,s=this.length;++o<s;){a.push(e=[]),e.parentNode=(n=this[o]).parentNode;for(var l=-1,c=n.length;++l<c;)(i=n[l])?(e.push(r=t.call(i,i.__data__,l,o)),r&&"__data__"in i&&(r.__data__=i.__data__)):e.push(null)}return V(a)},Y.selectAll=function(t){var e,r,i=[];t=X(t);for(var a=-1,o=this.length;++a<o;)for(var s=this[a],l=-1,c=s.length;++l<c;)(r=s[l])&&(i.push(e=n(t.call(r,r.__data__,l,a))),e.parentNode=r);return V(i)};var Z="http://www.w3.org/1999/xhtml",J={svg:"http://www.w3.org/2000/svg",xhtml:Z,xlink:"http://www.w3.org/1999/xlink",xml:"http://www.w3.org/XML/1998/namespace",xmlns:"http://www.w3.org/2000/xmlns/"};function K(e,r){return e=t.ns.qualify(e),null==r?e.local?function(){this.removeAttributeNS(e.space,e.local)}:function(){this.removeAttribute(e)}:"function"==typeof r?e.local?function(){var t=r.apply(this,arguments);null==t?this.removeAttributeNS(e.space,e.local):this.setAttributeNS(e.space,e.local,t)}:function(){var t=r.apply(this,arguments);null==t?this.removeAttribute(e):this.setAttribute(e,t)}:e.local?function(){this.setAttributeNS(e.space,e.local,r)}:function(){this.setAttribute(e,r)}}function Q(t){return t.trim().replace(/\s+/g," ")}function $(e){return new RegExp("(?:^|\\s+)"+t.requote(e)+"(?:\\s+|$)","g")}function tt(t){return(t+"").trim().split(/^|\s+/)}function et(t,e){var r=(t=tt(t).map(rt)).length;return"function"==typeof e?function(){for(var n=-1,i=e.apply(this,arguments);++n<r;)t[n](this,i)}:function(){for(var n=-1;++n<r;)t[n](this,e)}}function rt(t){var e=$(t);return function(r,n){if(i=r.classList)return n?i.add(t):i.remove(t);var i=r.getAttribute("class")||"";n?(e.lastIndex=0,e.test(i)||r.setAttribute("class",Q(i+" "+t))):r.setAttribute("class",Q(i.replace(e," ")))}}function nt(t,e,r){return null==e?function(){this.style.removeProperty(t)}:"function"==typeof e?function(){var n=e.apply(this,arguments);null==n?this.style.removeProperty(t):this.style.setProperty(t,n,r)}:function(){this.style.setProperty(t,e,r)}}function it(t,e){return null==e?function(){delete this[t]}:"function"==typeof e?function(){var r=e.apply(this,arguments);null==r?delete this[t]:this[t]=r}:function(){this[t]=e}}function at(e){return"function"==typeof e?e:(e=t.ns.qualify(e)).local?function(){return this.ownerDocument.createElementNS(e.space,e.local)}:function(){var t=this.ownerDocument,r=this.namespaceURI;return r===Z&&t.documentElement.namespaceURI===Z?t.createElement(e):t.createElementNS(r,e)}}function ot(){var t=this.parentNode;t&&t.removeChild(this)}function st(t){return{__data__:t}}function lt(t){return function(){return G(this,t)}}function ct(t){return arguments.length||(t=h),function(e,r){return e&&r?t(e.__data__,r.__data__):!e-!r}}function ut(t,e){for(var r=0,n=t.length;r<n;r++)for(var i,a=t[r],o=0,s=a.length;o<s;o++)(i=a[o])&&e(i,o,r);return t}function ft(t){return U(t,ht),t}t.ns={prefix:J,qualify:function(t){var e=t.indexOf(":"),r=t;return e>=0&&"xmlns"!==(r=t.slice(0,e))&&(t=t.slice(e+1)),J.hasOwnProperty(r)?{space:J[r],local:t}:t}},Y.attr=function(e,r){if(arguments.length<2){if("string"==typeof e){var n=this.node();return(e=t.ns.qualify(e)).local?n.getAttributeNS(e.space,e.local):n.getAttribute(e)}for(r in e)this.each(K(r,e[r]));return this}return this.each(K(e,r))},Y.classed=function(t,e){if(arguments.length<2){if("string"==typeof t){var r=this.node(),n=(t=tt(t)).length,i=-1;if(e=r.classList){for(;++i<n;)if(!e.contains(t[i]))return!1}else for(e=r.getAttribute("class");++i<n;)if(!$(t[i]).test(e))return!1;return!0}for(e in t)this.each(et(e,t[e]));return this}return this.each(et(t,e))},Y.style=function(t,e,r){var n=arguments.length;if(n<3){if("string"!=typeof t){for(r in n<2&&(e=""),t)this.each(nt(r,t[r],e));return this}if(n<2){var i=this.node();return o(i).getComputedStyle(i,null).getPropertyValue(t)}r=""}return this.each(nt(t,e,r))},Y.property=function(t,e){if(arguments.length<2){if("string"==typeof t)return this.node()[t];for(e in t)this.each(it(e,t[e]));return this}return this.each(it(t,e))},Y.text=function(t){return arguments.length?this.each("function"==typeof t?function(){var e=t.apply(this,arguments);this.textContent=null==e?"":e}:null==t?function(){this.textContent=""}:function(){this.textContent=t}):this.node().textContent},Y.html=function(t){return arguments.length?this.each("function"==typeof t?function(){var e=t.apply(this,arguments);this.innerHTML=null==e?"":e}:null==t?function(){this.innerHTML=""}:function(){this.innerHTML=t}):this.node().innerHTML},Y.append=function(t){return t=at(t),this.select((function(){return this.appendChild(t.apply(this,arguments))}))},Y.insert=function(t,e){return t=at(t),e=W(e),this.select((function(){return this.insertBefore(t.apply(this,arguments),e.apply(this,arguments)||null)}))},Y.remove=function(){return this.each(ot)},Y.data=function(t,e){var r,n,i=-1,a=this.length;if(!arguments.length){for(t=new Array(a=(r=this[0]).length);++i<a;)(n=r[i])&&(t[i]=n.__data__);return t}function o(t,r){var n,i,a,o=t.length,u=r.length,f=Math.min(o,u),h=new Array(u),p=new Array(u),d=new Array(o);if(e){var m,g=new _,v=new Array(o);for(n=-1;++n<o;)(i=t[n])&&(g.has(m=e.call(i,i.__data__,n))?d[n]=i:g.set(m,i),v[n]=m);for(n=-1;++n<u;)(i=g.get(m=e.call(r,a=r[n],n)))?!0!==i&&(h[n]=i,i.__data__=a):p[n]=st(a),g.set(m,!0);for(n=-1;++n<o;)n in v&&!0!==g.get(v[n])&&(d[n]=t[n])}else{for(n=-1;++n<f;)i=t[n],a=r[n],i?(i.__data__=a,h[n]=i):p[n]=st(a);for(;n<u;++n)p[n]=st(r[n]);for(;n<o;++n)d[n]=t[n]}p.update=h,p.parentNode=h.parentNode=d.parentNode=t.parentNode,s.push(p),l.push(h),c.push(d)}var s=ft([]),l=V([]),c=V([]);if("function"==typeof t)for(;++i<a;)o(r=this[i],t.call(r,r.parentNode.__data__,i));else for(;++i<a;)o(r=this[i],t);return l.enter=function(){return s},l.exit=function(){return c},l},Y.datum=function(t){return arguments.length?this.property("__data__",t):this.property("__data__")},Y.filter=function(t){var e,r,n,i=[];"function"!=typeof t&&(t=lt(t));for(var a=0,o=this.length;a<o;a++){i.push(e=[]),e.parentNode=(r=this[a]).parentNode;for(var s=0,l=r.length;s<l;s++)(n=r[s])&&t.call(n,n.__data__,s,a)&&e.push(n)}return V(i)},Y.order=function(){for(var t=-1,e=this.length;++t<e;)for(var r,n=this[t],i=n.length-1,a=n[i];--i>=0;)(r=n[i])&&(a&&a!==r.nextSibling&&a.parentNode.insertBefore(r,a),a=r);return this},Y.sort=function(t){t=ct.apply(this,arguments);for(var e=-1,r=this.length;++e<r;)this[e].sort(t);return this.order()},Y.each=function(t){return ut(this,(function(e,r,n){t.call(e,e.__data__,r,n)}))},Y.call=function(t){var e=n(arguments);return t.apply(e[0]=this,e),this},Y.empty=function(){return!this.node()},Y.node=function(){for(var t=0,e=this.length;t<e;t++)for(var r=this[t],n=0,i=r.length;n<i;n++){var a=r[n];if(a)return a}return null},Y.size=function(){var t=0;return ut(this,(function(){++t})),t};var ht=[];function pt(t){var e,r;return function(n,i,a){var o,s=t[a].update,l=s.length;for(a!=r&&(r=a,e=0),i>=e&&(e=i+1);!(o=s[e])&&++e<l;);return o}}function dt(e,r,i){var a="__on"+e,o=e.indexOf("."),s=gt;o>0&&(e=e.slice(0,o));var l=mt.get(e);function c(){var t=this[a];t&&(this.removeEventListener(e,t,t.$),delete this[a])}return l&&(e=l,s=vt),o?r?function(){var t=s(r,n(arguments));c.call(this),this.addEventListener(e,this[a]=t,t.$=i),t._=r}:c:r?z:function(){var r,n=new RegExp("^__on([^.]+)"+t.requote(e)+"$");for(var i in this)if(r=i.match(n)){var a=this[i];this.removeEventListener(r[1],a,a.$),delete this[i]}}}t.selection.enter=ft,t.selection.enter.prototype=ht,ht.append=Y.append,ht.empty=Y.empty,ht.node=Y.node,ht.call=Y.call,ht.size=Y.size,ht.select=function(t){for(var e,r,n,i,a,o=[],s=-1,l=this.length;++s<l;){n=(i=this[s]).update,o.push(e=[]),e.parentNode=i.parentNode;for(var c=-1,u=i.length;++c<u;)(a=i[c])?(e.push(n[c]=r=t.call(i.parentNode,a.__data__,c,s)),r.__data__=a.__data__):e.push(null)}return V(o)},ht.insert=function(t,e){return arguments.length<2&&(e=pt(this)),Y.insert.call(this,t,e)},t.select=function(t){var e;return"string"==typeof t?(e=[H(t,i)]).parentNode=i.documentElement:(e=[t]).parentNode=a(t),V([e])},t.selectAll=function(t){var e;return"string"==typeof t?(e=n(q(t,i))).parentNode=i.documentElement:(e=n(t)).parentNode=null,V([e])},Y.on=function(t,e,r){var n=arguments.length;if(n<3){if("string"!=typeof t){for(r in n<2&&(e=!1),t)this.each(dt(r,t[r],e));return this}if(n<2)return(n=this.node()["__on"+t])&&n._;r=!1}return this.each(dt(t,e,r))};var mt=t.map({mouseenter:"mouseover",mouseleave:"mouseout"});function gt(e,r){return function(n){var i=t.event;t.event=n,r[0]=this.__data__;try{e.apply(this,r)}finally{t.event=i}}}function vt(t,e){var r=gt(t,e);return function(t){var e=t.relatedTarget;e&&(e===this||8&e.compareDocumentPosition(this))||r.call(this,t)}}i&&mt.forEach((function(t){"on"+t in i&&mt.remove(t)}));var yt,xt=0;function bt(e){var r=".dragsuppress-"+ ++xt,n="click"+r,i=t.select(o(e)).on("touchmove"+r,F).on("dragstart"+r,F).on("selectstart"+r,F);if(null==yt&&(yt=!("onselectstart"in e)&&I(e.style,"userSelect")),yt){var s=a(e).style,l=s[yt];s[yt]="none"}return function(t){if(i.on(r,null),yt&&(s[yt]=l),t){var e=function(){i.on(n,null)};i.on(n,(function(){F(),e()}),!0),setTimeout(e,0)}}}t.mouse=function(t){return wt(t,B())};var _t=this.navigator&&/WebKit/.test(this.navigator.userAgent)?-1:0;function wt(e,r){r.changedTouches&&(r=r.changedTouches[0]);var n=e.ownerSVGElement||e;if(n.createSVGPoint){var i=n.createSVGPoint();if(_t<0){var a=o(e);if(a.scrollX||a.scrollY){var s=(n=t.select("body").append("svg").style({position:"absolute",top:0,left:0,margin:0,padding:0,border:"none"},"important"))[0][0].getScreenCTM();_t=!(s.f||s.e),n.remove()}}return _t?(i.x=r.pageX,i.y=r.pageY):(i.x=r.clientX,i.y=r.clientY),[(i=i.matrixTransform(e.getScreenCTM().inverse())).x,i.y]}var l=e.getBoundingClientRect();return[r.clientX-l.left-e.clientLeft,r.clientY-l.top-e.clientTop]}function Tt(){return t.event.changedTouches[0].identifier}t.touch=function(t,e,r){if(arguments.length<3&&(r=e,e=B().changedTouches),e)for(var n,i=0,a=e.length;i<a;++i)if((n=e[i]).identifier===r)return wt(t,n)},t.behavior.drag=function(){var e=N(a,"drag","dragstart","dragend"),r=null,n=s(z,t.mouse,o,"mousemove","mouseup"),i=s(Tt,t.touch,C,"touchmove","touchend");function a(){this.on("mousedown.drag",n).on("touchstart.drag",i)}function s(n,i,a,o,s){return function(){var l,c=this,u=t.event.target.correspondingElement||t.event.target,f=c.parentNode,h=e.of(c,arguments),p=0,d=n(),m=".drag"+(null==d?"":"-"+d),g=t.select(a(u)).on(o+m,x).on(s+m,b),v=bt(u),y=i(f,d);function x(){var t,e,r=i(f,d);r&&(t=r[0]-y[0],e=r[1]-y[1],p|=t|e,y=r,h({type:"drag",x:r[0]+l[0],y:r[1]+l[1],dx:t,dy:e}))}function b(){i(f,d)&&(g.on(o+m,null).on(s+m,null),v(p),h({type:"dragend"}))}l=r?[(l=r.apply(c,arguments)).x-y[0],l.y-y[1]]:[0,0],h({type:"dragstart"})}}return a.origin=function(t){return arguments.length?(r=t,a):r},t.rebind(a,e,"on")},t.touches=function(t,e){return arguments.length<2&&(e=B().touches),e?n(e).map((function(e){var r=wt(t,e);return r.identifier=e.identifier,r})):[]};var kt=1e-6,At=Math.PI,Mt=2*At,St=Mt-kt,Et=At/2,Lt=At/180,Ct=180/At;function Pt(t){return t>1?Et:t<-1?-Et:Math.asin(t)}function It(t){return((t=Math.exp(t))+1/t)/2}var Ot=Math.SQRT2;t.interpolateZoom=function(t,e){var r,n,i=t[0],a=t[1],o=t[2],s=e[0],l=e[1],c=e[2],u=s-i,f=l-a,h=u*u+f*f;if(h<1e-12)n=Math.log(c/o)/Ot,r=function(t){return[i+t*u,a+t*f,o*Math.exp(Ot*t*n)]};else{var p=Math.sqrt(h),d=(c*c-o*o+4*h)/(2*o*2*p),m=(c*c-o*o-4*h)/(2*c*2*p),g=Math.log(Math.sqrt(d*d+1)-d),v=Math.log(Math.sqrt(m*m+1)-m);n=(v-g)/Ot,r=function(t){var e,r=t*n,s=It(g),l=o/(2*p)*(s*(e=Ot*r+g,((e=Math.exp(2*e))-1)/(e+1))-function(t){return((t=Math.exp(t))-1/t)/2}(g));return[i+l*u,a+l*f,o*s/It(Ot*r+g)]}}return r.duration=1e3*n,r},t.behavior.zoom=function(){var e,r,n,a,s,l,c,u,f,h={x:0,y:0,k:1},p=[960,500],d=Rt,m=250,g=0,v="mousedown.zoom",y="mousemove.zoom",x="mouseup.zoom",b="touchstart.zoom",_=N(w,"zoomstart","zoom","zoomend");function w(t){t.on(v,P).on(Dt+".zoom",O).on("dblclick.zoom",z).on(b,I)}function T(t){return[(t[0]-h.x)/h.k,(t[1]-h.y)/h.k]}function k(t){h.k=Math.max(d[0],Math.min(d[1],t))}function A(t,e){e=function(t){return[t[0]*h.k+h.x,t[1]*h.k+h.y]}(e),h.x+=t[0]-e[0],h.y+=t[1]-e[1]}function M(e,n,i,a){e.__chart__={x:h.x,y:h.y,k:h.k},k(Math.pow(2,a)),A(r=n,i),e=t.select(e),m>0&&(e=e.transition().duration(m)),e.call(w.event)}function S(){c&&c.domain(l.range().map((function(t){return(t-h.x)/h.k})).map(l.invert)),f&&f.domain(u.range().map((function(t){return(t-h.y)/h.k})).map(u.invert))}function E(t){g++||t({type:"zoomstart"})}function L(t){S(),t({type:"zoom",scale:h.k,translate:[h.x,h.y]})}function C(t){--g||(t({type:"zoomend"}),r=null)}function P(){var e=this,r=_.of(e,arguments),n=0,i=t.select(o(e)).on(y,l).on(x,c),a=T(t.mouse(e)),s=bt(e);function l(){n=1,A(t.mouse(e),a),L(r)}function c(){i.on(y,null).on(x,null),s(n),C(r)}Di.call(e),E(r)}function I(){var e,r=this,n=_.of(r,arguments),i={},a=0,o=".zoom-"+t.event.changedTouches[0].identifier,l="touchmove"+o,c="touchend"+o,u=[],f=t.select(r),p=bt(r);function d(){var n=t.touches(r);return e=h.k,n.forEach((function(t){t.identifier in i&&(i[t.identifier]=T(t))})),n}function m(){var e=t.event.target;t.select(e).on(l,g).on(c,y),u.push(e);for(var n=t.event.changedTouches,o=0,f=n.length;o<f;++o)i[n[o].identifier]=null;var p=d(),m=Date.now();if(1===p.length){if(m-s<500){var v=p[0];M(r,v,i[v.identifier],Math.floor(Math.log(h.k)/Math.LN2)+1),F()}s=m}else if(p.length>1){v=p[0];var x=p[1],b=v[0]-x[0],_=v[1]-x[1];a=b*b+_*_}}function g(){var o,l,c,u,f=t.touches(r);Di.call(r);for(var h=0,p=f.length;h<p;++h,u=null)if(c=f[h],u=i[c.identifier]){if(l)break;o=c,l=u}if(u){var d=(d=c[0]-o[0])*d+(d=c[1]-o[1])*d,m=a&&Math.sqrt(d/a);o=[(o[0]+c[0])/2,(o[1]+c[1])/2],l=[(l[0]+u[0])/2,(l[1]+u[1])/2],k(m*e)}s=null,A(o,l),L(n)}function y(){if(t.event.touches.length){for(var e=t.event.changedTouches,r=0,a=e.length;r<a;++r)delete i[e[r].identifier];for(var s in i)return void d()}t.selectAll(u).on(o,null),f.on(v,P).on(b,I),p(),C(n)}m(),E(n),f.on(v,null).on(b,m)}function O(){var i=_.of(this,arguments);a?clearTimeout(a):(Di.call(this),e=T(r=n||t.mouse(this)),E(i)),a=setTimeout((function(){a=null,C(i)}),50),F(),k(Math.pow(2,.002*zt())*h.k),A(r,e),L(i)}function z(){var e=t.mouse(this),r=Math.log(h.k)/Math.LN2;M(this,e,T(e),t.event.shiftKey?Math.ceil(r)-1:Math.floor(r)+1)}return Dt||(Dt="onwheel"in i?(zt=function(){return-t.event.deltaY*(t.event.deltaMode?120:1)},"wheel"):"onmousewheel"in i?(zt=function(){return t.event.wheelDelta},"mousewheel"):(zt=function(){return-t.event.detail},"MozMousePixelScroll")),w.event=function(e){e.each((function(){var e=_.of(this,arguments),n=h;Bi?t.select(this).transition().each("start.zoom",(function(){h=this.__chart__||{x:0,y:0,k:1},E(e)})).tween("zoom:zoom",(function(){var i=p[0],a=p[1],o=r?r[0]:i/2,s=r?r[1]:a/2,l=t.interpolateZoom([(o-h.x)/h.k,(s-h.y)/h.k,i/h.k],[(o-n.x)/n.k,(s-n.y)/n.k,i/n.k]);return function(t){var r=l(t),n=i/r[2];this.__chart__=h={x:o-r[0]*n,y:s-r[1]*n,k:n},L(e)}})).each("interrupt.zoom",(function(){C(e)})).each("end.zoom",(function(){C(e)})):(this.__chart__=h,E(e),L(e),C(e))}))},w.translate=function(t){return arguments.length?(h={x:+t[0],y:+t[1],k:h.k},S(),w):[h.x,h.y]},w.scale=function(t){return arguments.length?(h={x:h.x,y:h.y,k:null},k(+t),S(),w):h.k},w.scaleExtent=function(t){return arguments.length?(d=null==t?Rt:[+t[0],+t[1]],w):d},w.center=function(t){return arguments.length?(n=t&&[+t[0],+t[1]],w):n},w.size=function(t){return arguments.length?(p=t&&[+t[0],+t[1]],w):p},w.duration=function(t){return arguments.length?(m=+t,w):m},w.x=function(t){return arguments.length?(c=t,l=t.copy(),h={x:0,y:0,k:1},w):c},w.y=function(t){return arguments.length?(f=t,u=t.copy(),h={x:0,y:0,k:1},w):f},t.rebind(w,_,"on")};var zt,Dt,Rt=[0,1/0];function Ft(){}function Bt(t,e,r){return this instanceof Bt?(this.h=+t,this.s=+e,void(this.l=+r)):arguments.length<2?t instanceof Bt?new Bt(t.h,t.s,t.l):ne(""+t,ie,Bt):new Bt(t,e,r)}t.color=Ft,Ft.prototype.toString=function(){return this.rgb()+""},t.hsl=Bt;var Nt=Bt.prototype=new Ft;function jt(t,e,r){var n,i;function a(t){return Math.round(255*function(t){return t>360?t-=360:t<0&&(t+=360),t<60?n+(i-n)*t/60:t<180?i:t<240?n+(i-n)*(240-t)/60:n}(t))}return t=isNaN(t)?0:(t%=360)<0?t+360:t,e=isNaN(e)||e<0?0:e>1?1:e,n=2*(r=r<0?0:r>1?1:r)-(i=r<=.5?r*(1+e):r+e-r*e),new Qt(a(t+120),a(t),a(t-120))}function Ut(e,r,n){return this instanceof Ut?(this.h=+e,this.c=+r,void(this.l=+n)):arguments.length<2?e instanceof Ut?new Ut(e.h,e.c,e.l):Xt(e instanceof qt?e.l:(e=ae((e=t.rgb(e)).r,e.g,e.b)).l,e.a,e.b):new Ut(e,r,n)}Nt.brighter=function(t){return t=Math.pow(.7,arguments.length?t:1),new Bt(this.h,this.s,this.l/t)},Nt.darker=function(t){return t=Math.pow(.7,arguments.length?t:1),new Bt(this.h,this.s,t*this.l)},Nt.rgb=function(){return jt(this.h,this.s,this.l)},t.hcl=Ut;var Vt=Ut.prototype=new Ft;function Ht(t,e,r){return isNaN(t)&&(t=0),isNaN(e)&&(e=0),new qt(r,Math.cos(t*=Lt)*e,Math.sin(t)*e)}function qt(t,e,r){return this instanceof qt?(this.l=+t,this.a=+e,void(this.b=+r)):arguments.length<2?t instanceof qt?new qt(t.l,t.a,t.b):t instanceof Ut?Ht(t.h,t.c,t.l):ae((t=Qt(t)).r,t.g,t.b):new qt(t,e,r)}Vt.brighter=function(t){return new Ut(this.h,this.c,Math.min(100,this.l+Gt*(arguments.length?t:1)))},Vt.darker=function(t){return new Ut(this.h,this.c,Math.max(0,this.l-Gt*(arguments.length?t:1)))},Vt.rgb=function(){return Ht(this.h,this.c,this.l).rgb()},t.lab=qt;var Gt=18,Yt=qt.prototype=new Ft;function Wt(t,e,r){var n=(t+16)/116,i=n+e/500,a=n-r/200;return new Qt(Kt(3.2404542*(i=.95047*Zt(i))-1.5371385*(n=1*Zt(n))-.4985314*(a=1.08883*Zt(a))),Kt(-.969266*i+1.8760108*n+.041556*a),Kt(.0556434*i-.2040259*n+1.0572252*a))}function Xt(t,e,r){return t>0?new Ut(Math.atan2(r,e)*Ct,Math.sqrt(e*e+r*r),t):new Ut(NaN,NaN,t)}function Zt(t){return t>.206893034?t*t*t:(t-4/29)/7.787037}function Jt(t){return t>.008856?Math.pow(t,1/3):7.787037*t+4/29}function Kt(t){return Math.round(255*(t<=.00304?12.92*t:1.055*Math.pow(t,1/2.4)-.055))}function Qt(t,e,r){return this instanceof Qt?(this.r=~~t,this.g=~~e,void(this.b=~~r)):arguments.length<2?t instanceof Qt?new Qt(t.r,t.g,t.b):ne(""+t,Qt,jt):new Qt(t,e,r)}function $t(t){return new Qt(t>>16,t>>8&255,255&t)}function te(t){return $t(t)+""}Yt.brighter=function(t){return new qt(Math.min(100,this.l+Gt*(arguments.length?t:1)),this.a,this.b)},Yt.darker=function(t){return new qt(Math.max(0,this.l-Gt*(arguments.length?t:1)),this.a,this.b)},Yt.rgb=function(){return Wt(this.l,this.a,this.b)},t.rgb=Qt;var ee=Qt.prototype=new Ft;function re(t){return t<16?"0"+Math.max(0,t).toString(16):Math.min(255,t).toString(16)}function ne(t,e,r){var n,i,a,o=0,s=0,l=0;if(n=/([a-z]+)\((.*)\)/.exec(t=t.toLowerCase()))switch(i=n[2].split(","),n[1]){case"hsl":return r(parseFloat(i[0]),parseFloat(i[1])/100,parseFloat(i[2])/100);case"rgb":return e(se(i[0]),se(i[1]),se(i[2]))}return(a=le.get(t))?e(a.r,a.g,a.b):(null==t||"#"!==t.charAt(0)||isNaN(a=parseInt(t.slice(1),16))||(4===t.length?(o=(3840&a)>>4,o|=o>>4,s=240&a,s|=s>>4,l=15&a,l|=l<<4):7===t.length&&(o=(16711680&a)>>16,s=(65280&a)>>8,l=255&a)),e(o,s,l))}function ie(t,e,r){var n,i,a=Math.min(t/=255,e/=255,r/=255),o=Math.max(t,e,r),s=o-a,l=(o+a)/2;return s?(i=l<.5?s/(o+a):s/(2-o-a),n=t==o?(e-r)/s+(e<r?6:0):e==o?(r-t)/s+2:(t-e)/s+4,n*=60):(n=NaN,i=l>0&&l<1?0:n),new Bt(n,i,l)}function ae(t,e,r){var n=Jt((.4124564*(t=oe(t))+.3575761*(e=oe(e))+.1804375*(r=oe(r)))/.95047),i=Jt((.2126729*t+.7151522*e+.072175*r)/1);return qt(116*i-16,500*(n-i),200*(i-Jt((.0193339*t+.119192*e+.9503041*r)/1.08883)))}function oe(t){return(t/=255)<=.04045?t/12.92:Math.pow((t+.055)/1.055,2.4)}function se(t){var e=parseFloat(t);return"%"===t.charAt(t.length-1)?Math.round(2.55*e):e}ee.brighter=function(t){t=Math.pow(.7,arguments.length?t:1);var e=this.r,r=this.g,n=this.b,i=30;return e||r||n?(e&&e<i&&(e=i),r&&r<i&&(r=i),n&&n<i&&(n=i),new Qt(Math.min(255,e/t),Math.min(255,r/t),Math.min(255,n/t))):new Qt(i,i,i)},ee.darker=function(t){return new Qt((t=Math.pow(.7,arguments.length?t:1))*this.r,t*this.g,t*this.b)},ee.hsl=function(){return ie(this.r,this.g,this.b)},ee.toString=function(){return"#"+re(this.r)+re(this.g)+re(this.b)};var le=t.map({aliceblue:15792383,antiquewhite:16444375,aqua:65535,aquamarine:8388564,azure:15794175,beige:16119260,bisque:16770244,black:0,blanchedalmond:16772045,blue:255,blueviolet:9055202,brown:10824234,burlywood:14596231,cadetblue:6266528,chartreuse:8388352,chocolate:13789470,coral:16744272,cornflowerblue:6591981,cornsilk:16775388,crimson:14423100,cyan:65535,darkblue:139,darkcyan:35723,darkgoldenrod:12092939,darkgray:11119017,darkgreen:25600,darkgrey:11119017,darkkhaki:12433259,darkmagenta:9109643,darkolivegreen:5597999,darkorange:16747520,darkorchid:10040012,darkred:9109504,darksalmon:15308410,darkseagreen:9419919,darkslateblue:4734347,darkslategray:3100495,darkslategrey:3100495,darkturquoise:52945,darkviolet:9699539,deeppink:16716947,deepskyblue:49151,dimgray:6908265,dimgrey:6908265,dodgerblue:2003199,firebrick:11674146,floralwhite:16775920,forestgreen:2263842,fuchsia:16711935,gainsboro:14474460,ghostwhite:16316671,gold:16766720,goldenrod:14329120,gray:8421504,green:32768,greenyellow:11403055,grey:8421504,honeydew:15794160,hotpink:16738740,indianred:13458524,indigo:4915330,ivory:16777200,khaki:15787660,lavender:15132410,lavenderblush:16773365,lawngreen:8190976,lemonchiffon:16775885,lightblue:11393254,lightcoral:15761536,lightcyan:14745599,lightgoldenrodyellow:16448210,lightgray:13882323,lightgreen:9498256,lightgrey:13882323,lightpink:16758465,lightsalmon:16752762,lightseagreen:2142890,lightskyblue:8900346,lightslategray:7833753,lightslategrey:7833753,lightsteelblue:11584734,lightyellow:16777184,lime:65280,limegreen:3329330,linen:16445670,magenta:16711935,maroon:8388608,mediumaquamarine:6737322,mediumblue:205,mediumorchid:12211667,mediumpurple:9662683,mediumseagreen:3978097,mediumslateblue:8087790,mediumspringgreen:64154,mediumturquoise:4772300,mediumvioletred:13047173,midnightblue:1644912,mintcream:16121850,mistyrose:16770273,moccasin:16770229,navajowhite:16768685,navy:128,oldlace:16643558,olive:8421376,olivedrab:7048739,orange:16753920,orangered:16729344,orchid:14315734,palegoldenrod:15657130,palegreen:10025880,paleturquoise:11529966,palevioletred:14381203,papayawhip:16773077,peachpuff:16767673,peru:13468991,pink:16761035,plum:14524637,powderblue:11591910,purple:8388736,rebeccapurple:6697881,red:16711680,rosybrown:12357519,royalblue:4286945,saddlebrown:9127187,salmon:16416882,sandybrown:16032864,seagreen:3050327,seashell:16774638,sienna:10506797,silver:12632256,skyblue:8900331,slateblue:6970061,slategray:7372944,slategrey:7372944,snow:16775930,springgreen:65407,steelblue:4620980,tan:13808780,teal:32896,thistle:14204888,tomato:16737095,turquoise:4251856,violet:15631086,wheat:16113331,white:16777215,whitesmoke:16119285,yellow:16776960,yellowgreen:10145074});function ce(t){return"function"==typeof t?t:function(){return t}}function ue(t){return function(e,r,n){return 2===arguments.length&&"function"==typeof r&&(n=r,r=null),fe(e,r,t,n)}}function fe(e,r,i,a){var o={},s=t.dispatch("beforesend","progress","load","error"),l={},c=new XMLHttpRequest,u=null;function f(){var t,e=c.status;if(!e&&function(t){var e=t.responseType;return e&&"text"!==e?t.response:t.responseText}(c)||e>=200&&e<300||304===e){try{t=i.call(o,c)}catch(t){return void s.error.call(o,t)}s.load.call(o,t)}else s.error.call(o,c)}return self.XDomainRequest&&!("withCredentials"in c)&&/^(http(s)?:)?\/\//.test(e)&&(c=new XDomainRequest),"onload"in c?c.onload=c.onerror=f:c.onreadystatechange=function(){c.readyState>3&&f()},c.onprogress=function(e){var r=t.event;t.event=e;try{s.progress.call(o,c)}finally{t.event=r}},o.header=function(t,e){return t=(t+"").toLowerCase(),arguments.length<2?l[t]:(null==e?delete l[t]:l[t]=e+"",o)},o.mimeType=function(t){return arguments.length?(r=null==t?null:t+"",o):r},o.responseType=function(t){return arguments.length?(u=t,o):u},o.response=function(t){return i=t,o},["get","post"].forEach((function(t){o[t]=function(){return o.send.apply(o,[t].concat(n(arguments)))}})),o.send=function(t,n,i){if(2===arguments.length&&"function"==typeof n&&(i=n,n=null),c.open(t,e,!0),null==r||"accept"in l||(l.accept=r+",*/*"),c.setRequestHeader)for(var a in l)c.setRequestHeader(a,l[a]);return null!=r&&c.overrideMimeType&&c.overrideMimeType(r),null!=u&&(c.responseType=u),null!=i&&o.on("error",i).on("load",(function(t){i(null,t)})),s.beforesend.call(o,c),c.send(null==n?null:n),o},o.abort=function(){return c.abort(),o},t.rebind(o,s,"on"),null==a?o:o.get(function(t){return 1===t.length?function(e,r){t(null==e?r:null)}:t}(a))}le.forEach((function(t,e){le.set(t,$t(e))})),t.functor=ce,t.xhr=ue(C),t.dsv=function(t,e){var r=new RegExp('["'+t+"\n]"),n=t.charCodeAt(0);function i(t,r,n){arguments.length<3&&(n=r,r=null);var i=fe(t,e,null==r?a:o(r),n);return i.row=function(t){return arguments.length?i.response(null==(r=t)?a:o(t)):r},i}function a(t){return i.parse(t.responseText)}function o(t){return function(e){return i.parse(e.responseText,t)}}function s(e){return e.map(l).join(t)}function l(t){return r.test(t)?'"'+t.replace(/\"/g,'""')+'"':t}return i.parse=function(t,e){var r;return i.parseRows(t,(function(t,n){if(r)return r(t,n-1);var i=function(e){for(var r={},n=t.length,i=0;i<n;++i)r[t[i]]=e[i];return r};r=e?function(t,r){return e(i(t),r)}:i}))},i.parseRows=function(t,e){var r,i,a={},o={},s=[],l=t.length,c=0,u=0;function f(){if(c>=l)return o;if(i)return i=!1,a;var e=c;if(34===t.charCodeAt(e)){for(var r=e;r++<l;)if(34===t.charCodeAt(r)){if(34!==t.charCodeAt(r+1))break;++r}return c=r+2,13===(s=t.charCodeAt(r+1))?(i=!0,10===t.charCodeAt(r+2)&&++c):10===s&&(i=!0),t.slice(e+1,r).replace(/""/g,'"')}for(;c<l;){var s,u=1;if(10===(s=t.charCodeAt(c++)))i=!0;else if(13===s)i=!0,10===t.charCodeAt(c)&&(++c,++u);else if(s!==n)continue;return t.slice(e,c-u)}return t.slice(e)}for(;(r=f())!==o;){for(var h=[];r!==a&&r!==o;)h.push(r),r=f();e&&null==(h=e(h,u++))||s.push(h)}return s},i.format=function(e){if(Array.isArray(e[0]))return i.formatRows(e);var r=new L,n=[];return e.forEach((function(t){for(var e in t)r.has(e)||n.push(r.add(e))})),[n.map(l).join(t)].concat(e.map((function(e){return n.map((function(t){return l(e[t])})).join(t)}))).join("\n")},i.formatRows=function(t){return t.map(s).join("\n")},i},t.csv=t.dsv(",","text/csv"),t.tsv=t.dsv("\t","text/tab-separated-values");var he,pe,de,me,ge=this[I(this,"requestAnimationFrame")]||function(t){setTimeout(t,17)};function ve(t,e,r){var n=arguments.length;n<2&&(e=0),n<3&&(r=Date.now());var i=r+e,a={c:t,t:i,n:null};return pe?pe.n=a:he=a,pe=a,de||(me=clearTimeout(me),de=1,ge(ye)),a}function ye(){var t=xe(),e=be()-t;e>24?(isFinite(e)&&(clearTimeout(me),me=setTimeout(ye,e)),de=0):(de=1,ge(ye))}function xe(){for(var t=Date.now(),e=he;e;)t>=e.t&&e.c(t-e.t)&&(e.c=null),e=e.n;return t}function be(){for(var t,e=he,r=1/0;e;)e.c?(e.t<r&&(r=e.t),e=(t=e).n):e=t?t.n=e.n:he=e.n;return pe=t,r}function _e(t){return t[0]}function we(t){return t[1]}function Te(t){for(var e,r,n,i=t.length,a=[0,1],o=2,s=2;s<i;s++){for(;o>1&&(e=t[a[o-2]],r=t[a[o-1]],n=t[s],(r[0]-e[0])*(n[1]-e[1])-(r[1]-e[1])*(n[0]-e[0])<=0);)--o;a[o++]=s}return a.slice(0,o)}function ke(t,e){return t[0]-e[0]||t[1]-e[1]}t.timer=function(){ve.apply(this,arguments)},t.timer.flush=function(){xe(),be()},t.round=function(t,e){return e?Math.round(t*(e=Math.pow(10,e)))/e:Math.round(t)},t.geom={},t.geom.hull=function(t){var e=_e,r=we;if(arguments.length)return n(t);function n(t){if(t.length<3)return[];var n,i=ce(e),a=ce(r),o=t.length,s=[],l=[];for(n=0;n<o;n++)s.push([+i.call(this,t[n],n),+a.call(this,t[n],n),n]);for(s.sort(ke),n=0;n<o;n++)l.push([s[n][0],-s[n][1]]);var c=Te(s),u=Te(l),f=u[0]===c[0],h=u[u.length-1]===c[c.length-1],p=[];for(n=c.length-1;n>=0;--n)p.push(t[s[c[n]][2]]);for(n=+f;n<u.length-h;++n)p.push(t[s[u[n]][2]]);return p}return n.x=function(t){return arguments.length?(e=t,n):e},n.y=function(t){return arguments.length?(r=t,n):r},n},t.geom.polygon=function(t){return U(t,Ae),t};var Ae=t.geom.polygon.prototype=[];function Me(t,e,r){return(r[0]-e[0])*(t[1]-e[1])<(r[1]-e[1])*(t[0]-e[0])}function Se(t,e,r,n){var i=t[0],a=r[0],o=e[0]-i,s=n[0]-a,l=t[1],c=r[1],u=e[1]-l,f=n[1]-c,h=(s*(l-c)-f*(i-a))/(f*o-s*u);return[i+h*o,l+h*u]}function Ee(t){var e=t[0],r=t[t.length-1];return!(e[0]-r[0]||e[1]-r[1])}Ae.area=function(){for(var t,e=-1,r=this.length,n=this[r-1],i=0;++e<r;)t=n,n=this[e],i+=t[1]*n[0]-t[0]*n[1];return.5*i},Ae.centroid=function(t){var e,r,n=-1,i=this.length,a=0,o=0,s=this[i-1];for(arguments.length||(t=-1/(6*this.area()));++n<i;)e=s,s=this[n],r=e[0]*s[1]-s[0]*e[1],a+=(e[0]+s[0])*r,o+=(e[1]+s[1])*r;return[a*t,o*t]},Ae.clip=function(t){for(var e,r,n,i,a,o,s=Ee(t),l=-1,c=this.length-Ee(this),u=this[c-1];++l<c;){for(e=t.slice(),t.length=0,i=this[l],a=e[(n=e.length-s)-1],r=-1;++r<n;)Me(o=e[r],u,i)?(Me(a,u,i)||t.push(Se(a,o,u,i)),t.push(o)):Me(a,u,i)&&t.push(Se(a,o,u,i)),a=o;s&&t.push(t[0]),u=i}return t};var Le,Ce,Pe,Ie,Oe,ze=[],De=[];function Re(){er(this),this.edge=this.site=this.circle=null}function Fe(t){var e=ze.pop()||new Re;return e.site=t,e}function Be(t){We(t),Pe.remove(t),ze.push(t),er(t)}function Ne(t){var e=t.circle,r=e.x,n=e.cy,i={x:r,y:n},a=t.P,o=t.N,s=[t];Be(t);for(var l=a;l.circle&&y(r-l.circle.x)<kt&&y(n-l.circle.cy)<kt;)a=l.P,s.unshift(l),Be(l),l=a;s.unshift(l),We(l);for(var c=o;c.circle&&y(r-c.circle.x)<kt&&y(n-c.circle.cy)<kt;)o=c.N,s.push(c),Be(c),c=o;s.push(c),We(c);var u,f=s.length;for(u=1;u<f;++u)c=s[u],l=s[u-1],Qe(c.edge,l.site,c.site,i);l=s[0],(c=s[f-1]).edge=Je(l.site,c.site,null,i),Ye(l),Ye(c)}function je(t){for(var e,r,n,i,a=t.x,o=t.y,s=Pe._;s;)if((n=Ue(s,o)-a)>kt)s=s.L;else{if(!((i=a-Ve(s,o))>kt)){n>-kt?(e=s.P,r=s):i>-kt?(e=s,r=s.N):e=r=s;break}if(!s.R){e=s;break}s=s.R}var l=Fe(t);if(Pe.insert(e,l),e||r){if(e===r)return We(e),r=Fe(e.site),Pe.insert(l,r),l.edge=r.edge=Je(e.site,l.site),Ye(e),void Ye(r);if(r){We(e),We(r);var c=e.site,u=c.x,f=c.y,h=t.x-u,p=t.y-f,d=r.site,m=d.x-u,g=d.y-f,v=2*(h*g-p*m),y=h*h+p*p,x=m*m+g*g,b={x:(g*y-p*x)/v+u,y:(h*x-m*y)/v+f};Qe(r.edge,c,d,b),l.edge=Je(c,t,null,b),r.edge=Je(t,d,null,b),Ye(e),Ye(r)}else l.edge=Je(e.site,l.site)}}function Ue(t,e){var r=t.site,n=r.x,i=r.y,a=i-e;if(!a)return n;var o=t.P;if(!o)return-1/0;var s=(r=o.site).x,l=r.y,c=l-e;if(!c)return s;var u=s-n,f=1/a-1/c,h=u/c;return f?(-h+Math.sqrt(h*h-2*f*(u*u/(-2*c)-l+c/2+i-a/2)))/f+n:(n+s)/2}function Ve(t,e){var r=t.N;if(r)return Ue(r,e);var n=t.site;return n.y===e?n.x:1/0}function He(t){this.site=t,this.edges=[]}function qe(t,e){return e.angle-t.angle}function Ge(){er(this),this.x=this.y=this.arc=this.site=this.cy=null}function Ye(t){var e=t.P,r=t.N;if(e&&r){var n=e.site,i=t.site,a=r.site;if(n!==a){var o=i.x,s=i.y,l=n.x-o,c=n.y-s,u=a.x-o,f=2*(l*(g=a.y-s)-c*u);if(!(f>=-1e-12)){var h=l*l+c*c,p=u*u+g*g,d=(g*h-c*p)/f,m=(l*p-u*h)/f,g=m+s,v=De.pop()||new Ge;v.arc=t,v.site=i,v.x=d+o,v.y=g+Math.sqrt(d*d+m*m),v.cy=g,t.circle=v;for(var y=null,x=Oe._;x;)if(v.y<x.y||v.y===x.y&&v.x<=x.x){if(!x.L){y=x.P;break}x=x.L}else{if(!x.R){y=x;break}x=x.R}Oe.insert(y,v),y||(Ie=v)}}}}function We(t){var e=t.circle;e&&(e.P||(Ie=e.N),Oe.remove(e),De.push(e),er(e),t.circle=null)}function Xe(t,e){var r=t.b;if(r)return!0;var n,i,a=t.a,o=e[0][0],s=e[1][0],l=e[0][1],c=e[1][1],u=t.l,f=t.r,h=u.x,p=u.y,d=f.x,m=f.y,g=(h+d)/2,v=(p+m)/2;if(m===p){if(g<o||g>=s)return;if(h>d){if(a){if(a.y>=c)return}else a={x:g,y:l};r={x:g,y:c}}else{if(a){if(a.y<l)return}else a={x:g,y:c};r={x:g,y:l}}}else if(i=v-(n=(h-d)/(m-p))*g,n<-1||n>1)if(h>d){if(a){if(a.y>=c)return}else a={x:(l-i)/n,y:l};r={x:(c-i)/n,y:c}}else{if(a){if(a.y<l)return}else a={x:(c-i)/n,y:c};r={x:(l-i)/n,y:l}}else if(p<m){if(a){if(a.x>=s)return}else a={x:o,y:n*o+i};r={x:s,y:n*s+i}}else{if(a){if(a.x<o)return}else a={x:s,y:n*s+i};r={x:o,y:n*o+i}}return t.a=a,t.b=r,!0}function Ze(t,e){this.l=t,this.r=e,this.a=this.b=null}function Je(t,e,r,n){var i=new Ze(t,e);return Le.push(i),r&&Qe(i,t,e,r),n&&Qe(i,e,t,n),Ce[t.i].edges.push(new $e(i,t,e)),Ce[e.i].edges.push(new $e(i,e,t)),i}function Ke(t,e,r){var n=new Ze(t,null);return n.a=e,n.b=r,Le.push(n),n}function Qe(t,e,r,n){t.a||t.b?t.l===r?t.b=n:t.a=n:(t.a=n,t.l=e,t.r=r)}function $e(t,e,r){var n=t.a,i=t.b;this.edge=t,this.site=e,this.angle=r?Math.atan2(r.y-e.y,r.x-e.x):t.l===e?Math.atan2(i.x-n.x,n.y-i.y):Math.atan2(n.x-i.x,i.y-n.y)}function tr(){this._=null}function er(t){t.U=t.C=t.L=t.R=t.P=t.N=null}function rr(t,e){var r=e,n=e.R,i=r.U;i?i.L===r?i.L=n:i.R=n:t._=n,n.U=i,r.U=n,r.R=n.L,r.R&&(r.R.U=r),n.L=r}function nr(t,e){var r=e,n=e.L,i=r.U;i?i.L===r?i.L=n:i.R=n:t._=n,n.U=i,r.U=n,r.L=n.R,r.L&&(r.L.U=r),n.R=r}function ir(t){for(;t.L;)t=t.L;return t}function ar(t,e){var r,n,i,a=t.sort(or).pop();for(Le=[],Ce=new Array(t.length),Pe=new tr,Oe=new tr;;)if(i=Ie,a&&(!i||a.y<i.y||a.y===i.y&&a.x<i.x))a.x===r&&a.y===n||(Ce[a.i]=new He(a),je(a),r=a.x,n=a.y),a=t.pop();else{if(!i)break;Ne(i.arc)}e&&(function(t){for(var e,r,n,i,a,o=Le,s=(r=t[0][0],n=t[0][1],i=t[1][0],a=t[1][1],function(t){var e,o=t.a,s=t.b,l=o.x,c=o.y,u=0,f=1,h=s.x-l,p=s.y-c;if(e=r-l,h||!(e>0)){if(e/=h,h<0){if(e<u)return;e<f&&(f=e)}else if(h>0){if(e>f)return;e>u&&(u=e)}if(e=i-l,h||!(e<0)){if(e/=h,h<0){if(e>f)return;e>u&&(u=e)}else if(h>0){if(e<u)return;e<f&&(f=e)}if(e=n-c,p||!(e>0)){if(e/=p,p<0){if(e<u)return;e<f&&(f=e)}else if(p>0){if(e>f)return;e>u&&(u=e)}if(e=a-c,p||!(e<0)){if(e/=p,p<0){if(e>f)return;e>u&&(u=e)}else if(p>0){if(e<u)return;e<f&&(f=e)}return u>0&&(t.a={x:l+u*h,y:c+u*p}),f<1&&(t.b={x:l+f*h,y:c+f*p}),t}}}}}),l=o.length;l--;)(!Xe(e=o[l],t)||!s(e)||y(e.a.x-e.b.x)<kt&&y(e.a.y-e.b.y)<kt)&&(e.a=e.b=null,o.splice(l,1))}(e),function(t){for(var e,r,n,i,a,o,s,l,c,u,f=t[0][0],h=t[1][0],p=t[0][1],d=t[1][1],m=Ce,g=m.length;g--;)if((a=m[g])&&a.prepare())for(l=(s=a.edges).length,o=0;o<l;)n=(u=s[o].end()).x,i=u.y,e=(c=s[++o%l].start()).x,r=c.y,(y(n-e)>kt||y(i-r)>kt)&&(s.splice(o,0,new $e(Ke(a.site,u,y(n-f)<kt&&d-i>kt?{x:f,y:y(e-f)<kt?r:d}:y(i-d)<kt&&h-n>kt?{x:y(r-d)<kt?e:h,y:d}:y(n-h)<kt&&i-p>kt?{x:h,y:y(e-h)<kt?r:p}:y(i-p)<kt&&n-f>kt?{x:y(r-p)<kt?e:f,y:p}:null),a.site,null)),++l)}(e));var o={cells:Ce,edges:Le};return Pe=Oe=Le=Ce=null,o}function or(t,e){return e.y-t.y||e.x-t.x}He.prototype.prepare=function(){for(var t,e=this.edges,r=e.length;r--;)(t=e[r].edge).b&&t.a||e.splice(r,1);return e.sort(qe),e.length},$e.prototype={start:function(){return this.edge.l===this.site?this.edge.a:this.edge.b},end:function(){return this.edge.l===this.site?this.edge.b:this.edge.a}},tr.prototype={insert:function(t,e){var r,n,i;if(t){if(e.P=t,e.N=t.N,t.N&&(t.N.P=e),t.N=e,t.R){for(t=t.R;t.L;)t=t.L;t.L=e}else t.R=e;r=t}else this._?(t=ir(this._),e.P=null,e.N=t,t.P=t.L=e,r=t):(e.P=e.N=null,this._=e,r=null);for(e.L=e.R=null,e.U=r,e.C=!0,t=e;r&&r.C;)r===(n=r.U).L?(i=n.R)&&i.C?(r.C=i.C=!1,n.C=!0,t=n):(t===r.R&&(rr(this,r),r=(t=r).U),r.C=!1,n.C=!0,nr(this,n)):(i=n.L)&&i.C?(r.C=i.C=!1,n.C=!0,t=n):(t===r.L&&(nr(this,r),r=(t=r).U),r.C=!1,n.C=!0,rr(this,n)),r=t.U;this._.C=!1},remove:function(t){t.N&&(t.N.P=t.P),t.P&&(t.P.N=t.N),t.N=t.P=null;var e,r,n,i=t.U,a=t.L,o=t.R;if(r=a?o?ir(o):a:o,i?i.L===t?i.L=r:i.R=r:this._=r,a&&o?(n=r.C,r.C=t.C,r.L=a,a.U=r,r!==o?(i=r.U,r.U=t.U,t=r.R,i.L=t,r.R=o,o.U=r):(r.U=i,i=r,t=r.R)):(n=t.C,t=r),t&&(t.U=i),!n)if(t&&t.C)t.C=!1;else{do{if(t===this._)break;if(t===i.L){if((e=i.R).C&&(e.C=!1,i.C=!0,rr(this,i),e=i.R),e.L&&e.L.C||e.R&&e.R.C){e.R&&e.R.C||(e.L.C=!1,e.C=!0,nr(this,e),e=i.R),e.C=i.C,i.C=e.R.C=!1,rr(this,i),t=this._;break}}else if((e=i.L).C&&(e.C=!1,i.C=!0,nr(this,i),e=i.L),e.L&&e.L.C||e.R&&e.R.C){e.L&&e.L.C||(e.R.C=!1,e.C=!0,rr(this,e),e=i.L),e.C=i.C,i.C=e.L.C=!1,nr(this,i),t=this._;break}e.C=!0,t=i,i=i.U}while(!t.C);t&&(t.C=!1)}}},t.geom.voronoi=function(t){var e=_e,r=we,n=e,i=r,a=sr;if(t)return o(t);function o(t){var e=new Array(t.length),r=a[0][0],n=a[0][1],i=a[1][0],o=a[1][1];return ar(s(t),a).cells.forEach((function(a,s){var l=a.edges,c=a.site;(e[s]=l.length?l.map((function(t){var e=t.start();return[e.x,e.y]})):c.x>=r&&c.x<=i&&c.y>=n&&c.y<=o?[[r,o],[i,o],[i,n],[r,n]]:[]).point=t[s]})),e}function s(t){return t.map((function(t,e){return{x:Math.round(n(t,e)/kt)*kt,y:Math.round(i(t,e)/kt)*kt,i:e}}))}return o.links=function(t){return ar(s(t)).edges.filter((function(t){return t.l&&t.r})).map((function(e){return{source:t[e.l.i],target:t[e.r.i]}}))},o.triangles=function(t){var e=[];return ar(s(t)).cells.forEach((function(r,n){for(var i,a,o,s,l=r.site,c=r.edges.sort(qe),u=-1,f=c.length,h=c[f-1].edge,p=h.l===l?h.r:h.l;++u<f;)h,i=p,p=(h=c[u].edge).l===l?h.r:h.l,n<i.i&&n<p.i&&(o=i,s=p,((a=l).x-s.x)*(o.y-a.y)-(a.x-o.x)*(s.y-a.y)<0)&&e.push([t[n],t[i.i],t[p.i]])})),e},o.x=function(t){return arguments.length?(n=ce(e=t),o):e},o.y=function(t){return arguments.length?(i=ce(r=t),o):r},o.clipExtent=function(t){return arguments.length?(a=null==t?sr:t,o):a===sr?null:a},o.size=function(t){return arguments.length?o.clipExtent(t&&[[0,0],t]):a===sr?null:a&&a[1]},o};var sr=[[-1e6,-1e6],[1e6,1e6]];function lr(t){return t.x}function cr(t){return t.y}function ur(t,e,r,n,i,a){if(!t(e,r,n,i,a)){var o=.5*(r+i),s=.5*(n+a),l=e.nodes;l[0]&&ur(t,l[0],r,n,o,s),l[1]&&ur(t,l[1],o,n,i,s),l[2]&&ur(t,l[2],r,s,o,a),l[3]&&ur(t,l[3],o,s,i,a)}}function fr(t,e,r,n,i,a,o){var s,l=1/0;return function t(c,u,f,h,p){if(!(u>a||f>o||h<n||p<i)){if(d=c.point){var d,m=e-c.x,g=r-c.y,v=m*m+g*g;if(v<l){var y=Math.sqrt(l=v);n=e-y,i=r-y,a=e+y,o=r+y,s=d}}for(var x=c.nodes,b=.5*(u+h),_=.5*(f+p),w=(r>=_)<<1|e>=b,T=w+4;w<T;++w)if(c=x[3&w])switch(3&w){case 0:t(c,u,f,b,_);break;case 1:t(c,b,f,h,_);break;case 2:t(c,u,_,b,p);break;case 3:t(c,b,_,h,p)}}}(t,n,i,a,o),s}function hr(e,r){e=t.rgb(e),r=t.rgb(r);var n=e.r,i=e.g,a=e.b,o=r.r-n,s=r.g-i,l=r.b-a;return function(t){return"#"+re(Math.round(n+o*t))+re(Math.round(i+s*t))+re(Math.round(a+l*t))}}function pr(t,e){var r,n={},i={};for(r in t)r in e?n[r]=yr(t[r],e[r]):i[r]=t[r];for(r in e)r in t||(i[r]=e[r]);return function(t){for(r in n)i[r]=n[r](t);return i}}function dr(t,e){return t=+t,e=+e,function(r){return t*(1-r)+e*r}}function mr(t,e){var r,n,i,a=gr.lastIndex=vr.lastIndex=0,o=-1,s=[],l=[];for(t+="",e+="";(r=gr.exec(t))&&(n=vr.exec(e));)(i=n.index)>a&&(i=e.slice(a,i),s[o]?s[o]+=i:s[++o]=i),(r=r[0])===(n=n[0])?s[o]?s[o]+=n:s[++o]=n:(s[++o]=null,l.push({i:o,x:dr(r,n)})),a=vr.lastIndex;return a<e.length&&(i=e.slice(a),s[o]?s[o]+=i:s[++o]=i),s.length<2?l[0]?(e=l[0].x,function(t){return e(t)+""}):function(){return e}:(e=l.length,function(t){for(var r,n=0;n<e;++n)s[(r=l[n]).i]=r.x(t);return s.join("")})}t.geom.delaunay=function(e){return t.geom.voronoi().triangles(e)},t.geom.quadtree=function(t,e,r,n,i){var a,o=_e,s=we;if(a=arguments.length)return o=lr,s=cr,3===a&&(i=r,n=e,r=e=0),l(t);function l(t){var l,c,u,f,h,p,d,m,g,v=ce(o),x=ce(s);if(null!=e)p=e,d=r,m=n,g=i;else if(m=g=-(p=d=1/0),c=[],u=[],h=t.length,a)for(f=0;f<h;++f)(l=t[f]).x<p&&(p=l.x),l.y<d&&(d=l.y),l.x>m&&(m=l.x),l.y>g&&(g=l.y),c.push(l.x),u.push(l.y);else for(f=0;f<h;++f){var b=+v(l=t[f],f),_=+x(l,f);b<p&&(p=b),_<d&&(d=_),b>m&&(m=b),_>g&&(g=_),c.push(b),u.push(_)}var w=m-p,T=g-d;function k(t,e,r,n,i,a,o,s){if(!isNaN(r)&&!isNaN(n))if(t.leaf){var l=t.x,c=t.y;if(null!=l)if(y(l-r)+y(c-n)<.01)A(t,e,r,n,i,a,o,s);else{var u=t.point;t.x=t.y=t.point=null,A(t,u,l,c,i,a,o,s),A(t,e,r,n,i,a,o,s)}else t.x=r,t.y=n,t.point=e}else A(t,e,r,n,i,a,o,s)}function A(t,e,r,n,i,a,o,s){var l=.5*(i+o),c=.5*(a+s),u=r>=l,f=n>=c,h=f<<1|u;t.leaf=!1,u?i=l:o=l,f?a=c:s=c,k(t=t.nodes[h]||(t.nodes[h]={leaf:!0,nodes:[],point:null,x:null,y:null}),e,r,n,i,a,o,s)}w>T?g=d+w:m=p+T;var M={leaf:!0,nodes:[],point:null,x:null,y:null,add:function(t){k(M,t,+v(t,++f),+x(t,f),p,d,m,g)},visit:function(t){ur(t,M,p,d,m,g)},find:function(t){return fr(M,t[0],t[1],p,d,m,g)}};if(f=-1,null==e){for(;++f<h;)k(M,t[f],c[f],u[f],p,d,m,g);--f}else t.forEach(M.add);return c=u=t=l=null,M}return l.x=function(t){return arguments.length?(o=t,l):o},l.y=function(t){return arguments.length?(s=t,l):s},l.extent=function(t){return arguments.length?(null==t?e=r=n=i=null:(e=+t[0][0],r=+t[0][1],n=+t[1][0],i=+t[1][1]),l):null==e?null:[[e,r],[n,i]]},l.size=function(t){return arguments.length?(null==t?e=r=n=i=null:(e=r=0,n=+t[0],i=+t[1]),l):null==e?null:[n-e,i-r]},l},t.interpolateRgb=hr,t.interpolateObject=pr,t.interpolateNumber=dr,t.interpolateString=mr;var gr=/[-+]?(?:\d+\.?\d*|\.?\d+)(?:[eE][-+]?\d+)?/g,vr=new RegExp(gr.source,"g");function yr(e,r){for(var n,i=t.interpolators.length;--i>=0&&!(n=t.interpolators[i](e,r)););return n}function xr(t,e){var r,n=[],i=[],a=t.length,o=e.length,s=Math.min(t.length,e.length);for(r=0;r<s;++r)n.push(yr(t[r],e[r]));for(;r<a;++r)i[r]=t[r];for(;r<o;++r)i[r]=e[r];return function(t){for(r=0;r<s;++r)i[r]=n[r](t);return i}}t.interpolate=yr,t.interpolators=[function(t,e){var r=typeof e;return("string"===r?le.has(e.toLowerCase())||/^(#|rgb\(|hsl\()/i.test(e)?hr:mr:e instanceof Ft?hr:Array.isArray(e)?xr:"object"===r&&isNaN(e)?pr:dr)(t,e)}],t.interpolateArray=xr;var br=function(){return C},_r=t.map({linear:br,poly:function(t){return function(e){return Math.pow(e,t)}},quad:function(){return Mr},cubic:function(){return Sr},sin:function(){return Lr},exp:function(){return Cr},circle:function(){return Pr},elastic:function(t,e){var r;arguments.length<2&&(e=.45);arguments.length?r=e/Mt*Math.asin(1/t):(t=1,r=e/4);return function(n){return 1+t*Math.pow(2,-10*n)*Math.sin((n-r)*Mt/e)}},back:function(t){t||(t=1.70158);return function(e){return e*e*((t+1)*e-t)}},bounce:function(){return Ir}}),wr=t.map({in:C,out:kr,"in-out":Ar,"out-in":function(t){return Ar(kr(t))}});function Tr(t){return function(e){return e<=0?0:e>=1?1:t(e)}}function kr(t){return function(e){return 1-t(1-e)}}function Ar(t){return function(e){return.5*(e<.5?t(2*e):2-t(2-2*e))}}function Mr(t){return t*t}function Sr(t){return t*t*t}function Er(t){if(t<=0)return 0;if(t>=1)return 1;var e=t*t,r=e*t;return 4*(t<.5?r:3*(t-e)+r-.75)}function Lr(t){return 1-Math.cos(t*Et)}function Cr(t){return Math.pow(2,10*(t-1))}function Pr(t){return 1-Math.sqrt(1-t*t)}function Ir(t){return t<1/2.75?7.5625*t*t:t<2/2.75?7.5625*(t-=1.5/2.75)*t+.75:t<2.5/2.75?7.5625*(t-=2.25/2.75)*t+.9375:7.5625*(t-=2.625/2.75)*t+.984375}function Or(t,e){return e-=t,function(r){return Math.round(t+e*r)}}function zr(t){var e,r,n,i=[t.a,t.b],a=[t.c,t.d],o=Rr(i),s=Dr(i,a),l=Rr(((e=a)[0]+=(n=-s)*(r=i)[0],e[1]+=n*r[1],e))||0;i[0]*a[1]<a[0]*i[1]&&(i[0]*=-1,i[1]*=-1,o*=-1,s*=-1),this.rotate=(o?Math.atan2(i[1],i[0]):Math.atan2(-a[0],a[1]))*Ct,this.translate=[t.e,t.f],this.scale=[o,l],this.skew=l?Math.atan2(s,l)*Ct:0}function Dr(t,e){return t[0]*e[0]+t[1]*e[1]}function Rr(t){var e=Math.sqrt(Dr(t,t));return e&&(t[0]/=e,t[1]/=e),e}t.ease=function(t){var e=t.indexOf("-"),n=e>=0?t.slice(0,e):t,i=e>=0?t.slice(e+1):"in";return n=_r.get(n)||br,Tr((i=wr.get(i)||C)(n.apply(null,r.call(arguments,1))))},t.interpolateHcl=function(e,r){e=t.hcl(e),r=t.hcl(r);var n=e.h,i=e.c,a=e.l,o=r.h-n,s=r.c-i,l=r.l-a;isNaN(s)&&(s=0,i=isNaN(i)?r.c:i);isNaN(o)?(o=0,n=isNaN(n)?r.h:n):o>180?o-=360:o<-180&&(o+=360);return function(t){return Ht(n+o*t,i+s*t,a+l*t)+""}},t.interpolateHsl=function(e,r){e=t.hsl(e),r=t.hsl(r);var n=e.h,i=e.s,a=e.l,o=r.h-n,s=r.s-i,l=r.l-a;isNaN(s)&&(s=0,i=isNaN(i)?r.s:i);isNaN(o)?(o=0,n=isNaN(n)?r.h:n):o>180?o-=360:o<-180&&(o+=360);return function(t){return jt(n+o*t,i+s*t,a+l*t)+""}},t.interpolateLab=function(e,r){e=t.lab(e),r=t.lab(r);var n=e.l,i=e.a,a=e.b,o=r.l-n,s=r.a-i,l=r.b-a;return function(t){return Wt(n+o*t,i+s*t,a+l*t)+""}},t.interpolateRound=Or,t.transform=function(e){var r=i.createElementNS(t.ns.prefix.svg,"g");return(t.transform=function(t){if(null!=t){r.setAttribute("transform",t);var e=r.transform.baseVal.consolidate()}return new zr(e?e.matrix:Fr)})(e)},zr.prototype.toString=function(){return"translate("+this.translate+")rotate("+this.rotate+")skewX("+this.skew+")scale("+this.scale+")"};var Fr={a:1,b:0,c:0,d:1,e:0,f:0};function Br(t){return t.length?t.pop()+",":""}function Nr(e,r){var n=[],i=[];return e=t.transform(e),r=t.transform(r),function(t,e,r,n){if(t[0]!==e[0]||t[1]!==e[1]){var i=r.push("translate(",null,",",null,")");n.push({i:i-4,x:dr(t[0],e[0])},{i:i-2,x:dr(t[1],e[1])})}else(e[0]||e[1])&&r.push("translate("+e+")")}(e.translate,r.translate,n,i),function(t,e,r,n){t!==e?(t-e>180?e+=360:e-t>180&&(t+=360),n.push({i:r.push(Br(r)+"rotate(",null,")")-2,x:dr(t,e)})):e&&r.push(Br(r)+"rotate("+e+")")}(e.rotate,r.rotate,n,i),function(t,e,r,n){t!==e?n.push({i:r.push(Br(r)+"skewX(",null,")")-2,x:dr(t,e)}):e&&r.push(Br(r)+"skewX("+e+")")}(e.skew,r.skew,n,i),function(t,e,r,n){if(t[0]!==e[0]||t[1]!==e[1]){var i=r.push(Br(r)+"scale(",null,",",null,")");n.push({i:i-4,x:dr(t[0],e[0])},{i:i-2,x:dr(t[1],e[1])})}else 1===e[0]&&1===e[1]||r.push(Br(r)+"scale("+e+")")}(e.scale,r.scale,n,i),e=r=null,function(t){for(var e,r=-1,a=i.length;++r<a;)n[(e=i[r]).i]=e.x(t);return n.join("")}}function jr(t,e){return e=(e-=t=+t)||1/e,function(r){return(r-t)/e}}function Ur(t,e){return e=(e-=t=+t)||1/e,function(r){return Math.max(0,Math.min(1,(r-t)/e))}}function Vr(t){for(var e=t.source,r=t.target,n=function(t,e){if(t===e)return t;var r=Hr(t),n=Hr(e),i=r.pop(),a=n.pop(),o=null;for(;i===a;)o=i,i=r.pop(),a=n.pop();return o}(e,r),i=[e];e!==n;)e=e.parent,i.push(e);for(var a=i.length;r!==n;)i.splice(a,0,r),r=r.parent;return i}function Hr(t){for(var e=[],r=t.parent;null!=r;)e.push(t),t=r,r=r.parent;return e.push(t),e}function qr(t){t.fixed|=2}function Gr(t){t.fixed&=-7}function Yr(t){t.fixed|=4,t.px=t.x,t.py=t.y}function Wr(t){t.fixed&=-5}t.interpolateTransform=Nr,t.layout={},t.layout.bundle=function(){return function(t){for(var e=[],r=-1,n=t.length;++r<n;)e.push(Vr(t[r]));return e}},t.layout.chord=function(){var e,r,n,i,a,o,s,l={},c=0;function u(){var l,u,h,p,d,m={},g=[],v=t.range(i),y=[];for(e=[],r=[],l=0,p=-1;++p<i;){for(u=0,d=-1;++d<i;)u+=n[p][d];g.push(u),y.push(t.range(i)),l+=u}for(a&&v.sort((function(t,e){return a(g[t],g[e])})),o&&y.forEach((function(t,e){t.sort((function(t,r){return o(n[e][t],n[e][r])}))})),l=(Mt-c*i)/l,u=0,p=-1;++p<i;){for(h=u,d=-1;++d<i;){var x=v[p],b=y[x][d],_=n[x][b],w=u,T=u+=_*l;m[x+"-"+b]={index:x,subindex:b,startAngle:w,endAngle:T,value:_}}r[x]={index:x,startAngle:h,endAngle:u,value:g[x]},u+=c}for(p=-1;++p<i;)for(d=p-1;++d<i;){var k=m[p+"-"+d],A=m[d+"-"+p];(k.value||A.value)&&e.push(k.value<A.value?{source:A,target:k}:{source:k,target:A})}s&&f()}function f(){e.sort((function(t,e){return s((t.source.value+t.target.value)/2,(e.source.value+e.target.value)/2)}))}return l.matrix=function(t){return arguments.length?(i=(n=t)&&n.length,e=r=null,l):n},l.padding=function(t){return arguments.length?(c=t,e=r=null,l):c},l.sortGroups=function(t){return arguments.length?(a=t,e=r=null,l):a},l.sortSubgroups=function(t){return arguments.length?(o=t,e=null,l):o},l.sortChords=function(t){return arguments.length?(s=t,e&&f(),l):s},l.chords=function(){return e||u(),e},l.groups=function(){return r||u(),r},l},t.layout.force=function(){var e,r,n,i,a,o,s={},l=t.dispatch("start","tick","end"),c=[1,1],u=.9,f=Xr,h=Zr,p=-30,d=Jr,m=.1,g=.64,v=[],y=[];function x(t){return function(e,r,n,i){if(e.point!==t){var a=e.cx-t.x,o=e.cy-t.y,s=i-r,l=a*a+o*o;if(s*s/g<l){if(l<d){var c=e.charge/l;t.px-=a*c,t.py-=o*c}return!0}if(e.point&&l&&l<d){c=e.pointCharge/l;t.px-=a*c,t.py-=o*c}}return!e.charge}}function b(e){e.px=t.event.x,e.py=t.event.y,s.resume()}return s.tick=function(){if((n*=.99)<.005)return e=null,l.end({type:"end",alpha:n=0}),!0;var r,s,f,h,d,g,b,_,w,T=v.length,k=y.length;for(s=0;s<k;++s)h=(f=y[s]).source,(g=(_=(d=f.target).x-h.x)*_+(w=d.y-h.y)*w)&&(_*=g=n*a[s]*((g=Math.sqrt(g))-i[s])/g,w*=g,d.x-=_*(b=h.weight+d.weight?h.weight/(h.weight+d.weight):.5),d.y-=w*b,h.x+=_*(b=1-b),h.y+=w*b);if((b=n*m)&&(_=c[0]/2,w=c[1]/2,s=-1,b))for(;++s<T;)(f=v[s]).x+=(_-f.x)*b,f.y+=(w-f.y)*b;if(p)for(!function t(e,r,n){var i=0,a=0;if(e.charge=0,!e.leaf)for(var o,s=e.nodes,l=s.length,c=-1;++c<l;)null!=(o=s[c])&&(t(o,r,n),e.charge+=o.charge,i+=o.charge*o.cx,a+=o.charge*o.cy);if(e.point){e.leaf||(e.point.x+=Math.random()-.5,e.point.y+=Math.random()-.5);var u=r*n[e.point.index];e.charge+=e.pointCharge=u,i+=u*e.point.x,a+=u*e.point.y}e.cx=i/e.charge,e.cy=a/e.charge}(r=t.geom.quadtree(v),n,o),s=-1;++s<T;)(f=v[s]).fixed||r.visit(x(f));for(s=-1;++s<T;)(f=v[s]).fixed?(f.x=f.px,f.y=f.py):(f.x-=(f.px-(f.px=f.x))*u,f.y-=(f.py-(f.py=f.y))*u);l.tick({type:"tick",alpha:n})},s.nodes=function(t){return arguments.length?(v=t,s):v},s.links=function(t){return arguments.length?(y=t,s):y},s.size=function(t){return arguments.length?(c=t,s):c},s.linkDistance=function(t){return arguments.length?(f="function"==typeof t?t:+t,s):f},s.distance=s.linkDistance,s.linkStrength=function(t){return arguments.length?(h="function"==typeof t?t:+t,s):h},s.friction=function(t){return arguments.length?(u=+t,s):u},s.charge=function(t){return arguments.length?(p="function"==typeof t?t:+t,s):p},s.chargeDistance=function(t){return arguments.length?(d=t*t,s):Math.sqrt(d)},s.gravity=function(t){return arguments.length?(m=+t,s):m},s.theta=function(t){return arguments.length?(g=t*t,s):Math.sqrt(g)},s.alpha=function(t){return arguments.length?(t=+t,n?t>0?n=t:(e.c=null,e.t=NaN,e=null,l.end({type:"end",alpha:n=0})):t>0&&(l.start({type:"start",alpha:n=t}),e=ve(s.tick)),s):n},s.start=function(){var t,e,r,n=v.length,l=y.length,u=c[0],d=c[1];for(t=0;t<n;++t)(r=v[t]).index=t,r.weight=0;for(t=0;t<l;++t)"number"==typeof(r=y[t]).source&&(r.source=v[r.source]),"number"==typeof r.target&&(r.target=v[r.target]),++r.source.weight,++r.target.weight;for(t=0;t<n;++t)r=v[t],isNaN(r.x)&&(r.x=m("x",u)),isNaN(r.y)&&(r.y=m("y",d)),isNaN(r.px)&&(r.px=r.x),isNaN(r.py)&&(r.py=r.y);if(i=[],"function"==typeof f)for(t=0;t<l;++t)i[t]=+f.call(this,y[t],t);else for(t=0;t<l;++t)i[t]=f;if(a=[],"function"==typeof h)for(t=0;t<l;++t)a[t]=+h.call(this,y[t],t);else for(t=0;t<l;++t)a[t]=h;if(o=[],"function"==typeof p)for(t=0;t<n;++t)o[t]=+p.call(this,v[t],t);else for(t=0;t<n;++t)o[t]=p;function m(r,i){if(!e){for(e=new Array(n),c=0;c<n;++c)e[c]=[];for(c=0;c<l;++c){var a=y[c];e[a.source.index].push(a.target),e[a.target.index].push(a.source)}}for(var o,s=e[t],c=-1,u=s.length;++c<u;)if(!isNaN(o=s[c][r]))return o;return Math.random()*i}return s.resume()},s.resume=function(){return s.alpha(.1)},s.stop=function(){return s.alpha(0)},s.drag=function(){if(r||(r=t.behavior.drag().origin(C).on("dragstart.force",qr).on("drag.force",b).on("dragend.force",Gr)),!arguments.length)return r;this.on("mouseover.force",Yr).on("mouseout.force",Wr).call(r)},t.rebind(s,l,"on")};var Xr=20,Zr=1,Jr=1/0;function Kr(e,r){return t.rebind(e,r,"sort","children","value"),e.nodes=e,e.links=nn,e}function Qr(t,e){for(var r=[t];null!=(t=r.pop());)if(e(t),(i=t.children)&&(n=i.length))for(var n,i;--n>=0;)r.push(i[n])}function $r(t,e){for(var r=[t],n=[];null!=(t=r.pop());)if(n.push(t),(a=t.children)&&(i=a.length))for(var i,a,o=-1;++o<i;)r.push(a[o]);for(;null!=(t=n.pop());)e(t)}function tn(t){return t.children}function en(t){return t.value}function rn(t,e){return e.value-t.value}function nn(e){return t.merge(e.map((function(t){return(t.children||[]).map((function(e){return{source:t,target:e}}))})))}t.layout.hierarchy=function(){var t=rn,e=tn,r=en;function n(i){var a,o=[i],s=[];for(i.depth=0;null!=(a=o.pop());)if(s.push(a),(c=e.call(n,a,a.depth))&&(l=c.length)){for(var l,c,u;--l>=0;)o.push(u=c[l]),u.parent=a,u.depth=a.depth+1;r&&(a.value=0),a.children=c}else r&&(a.value=+r.call(n,a,a.depth)||0),delete a.children;return $r(i,(function(e){var n,i;t&&(n=e.children)&&n.sort(t),r&&(i=e.parent)&&(i.value+=e.value)})),s}return n.sort=function(e){return arguments.length?(t=e,n):t},n.children=function(t){return arguments.length?(e=t,n):e},n.value=function(t){return arguments.length?(r=t,n):r},n.revalue=function(t){return r&&(Qr(t,(function(t){t.children&&(t.value=0)})),$r(t,(function(t){var e;t.children||(t.value=+r.call(n,t,t.depth)||0),(e=t.parent)&&(e.value+=t.value)}))),t},n},t.layout.partition=function(){var e=t.layout.hierarchy(),r=[1,1];function n(t,n){var i=e.call(this,t,n);return function t(e,r,n,i){var a=e.children;if(e.x=r,e.y=e.depth*i,e.dx=n,e.dy=i,a&&(o=a.length)){var o,s,l,c=-1;for(n=e.value?n/e.value:0;++c<o;)t(s=a[c],r,l=s.value*n,i),r+=l}}(i[0],0,r[0],r[1]/function t(e){var r=e.children,n=0;if(r&&(i=r.length))for(var i,a=-1;++a<i;)n=Math.max(n,t(r[a]));return 1+n}(i[0])),i}return n.size=function(t){return arguments.length?(r=t,n):r},Kr(n,e)},t.layout.pie=function(){var e=Number,r=an,n=0,i=Mt,a=0;function o(s){var l,c=s.length,u=s.map((function(t,r){return+e.call(o,t,r)})),f=+("function"==typeof n?n.apply(this,arguments):n),h=("function"==typeof i?i.apply(this,arguments):i)-f,p=Math.min(Math.abs(h)/c,+("function"==typeof a?a.apply(this,arguments):a)),d=p*(h<0?-1:1),m=t.sum(u),g=m?(h-c*d)/m:0,v=t.range(c),y=[];return null!=r&&v.sort(r===an?function(t,e){return u[e]-u[t]}:function(t,e){return r(s[t],s[e])}),v.forEach((function(t){y[t]={data:s[t],value:l=u[t],startAngle:f,endAngle:f+=l*g+d,padAngle:p}})),y}return o.value=function(t){return arguments.length?(e=t,o):e},o.sort=function(t){return arguments.length?(r=t,o):r},o.startAngle=function(t){return arguments.length?(n=t,o):n},o.endAngle=function(t){return arguments.length?(i=t,o):i},o.padAngle=function(t){return arguments.length?(a=t,o):a},o};var an={};function on(t){return t.x}function sn(t){return t.y}function ln(t,e,r){t.y0=e,t.y=r}t.layout.stack=function(){var e=C,r=fn,n=hn,i=ln,a=on,o=sn;function s(l,c){if(!(p=l.length))return l;var u=l.map((function(t,r){return e.call(s,t,r)})),f=u.map((function(t){return t.map((function(t,e){return[a.call(s,t,e),o.call(s,t,e)]}))})),h=r.call(s,f,c);u=t.permute(u,h),f=t.permute(f,h);var p,d,m,g,v=n.call(s,f,c),y=u[0].length;for(m=0;m<y;++m)for(i.call(s,u[0][m],g=v[m],f[0][m][1]),d=1;d<p;++d)i.call(s,u[d][m],g+=f[d-1][m][1],f[d][m][1]);return l}return s.values=function(t){return arguments.length?(e=t,s):e},s.order=function(t){return arguments.length?(r="function"==typeof t?t:cn.get(t)||fn,s):r},s.offset=function(t){return arguments.length?(n="function"==typeof t?t:un.get(t)||hn,s):n},s.x=function(t){return arguments.length?(a=t,s):a},s.y=function(t){return arguments.length?(o=t,s):o},s.out=function(t){return arguments.length?(i=t,s):i},s};var cn=t.map({"inside-out":function(e){var r,n,i=e.length,a=e.map(pn),o=e.map(dn),s=t.range(i).sort((function(t,e){return a[t]-a[e]})),l=0,c=0,u=[],f=[];for(r=0;r<i;++r)n=s[r],l<c?(l+=o[n],u.push(n)):(c+=o[n],f.push(n));return f.reverse().concat(u)},reverse:function(e){return t.range(e.length).reverse()},default:fn}),un=t.map({silhouette:function(t){var e,r,n,i=t.length,a=t[0].length,o=[],s=0,l=[];for(r=0;r<a;++r){for(e=0,n=0;e<i;e++)n+=t[e][r][1];n>s&&(s=n),o.push(n)}for(r=0;r<a;++r)l[r]=(s-o[r])/2;return l},wiggle:function(t){var e,r,n,i,a,o,s,l,c,u=t.length,f=t[0],h=f.length,p=[];for(p[0]=l=c=0,r=1;r<h;++r){for(e=0,i=0;e<u;++e)i+=t[e][r][1];for(e=0,a=0,s=f[r][0]-f[r-1][0];e<u;++e){for(n=0,o=(t[e][r][1]-t[e][r-1][1])/(2*s);n<e;++n)o+=(t[n][r][1]-t[n][r-1][1])/s;a+=o*t[e][r][1]}p[r]=l-=i?a/i*s:0,l<c&&(c=l)}for(r=0;r<h;++r)p[r]-=c;return p},expand:function(t){var e,r,n,i=t.length,a=t[0].length,o=1/i,s=[];for(r=0;r<a;++r){for(e=0,n=0;e<i;e++)n+=t[e][r][1];if(n)for(e=0;e<i;e++)t[e][r][1]/=n;else for(e=0;e<i;e++)t[e][r][1]=o}for(r=0;r<a;++r)s[r]=0;return s},zero:hn});function fn(e){return t.range(e.length)}function hn(t){for(var e=-1,r=t[0].length,n=[];++e<r;)n[e]=0;return n}function pn(t){for(var e,r=1,n=0,i=t[0][1],a=t.length;r<a;++r)(e=t[r][1])>i&&(n=r,i=e);return n}function dn(t){return t.reduce(mn,0)}function mn(t,e){return t+e[1]}function gn(t,e){return vn(t,Math.ceil(Math.log(e.length)/Math.LN2+1))}function vn(t,e){for(var r=-1,n=+t[0],i=(t[1]-n)/e,a=[];++r<=e;)a[r]=i*r+n;return a}function yn(e){return[t.min(e),t.max(e)]}function xn(t,e){return t.value-e.value}function bn(t,e){var r=t._pack_next;t._pack_next=e,e._pack_prev=t,e._pack_next=r,r._pack_prev=e}function _n(t,e){t._pack_next=e,e._pack_prev=t}function wn(t,e){var r=e.x-t.x,n=e.y-t.y,i=t.r+e.r;return.999*i*i>r*r+n*n}function Tn(t){if((e=t.children)&&(l=e.length)){var e,r,n,i,a,o,s,l,c=1/0,u=-1/0,f=1/0,h=-1/0;if(e.forEach(kn),(r=e[0]).x=-r.r,r.y=0,x(r),l>1&&((n=e[1]).x=n.r,n.y=0,x(n),l>2))for(Mn(r,n,i=e[2]),x(i),bn(r,i),r._pack_prev=i,bn(i,n),n=r._pack_next,a=3;a<l;a++){Mn(r,n,i=e[a]);var p=0,d=1,m=1;for(o=n._pack_next;o!==n;o=o._pack_next,d++)if(wn(o,i)){p=1;break}if(1==p)for(s=r._pack_prev;s!==o._pack_prev&&!wn(s,i);s=s._pack_prev,m++);p?(d<m||d==m&&n.r<r.r?_n(r,n=o):_n(r=s,n),a--):(bn(r,i),n=i,x(i))}var g=(c+u)/2,v=(f+h)/2,y=0;for(a=0;a<l;a++)(i=e[a]).x-=g,i.y-=v,y=Math.max(y,i.r+Math.sqrt(i.x*i.x+i.y*i.y));t.r=y,e.forEach(An)}function x(t){c=Math.min(t.x-t.r,c),u=Math.max(t.x+t.r,u),f=Math.min(t.y-t.r,f),h=Math.max(t.y+t.r,h)}}function kn(t){t._pack_next=t._pack_prev=t}function An(t){delete t._pack_next,delete t._pack_prev}function Mn(t,e,r){var n=t.r+r.r,i=e.x-t.x,a=e.y-t.y;if(n&&(i||a)){var o=e.r+r.r,s=i*i+a*a,l=.5+((n*=n)-(o*=o))/(2*s),c=Math.sqrt(Math.max(0,2*o*(n+s)-(n-=s)*n-o*o))/(2*s);r.x=t.x+l*i+c*a,r.y=t.y+l*a-c*i}else r.x=t.x+n,r.y=t.y}function Sn(t,e){return t.parent==e.parent?1:2}function En(t){var e=t.children;return e.length?e[0]:t.t}function Ln(t){var e,r=t.children;return(e=r.length)?r[e-1]:t.t}function Cn(t,e,r){var n=r/(e.i-t.i);e.c-=n,e.s+=r,t.c+=n,e.z+=r,e.m+=r}function Pn(t,e,r){return t.a.parent===e.parent?t.a:r}function In(t){return{x:t.x,y:t.y,dx:t.dx,dy:t.dy}}function On(t,e){var r=t.x+e[3],n=t.y+e[0],i=t.dx-e[1]-e[3],a=t.dy-e[0]-e[2];return i<0&&(r+=i/2,i=0),a<0&&(n+=a/2,a=0),{x:r,y:n,dx:i,dy:a}}function zn(t){var e=t[0],r=t[t.length-1];return e<r?[e,r]:[r,e]}function Dn(t){return t.rangeExtent?t.rangeExtent():zn(t.range())}function Rn(t,e,r,n){var i=r(t[0],t[1]),a=n(e[0],e[1]);return function(t){return a(i(t))}}function Fn(t,e){var r,n=0,i=t.length-1,a=t[n],o=t[i];return o<a&&(r=n,n=i,i=r,r=a,a=o,o=r),t[n]=e.floor(a),t[i]=e.ceil(o),t}function Bn(t){return t?{floor:function(e){return Math.floor(e/t)*t},ceil:function(e){return Math.ceil(e/t)*t}}:Nn}t.layout.histogram=function(){var e=!0,r=Number,n=yn,i=gn;function a(a,o){for(var s,l,c=[],u=a.map(r,this),f=n.call(this,u,o),h=i.call(this,f,u,o),p=(o=-1,u.length),d=h.length-1,m=e?1:1/p;++o<d;)(s=c[o]=[]).dx=h[o+1]-(s.x=h[o]),s.y=0;if(d>0)for(o=-1;++o<p;)(l=u[o])>=f[0]&&l<=f[1]&&((s=c[t.bisect(h,l,1,d)-1]).y+=m,s.push(a[o]));return c}return a.value=function(t){return arguments.length?(r=t,a):r},a.range=function(t){return arguments.length?(n=ce(t),a):n},a.bins=function(t){return arguments.length?(i="number"==typeof t?function(e){return vn(e,t)}:ce(t),a):i},a.frequency=function(t){return arguments.length?(e=!!t,a):e},a},t.layout.pack=function(){var e,r=t.layout.hierarchy().sort(xn),n=0,i=[1,1];function a(t,a){var o=r.call(this,t,a),s=o[0],l=i[0],c=i[1],u=null==e?Math.sqrt:"function"==typeof e?e:function(){return e};if(s.x=s.y=0,$r(s,(function(t){t.r=+u(t.value)})),$r(s,Tn),n){var f=n*(e?1:Math.max(2*s.r/l,2*s.r/c))/2;$r(s,(function(t){t.r+=f})),$r(s,Tn),$r(s,(function(t){t.r-=f}))}return function t(e,r,n,i){var a=e.children;if(e.x=r+=i*e.x,e.y=n+=i*e.y,e.r*=i,a)for(var o=-1,s=a.length;++o<s;)t(a[o],r,n,i)}(s,l/2,c/2,e?1:1/Math.max(2*s.r/l,2*s.r/c)),o}return a.size=function(t){return arguments.length?(i=t,a):i},a.radius=function(t){return arguments.length?(e=null==t||"function"==typeof t?t:+t,a):e},a.padding=function(t){return arguments.length?(n=+t,a):n},Kr(a,r)},t.layout.tree=function(){var e=t.layout.hierarchy().sort(null).value(null),r=Sn,n=[1,1],i=null;function a(t,a){var c=e.call(this,t,a),u=c[0],f=function(t){var e,r={A:null,children:[t]},n=[r];for(;null!=(e=n.pop());)for(var i,a=e.children,o=0,s=a.length;o<s;++o)n.push((a[o]=i={_:a[o],parent:e,children:(i=a[o].children)&&i.slice()||[],A:null,a:null,z:0,m:0,c:0,s:0,t:null,i:o}).a=i);return r.children[0]}(u);if($r(f,o),f.parent.m=-f.z,Qr(f,s),i)Qr(u,l);else{var h=u,p=u,d=u;Qr(u,(function(t){t.x<h.x&&(h=t),t.x>p.x&&(p=t),t.depth>d.depth&&(d=t)}));var m=r(h,p)/2-h.x,g=n[0]/(p.x+r(p,h)/2+m),v=n[1]/(d.depth||1);Qr(u,(function(t){t.x=(t.x+m)*g,t.y=t.depth*v}))}return c}function o(t){var e=t.children,n=t.parent.children,i=t.i?n[t.i-1]:null;if(e.length){!function(t){var e,r=0,n=0,i=t.children,a=i.length;for(;--a>=0;)(e=i[a]).z+=r,e.m+=r,r+=e.s+(n+=e.c)}(t);var a=(e[0].z+e[e.length-1].z)/2;i?(t.z=i.z+r(t._,i._),t.m=t.z-a):t.z=a}else i&&(t.z=i.z+r(t._,i._));t.parent.A=function(t,e,n){if(e){for(var i,a=t,o=t,s=e,l=a.parent.children[0],c=a.m,u=o.m,f=s.m,h=l.m;s=Ln(s),a=En(a),s&&a;)l=En(l),(o=Ln(o)).a=t,(i=s.z+f-a.z-c+r(s._,a._))>0&&(Cn(Pn(s,t,n),t,i),c+=i,u+=i),f+=s.m,c+=a.m,h+=l.m,u+=o.m;s&&!Ln(o)&&(o.t=s,o.m+=f-u),a&&!En(l)&&(l.t=a,l.m+=c-h,n=t)}return n}(t,i,t.parent.A||n[0])}function s(t){t._.x=t.z+t.parent.m,t.m+=t.parent.m}function l(t){t.x*=n[0],t.y=t.depth*n[1]}return a.separation=function(t){return arguments.length?(r=t,a):r},a.size=function(t){return arguments.length?(i=null==(n=t)?l:null,a):i?null:n},a.nodeSize=function(t){return arguments.length?(i=null==(n=t)?null:l,a):i?n:null},Kr(a,e)},t.layout.cluster=function(){var e=t.layout.hierarchy().sort(null).value(null),r=Sn,n=[1,1],i=!1;function a(a,o){var s,l=e.call(this,a,o),c=l[0],u=0;$r(c,(function(e){var n=e.children;n&&n.length?(e.x=function(t){return t.reduce((function(t,e){return t+e.x}),0)/t.length}(n),e.y=function(e){return 1+t.max(e,(function(t){return t.y}))}(n)):(e.x=s?u+=r(e,s):0,e.y=0,s=e)}));var f=function t(e){var r=e.children;return r&&r.length?t(r[0]):e}(c),h=function t(e){var r,n=e.children;return n&&(r=n.length)?t(n[r-1]):e}(c),p=f.x-r(f,h)/2,d=h.x+r(h,f)/2;return $r(c,i?function(t){t.x=(t.x-c.x)*n[0],t.y=(c.y-t.y)*n[1]}:function(t){t.x=(t.x-p)/(d-p)*n[0],t.y=(1-(c.y?t.y/c.y:1))*n[1]}),l}return a.separation=function(t){return arguments.length?(r=t,a):r},a.size=function(t){return arguments.length?(i=null==(n=t),a):i?null:n},a.nodeSize=function(t){return arguments.length?(i=null!=(n=t),a):i?n:null},Kr(a,e)},t.layout.treemap=function(){var e,r=t.layout.hierarchy(),n=Math.round,i=[1,1],a=null,o=In,s=!1,l="squarify",c=.5*(1+Math.sqrt(5));function u(t,e){for(var r,n,i=-1,a=t.length;++i<a;)n=(r=t[i]).value*(e<0?0:e),r.area=isNaN(n)||n<=0?0:n}function f(t){var e=t.children;if(e&&e.length){var r,n,i,a=o(t),s=[],c=e.slice(),h=1/0,m="slice"===l?a.dx:"dice"===l?a.dy:"slice-dice"===l?1&t.depth?a.dy:a.dx:Math.min(a.dx,a.dy);for(u(c,a.dx*a.dy/t.value),s.area=0;(i=c.length)>0;)s.push(r=c[i-1]),s.area+=r.area,"squarify"!==l||(n=p(s,m))<=h?(c.pop(),h=n):(s.area-=s.pop().area,d(s,m,a,!1),m=Math.min(a.dx,a.dy),s.length=s.area=0,h=1/0);s.length&&(d(s,m,a,!0),s.length=s.area=0),e.forEach(f)}}function h(t){var e=t.children;if(e&&e.length){var r,n=o(t),i=e.slice(),a=[];for(u(i,n.dx*n.dy/t.value),a.area=0;r=i.pop();)a.push(r),a.area+=r.area,null!=r.z&&(d(a,r.z?n.dx:n.dy,n,!i.length),a.length=a.area=0);e.forEach(h)}}function p(t,e){for(var r,n=t.area,i=0,a=1/0,o=-1,s=t.length;++o<s;)(r=t[o].area)&&(r<a&&(a=r),r>i&&(i=r));return e*=e,(n*=n)?Math.max(e*i*c/n,n/(e*a*c)):1/0}function d(t,e,r,i){var a,o=-1,s=t.length,l=r.x,c=r.y,u=e?n(t.area/e):0;if(e==r.dx){for((i||u>r.dy)&&(u=r.dy);++o<s;)(a=t[o]).x=l,a.y=c,a.dy=u,l+=a.dx=Math.min(r.x+r.dx-l,u?n(a.area/u):0);a.z=!0,a.dx+=r.x+r.dx-l,r.y+=u,r.dy-=u}else{for((i||u>r.dx)&&(u=r.dx);++o<s;)(a=t[o]).x=l,a.y=c,a.dx=u,c+=a.dy=Math.min(r.y+r.dy-c,u?n(a.area/u):0);a.z=!1,a.dy+=r.y+r.dy-c,r.x+=u,r.dx-=u}}function m(t){var n=e||r(t),a=n[0];return a.x=a.y=0,a.value?(a.dx=i[0],a.dy=i[1]):a.dx=a.dy=0,e&&r.revalue(a),u([a],a.dx*a.dy/a.value),(e?h:f)(a),s&&(e=n),n}return m.size=function(t){return arguments.length?(i=t,m):i},m.padding=function(t){if(!arguments.length)return a;function e(e){var r=t.call(m,e,e.depth);return null==r?In(e):On(e,"number"==typeof r?[r,r,r,r]:r)}function r(e){return On(e,t)}var n;return o=null==(a=t)?In:"function"==(n=typeof t)?e:"number"===n?(t=[t,t,t,t],r):r,m},m.round=function(t){return arguments.length?(n=t?Math.round:Number,m):n!=Number},m.sticky=function(t){return arguments.length?(s=t,e=null,m):s},m.ratio=function(t){return arguments.length?(c=t,m):c},m.mode=function(t){return arguments.length?(l=t+"",m):l},Kr(m,r)},t.random={normal:function(t,e){var r=arguments.length;return r<2&&(e=1),r<1&&(t=0),function(){var r,n,i;do{i=(r=2*Math.random()-1)*r+(n=2*Math.random()-1)*n}while(!i||i>1);return t+e*r*Math.sqrt(-2*Math.log(i)/i)}},logNormal:function(){var e=t.random.normal.apply(t,arguments);return function(){return Math.exp(e())}},bates:function(e){var r=t.random.irwinHall(e);return function(){return r()/e}},irwinHall:function(t){return function(){for(var e=0,r=0;r<t;r++)e+=Math.random();return e}}},t.scale={};var Nn={floor:C,ceil:C};function jn(e,r,n,i){var a=[],o=[],s=0,l=Math.min(e.length,r.length)-1;for(e[l]<e[0]&&(e=e.slice().reverse(),r=r.slice().reverse());++s<=l;)a.push(n(e[s-1],e[s])),o.push(i(r[s-1],r[s]));return function(r){var n=t.bisect(e,r,1,l)-1;return o[n](a[n](r))}}function Un(e,r){return t.rebind(e,r,"range","rangeRound","interpolate","clamp")}function Vn(t,e){return Fn(t,Bn(Hn(t,e)[2])),Fn(t,Bn(Hn(t,e)[2])),t}function Hn(t,e){null==e&&(e=10);var r=zn(t),n=r[1]-r[0],i=Math.pow(10,Math.floor(Math.log(n/e)/Math.LN10)),a=e/n*i;return a<=.15?i*=10:a<=.35?i*=5:a<=.75&&(i*=2),r[0]=Math.ceil(r[0]/i)*i,r[1]=Math.floor(r[1]/i)*i+.5*i,r[2]=i,r}function qn(e,r){return t.range.apply(t,Hn(e,r))}t.scale.linear=function(){return function t(e,r,n,i){var a,o;function s(){var t=Math.min(e.length,r.length)>2?jn:Rn,s=i?Ur:jr;return a=t(e,r,s,n),o=t(r,e,s,yr),l}function l(t){return a(t)}return l.invert=function(t){return o(t)},l.domain=function(t){return arguments.length?(e=t.map(Number),s()):e},l.range=function(t){return arguments.length?(r=t,s()):r},l.rangeRound=function(t){return l.range(t).interpolate(Or)},l.clamp=function(t){return arguments.length?(i=t,s()):i},l.interpolate=function(t){return arguments.length?(n=t,s()):n},l.ticks=function(t){return qn(e,t)},l.tickFormat=function(t,r){return d3_scale_linearTickFormat(e,t,r)},l.nice=function(t){return Vn(e,t),s()},l.copy=function(){return t(e,r,n,i)},s()}([0,1],[0,1],yr,!1)};t.scale.log=function(){return function t(e,r,n,i){function a(t){return(n?Math.log(t<0?0:t):-Math.log(t>0?0:-t))/Math.log(r)}function o(t){return n?Math.pow(r,t):-Math.pow(r,-t)}function s(t){return e(a(t))}return s.invert=function(t){return o(e.invert(t))},s.domain=function(t){return arguments.length?(n=t[0]>=0,e.domain((i=t.map(Number)).map(a)),s):i},s.base=function(t){return arguments.length?(r=+t,e.domain(i.map(a)),s):r},s.nice=function(){var t=Fn(i.map(a),n?Math:Gn);return e.domain(t),i=t.map(o),s},s.ticks=function(){var t=zn(i),e=[],s=t[0],l=t[1],c=Math.floor(a(s)),u=Math.ceil(a(l)),f=r%1?2:r;if(isFinite(u-c)){if(n){for(;c<u;c++)for(var h=1;h<f;h++)e.push(o(c)*h);e.push(o(c))}else for(e.push(o(c));c++<u;)for(h=f-1;h>0;h--)e.push(o(c)*h);for(c=0;e[c]<s;c++);for(u=e.length;e[u-1]>l;u--);e=e.slice(c,u)}return e},s.copy=function(){return t(e.copy(),r,n,i)},Un(s,e)}(t.scale.linear().domain([0,1]),10,!0,[1,10])};var Gn={floor:function(t){return-Math.ceil(-t)},ceil:function(t){return-Math.floor(-t)}};function Yn(t){return function(e){return e<0?-Math.pow(-e,t):Math.pow(e,t)}}t.scale.pow=function(){return function t(e,r,n){var i=Yn(r),a=Yn(1/r);function o(t){return e(i(t))}return o.invert=function(t){return a(e.invert(t))},o.domain=function(t){return arguments.length?(e.domain((n=t.map(Number)).map(i)),o):n},o.ticks=function(t){return qn(n,t)},o.tickFormat=function(t,e){return d3_scale_linearTickFormat(n,t,e)},o.nice=function(t){return o.domain(Vn(n,t))},o.exponent=function(t){return arguments.length?(i=Yn(r=t),a=Yn(1/r),e.domain(n.map(i)),o):r},o.copy=function(){return t(e.copy(),r,n)},Un(o,e)}(t.scale.linear(),1,[0,1])},t.scale.sqrt=function(){return t.scale.pow().exponent(.5)},t.scale.ordinal=function(){return function e(r,n){var i,a,o;function s(t){return a[((i.get(t)||("range"===n.t?i.set(t,r.push(t)):NaN))-1)%a.length]}function l(e,n){return t.range(r.length).map((function(t){return e+n*t}))}return s.domain=function(t){if(!arguments.length)return r;r=[],i=new _;for(var e,a=-1,o=t.length;++a<o;)i.has(e=t[a])||i.set(e,r.push(e));return s[n.t].apply(s,n.a)},s.range=function(t){return arguments.length?(a=t,o=0,n={t:"range",a:arguments},s):a},s.rangePoints=function(t,e){arguments.length<2&&(e=0);var i=t[0],c=t[1],u=r.length<2?(i=(i+c)/2,0):(c-i)/(r.length-1+e);return a=l(i+u*e/2,u),o=0,n={t:"rangePoints",a:arguments},s},s.rangeRoundPoints=function(t,e){arguments.length<2&&(e=0);var i=t[0],c=t[1],u=r.length<2?(i=c=Math.round((i+c)/2),0):(c-i)/(r.length-1+e)|0;return a=l(i+Math.round(u*e/2+(c-i-(r.length-1+e)*u)/2),u),o=0,n={t:"rangeRoundPoints",a:arguments},s},s.rangeBands=function(t,e,i){arguments.length<2&&(e=0),arguments.length<3&&(i=e);var c=t[1]<t[0],u=t[c-0],f=t[1-c],h=(f-u)/(r.length-e+2*i);return a=l(u+h*i,h),c&&a.reverse(),o=h*(1-e),n={t:"rangeBands",a:arguments},s},s.rangeRoundBands=function(t,e,i){arguments.length<2&&(e=0),arguments.length<3&&(i=e);var c=t[1]<t[0],u=t[c-0],f=t[1-c],h=Math.floor((f-u)/(r.length-e+2*i));return a=l(u+Math.round((f-u-(r.length-e)*h)/2),h),c&&a.reverse(),o=Math.round(h*(1-e)),n={t:"rangeRoundBands",a:arguments},s},s.rangeBand=function(){return o},s.rangeExtent=function(){return zn(n.a[0])},s.copy=function(){return e(r,n)},s.domain(r)}([],{t:"range",a:[[]]})},t.scale.category10=function(){return t.scale.ordinal().range(Wn)},t.scale.category20=function(){return t.scale.ordinal().range(Xn)},t.scale.category20b=function(){return t.scale.ordinal().range(Zn)},t.scale.category20c=function(){return t.scale.ordinal().range(Jn)};var Wn=[2062260,16744206,2924588,14034728,9725885,9197131,14907330,8355711,12369186,1556175].map(te),Xn=[2062260,11454440,16744206,16759672,2924588,10018698,14034728,16750742,9725885,12955861,9197131,12885140,14907330,16234194,8355711,13092807,12369186,14408589,1556175,10410725].map(te),Zn=[3750777,5395619,7040719,10264286,6519097,9216594,11915115,13556636,9202993,12426809,15186514,15190932,8666169,11356490,14049643,15177372,8077683,10834324,13528509,14589654].map(te),Jn=[3244733,7057110,10406625,13032431,15095053,16616764,16625259,16634018,3253076,7652470,10607003,13101504,7695281,10394312,12369372,14342891,6513507,9868950,12434877,14277081].map(te);function Kn(){return 0}t.scale.quantile=function(){return function e(r,n){var i;function a(){var e=0,a=n.length;for(i=[];++e<a;)i[e-1]=t.quantile(r,e/a);return o}function o(e){if(!isNaN(e=+e))return n[t.bisect(i,e)]}return o.domain=function(t){return arguments.length?(r=t.map(p).filter(d).sort(h),a()):r},o.range=function(t){return arguments.length?(n=t,a()):n},o.quantiles=function(){return i},o.invertExtent=function(t){return(t=n.indexOf(t))<0?[NaN,NaN]:[t>0?i[t-1]:r[0],t<i.length?i[t]:r[r.length-1]]},o.copy=function(){return e(r,n)},a()}([],[])},t.scale.quantize=function(){return function t(e,r,n){var i,a;function o(t){return n[Math.max(0,Math.min(a,Math.floor(i*(t-e))))]}function s(){return i=n.length/(r-e),a=n.length-1,o}return o.domain=function(t){return arguments.length?(e=+t[0],r=+t[t.length-1],s()):[e,r]},o.range=function(t){return arguments.length?(n=t,s()):n},o.invertExtent=function(t){return[t=(t=n.indexOf(t))<0?NaN:t/i+e,t+1/i]},o.copy=function(){return t(e,r,n)},s()}(0,1,[0,1])},t.scale.threshold=function(){return function e(r,n){function i(e){if(e<=e)return n[t.bisect(r,e)]}return i.domain=function(t){return arguments.length?(r=t,i):r},i.range=function(t){return arguments.length?(n=t,i):n},i.invertExtent=function(t){return t=n.indexOf(t),[r[t-1],r[t]]},i.copy=function(){return e(r,n)},i}([.5],[0,1])},t.scale.identity=function(){return function t(e){function r(t){return+t}return r.invert=r,r.domain=r.range=function(t){return arguments.length?(e=t.map(r),r):e},r.ticks=function(t){return qn(e,t)},r.tickFormat=function(t,r){return d3_scale_linearTickFormat(e,t,r)},r.copy=function(){return t(e)},r}([0,1])},t.svg={},t.svg.arc=function(){var t=$n,e=ti,r=Kn,n=Qn,i=ei,a=ri,o=ni;function s(){var s=Math.max(0,+t.apply(this,arguments)),c=Math.max(0,+e.apply(this,arguments)),u=i.apply(this,arguments)-Et,f=a.apply(this,arguments)-Et,h=Math.abs(f-u),p=u>f?0:1;if(c<s&&(d=c,c=s,s=d),h>=St)return l(c,p)+(s?l(s,1-p):"")+"Z";var d,m,g,v,y,x,b,_,w,T,k,A,M=0,S=0,E=[];if((v=(+o.apply(this,arguments)||0)/2)&&(g=n===Qn?Math.sqrt(s*s+c*c):+n.apply(this,arguments),p||(S*=-1),c&&(S=Pt(g/c*Math.sin(v))),s&&(M=Pt(g/s*Math.sin(v)))),c){y=c*Math.cos(u+S),x=c*Math.sin(u+S),b=c*Math.cos(f-S),_=c*Math.sin(f-S);var L=Math.abs(f-u-2*S)<=At?0:1;if(S&&ii(y,x,b,_)===p^L){var C=(u+f)/2;y=c*Math.cos(C),x=c*Math.sin(C),b=_=null}}else y=x=0;if(s){w=s*Math.cos(f-M),T=s*Math.sin(f-M),k=s*Math.cos(u+M),A=s*Math.sin(u+M);var P=Math.abs(u-f+2*M)<=At?0:1;if(M&&ii(w,T,k,A)===1-p^P){var I=(u+f)/2;w=s*Math.cos(I),T=s*Math.sin(I),k=A=null}}else w=T=0;if(h>kt&&(d=Math.min(Math.abs(c-s)/2,+r.apply(this,arguments)))>.001){m=s<c^p?0:1;var O=d,z=d;if(h<At){var D=null==k?[w,T]:null==b?[y,x]:Se([y,x],[k,A],[b,_],[w,T]),R=y-D[0],F=x-D[1],B=b-D[0],N=_-D[1],j=1/Math.sin(Math.acos((R*B+F*N)/(Math.sqrt(R*R+F*F)*Math.sqrt(B*B+N*N)))/2),U=Math.sqrt(D[0]*D[0]+D[1]*D[1]);z=Math.min(d,(s-U)/(j-1)),O=Math.min(d,(c-U)/(j+1))}if(null!=b){var V=ai(null==k?[w,T]:[k,A],[y,x],c,O,p),H=ai([b,_],[w,T],c,O,p);d===O?E.push("M",V[0],"A",O,",",O," 0 0,",m," ",V[1],"A",c,",",c," 0 ",1-p^ii(V[1][0],V[1][1],H[1][0],H[1][1]),",",p," ",H[1],"A",O,",",O," 0 0,",m," ",H[0]):E.push("M",V[0],"A",O,",",O," 0 1,",m," ",H[0])}else E.push("M",y,",",x);if(null!=k){var q=ai([y,x],[k,A],s,-z,p),G=ai([w,T],null==b?[y,x]:[b,_],s,-z,p);d===z?E.push("L",G[0],"A",z,",",z," 0 0,",m," ",G[1],"A",s,",",s," 0 ",p^ii(G[1][0],G[1][1],q[1][0],q[1][1]),",",1-p," ",q[1],"A",z,",",z," 0 0,",m," ",q[0]):E.push("L",G[0],"A",z,",",z," 0 0,",m," ",q[0])}else E.push("L",w,",",T)}else E.push("M",y,",",x),null!=b&&E.push("A",c,",",c," 0 ",L,",",p," ",b,",",_),E.push("L",w,",",T),null!=k&&E.push("A",s,",",s," 0 ",P,",",1-p," ",k,",",A);return E.push("Z"),E.join("")}function l(t,e){return"M0,"+t+"A"+t+","+t+" 0 1,"+e+" 0,"+-t+"A"+t+","+t+" 0 1,"+e+" 0,"+t}return s.innerRadius=function(e){return arguments.length?(t=ce(e),s):t},s.outerRadius=function(t){return arguments.length?(e=ce(t),s):e},s.cornerRadius=function(t){return arguments.length?(r=ce(t),s):r},s.padRadius=function(t){return arguments.length?(n=t==Qn?Qn:ce(t),s):n},s.startAngle=function(t){return arguments.length?(i=ce(t),s):i},s.endAngle=function(t){return arguments.length?(a=ce(t),s):a},s.padAngle=function(t){return arguments.length?(o=ce(t),s):o},s.centroid=function(){var r=(+t.apply(this,arguments)+ +e.apply(this,arguments))/2,n=(+i.apply(this,arguments)+ +a.apply(this,arguments))/2-Et;return[Math.cos(n)*r,Math.sin(n)*r]},s};var Qn="auto";function $n(t){return t.innerRadius}function ti(t){return t.outerRadius}function ei(t){return t.startAngle}function ri(t){return t.endAngle}function ni(t){return t&&t.padAngle}function ii(t,e,r,n){return(t-r)*e-(e-n)*t>0?0:1}function ai(t,e,r,n,i){var a=t[0]-e[0],o=t[1]-e[1],s=(i?n:-n)/Math.sqrt(a*a+o*o),l=s*o,c=-s*a,u=t[0]+l,f=t[1]+c,h=e[0]+l,p=e[1]+c,d=(u+h)/2,m=(f+p)/2,g=h-u,v=p-f,y=g*g+v*v,x=r-n,b=u*p-h*f,_=(v<0?-1:1)*Math.sqrt(Math.max(0,x*x*y-b*b)),w=(b*v-g*_)/y,T=(-b*g-v*_)/y,k=(b*v+g*_)/y,A=(-b*g+v*_)/y,M=w-d,S=T-m,E=k-d,L=A-m;return M*M+S*S>E*E+L*L&&(w=k,T=A),[[w-l,T-c],[w*r/x,T*r/x]]}function oi(){return!0}function si(t){var e=_e,r=we,n=oi,i=ci,a=i.key,o=.7;function s(a){var s,l=[],c=[],u=-1,f=a.length,h=ce(e),p=ce(r);function d(){l.push("M",i(t(c),o))}for(;++u<f;)n.call(this,s=a[u],u)?c.push([+h.call(this,s,u),+p.call(this,s,u)]):c.length&&(d(),c=[]);return c.length&&d(),l.length?l.join(""):null}return s.x=function(t){return arguments.length?(e=t,s):e},s.y=function(t){return arguments.length?(r=t,s):r},s.defined=function(t){return arguments.length?(n=t,s):n},s.interpolate=function(t){return arguments.length?(a="function"==typeof t?i=t:(i=li.get(t)||ci).key,s):a},s.tension=function(t){return arguments.length?(o=t,s):o},s}t.svg.line=function(){return si(C)};var li=t.map({linear:ci,"linear-closed":ui,step:function(t){var e=0,r=t.length,n=t[0],i=[n[0],",",n[1]];for(;++e<r;)i.push("H",(n[0]+(n=t[e])[0])/2,"V",n[1]);r>1&&i.push("H",n[0]);return i.join("")},"step-before":fi,"step-after":hi,basis:mi,"basis-open":function(t){if(t.length<4)return ci(t);var e,r=[],n=-1,i=t.length,a=[0],o=[0];for(;++n<3;)e=t[n],a.push(e[0]),o.push(e[1]);r.push(gi(xi,a)+","+gi(xi,o)),--n;for(;++n<i;)e=t[n],a.shift(),a.push(e[0]),o.shift(),o.push(e[1]),bi(r,a,o);return r.join("")},"basis-closed":function(t){var e,r,n=-1,i=t.length,a=i+4,o=[],s=[];for(;++n<4;)r=t[n%i],o.push(r[0]),s.push(r[1]);e=[gi(xi,o),",",gi(xi,s)],--n;for(;++n<a;)r=t[n%i],o.shift(),o.push(r[0]),s.shift(),s.push(r[1]),bi(e,o,s);return e.join("")},bundle:function(t,e){var r=t.length-1;if(r)for(var n,i,a=t[0][0],o=t[0][1],s=t[r][0]-a,l=t[r][1]-o,c=-1;++c<=r;)n=t[c],i=c/r,n[0]=e*n[0]+(1-e)*(a+i*s),n[1]=e*n[1]+(1-e)*(o+i*l);return mi(t)},cardinal:function(t,e){return t.length<3?ci(t):t[0]+pi(t,di(t,e))},"cardinal-open":function(t,e){return t.length<4?ci(t):t[1]+pi(t.slice(1,-1),di(t,e))},"cardinal-closed":function(t,e){return t.length<3?ui(t):t[0]+pi((t.push(t[0]),t),di([t[t.length-2]].concat(t,[t[1]]),e))},monotone:function(t){return t.length<3?ci(t):t[0]+pi(t,function(t){var e,r,n,i,a=[],o=function(t){var e=0,r=t.length-1,n=[],i=t[0],a=t[1],o=n[0]=_i(i,a);for(;++e<r;)n[e]=(o+(o=_i(i=a,a=t[e+1])))/2;return n[e]=o,n}(t),s=-1,l=t.length-1;for(;++s<l;)e=_i(t[s],t[s+1]),y(e)<kt?o[s]=o[s+1]=0:(r=o[s]/e,n=o[s+1]/e,(i=r*r+n*n)>9&&(i=3*e/Math.sqrt(i),o[s]=i*r,o[s+1]=i*n));s=-1;for(;++s<=l;)i=(t[Math.min(l,s+1)][0]-t[Math.max(0,s-1)][0])/(6*(1+o[s]*o[s])),a.push([i||0,o[s]*i||0]);return a}(t))}});function ci(t){return t.length>1?t.join("L"):t+"Z"}function ui(t){return t.join("L")+"Z"}function fi(t){for(var e=0,r=t.length,n=t[0],i=[n[0],",",n[1]];++e<r;)i.push("V",(n=t[e])[1],"H",n[0]);return i.join("")}function hi(t){for(var e=0,r=t.length,n=t[0],i=[n[0],",",n[1]];++e<r;)i.push("H",(n=t[e])[0],"V",n[1]);return i.join("")}function pi(t,e){if(e.length<1||t.length!=e.length&&t.length!=e.length+2)return ci(t);var r=t.length!=e.length,n="",i=t[0],a=t[1],o=e[0],s=o,l=1;if(r&&(n+="Q"+(a[0]-2*o[0]/3)+","+(a[1]-2*o[1]/3)+","+a[0]+","+a[1],i=t[1],l=2),e.length>1){s=e[1],a=t[l],l++,n+="C"+(i[0]+o[0])+","+(i[1]+o[1])+","+(a[0]-s[0])+","+(a[1]-s[1])+","+a[0]+","+a[1];for(var c=2;c<e.length;c++,l++)a=t[l],s=e[c],n+="S"+(a[0]-s[0])+","+(a[1]-s[1])+","+a[0]+","+a[1]}if(r){var u=t[l];n+="Q"+(a[0]+2*s[0]/3)+","+(a[1]+2*s[1]/3)+","+u[0]+","+u[1]}return n}function di(t,e){for(var r,n=[],i=(1-e)/2,a=t[0],o=t[1],s=1,l=t.length;++s<l;)r=a,a=o,o=t[s],n.push([i*(o[0]-r[0]),i*(o[1]-r[1])]);return n}function mi(t){if(t.length<3)return ci(t);var e=1,r=t.length,n=t[0],i=n[0],a=n[1],o=[i,i,i,(n=t[1])[0]],s=[a,a,a,n[1]],l=[i,",",a,"L",gi(xi,o),",",gi(xi,s)];for(t.push(t[r-1]);++e<=r;)n=t[e],o.shift(),o.push(n[0]),s.shift(),s.push(n[1]),bi(l,o,s);return t.pop(),l.push("L",n),l.join("")}function gi(t,e){return t[0]*e[0]+t[1]*e[1]+t[2]*e[2]+t[3]*e[3]}li.forEach((function(t,e){e.key=t,e.closed=/-closed$/.test(t)}));var vi=[0,2/3,1/3,0],yi=[0,1/3,2/3,0],xi=[0,1/6,2/3,1/6];function bi(t,e,r){t.push("C",gi(vi,e),",",gi(vi,r),",",gi(yi,e),",",gi(yi,r),",",gi(xi,e),",",gi(xi,r))}function _i(t,e){return(e[1]-t[1])/(e[0]-t[0])}function wi(t){for(var e,r,n,i=-1,a=t.length;++i<a;)r=(e=t[i])[0],n=e[1]-Et,e[0]=r*Math.cos(n),e[1]=r*Math.sin(n);return t}function Ti(t){var e=_e,r=_e,n=0,i=we,a=oi,o=ci,s=o.key,l=o,c="L",u=.7;function f(s){var f,h,p,d=[],m=[],g=[],v=-1,y=s.length,x=ce(e),b=ce(n),_=e===r?function(){return h}:ce(r),w=n===i?function(){return p}:ce(i);function T(){d.push("M",o(t(g),u),c,l(t(m.reverse()),u),"Z")}for(;++v<y;)a.call(this,f=s[v],v)?(m.push([h=+x.call(this,f,v),p=+b.call(this,f,v)]),g.push([+_.call(this,f,v),+w.call(this,f,v)])):m.length&&(T(),m=[],g=[]);return m.length&&T(),d.length?d.join(""):null}return f.x=function(t){return arguments.length?(e=r=t,f):r},f.x0=function(t){return arguments.length?(e=t,f):e},f.x1=function(t){return arguments.length?(r=t,f):r},f.y=function(t){return arguments.length?(n=i=t,f):i},f.y0=function(t){return arguments.length?(n=t,f):n},f.y1=function(t){return arguments.length?(i=t,f):i},f.defined=function(t){return arguments.length?(a=t,f):a},f.interpolate=function(t){return arguments.length?(s="function"==typeof t?o=t:(o=li.get(t)||ci).key,l=o.reverse||o,c=o.closed?"M":"L",f):s},f.tension=function(t){return arguments.length?(u=t,f):u},f}function ki(t){return t.source}function Ai(t){return t.target}function Mi(t){return t.radius}function Si(t){return[t.x,t.y]}function Ei(t){return function(){var e=t.apply(this,arguments),r=e[0],n=e[1]-Et;return[r*Math.cos(n),r*Math.sin(n)]}}function Li(){return 64}function Ci(){return"circle"}function Pi(t){var e=Math.sqrt(t/At);return"M0,"+e+"A"+e+","+e+" 0 1,1 0,"+-e+"A"+e+","+e+" 0 1,1 0,"+e+"Z"}t.svg.line.radial=function(){var t=si(wi);return t.radius=t.x,delete t.x,t.angle=t.y,delete t.y,t},fi.reverse=hi,hi.reverse=fi,t.svg.area=function(){return Ti(C)},t.svg.area.radial=function(){var t=Ti(wi);return t.radius=t.x,delete t.x,t.innerRadius=t.x0,delete t.x0,t.outerRadius=t.x1,delete t.x1,t.angle=t.y,delete t.y,t.startAngle=t.y0,delete t.y0,t.endAngle=t.y1,delete t.y1,t},t.svg.chord=function(){var t=ki,e=Ai,r=Mi,n=ei,i=ri;function a(r,n){var i,a,c=o(this,t,r,n),u=o(this,e,r,n);return"M"+c.p0+s(c.r,c.p1,c.a1-c.a0)+(a=u,((i=c).a0==a.a0&&i.a1==a.a1?l(c.r,c.p1,c.r,c.p0):l(c.r,c.p1,u.r,u.p0)+s(u.r,u.p1,u.a1-u.a0)+l(u.r,u.p1,c.r,c.p0))+"Z")}function o(t,e,a,o){var s=e.call(t,a,o),l=r.call(t,s,o),c=n.call(t,s,o)-Et,u=i.call(t,s,o)-Et;return{r:l,a0:c,a1:u,p0:[l*Math.cos(c),l*Math.sin(c)],p1:[l*Math.cos(u),l*Math.sin(u)]}}function s(t,e,r){return"A"+t+","+t+" 0 "+ +(r>At)+",1 "+e}function l(t,e,r,n){return"Q 0,0 "+n}return a.radius=function(t){return arguments.length?(r=ce(t),a):r},a.source=function(e){return arguments.length?(t=ce(e),a):t},a.target=function(t){return arguments.length?(e=ce(t),a):e},a.startAngle=function(t){return arguments.length?(n=ce(t),a):n},a.endAngle=function(t){return arguments.length?(i=ce(t),a):i},a},t.svg.diagonal=function(){var t=ki,e=Ai,r=Si;function n(n,i){var a=t.call(this,n,i),o=e.call(this,n,i),s=(a.y+o.y)/2,l=[a,{x:a.x,y:s},{x:o.x,y:s},o];return"M"+(l=l.map(r))[0]+"C"+l[1]+" "+l[2]+" "+l[3]}return n.source=function(e){return arguments.length?(t=ce(e),n):t},n.target=function(t){return arguments.length?(e=ce(t),n):e},n.projection=function(t){return arguments.length?(r=t,n):r},n},t.svg.diagonal.radial=function(){var e=t.svg.diagonal(),r=Si,n=e.projection;return e.projection=function(t){return arguments.length?n(Ei(r=t)):r},e},t.svg.symbol=function(){var t=Ci,e=Li;function r(r,n){return(Ii.get(t.call(this,r,n))||Pi)(e.call(this,r,n))}return r.type=function(e){return arguments.length?(t=ce(e),r):t},r.size=function(t){return arguments.length?(e=ce(t),r):e},r};var Ii=t.map({circle:Pi,cross:function(t){var e=Math.sqrt(t/5)/2;return"M"+-3*e+","+-e+"H"+-e+"V"+-3*e+"H"+e+"V"+-e+"H"+3*e+"V"+e+"H"+e+"V"+3*e+"H"+-e+"V"+e+"H"+-3*e+"Z"},diamond:function(t){var e=Math.sqrt(t/(2*zi)),r=e*zi;return"M0,"+-e+"L"+r+",0 0,"+e+" "+-r+",0Z"},square:function(t){var e=Math.sqrt(t)/2;return"M"+-e+","+-e+"L"+e+","+-e+" "+e+","+e+" "+-e+","+e+"Z"},"triangle-down":function(t){var e=Math.sqrt(t/Oi),r=e*Oi/2;return"M0,"+r+"L"+e+","+-r+" "+-e+","+-r+"Z"},"triangle-up":function(t){var e=Math.sqrt(t/Oi),r=e*Oi/2;return"M0,"+-r+"L"+e+","+r+" "+-e+","+r+"Z"}});t.svg.symbolTypes=Ii.keys();var Oi=Math.sqrt(3),zi=Math.tan(30*Lt);Y.transition=function(t){for(var e,r,n=Bi||++Ui,i=qi(t),a=[],o=Ni||{time:Date.now(),ease:Er,delay:0,duration:250},s=-1,l=this.length;++s<l;){a.push(e=[]);for(var c=this[s],u=-1,f=c.length;++u<f;)(r=c[u])&&Gi(r,u,i,n,o),e.push(r)}return Fi(a,i,n)},Y.interrupt=function(t){return this.each(null==t?Di:Ri(qi(t)))};var Di=Ri(qi());function Ri(t){return function(){var e,r,n;(e=this[t])&&(n=e[r=e.active])&&(n.timer.c=null,n.timer.t=NaN,--e.count?delete e[r]:delete this[t],e.active+=.5,n.event&&n.event.interrupt.call(this,this.__data__,n.index))}}function Fi(t,e,r){return U(t,ji),t.namespace=e,t.id=r,t}var Bi,Ni,ji=[],Ui=0;function Vi(t,e,r,n){var i=t.id,a=t.namespace;return ut(t,"function"==typeof r?function(t,o,s){t[a][i].tween.set(e,n(r.call(t,t.__data__,o,s)))}:(r=n(r),function(t){t[a][i].tween.set(e,r)}))}function Hi(t){return null==t&&(t=""),function(){this.textContent=t}}function qi(t){return null==t?"__transition__":"__transition_"+t+"__"}function Gi(t,e,r,n,i){var a,o,s,l,c,u=t[r]||(t[r]={active:0,count:0}),f=u[n];function h(r){var i=u.active,h=u[i];for(var d in h&&(h.timer.c=null,h.timer.t=NaN,--u.count,delete u[i],h.event&&h.event.interrupt.call(t,t.__data__,h.index)),u)if(+d<n){var m=u[d];m.timer.c=null,m.timer.t=NaN,--u.count,delete u[d]}o.c=p,ve((function(){return o.c&&p(r||1)&&(o.c=null,o.t=NaN),1}),0,a),u.active=n,f.event&&f.event.start.call(t,t.__data__,e),c=[],f.tween.forEach((function(r,n){(n=n.call(t,t.__data__,e))&&c.push(n)})),l=f.ease,s=f.duration}function p(i){for(var a=i/s,o=l(a),h=c.length;h>0;)c[--h].call(t,o);if(a>=1)return f.event&&f.event.end.call(t,t.__data__,e),--u.count?delete u[n]:delete t[r],1}f||(a=i.time,o=ve((function(t){var e=f.delay;if(o.t=e+a,e<=t)return h(t-e);o.c=h}),0,a),f=u[n]={tween:new _,time:a,timer:o,delay:i.delay,duration:i.duration,ease:i.ease,index:e},i=null,++u.count)}ji.call=Y.call,ji.empty=Y.empty,ji.node=Y.node,ji.size=Y.size,t.transition=function(e,r){return e&&e.transition?Bi?e.transition(r):e:t.selection().transition(e)},t.transition.prototype=ji,ji.select=function(t){var e,r,n,i=this.id,a=this.namespace,o=[];t=W(t);for(var s=-1,l=this.length;++s<l;){o.push(e=[]);for(var c=this[s],u=-1,f=c.length;++u<f;)(n=c[u])&&(r=t.call(n,n.__data__,u,s))?("__data__"in n&&(r.__data__=n.__data__),Gi(r,u,a,i,n[a][i]),e.push(r)):e.push(null)}return Fi(o,a,i)},ji.selectAll=function(t){var e,r,n,i,a,o=this.id,s=this.namespace,l=[];t=X(t);for(var c=-1,u=this.length;++c<u;)for(var f=this[c],h=-1,p=f.length;++h<p;)if(n=f[h]){a=n[s][o],r=t.call(n,n.__data__,h,c),l.push(e=[]);for(var d=-1,m=r.length;++d<m;)(i=r[d])&&Gi(i,d,s,o,a),e.push(i)}return Fi(l,s,o)},ji.filter=function(t){var e,r,n=[];"function"!=typeof t&&(t=lt(t));for(var i=0,a=this.length;i<a;i++){n.push(e=[]);for(var o,s=0,l=(o=this[i]).length;s<l;s++)(r=o[s])&&t.call(r,r.__data__,s,i)&&e.push(r)}return Fi(n,this.namespace,this.id)},ji.tween=function(t,e){var r=this.id,n=this.namespace;return arguments.length<2?this.node()[n][r].tween.get(t):ut(this,null==e?function(e){e[n][r].tween.remove(t)}:function(i){i[n][r].tween.set(t,e)})},ji.attr=function(e,r){if(arguments.length<2){for(r in e)this.attr(r,e[r]);return this}var n="transform"==e?Nr:yr,i=t.ns.qualify(e);function a(){this.removeAttribute(i)}function o(){this.removeAttributeNS(i.space,i.local)}function s(t){return null==t?a:(t+="",function(){var e,r=this.getAttribute(i);return r!==t&&(e=n(r,t),function(t){this.setAttribute(i,e(t))})})}function l(t){return null==t?o:(t+="",function(){var e,r=this.getAttributeNS(i.space,i.local);return r!==t&&(e=n(r,t),function(t){this.setAttributeNS(i.space,i.local,e(t))})})}return Vi(this,"attr."+e,r,i.local?l:s)},ji.attrTween=function(e,r){var n=t.ns.qualify(e);return this.tween("attr."+e,n.local?function(t,e){var i=r.call(this,t,e,this.getAttributeNS(n.space,n.local));return i&&function(t){this.setAttributeNS(n.space,n.local,i(t))}}:function(t,e){var i=r.call(this,t,e,this.getAttribute(n));return i&&function(t){this.setAttribute(n,i(t))}})},ji.style=function(t,e,r){var n=arguments.length;if(n<3){if("string"!=typeof t){for(r in n<2&&(e=""),t)this.style(r,t[r],e);return this}r=""}function i(){this.style.removeProperty(t)}function a(e){return null==e?i:(e+="",function(){var n,i=o(this).getComputedStyle(this,null).getPropertyValue(t);return i!==e&&(n=yr(i,e),function(e){this.style.setProperty(t,n(e),r)})})}return Vi(this,"style."+t,e,a)},ji.styleTween=function(t,e,r){function n(n,i){var a=e.call(this,n,i,o(this).getComputedStyle(this,null).getPropertyValue(t));return a&&function(e){this.style.setProperty(t,a(e),r)}}return arguments.length<3&&(r=""),this.tween("style."+t,n)},ji.text=function(t){return Vi(this,"text",t,Hi)},ji.remove=function(){var t=this.namespace;return this.each("end.transition",(function(){var e;this[t].count<2&&(e=this.parentNode)&&e.removeChild(this)}))},ji.ease=function(e){var r=this.id,n=this.namespace;return arguments.length<1?this.node()[n][r].ease:("function"!=typeof e&&(e=t.ease.apply(t,arguments)),ut(this,(function(t){t[n][r].ease=e})))},ji.delay=function(t){var e=this.id,r=this.namespace;return arguments.length<1?this.node()[r][e].delay:ut(this,"function"==typeof t?function(n,i,a){n[r][e].delay=+t.call(n,n.__data__,i,a)}:(t=+t,function(n){n[r][e].delay=t}))},ji.duration=function(t){var e=this.id,r=this.namespace;return arguments.length<1?this.node()[r][e].duration:ut(this,"function"==typeof t?function(n,i,a){n[r][e].duration=Math.max(1,t.call(n,n.__data__,i,a))}:(t=Math.max(1,t),function(n){n[r][e].duration=t}))},ji.each=function(e,r){var n=this.id,i=this.namespace;if(arguments.length<2){var a=Ni,o=Bi;try{Bi=n,ut(this,(function(t,r,a){Ni=t[i][n],e.call(t,t.__data__,r,a)}))}finally{Ni=a,Bi=o}}else ut(this,(function(a){var o=a[i][n];(o.event||(o.event=t.dispatch("start","end","interrupt"))).on(e,r)}));return this},ji.transition=function(){for(var t,e,r,n=this.id,i=++Ui,a=this.namespace,o=[],s=0,l=this.length;s<l;s++){o.push(t=[]);for(var c,u=0,f=(c=this[s]).length;u<f;u++)(e=c[u])&&Gi(e,u,a,i,{time:(r=e[a][n]).time,ease:r.ease,delay:r.delay+r.duration,duration:r.duration}),t.push(e)}return Fi(o,a,i)},t.svg.axis=function(){var e,r=t.scale.linear(),i=Yi,a=6,o=6,s=3,l=[10],c=null;function u(n){n.each((function(){var n,u=t.select(this),f=this.__chart__||r,h=this.__chart__=r.copy(),p=null==c?h.ticks?h.ticks.apply(h,l):h.domain():c,d=null==e?h.tickFormat?h.tickFormat.apply(h,l):C:e,m=u.selectAll(".tick").data(p,h),g=m.enter().insert("g",".domain").attr("class","tick").style("opacity",kt),v=t.transition(m.exit()).style("opacity",kt).remove(),y=t.transition(m.order()).style("opacity",1),x=Math.max(a,0)+s,b=Dn(h),_=u.selectAll(".domain").data([0]),w=(_.enter().append("path").attr("class","domain"),t.transition(_));g.append("line"),g.append("text");var T,k,A,M,S=g.select("line"),E=y.select("line"),L=m.select("text").text(d),P=g.select("text"),I=y.select("text"),O="top"===i||"left"===i?-1:1;if("bottom"===i||"top"===i?(n=Xi,T="x",A="y",k="x2",M="y2",L.attr("dy",O<0?"0em":".71em").style("text-anchor","middle"),w.attr("d","M"+b[0]+","+O*o+"V0H"+b[1]+"V"+O*o)):(n=Zi,T="y",A="x",k="y2",M="x2",L.attr("dy",".32em").style("text-anchor",O<0?"end":"start"),w.attr("d","M"+O*o+","+b[0]+"H0V"+b[1]+"H"+O*o)),S.attr(M,O*a),P.attr(A,O*x),E.attr(k,0).attr(M,O*a),I.attr(T,0).attr(A,O*x),h.rangeBand){var z=h,D=z.rangeBand()/2;f=h=function(t){return z(t)+D}}else f.rangeBand?f=h:v.call(n,h,f);g.call(n,f,h),y.call(n,h,h)}))}return u.scale=function(t){return arguments.length?(r=t,u):r},u.orient=function(t){return arguments.length?(i=t in Wi?t+"":Yi,u):i},u.ticks=function(){return arguments.length?(l=n(arguments),u):l},u.tickValues=function(t){return arguments.length?(c=t,u):c},u.tickFormat=function(t){return arguments.length?(e=t,u):e},u.tickSize=function(t){var e=arguments.length;return e?(a=+t,o=+arguments[e-1],u):a},u.innerTickSize=function(t){return arguments.length?(a=+t,u):a},u.outerTickSize=function(t){return arguments.length?(o=+t,u):o},u.tickPadding=function(t){return arguments.length?(s=+t,u):s},u.tickSubdivide=function(){return arguments.length&&u},u};var Yi="bottom",Wi={top:1,right:1,bottom:1,left:1};function Xi(t,e,r){t.attr("transform",(function(t){var n=e(t);return"translate("+(isFinite(n)?n:r(t))+",0)"}))}function Zi(t,e,r){t.attr("transform",(function(t){var n=e(t);return"translate(0,"+(isFinite(n)?n:r(t))+")"}))}t.svg.brush=function(){var e,r,n=N(h,"brushstart","brush","brushend"),i=null,a=null,s=[0,0],l=[0,0],c=!0,u=!0,f=Ki[0];function h(e){e.each((function(){var e=t.select(this).style("pointer-events","all").style("-webkit-tap-highlight-color","rgba(0,0,0,0)").on("mousedown.brush",g).on("touchstart.brush",g),r=e.selectAll(".background").data([0]);r.enter().append("rect").attr("class","background").style("visibility","hidden").style("cursor","crosshair"),e.selectAll(".extent").data([0]).enter().append("rect").attr("class","extent").style("cursor","move");var n=e.selectAll(".resize").data(f,C);n.exit().remove(),n.enter().append("g").attr("class",(function(t){return"resize "+t})).style("cursor",(function(t){return Ji[t]})).append("rect").attr("x",(function(t){return/[ew]$/.test(t)?-3:null})).attr("y",(function(t){return/^[ns]/.test(t)?-3:null})).attr("width",6).attr("height",6).style("visibility","hidden"),n.style("display",h.empty()?"none":null);var o,s=t.transition(e),l=t.transition(r);i&&(o=Dn(i),l.attr("x",o[0]).attr("width",o[1]-o[0]),d(s)),a&&(o=Dn(a),l.attr("y",o[0]).attr("height",o[1]-o[0]),m(s)),p(s)}))}function p(t){t.selectAll(".resize").attr("transform",(function(t){return"translate("+s[+/e$/.test(t)]+","+l[+/^s/.test(t)]+")"}))}function d(t){t.select(".extent").attr("x",s[0]),t.selectAll(".extent,.n>rect,.s>rect").attr("width",s[1]-s[0])}function m(t){t.select(".extent").attr("y",l[0]),t.selectAll(".extent,.e>rect,.w>rect").attr("height",l[1]-l[0])}function g(){var f,g,v=this,y=t.select(t.event.target),x=n.of(v,arguments),b=t.select(v),_=y.datum(),w=!/^(n|s)$/.test(_)&&i,T=!/^(e|w)$/.test(_)&&a,k=y.classed("extent"),A=bt(v),M=t.mouse(v),S=t.select(o(v)).on("keydown.brush",C).on("keyup.brush",P);if(t.event.changedTouches?S.on("touchmove.brush",I).on("touchend.brush",z):S.on("mousemove.brush",I).on("mouseup.brush",z),b.interrupt().selectAll("*").interrupt(),k)M[0]=s[0]-M[0],M[1]=l[0]-M[1];else if(_){var E=+/w$/.test(_),L=+/^n/.test(_);g=[s[1-E]-M[0],l[1-L]-M[1]],M[0]=s[E],M[1]=l[L]}else t.event.altKey&&(f=M.slice());function C(){32==t.event.keyCode&&(k||(f=null,M[0]-=s[1],M[1]-=l[1],k=2),F())}function P(){32==t.event.keyCode&&2==k&&(M[0]+=s[1],M[1]+=l[1],k=0,F())}function I(){var e=t.mouse(v),r=!1;g&&(e[0]+=g[0],e[1]+=g[1]),k||(t.event.altKey?(f||(f=[(s[0]+s[1])/2,(l[0]+l[1])/2]),M[0]=s[+(e[0]<f[0])],M[1]=l[+(e[1]<f[1])]):f=null),w&&O(e,i,0)&&(d(b),r=!0),T&&O(e,a,1)&&(m(b),r=!0),r&&(p(b),x({type:"brush",mode:k?"move":"resize"}))}function O(t,n,i){var a,o,h=Dn(n),p=h[0],d=h[1],m=M[i],g=i?l:s,v=g[1]-g[0];if(k&&(p-=m,d-=v+m),a=(i?u:c)?Math.max(p,Math.min(d,t[i])):t[i],k?o=(a+=m)+v:(f&&(m=Math.max(p,Math.min(d,2*f[i]-a))),m<a?(o=a,a=m):o=m),g[0]!=a||g[1]!=o)return i?r=null:e=null,g[0]=a,g[1]=o,!0}function z(){I(),b.style("pointer-events","all").selectAll(".resize").style("display",h.empty()?"none":null),t.select("body").style("cursor",null),S.on("mousemove.brush",null).on("mouseup.brush",null).on("touchmove.brush",null).on("touchend.brush",null).on("keydown.brush",null).on("keyup.brush",null),A(),x({type:"brushend"})}b.style("pointer-events","none").selectAll(".resize").style("display",null),t.select("body").style("cursor",y.style("cursor")),x({type:"brushstart"}),I()}return h.event=function(i){i.each((function(){var i=n.of(this,arguments),a={x:s,y:l,i:e,j:r},o=this.__chart__||a;this.__chart__=a,Bi?t.select(this).transition().each("start.brush",(function(){e=o.i,r=o.j,s=o.x,l=o.y,i({type:"brushstart"})})).tween("brush:brush",(function(){var t=xr(s,a.x),n=xr(l,a.y);return e=r=null,function(e){s=a.x=t(e),l=a.y=n(e),i({type:"brush",mode:"resize"})}})).each("end.brush",(function(){e=a.i,r=a.j,i({type:"brush",mode:"resize"}),i({type:"brushend"})})):(i({type:"brushstart"}),i({type:"brush",mode:"resize"}),i({type:"brushend"}))}))},h.x=function(t){return arguments.length?(f=Ki[!(i=t)<<1|!a],h):i},h.y=function(t){return arguments.length?(f=Ki[!i<<1|!(a=t)],h):a},h.clamp=function(t){return arguments.length?(i&&a?(c=!!t[0],u=!!t[1]):i?c=!!t:a&&(u=!!t),h):i&&a?[c,u]:i?c:a?u:null},h.extent=function(t){var n,o,c,u,f;return arguments.length?(i&&(n=t[0],o=t[1],a&&(n=n[0],o=o[0]),e=[n,o],i.invert&&(n=i(n),o=i(o)),o<n&&(f=n,n=o,o=f),n==s[0]&&o==s[1]||(s=[n,o])),a&&(c=t[0],u=t[1],i&&(c=c[1],u=u[1]),r=[c,u],a.invert&&(c=a(c),u=a(u)),u<c&&(f=c,c=u,u=f),c==l[0]&&u==l[1]||(l=[c,u])),h):(i&&(e?(n=e[0],o=e[1]):(n=s[0],o=s[1],i.invert&&(n=i.invert(n),o=i.invert(o)),o<n&&(f=n,n=o,o=f))),a&&(r?(c=r[0],u=r[1]):(c=l[0],u=l[1],a.invert&&(c=a.invert(c),u=a.invert(u)),u<c&&(f=c,c=u,u=f))),i&&a?[[n,c],[o,u]]:i?[n,o]:a&&[c,u])},h.clear=function(){return h.empty()||(s=[0,0],l=[0,0],e=r=null),h},h.empty=function(){return!!i&&s[0]==s[1]||!!a&&l[0]==l[1]},t.rebind(h,n,"on")};var Ji={n:"ns-resize",e:"ew-resize",s:"ns-resize",w:"ew-resize",nw:"nwse-resize",ne:"nesw-resize",se:"nwse-resize",sw:"nesw-resize"},Ki=[["n","e","s","w","nw","ne","se","sw"],["e","w"],["n","s"],[]];function Qi(t){return JSON.parse(t.responseText)}function $i(t){var e=i.createRange();return e.selectNode(i.body),e.createContextualFragment(t.responseText)}t.text=ue((function(t){return t.responseText})),t.json=function(t,e){return fe(t,"application/json",Qi,e)},t.html=function(t,e){return fe(t,"text/html",$i,e)},t.xml=ue((function(t){return t.responseXML})),"object"==typeof e&&e.exports?e.exports=t:this.d3=t}).apply(self)},{}],59:[function(t,e,r){"use strict";e.exports=t("./quad")},{"./quad":60}],60:[function(t,e,r){"use strict";var n=t("binary-search-bounds"),i=t("clamp"),a=t("parse-rect"),o=t("array-bounds"),s=t("pick-by-alias"),l=t("defined"),c=t("flatten-vertex-data"),u=t("is-obj"),f=t("dtype"),h=t("math-log2");function p(t,e){for(var r=e[0],n=e[1],a=1/(e[2]-r),o=1/(e[3]-n),s=new Array(t.length),l=0,c=t.length/2;l<c;l++)s[2*l]=i((t[2*l]-r)*a,0,1),s[2*l+1]=i((t[2*l+1]-n)*o,0,1);return s}e.exports=function(t,e){e||(e={}),t=c(t,"float64"),e=s(e,{bounds:"range bounds dataBox databox",maxDepth:"depth maxDepth maxdepth level maxLevel maxlevel levels",dtype:"type dtype format out dst output destination"});var r=l(e.maxDepth,255),i=l(e.bounds,o(t,2));i[0]===i[2]&&i[2]++,i[1]===i[3]&&i[3]++;var d,m=p(t,i),g=t.length>>>1;e.dtype||(e.dtype="array"),"string"==typeof e.dtype?d=new(f(e.dtype))(g):e.dtype&&(d=e.dtype,Array.isArray(d)&&(d.length=g));for(var v=0;v<g;++v)d[v]=v;var y=[],x=[],b=[],_=[];!function t(e,n,i,a,o,s){if(!a.length)return null;var l=y[o]||(y[o]=[]),c=b[o]||(b[o]=[]),u=x[o]||(x[o]=[]),f=l.length;if(++o>r||s>1073741824){for(var h=0;h<a.length;h++)l.push(a[h]),c.push(s),u.push(null,null,null,null);return f}if(l.push(a[0]),c.push(s),a.length<=1)return u.push(null,null,null,null),f;for(var p=.5*i,d=e+p,g=n+p,v=[],_=[],w=[],T=[],k=1,A=a.length;k<A;k++){var M=a[k],S=m[2*M],E=m[2*M+1];S<d?E<g?v.push(M):_.push(M):E<g?w.push(M):T.push(M)}return s<<=2,u.push(t(e,n,p,v,o,s),t(e,g,p,_,o,s+1),t(d,n,p,w,o,s+2),t(d,g,p,T,o,s+3)),f}(0,0,1,d,0,1);for(var w=0,T=0;T<y.length;T++){var k=y[T];if(d.set)d.set(k,w);else for(var A=0,M=k.length;A<M;A++)d[A+w]=k[A];var S=w+y[T].length;_[T]=[w,S],w=S}return d.range=function(){var e,r=[],n=arguments.length;for(;n--;)r[n]=arguments[n];if(u(r[r.length-1])){var o=r.pop();r.length||null==o.x&&null==o.l&&null==o.left||(r=[o],e={}),e=s(o,{level:"level maxLevel",d:"d diam diameter r radius px pxSize pixel pixelSize maxD size minSize",lod:"lod details ranges offsets"})}else e={};r.length||(r=i);var c=a.apply(void 0,r),f=[Math.min(c.x,c.x+c.width),Math.min(c.y,c.y+c.height),Math.max(c.x,c.x+c.width),Math.max(c.y,c.y+c.height)],d=f[0],m=f[1],g=f[2],v=f[3],b=p([d,m,g,v],i),_=b[0],w=b[1],T=b[2],k=b[3],A=l(e.level,y.length);if(null!=e.d){var M;"number"==typeof e.d?M=[e.d,e.d]:e.d.length&&(M=e.d),A=Math.min(Math.max(Math.ceil(-h(Math.abs(M[0])/(i[2]-i[0]))),Math.ceil(-h(Math.abs(M[1])/(i[3]-i[1])))),A)}if(A=Math.min(A,y.length),e.lod)return E(_,w,T,k,A);var S=[];function L(e,r,n,i,a,o){if(null!==a&&null!==o&&!(_>e+n||w>r+n||T<e||k<r||i>=A||a===o)){var s=y[i];void 0===o&&(o=s.length);for(var l=a;l<o;l++){var c=s[l],u=t[2*c],f=t[2*c+1];u>=d&&u<=g&&f>=m&&f<=v&&S.push(c)}var h=x[i],p=h[4*a+0],b=h[4*a+1],M=h[4*a+2],E=h[4*a+3],P=C(h,a+1),I=.5*n,O=i+1;L(e,r,I,O,p,b||M||E||P),L(e,r+I,I,O,b,M||E||P),L(e+I,r,I,O,M,E||P),L(e+I,r+I,I,O,E,P)}}function C(t,e){for(var r=null,n=0;null===r;)if(r=t[4*e+n],++n>t.length)return null;return r}return L(0,0,1,0,0,1),S},d;function E(t,e,r,i,a){for(var o=[],s=0;s<a;s++){var l=b[s],c=_[s][0],u=L(t,e,s),f=L(r,i,s),h=n.ge(l,u),p=n.gt(l,f,h,l.length-1);o[s]=[h+c,p+c]}return o}function L(t,e,r){for(var n=1,i=.5,a=.5,o=.5,s=0;s<r;s++)n<<=2,n+=t<i?e<a?0:1:e<a?2:3,o*=.5,i+=t<i?-o:o,a+=e<a?-o:o;return n}}},{"array-bounds":67,"binary-search-bounds":76,clamp:81,defined:118,dtype:121,"flatten-vertex-data":186,"is-obj":230,"math-log2":235,"parse-rect":244,"pick-by-alias":248}],61:[function(t,e,r){"use strict";Object.defineProperty(r,"__esModule",{value:!0});var n=t("@turf/meta");function i(t){var e=0;if(t&&t.length>0){e+=Math.abs(a(t[0]));for(var r=1;r<t.length;r++)e-=Math.abs(a(t[r]))}return e}function a(t){var e,r,n,i,a,s,l=0,c=t.length;if(c>2){for(s=0;s<c;s++)s===c-2?(n=c-2,i=c-1,a=0):s===c-1?(n=c-1,i=0,a=1):(n=s,i=s+1,a=s+2),e=t[n],r=t[i],l+=(o(t[a][0])-o(e[0]))*Math.sin(o(r[1]));l=6378137*l*6378137/2}return l}function o(t){return t*Math.PI/180}r.default=function(t){return n.geomReduce(t,(function(t,e){return t+function(t){var e,r=0;switch(t.type){case"Polygon":return i(t.coordinates);case"MultiPolygon":for(e=0;e<t.coordinates.length;e++)r+=i(t.coordinates[e]);return r;case"Point":case"MultiPoint":case"LineString":case"MultiLineString":return 0}return 0}(e)}),0)}},{"@turf/meta":65}],62:[function(t,e,r){"use strict";Object.defineProperty(r,"__esModule",{value:!0});var n=t("@turf/meta");function i(t){var e=[1/0,1/0,-1/0,-1/0];return n.coordEach(t,(function(t){e[0]>t[0]&&(e[0]=t[0]),e[1]>t[1]&&(e[1]=t[1]),e[2]<t[0]&&(e[2]=t[0]),e[3]<t[1]&&(e[3]=t[1])})),e}i.default=i,r.default=i},{"@turf/meta":65}],63:[function(t,e,r){"use strict";Object.defineProperty(r,"__esModule",{value:!0});var n=t("@turf/meta"),i=t("@turf/helpers");r.default=function(t,e){void 0===e&&(e={});var r=0,a=0,o=0;return n.coordEach(t,(function(t){r+=t[0],a+=t[1],o++}),!0),i.point([r/o,a/o],e.properties)}},{"@turf/helpers":64,"@turf/meta":65}],64:[function(t,e,r){"use strict";function n(t,e,r){void 0===r&&(r={});var n={type:"Feature"};return(0===r.id||r.id)&&(n.id=r.id),r.bbox&&(n.bbox=r.bbox),n.properties=e||{},n.geometry=t,n}function i(t,e,r){if(void 0===r&&(r={}),!t)throw new Error("coordinates is required");if(!Array.isArray(t))throw new Error("coordinates must be an Array");if(t.length<2)throw new Error("coordinates must be at least 2 numbers long");if(!d(t[0])||!d(t[1]))throw new Error("coordinates must contain numbers");return n({type:"Point",coordinates:t},e,r)}function a(t,e,r){void 0===r&&(r={});for(var i=0,a=t;i<a.length;i++){var o=a[i];if(o.length<4)throw new Error("Each LinearRing of a Polygon must have 4 or more Positions.");for(var s=0;s<o[o.length-1].length;s++)if(o[o.length-1][s]!==o[0][s])throw new Error("First and last Position are not equivalent.")}return n({type:"Polygon",coordinates:t},e,r)}function o(t,e,r){if(void 0===r&&(r={}),t.length<2)throw new Error("coordinates must be an array of two or more positions");return n({type:"LineString",coordinates:t},e,r)}function s(t,e){void 0===e&&(e={});var r={type:"FeatureCollection"};return e.id&&(r.id=e.id),e.bbox&&(r.bbox=e.bbox),r.features=t,r}function l(t,e,r){return void 0===r&&(r={}),n({type:"MultiLineString",coordinates:t},e,r)}function c(t,e,r){return void 0===r&&(r={}),n({type:"MultiPoint",coordinates:t},e,r)}function u(t,e,r){return void 0===r&&(r={}),n({type:"MultiPolygon",coordinates:t},e,r)}function f(t,e){void 0===e&&(e="kilometers");var n=r.factors[e];if(!n)throw new Error(e+" units is invalid");return t*n}function h(t,e){void 0===e&&(e="kilometers");var n=r.factors[e];if(!n)throw new Error(e+" units is invalid");return t/n}function p(t){return 180*(t%(2*Math.PI))/Math.PI}function d(t){return!isNaN(t)&&null!==t&&!Array.isArray(t)}Object.defineProperty(r,"__esModule",{value:!0}),r.earthRadius=6371008.8,r.factors={centimeters:100*r.earthRadius,centimetres:100*r.earthRadius,degrees:r.earthRadius/111325,feet:3.28084*r.earthRadius,inches:39.37*r.earthRadius,kilometers:r.earthRadius/1e3,kilometres:r.earthRadius/1e3,meters:r.earthRadius,metres:r.earthRadius,miles:r.earthRadius/1609.344,millimeters:1e3*r.earthRadius,millimetres:1e3*r.earthRadius,nauticalmiles:r.earthRadius/1852,radians:1,yards:1.0936*r.earthRadius},r.unitsFactors={centimeters:100,centimetres:100,degrees:1/111325,feet:3.28084,inches:39.37,kilometers:.001,kilometres:.001,meters:1,metres:1,miles:1/1609.344,millimeters:1e3,millimetres:1e3,nauticalmiles:1/1852,radians:1/r.earthRadius,yards:1.0936133},r.areaFactors={acres:247105e-9,centimeters:1e4,centimetres:1e4,feet:10.763910417,hectares:1e-4,inches:1550.003100006,kilometers:1e-6,kilometres:1e-6,meters:1,metres:1,miles:386e-9,millimeters:1e6,millimetres:1e6,yards:1.195990046},r.feature=n,r.geometry=function(t,e,r){switch(void 0===r&&(r={}),t){case"Point":return i(e).geometry;case"LineString":return o(e).geometry;case"Polygon":return a(e).geometry;case"MultiPoint":return c(e).geometry;case"MultiLineString":return l(e).geometry;case"MultiPolygon":return u(e).geometry;default:throw new Error(t+" is invalid")}},r.point=i,r.points=function(t,e,r){return void 0===r&&(r={}),s(t.map((function(t){return i(t,e)})),r)},r.polygon=a,r.polygons=function(t,e,r){return void 0===r&&(r={}),s(t.map((function(t){return a(t,e)})),r)},r.lineString=o,r.lineStrings=function(t,e,r){return void 0===r&&(r={}),s(t.map((function(t){return o(t,e)})),r)},r.featureCollection=s,r.multiLineString=l,r.multiPoint=c,r.multiPolygon=u,r.geometryCollection=function(t,e,r){return void 0===r&&(r={}),n({type:"GeometryCollection",geometries:t},e,r)},r.round=function(t,e){if(void 0===e&&(e=0),e&&!(e>=0))throw new Error("precision must be a positive number");var r=Math.pow(10,e||0);return Math.round(t*r)/r},r.radiansToLength=f,r.lengthToRadians=h,r.lengthToDegrees=function(t,e){return p(h(t,e))},r.bearingToAzimuth=function(t){var e=t%360;return e<0&&(e+=360),e},r.radiansToDegrees=p,r.degreesToRadians=function(t){return t%360*Math.PI/180},r.convertLength=function(t,e,r){if(void 0===e&&(e="kilometers"),void 0===r&&(r="kilometers"),!(t>=0))throw new Error("length must be a positive number");return f(h(t,e),r)},r.convertArea=function(t,e,n){if(void 0===e&&(e="meters"),void 0===n&&(n="kilometers"),!(t>=0))throw new Error("area must be a positive number");var i=r.areaFactors[e];if(!i)throw new Error("invalid original units");var a=r.areaFactors[n];if(!a)throw new Error("invalid final units");return t/i*a},r.isNumber=d,r.isObject=function(t){return!!t&&t.constructor===Object},r.validateBBox=function(t){if(!t)throw new Error("bbox is required");if(!Array.isArray(t))throw new Error("bbox must be an Array");if(4!==t.length&&6!==t.length)throw new Error("bbox must be an Array of 4 or 6 numbers");t.forEach((function(t){if(!d(t))throw new Error("bbox must only contain numbers")}))},r.validateId=function(t){if(!t)throw new Error("id is required");if(-1===["string","number"].indexOf(typeof t))throw new Error("id must be a number or a string")}},{}],65:[function(t,e,r){"use strict";Object.defineProperty(r,"__esModule",{value:!0});var n=t("@turf/helpers");function i(t,e,r){if(null!==t)for(var n,a,o,s,l,c,u,f,h=0,p=0,d=t.type,m="FeatureCollection"===d,g="Feature"===d,v=m?t.features.length:1,y=0;y<v;y++){l=(f=!!(u=m?t.features[y].geometry:g?t.geometry:t)&&"GeometryCollection"===u.type)?u.geometries.length:1;for(var x=0;x<l;x++){var b=0,_=0;if(null!==(s=f?u.geometries[x]:u)){c=s.coordinates;var w=s.type;switch(h=!r||"Polygon"!==w&&"MultiPolygon"!==w?0:1,w){case null:break;case"Point":if(!1===e(c,p,y,b,_))return!1;p++,b++;break;case"LineString":case"MultiPoint":for(n=0;n<c.length;n++){if(!1===e(c[n],p,y,b,_))return!1;p++,"MultiPoint"===w&&b++}"LineString"===w&&b++;break;case"Polygon":case"MultiLineString":for(n=0;n<c.length;n++){for(a=0;a<c[n].length-h;a++){if(!1===e(c[n][a],p,y,b,_))return!1;p++}"MultiLineString"===w&&b++,"Polygon"===w&&_++}"Polygon"===w&&b++;break;case"MultiPolygon":for(n=0;n<c.length;n++){for(_=0,a=0;a<c[n].length;a++){for(o=0;o<c[n][a].length-h;o++){if(!1===e(c[n][a][o],p,y,b,_))return!1;p++}_++}b++}break;case"GeometryCollection":for(n=0;n<s.geometries.length;n++)if(!1===i(s.geometries[n],e,r))return!1;break;default:throw new Error("Unknown Geometry Type")}}}}}function a(t,e){var r;switch(t.type){case"FeatureCollection":for(r=0;r<t.features.length&&!1!==e(t.features[r].properties,r);r++);break;case"Feature":e(t.properties,0)}}function o(t,e){if("Feature"===t.type)e(t,0);else if("FeatureCollection"===t.type)for(var r=0;r<t.features.length&&!1!==e(t.features[r],r);r++);}function s(t,e){var r,n,i,a,o,s,l,c,u,f,h=0,p="FeatureCollection"===t.type,d="Feature"===t.type,m=p?t.features.length:1;for(r=0;r<m;r++){for(s=p?t.features[r].geometry:d?t.geometry:t,c=p?t.features[r].properties:d?t.properties:{},u=p?t.features[r].bbox:d?t.bbox:void 0,f=p?t.features[r].id:d?t.id:void 0,o=(l=!!s&&"GeometryCollection"===s.type)?s.geometries.length:1,i=0;i<o;i++)if(null!==(a=l?s.geometries[i]:s))switch(a.type){case"Point":case"LineString":case"MultiPoint":case"Polygon":case"MultiLineString":case"MultiPolygon":if(!1===e(a,h,c,u,f))return!1;break;case"GeometryCollection":for(n=0;n<a.geometries.length;n++)if(!1===e(a.geometries[n],h,c,u,f))return!1;break;default:throw new Error("Unknown Geometry Type")}else if(!1===e(null,h,c,u,f))return!1;h++}}function l(t,e){s(t,(function(t,r,i,a,o){var s,l=null===t?null:t.type;switch(l){case null:case"Point":case"LineString":case"Polygon":return!1!==e(n.feature(t,i,{bbox:a,id:o}),r,0)&&void 0}switch(l){case"MultiPoint":s="Point";break;case"MultiLineString":s="LineString";break;case"MultiPolygon":s="Polygon"}for(var c=0;c<t.coordinates.length;c++){var u={type:s,coordinates:t.coordinates[c]};if(!1===e(n.feature(u,i),r,c))return!1}}))}function c(t,e){l(t,(function(t,r,a){var o=0;if(t.geometry){var s=t.geometry.type;if("Point"!==s&&"MultiPoint"!==s){var l,c=0,u=0,f=0;return!1!==i(t,(function(i,s,h,p,d){if(void 0===l||r>c||p>u||d>f)return l=i,c=r,u=p,f=d,void(o=0);var m=n.lineString([l,i],t.properties);if(!1===e(m,r,a,d,o))return!1;o++,l=i}))&&void 0}}}))}function u(t,e){if(!t)throw new Error("geojson is required");l(t,(function(t,r,i){if(null!==t.geometry){var a=t.geometry.type,o=t.geometry.coordinates;switch(a){case"LineString":if(!1===e(t,r,i,0,0))return!1;break;case"Polygon":for(var s=0;s<o.length;s++)if(!1===e(n.lineString(o[s],t.properties),r,i,s))return!1}}}))}r.coordAll=function(t){var e=[];return i(t,(function(t){e.push(t)})),e},r.coordEach=i,r.coordReduce=function(t,e,r,n){var a=r;return i(t,(function(t,n,i,o,s){a=0===n&&void 0===r?t:e(a,t,n,i,o,s)}),n),a},r.featureEach=o,r.featureReduce=function(t,e,r){var n=r;return o(t,(function(t,i){n=0===i&&void 0===r?t:e(n,t,i)})),n},r.findPoint=function(t,e){if(e=e||{},!n.isObject(e))throw new Error("options is invalid");var r,i=e.featureIndex||0,a=e.multiFeatureIndex||0,o=e.geometryIndex||0,s=e.coordIndex||0,l=e.properties;switch(t.type){case"FeatureCollection":i<0&&(i=t.features.length+i),l=l||t.features[i].properties,r=t.features[i].geometry;break;case"Feature":l=l||t.properties,r=t.geometry;break;case"Point":case"MultiPoint":return null;case"LineString":case"Polygon":case"MultiLineString":case"MultiPolygon":r=t;break;default:throw new Error("geojson is invalid")}if(null===r)return null;var c=r.coordinates;switch(r.type){case"Point":return n.point(c,l,e);case"MultiPoint":return a<0&&(a=c.length+a),n.point(c[a],l,e);case"LineString":return s<0&&(s=c.length+s),n.point(c[s],l,e);case"Polygon":return o<0&&(o=c.length+o),s<0&&(s=c[o].length+s),n.point(c[o][s],l,e);case"MultiLineString":return a<0&&(a=c.length+a),s<0&&(s=c[a].length+s),n.point(c[a][s],l,e);case"MultiPolygon":return a<0&&(a=c.length+a),o<0&&(o=c[a].length+o),s<0&&(s=c[a][o].length-s),n.point(c[a][o][s],l,e)}throw new Error("geojson is invalid")},r.findSegment=function(t,e){if(e=e||{},!n.isObject(e))throw new Error("options is invalid");var r,i=e.featureIndex||0,a=e.multiFeatureIndex||0,o=e.geometryIndex||0,s=e.segmentIndex||0,l=e.properties;switch(t.type){case"FeatureCollection":i<0&&(i=t.features.length+i),l=l||t.features[i].properties,r=t.features[i].geometry;break;case"Feature":l=l||t.properties,r=t.geometry;break;case"Point":case"MultiPoint":return null;case"LineString":case"Polygon":case"MultiLineString":case"MultiPolygon":r=t;break;default:throw new Error("geojson is invalid")}if(null===r)return null;var c=r.coordinates;switch(r.type){case"Point":case"MultiPoint":return null;case"LineString":return s<0&&(s=c.length+s-1),n.lineString([c[s],c[s+1]],l,e);case"Polygon":return o<0&&(o=c.length+o),s<0&&(s=c[o].length+s-1),n.lineString([c[o][s],c[o][s+1]],l,e);case"MultiLineString":return a<0&&(a=c.length+a),s<0&&(s=c[a].length+s-1),n.lineString([c[a][s],c[a][s+1]],l,e);case"MultiPolygon":return a<0&&(a=c.length+a),o<0&&(o=c[a].length+o),s<0&&(s=c[a][o].length-s-1),n.lineString([c[a][o][s],c[a][o][s+1]],l,e)}throw new Error("geojson is invalid")},r.flattenEach=l,r.flattenReduce=function(t,e,r){var n=r;return l(t,(function(t,i,a){n=0===i&&0===a&&void 0===r?t:e(n,t,i,a)})),n},r.geomEach=s,r.geomReduce=function(t,e,r){var n=r;return s(t,(function(t,i,a,o,s){n=0===i&&void 0===r?t:e(n,t,i,a,o,s)})),n},r.lineEach=u,r.lineReduce=function(t,e,r){var n=r;return u(t,(function(t,i,a,o){n=0===i&&void 0===r?t:e(n,t,i,a,o)})),n},r.propEach=a,r.propReduce=function(t,e,r){var n=r;return a(t,(function(t,i){n=0===i&&void 0===r?t:e(n,t,i)})),n},r.segmentEach=c,r.segmentReduce=function(t,e,r){var n=r,i=!1;return c(t,(function(t,a,o,s,l){n=!1===i&&void 0===r?t:e(n,t,a,o,s,l),i=!0})),n}},{"@turf/helpers":64}],66:[function(t,e,r){e.exports=function(t){var e=0,r=0,n=0,i=0;return t.map((function(t){var a=(t=t.slice())[0],o=a.toUpperCase();if(a!=o)switch(t[0]=o,a){case"a":t[6]+=n,t[7]+=i;break;case"v":t[1]+=i;break;case"h":t[1]+=n;break;default:for(var s=1;s<t.length;)t[s++]+=n,t[s++]+=i}switch(o){case"Z":n=e,i=r;break;case"H":n=t[1];break;case"V":i=t[1];break;case"M":n=e=t[1],i=r=t[2];break;default:n=t[t.length-2],i=t[t.length-1]}return t}))}},{}],67:[function(t,e,r){"use strict";e.exports=function(t,e){if(!t||null==t.length)throw Error("Argument should be an array");e=null==e?1:Math.floor(e);for(var r=Array(2*e),n=0;n<e;n++){for(var i=-1/0,a=1/0,o=n,s=t.length;o<s;o+=e)t[o]>i&&(i=t[o]),t[o]<a&&(a=t[o]);r[n]=a,r[e+n]=i}return r}},{}],68:[function(t,e,r){"use strict";e.exports=function(t,e,r){if("function"==typeof Array.prototype.findIndex)return t.findIndex(e,r);if("function"!=typeof e)throw new TypeError("predicate must be a function");var n=Object(t),i=n.length;if(0===i)return-1;for(var a=0;a<i;a++)if(e.call(r,n[a],a,n))return a;return-1}},{}],69:[function(t,e,r){"use strict";var n=t("array-bounds");e.exports=function(t,e,r){if(!t||null==t.length)throw Error("Argument should be an array");null==e&&(e=1);null==r&&(r=n(t,e));for(var i=0;i<e;i++){var a=r[e+i],o=r[i],s=i,l=t.length;if(a===1/0&&o===-1/0)for(s=i;s<l;s+=e)t[s]=t[s]===a?1:t[s]===o?0:.5;else if(a===1/0)for(s=i;s<l;s+=e)t[s]=t[s]===a?1:0;else if(o===-1/0)for(s=i;s<l;s+=e)t[s]=t[s]===o?0:1;else{var c=a-o;for(s=i;s<l;s+=e)isNaN(t[s])||(t[s]=0===c?.5:(t[s]-o)/c)}}return t}},{"array-bounds":67}],70:[function(t,e,r){e.exports=function(t,e){var r="number"==typeof t,n="number"==typeof e;r&&!n?(e=t,t=0):r||n||(t=0,e=0);var i=(e|=0)-(t|=0);if(i<0)throw new Error("array length must be positive");for(var a=new Array(i),o=0,s=t;o<i;o++,s++)a[o]=s;return a}},{}],71:[function(t,e,r){(function(r){(function(){"use strict";var n=t("object-assign");
/*!
 * The buffer module from node.js, for the browser.
 *
 * @author   Feross Aboukhadijeh <> <http://feross.org>
 * @license  MIT
 */function i(t,e){if(t===e)return 0;for(var r=t.length,n=e.length,i=0,a=Math.min(r,n);i<a;++i)if(t[i]!==e[i]){r=t[i],n=e[i];break}return r<n?-1:n<r?1:0}function a(t){return r.Buffer&&"function"==typeof r.Buffer.isBuffer?r.Buffer.isBuffer(t):!(null==t||!t._isBuffer)}var o=t("util/"),s=Object.prototype.hasOwnProperty,l=Array.prototype.slice,c="foo"===function(){}.name;function u(t){return Object.prototype.toString.call(t)}function f(t){return!a(t)&&("function"==typeof r.ArrayBuffer&&("function"==typeof ArrayBuffer.isView?ArrayBuffer.isView(t):!!t&&(t instanceof DataView||!!(t.buffer&&t.buffer instanceof ArrayBuffer))))}var h=e.exports=y,p=/\s*function\s+([^\(\s]*)\s*/;function d(t){if(o.isFunction(t)){if(c)return t.name;var e=t.toString().match(p);return e&&e[1]}}function m(t,e){return"string"==typeof t?t.length<e?t:t.slice(0,e):t}function g(t){if(c||!o.isFunction(t))return o.inspect(t);var e=d(t);return"[Function"+(e?": "+e:"")+"]"}function v(t,e,r,n,i){throw new h.AssertionError({message:r,actual:t,expected:e,operator:n,stackStartFunction:i})}function y(t,e){t||v(t,!0,e,"==",h.ok)}function x(t,e,r,n){if(t===e)return!0;if(a(t)&&a(e))return 0===i(t,e);if(o.isDate(t)&&o.isDate(e))return t.getTime()===e.getTime();if(o.isRegExp(t)&&o.isRegExp(e))return t.source===e.source&&t.global===e.global&&t.multiline===e.multiline&&t.lastIndex===e.lastIndex&&t.ignoreCase===e.ignoreCase;if(null!==t&&"object"==typeof t||null!==e&&"object"==typeof e){if(f(t)&&f(e)&&u(t)===u(e)&&!(t instanceof Float32Array||t instanceof Float64Array))return 0===i(new Uint8Array(t.buffer),new Uint8Array(e.buffer));if(a(t)!==a(e))return!1;var s=(n=n||{actual:[],expected:[]}).actual.indexOf(t);return-1!==s&&s===n.expected.indexOf(e)||(n.actual.push(t),n.expected.push(e),function(t,e,r,n){if(null==t||null==e)return!1;if(o.isPrimitive(t)||o.isPrimitive(e))return t===e;if(r&&Object.getPrototypeOf(t)!==Object.getPrototypeOf(e))return!1;var i=b(t),a=b(e);if(i&&!a||!i&&a)return!1;if(i)return t=l.call(t),e=l.call(e),x(t,e,r);var s,c,u=T(t),f=T(e);if(u.length!==f.length)return!1;for(u.sort(),f.sort(),c=u.length-1;c>=0;c--)if(u[c]!==f[c])return!1;for(c=u.length-1;c>=0;c--)if(s=u[c],!x(t[s],e[s],r,n))return!1;return!0}(t,e,r,n))}return r?t===e:t==e}function b(t){return"[object Arguments]"==Object.prototype.toString.call(t)}function _(t,e){if(!t||!e)return!1;if("[object RegExp]"==Object.prototype.toString.call(e))return e.test(t);try{if(t instanceof e)return!0}catch(t){}return!Error.isPrototypeOf(e)&&!0===e.call({},t)}function w(t,e,r,n){var i;if("function"!=typeof e)throw new TypeError('"block" argument must be a function');"string"==typeof r&&(n=r,r=null),i=function(t){var e;try{t()}catch(t){e=t}return e}(e),n=(r&&r.name?" ("+r.name+").":".")+(n?" "+n:"."),t&&!i&&v(i,r,"Missing expected exception"+n);var a="string"==typeof n,s=!t&&i&&!r;if((!t&&o.isError(i)&&a&&_(i,r)||s)&&v(i,r,"Got unwanted exception"+n),t&&i&&r&&!_(i,r)||!t&&i)throw i}h.AssertionError=function(t){this.name="AssertionError",this.actual=t.actual,this.expected=t.expected,this.operator=t.operator,t.message?(this.message=t.message,this.generatedMessage=!1):(this.message=function(t){return m(g(t.actual),128)+" "+t.operator+" "+m(g(t.expected),128)}(this),this.generatedMessage=!0);var e=t.stackStartFunction||v;if(Error.captureStackTrace)Error.captureStackTrace(this,e);else{var r=new Error;if(r.stack){var n=r.stack,i=d(e),a=n.indexOf("\n"+i);if(a>=0){var o=n.indexOf("\n",a+1);n=n.substring(o+1)}this.stack=n}}},o.inherits(h.AssertionError,Error),h.fail=v,h.ok=y,h.equal=function(t,e,r){t!=e&&v(t,e,r,"==",h.equal)},h.notEqual=function(t,e,r){t==e&&v(t,e,r,"!=",h.notEqual)},h.deepEqual=function(t,e,r){x(t,e,!1)||v(t,e,r,"deepEqual",h.deepEqual)},h.deepStrictEqual=function(t,e,r){x(t,e,!0)||v(t,e,r,"deepStrictEqual",h.deepStrictEqual)},h.notDeepEqual=function(t,e,r){x(t,e,!1)&&v(t,e,r,"notDeepEqual",h.notDeepEqual)},h.notDeepStrictEqual=function t(e,r,n){x(e,r,!0)&&v(e,r,n,"notDeepStrictEqual",t)},h.strictEqual=function(t,e,r){t!==e&&v(t,e,r,"===",h.strictEqual)},h.notStrictEqual=function(t,e,r){t===e&&v(t,e,r,"!==",h.notStrictEqual)},h.throws=function(t,e,r){w(!0,t,e,r)},h.doesNotThrow=function(t,e,r){w(!1,t,e,r)},h.ifError=function(t){if(t)throw t},h.strict=n((function t(e,r){e||v(e,!0,r,"==",t)}),h,{equal:h.strictEqual,deepEqual:h.deepStrictEqual,notEqual:h.notStrictEqual,notDeepEqual:h.notDeepStrictEqual}),h.strict.strict=h.strict;var T=Object.keys||function(t){var e=[];for(var r in t)s.call(t,r)&&e.push(r);return e}}).call(this)}).call(this,"undefined"!=typeof global?global:"undefined"!=typeof self?self:"undefined"!=typeof window?window:{})},{"object-assign":242,"util/":74}],72:[function(t,e,r){"function"==typeof Object.create?e.exports=function(t,e){t.super_=e,t.prototype=Object.create(e.prototype,{constructor:{value:t,enumerable:!1,writable:!0,configurable:!0}})}:e.exports=function(t,e){t.super_=e;var r=function(){};r.prototype=e.prototype,t.prototype=new r,t.prototype.constructor=t}},{}],73:[function(t,e,r){e.exports=function(t){return t&&"object"==typeof t&&"function"==typeof t.copy&&"function"==typeof t.fill&&"function"==typeof t.readUInt8}},{}],74:[function(t,e,r){(function(e,n){(function(){var i=/%[sdj%]/g;r.format=function(t){if(!v(t)){for(var e=[],r=0;r<arguments.length;r++)e.push(s(arguments[r]));return e.join(" ")}r=1;for(var n=arguments,a=n.length,o=String(t).replace(i,(function(t){if("%%"===t)return"%";if(r>=a)return t;switch(t){case"%s":return String(n[r++]);case"%d":return Number(n[r++]);case"%j":try{return JSON.stringify(n[r++])}catch(t){return"[Circular]"}default:return t}})),l=n[r];r<a;l=n[++r])m(l)||!b(l)?o+=" "+l:o+=" "+s(l);return o},r.deprecate=function(t,i){if(y(n.process))return function(){return r.deprecate(t,i).apply(this,arguments)};if(!0===e.noDeprecation)return t;var a=!1;return function(){if(!a){if(e.throwDeprecation)throw new Error(i);e.traceDeprecation?console.trace(i):console.error(i),a=!0}return t.apply(this,arguments)}};var a,o={};function s(t,e){var n={seen:[],stylize:c};return arguments.length>=3&&(n.depth=arguments[2]),arguments.length>=4&&(n.colors=arguments[3]),d(e)?n.showHidden=e:e&&r._extend(n,e),y(n.showHidden)&&(n.showHidden=!1),y(n.depth)&&(n.depth=2),y(n.colors)&&(n.colors=!1),y(n.customInspect)&&(n.customInspect=!0),n.colors&&(n.stylize=l),u(n,t,n.depth)}function l(t,e){var r=s.styles[e];return r?"\x1b["+s.colors[r][0]+"m"+t+"\x1b["+s.colors[r][1]+"m":t}function c(t,e){return t}function u(t,e,n){if(t.customInspect&&e&&T(e.inspect)&&e.inspect!==r.inspect&&(!e.constructor||e.constructor.prototype!==e)){var i=e.inspect(n,t);return v(i)||(i=u(t,i,n)),i}var a=function(t,e){if(y(e))return t.stylize("undefined","undefined");if(v(e)){var r="'"+JSON.stringify(e).replace(/^"|"$/g,"").replace(/'/g,"\\'").replace(/\\"/g,'"')+"'";return t.stylize(r,"string")}if(g(e))return t.stylize(""+e,"number");if(d(e))return t.stylize(""+e,"boolean");if(m(e))return t.stylize("null","null")}(t,e);if(a)return a;var o=Object.keys(e),s=function(t){var e={};return t.forEach((function(t,r){e[t]=!0})),e}(o);if(t.showHidden&&(o=Object.getOwnPropertyNames(e)),w(e)&&(o.indexOf("message")>=0||o.indexOf("description")>=0))return f(e);if(0===o.length){if(T(e)){var l=e.name?": "+e.name:"";return t.stylize("[Function"+l+"]","special")}if(x(e))return t.stylize(RegExp.prototype.toString.call(e),"regexp");if(_(e))return t.stylize(Date.prototype.toString.call(e),"date");if(w(e))return f(e)}var c,b="",k=!1,A=["{","}"];(p(e)&&(k=!0,A=["[","]"]),T(e))&&(b=" [Function"+(e.name?": "+e.name:"")+"]");return x(e)&&(b=" "+RegExp.prototype.toString.call(e)),_(e)&&(b=" "+Date.prototype.toUTCString.call(e)),w(e)&&(b=" "+f(e)),0!==o.length||k&&0!=e.length?n<0?x(e)?t.stylize(RegExp.prototype.toString.call(e),"regexp"):t.stylize("[Object]","special"):(t.seen.push(e),c=k?function(t,e,r,n,i){for(var a=[],o=0,s=e.length;o<s;++o)E(e,String(o))?a.push(h(t,e,r,n,String(o),!0)):a.push("");return i.forEach((function(i){i.match(/^\d+$/)||a.push(h(t,e,r,n,i,!0))})),a}(t,e,n,s,o):o.map((function(r){return h(t,e,n,s,r,k)})),t.seen.pop(),function(t,e,r){if(t.reduce((function(t,e){return e.indexOf("\n")>=0&&0,t+e.replace(/\u001b\[\d\d?m/g,"").length+1}),0)>60)return r[0]+(""===e?"":e+"\n ")+" "+t.join(",\n  ")+" "+r[1];return r[0]+e+" "+t.join(", ")+" "+r[1]}(c,b,A)):A[0]+b+A[1]}function f(t){return"["+Error.prototype.toString.call(t)+"]"}function h(t,e,r,n,i,a){var o,s,l;if((l=Object.getOwnPropertyDescriptor(e,i)||{value:e[i]}).get?s=l.set?t.stylize("[Getter/Setter]","special"):t.stylize("[Getter]","special"):l.set&&(s=t.stylize("[Setter]","special")),E(n,i)||(o="["+i+"]"),s||(t.seen.indexOf(l.value)<0?(s=m(r)?u(t,l.value,null):u(t,l.value,r-1)).indexOf("\n")>-1&&(s=a?s.split("\n").map((function(t){return"  "+t})).join("\n").substr(2):"\n"+s.split("\n").map((function(t){return"   "+t})).join("\n")):s=t.stylize("[Circular]","special")),y(o)){if(a&&i.match(/^\d+$/))return s;(o=JSON.stringify(""+i)).match(/^"([a-zA-Z_][a-zA-Z_0-9]*)"$/)?(o=o.substr(1,o.length-2),o=t.stylize(o,"name")):(o=o.replace(/'/g,"\\'").replace(/\\"/g,'"').replace(/(^"|"$)/g,"'"),o=t.stylize(o,"string"))}return o+": "+s}function p(t){return Array.isArray(t)}function d(t){return"boolean"==typeof t}function m(t){return null===t}function g(t){return"number"==typeof t}function v(t){return"string"==typeof t}function y(t){return void 0===t}function x(t){return b(t)&&"[object RegExp]"===k(t)}function b(t){return"object"==typeof t&&null!==t}function _(t){return b(t)&&"[object Date]"===k(t)}function w(t){return b(t)&&("[object Error]"===k(t)||t instanceof Error)}function T(t){return"function"==typeof t}function k(t){return Object.prototype.toString.call(t)}function A(t){return t<10?"0"+t.toString(10):t.toString(10)}r.debuglog=function(t){if(y(a)&&(a=e.env.NODE_DEBUG||""),t=t.toUpperCase(),!o[t])if(new RegExp("\\b"+t+"\\b","i").test(a)){var n=e.pid;o[t]=function(){var e=r.format.apply(r,arguments);console.error("%s %d: %s",t,n,e)}}else o[t]=function(){};return o[t]},r.inspect=s,s.colors={bold:[1,22],italic:[3,23],underline:[4,24],inverse:[7,27],white:[37,39],grey:[90,39],black:[30,39],blue:[34,39],cyan:[36,39],green:[32,39],magenta:[35,39],red:[31,39],yellow:[33,39]},s.styles={special:"cyan",number:"yellow",boolean:"yellow",undefined:"grey",null:"bold",string:"green",date:"magenta",regexp:"red"},r.isArray=p,r.isBoolean=d,r.isNull=m,r.isNullOrUndefined=function(t){return null==t},r.isNumber=g,r.isString=v,r.isSymbol=function(t){return"symbol"==typeof t},r.isUndefined=y,r.isRegExp=x,r.isObject=b,r.isDate=_,r.isError=w,r.isFunction=T,r.isPrimitive=function(t){return null===t||"boolean"==typeof t||"number"==typeof t||"string"==typeof t||"symbol"==typeof t||void 0===t},r.isBuffer=t("./support/isBuffer");var M=["Jan","Feb","Mar","Apr","May","Jun","Jul","Aug","Sep","Oct","Nov","Dec"];function S(){var t=new Date,e=[A(t.getHours()),A(t.getMinutes()),A(t.getSeconds())].join(":");return[t.getDate(),M[t.getMonth()],e].join(" ")}function E(t,e){return Object.prototype.hasOwnProperty.call(t,e)}r.log=function(){console.log("%s - %s",S(),r.format.apply(r,arguments))},r.inherits=t("inherits"),r._extend=function(t,e){if(!e||!b(e))return t;for(var r=Object.keys(e),n=r.length;n--;)t[r[n]]=e[r[n]];return t}}).call(this)}).call(this,t("_process"),"undefined"!=typeof global?global:"undefined"!=typeof self?self:"undefined"!=typeof window?window:{})},{"./support/isBuffer":73,_process:272,inherits:72}],75:[function(t,e,r){"use strict";r.byteLength=function(t){var e=c(t),r=e[0],n=e[1];return 3*(r+n)/4-n},r.toByteArray=function(t){var e,r,n=c(t),o=n[0],s=n[1],l=new a(function(t,e,r){return 3*(e+r)/4-r}(0,o,s)),u=0,f=s>0?o-4:o;for(r=0;r<f;r+=4)e=i[t.charCodeAt(r)]<<18|i[t.charCodeAt(r+1)]<<12|i[t.charCodeAt(r+2)]<<6|i[t.charCodeAt(r+3)],l[u++]=e>>16&255,l[u++]=e>>8&255,l[u++]=255&e;2===s&&(e=i[t.charCodeAt(r)]<<2|i[t.charCodeAt(r+1)]>>4,l[u++]=255&e);1===s&&(e=i[t.charCodeAt(r)]<<10|i[t.charCodeAt(r+1)]<<4|i[t.charCodeAt(r+2)]>>2,l[u++]=e>>8&255,l[u++]=255&e);return l},r.fromByteArray=function(t){for(var e,r=t.length,i=r%3,a=[],o=0,s=r-i;o<s;o+=16383)a.push(u(t,o,o+16383>s?s:o+16383));1===i?(e=t[r-1],a.push(n[e>>2]+n[e<<4&63]+"==")):2===i&&(e=(t[r-2]<<8)+t[r-1],a.push(n[e>>10]+n[e>>4&63]+n[e<<2&63]+"="));return a.join("")};for(var n=[],i=[],a="undefined"!=typeof Uint8Array?Uint8Array:Array,o="ABCDEFGHIJKLMNOPQRSTUVWXYZabcdefghijklmnopqrstuvwxyz0123456789+/",s=0,l=o.length;s<l;++s)n[s]=o[s],i[o.charCodeAt(s)]=s;function c(t){var e=t.length;if(e%4>0)throw new Error("Invalid string. Length must be a multiple of 4");var r=t.indexOf("=");return-1===r&&(r=e),[r,r===e?0:4-r%4]}function u(t,e,r){for(var i,a,o=[],s=e;s<r;s+=3)i=(t[s]<<16&16711680)+(t[s+1]<<8&65280)+(255&t[s+2]),o.push(n[(a=i)>>18&63]+n[a>>12&63]+n[a>>6&63]+n[63&a]);return o.join("")}i["-".charCodeAt(0)]=62,i["_".charCodeAt(0)]=63},{}],76:[function(t,e,r){"use strict";function n(t,e,r,n,i){for(var a=i+1;n<=i;){var o=n+i>>>1,s=t[o];(void 0!==r?r(s,e):s-e)>=0?(a=o,i=o-1):n=o+1}return a}function i(t,e,r,n,i){for(var a=i+1;n<=i;){var o=n+i>>>1,s=t[o];(void 0!==r?r(s,e):s-e)>0?(a=o,i=o-1):n=o+1}return a}function a(t,e,r,n,i){for(var a=n-1;n<=i;){var o=n+i>>>1,s=t[o];(void 0!==r?r(s,e):s-e)<0?(a=o,n=o+1):i=o-1}return a}function o(t,e,r,n,i){for(var a=n-1;n<=i;){var o=n+i>>>1,s=t[o];(void 0!==r?r(s,e):s-e)<=0?(a=o,n=o+1):i=o-1}return a}function s(t,e,r,n,i){for(;n<=i;){var a=n+i>>>1,o=t[a],s=void 0!==r?r(o,e):o-e;if(0===s)return a;s<=0?n=a+1:i=a-1}return-1}function l(t,e,r,n,i,a){return"function"==typeof r?a(t,e,r,void 0===n?0:0|n,void 0===i?t.length-1:0|i):a(t,e,void 0,void 0===r?0:0|r,void 0===n?t.length-1:0|n)}e.exports={ge:function(t,e,r,i,a){return l(t,e,r,i,a,n)},gt:function(t,e,r,n,a){return l(t,e,r,n,a,i)},lt:function(t,e,r,n,i){return l(t,e,r,n,i,a)},le:function(t,e,r,n,i){return l(t,e,r,n,i,o)},eq:function(t,e,r,n,i){return l(t,e,r,n,i,s)}}},{}],77:[function(t,e,r){"use strict";function n(t){var e=32;return(t&=-t)&&e--,65535&t&&(e-=16),16711935&t&&(e-=8),252645135&t&&(e-=4),858993459&t&&(e-=2),1431655765&t&&(e-=1),e}r.INT_BITS=32,r.INT_MAX=2147483647,r.INT_MIN=-1<<31,r.sign=function(t){return(t>0)-(t<0)},r.abs=function(t){var e=t>>31;return(t^e)-e},r.min=function(t,e){return e^(t^e)&-(t<e)},r.max=function(t,e){return t^(t^e)&-(t<e)},r.isPow2=function(t){return!(t&t-1||!t)},r.log2=function(t){var e,r;return e=(t>65535)<<4,e|=r=((t>>>=e)>255)<<3,e|=r=((t>>>=r)>15)<<2,(e|=r=((t>>>=r)>3)<<1)|(t>>>=r)>>1},r.log10=function(t){return t>=1e9?9:t>=1e8?8:t>=1e7?7:t>=1e6?6:t>=1e5?5:t>=1e4?4:t>=1e3?3:t>=100?2:t>=10?1:0},r.popCount=function(t){return 16843009*((t=(858993459&(t-=t>>>1&1431655765))+(t>>>2&858993459))+(t>>>4)&252645135)>>>24},r.countTrailingZeros=n,r.nextPow2=function(t){return t+=0===t,--t,t|=t>>>1,t|=t>>>2,t|=t>>>4,t|=t>>>8,(t|=t>>>16)+1},r.prevPow2=function(t){return t|=t>>>1,t|=t>>>2,t|=t>>>4,t|=t>>>8,(t|=t>>>16)-(t>>>1)},r.parity=function(t){return t^=t>>>16,t^=t>>>8,t^=t>>>4,27030>>>(t&=15)&1};var i=new Array(256);!function(t){for(var e=0;e<256;++e){var r=e,n=e,i=7;for(r>>>=1;r;r>>>=1)n<<=1,n|=1&r,--i;t[e]=n<<i&255}}(i),r.reverse=function(t){return i[255&t]<<24|i[t>>>8&255]<<16|i[t>>>16&255]<<8|i[t>>>24&255]},r.interleave2=function(t,e){return(t=1431655765&((t=858993459&((t=252645135&((t=16711935&((t&=65535)|t<<8))|t<<4))|t<<2))|t<<1))|(e=1431655765&((e=858993459&((e=252645135&((e=16711935&((e&=65535)|e<<8))|e<<4))|e<<2))|e<<1))<<1},r.deinterleave2=function(t,e){return(t=65535&((t=16711935&((t=252645135&((t=858993459&((t=t>>>e&1431655765)|t>>>1))|t>>>2))|t>>>4))|t>>>16))<<16>>16},r.interleave3=function(t,e,r){return t=1227133513&((t=3272356035&((t=251719695&((t=4278190335&((t&=1023)|t<<16))|t<<8))|t<<4))|t<<2),(t|=(e=1227133513&((e=3272356035&((e=251719695&((e=4278190335&((e&=1023)|e<<16))|e<<8))|e<<4))|e<<2))<<1)|(r=1227133513&((r=3272356035&((r=251719695&((r=4278190335&((r&=1023)|r<<16))|r<<8))|r<<4))|r<<2))<<2},r.deinterleave3=function(t,e){return(t=1023&((t=4278190335&((t=251719695&((t=3272356035&((t=t>>>e&1227133513)|t>>>2))|t>>>4))|t>>>8))|t>>>16))<<22>>22},r.nextCombination=function(t){var e=t|t-1;return e+1|(~e&-~e)-1>>>n(t)+1}},{}],78:[function(t,e,r){"use strict";var n=t("clamp");e.exports=function(t,e){e||(e={});var r,o,s,l,c,u,f,h,p,d,m,g=null==e.cutoff?.25:e.cutoff,v=null==e.radius?8:e.radius,y=e.channel||0;if(ArrayBuffer.isView(t)||Array.isArray(t)){if(!e.width||!e.height)throw Error("For raw data width and height should be provided by options");r=e.width,o=e.height,l=t,u=e.stride?e.stride:Math.floor(t.length/r/o)}else window.HTMLCanvasElement&&t instanceof window.HTMLCanvasElement?(f=(h=t).getContext("2d"),r=h.width,o=h.height,p=f.getImageData(0,0,r,o),l=p.data,u=4):window.CanvasRenderingContext2D&&t instanceof window.CanvasRenderingContext2D?(h=t.canvas,f=t,r=h.width,o=h.height,p=f.getImageData(0,0,r,o),l=p.data,u=4):window.ImageData&&t instanceof window.ImageData&&(p=t,r=t.width,o=t.height,l=p.data,u=4);if(s=Math.max(r,o),window.Uint8ClampedArray&&l instanceof window.Uint8ClampedArray||window.Uint8Array&&l instanceof window.Uint8Array)for(c=l,l=Array(r*o),d=0,m=c.length;d<m;d++)l[d]=c[d*u+y]/255;else if(1!==u)throw Error("Raw data can have only 1 value per pixel");var x=Array(r*o),b=Array(r*o),_=Array(s),w=Array(s),T=Array(s+1),k=Array(s);for(d=0,m=r*o;d<m;d++){var A=l[d];x[d]=1===A?0:0===A?i:Math.pow(Math.max(0,.5-A),2),b[d]=1===A?i:0===A?0:Math.pow(Math.max(0,A-.5),2)}a(x,r,o,_,w,k,T),a(b,r,o,_,w,k,T);var M=window.Float32Array?new Float32Array(r*o):new Array(r*o);for(d=0,m=r*o;d<m;d++)M[d]=n(1-((x[d]-b[d])/v+g),0,1);return M};var i=1e20;function a(t,e,r,n,i,a,s){for(var l=0;l<e;l++){for(var c=0;c<r;c++)n[c]=t[c*e+l];for(o(n,i,a,s,r),c=0;c<r;c++)t[c*e+l]=i[c]}for(c=0;c<r;c++){for(l=0;l<e;l++)n[l]=t[c*e+l];for(o(n,i,a,s,e),l=0;l<e;l++)t[c*e+l]=Math.sqrt(i[l])}}function o(t,e,r,n,a){r[0]=0,n[0]=-i,n[1]=+i;for(var o=1,s=0;o<a;o++){for(var l=(t[o]+o*o-(t[r[s]]+r[s]*r[s]))/(2*o-2*r[s]);l<=n[s];)s--,l=(t[o]+o*o-(t[r[s]]+r[s]*r[s]))/(2*o-2*r[s]);r[++s]=o,n[s]=l,n[s+1]=+i}for(o=0,s=0;o<a;o++){for(;n[s+1]<o;)s++;e[o]=(o-r[s])*(o-r[s])+t[r[s]]}}},{clamp:81}],79:[function(t,e,r){},{}],80:[function(t,e,r){(function(e){(function(){
/*!
 * The buffer module from node.js, for the browser.
 *
 * @author   Feross Aboukhadijeh <https://feross.org>
 * @license  MIT
 */
"use strict";var e=t("base64-js"),n=t("ieee754");r.Buffer=a,r.SlowBuffer=function(t){+t!=t&&(t=0);return a.alloc(+t)},r.INSPECT_MAX_BYTES=50;function i(t){if(t>2147483647)throw new RangeError('The value "'+t+'" is invalid for option "size"');var e=new Uint8Array(t);return e.__proto__=a.prototype,e}function a(t,e,r){if("number"==typeof t){if("string"==typeof e)throw new TypeError('The "string" argument must be of type string. Received type number');return l(t)}return o(t,e,r)}function o(t,e,r){if("string"==typeof t)return function(t,e){"string"==typeof e&&""!==e||(e="utf8");if(!a.isEncoding(e))throw new TypeError("Unknown encoding: "+e);var r=0|f(t,e),n=i(r),o=n.write(t,e);o!==r&&(n=n.slice(0,o));return n}(t,e);if(ArrayBuffer.isView(t))return c(t);if(null==t)throw TypeError("The first argument must be one of type string, Buffer, ArrayBuffer, Array, or Array-like Object. Received type "+typeof t);if(B(t,ArrayBuffer)||t&&B(t.buffer,ArrayBuffer))return function(t,e,r){if(e<0||t.byteLength<e)throw new RangeError('"offset" is outside of buffer bounds');if(t.byteLength<e+(r||0))throw new RangeError('"length" is outside of buffer bounds');var n;n=void 0===e&&void 0===r?new Uint8Array(t):void 0===r?new Uint8Array(t,e):new Uint8Array(t,e,r);return n.__proto__=a.prototype,n}(t,e,r);if("number"==typeof t)throw new TypeError('The "value" argument must not be of type number. Received type number');var n=t.valueOf&&t.valueOf();if(null!=n&&n!==t)return a.from(n,e,r);var o=function(t){if(a.isBuffer(t)){var e=0|u(t.length),r=i(e);return 0===r.length||t.copy(r,0,0,e),r}if(void 0!==t.length)return"number"!=typeof t.length||N(t.length)?i(0):c(t);if("Buffer"===t.type&&Array.isArray(t.data))return c(t.data)}(t);if(o)return o;if("undefined"!=typeof Symbol&&null!=Symbol.toPrimitive&&"function"==typeof t[Symbol.toPrimitive])return a.from(t[Symbol.toPrimitive]("string"),e,r);throw new TypeError("The first argument must be one of type string, Buffer, ArrayBuffer, Array, or Array-like Object. Received type "+typeof t)}function s(t){if("number"!=typeof t)throw new TypeError('"size" argument must be of type number');if(t<0)throw new RangeError('The value "'+t+'" is invalid for option "size"')}function l(t){return s(t),i(t<0?0:0|u(t))}function c(t){for(var e=t.length<0?0:0|u(t.length),r=i(e),n=0;n<e;n+=1)r[n]=255&t[n];return r}function u(t){if(t>=2147483647)throw new RangeError("Attempt to allocate Buffer larger than maximum size: 0x"+2147483647..toString(16)+" bytes");return 0|t}function f(t,e){if(a.isBuffer(t))return t.length;if(ArrayBuffer.isView(t)||B(t,ArrayBuffer))return t.byteLength;if("string"!=typeof t)throw new TypeError('The "string" argument must be one of type string, Buffer, or ArrayBuffer. Received type '+typeof t);var r=t.length,n=arguments.length>2&&!0===arguments[2];if(!n&&0===r)return 0;for(var i=!1;;)switch(e){case"ascii":case"latin1":case"binary":return r;case"utf8":case"utf-8":return D(t).length;case"ucs2":case"ucs-2":case"utf16le":case"utf-16le":return 2*r;case"hex":return r>>>1;case"base64":return R(t).length;default:if(i)return n?-1:D(t).length;e=(""+e).toLowerCase(),i=!0}}function h(t,e,r){var n=!1;if((void 0===e||e<0)&&(e=0),e>this.length)return"";if((void 0===r||r>this.length)&&(r=this.length),r<=0)return"";if((r>>>=0)<=(e>>>=0))return"";for(t||(t="utf8");;)switch(t){case"hex":return M(this,e,r);case"utf8":case"utf-8":return T(this,e,r);case"ascii":return k(this,e,r);case"latin1":case"binary":return A(this,e,r);case"base64":return w(this,e,r);case"ucs2":case"ucs-2":case"utf16le":case"utf-16le":return S(this,e,r);default:if(n)throw new TypeError("Unknown encoding: "+t);t=(t+"").toLowerCase(),n=!0}}function p(t,e,r){var n=t[e];t[e]=t[r],t[r]=n}function d(t,e,r,n,i){if(0===t.length)return-1;if("string"==typeof r?(n=r,r=0):r>2147483647?r=2147483647:r<-2147483648&&(r=-2147483648),N(r=+r)&&(r=i?0:t.length-1),r<0&&(r=t.length+r),r>=t.length){if(i)return-1;r=t.length-1}else if(r<0){if(!i)return-1;r=0}if("string"==typeof e&&(e=a.from(e,n)),a.isBuffer(e))return 0===e.length?-1:m(t,e,r,n,i);if("number"==typeof e)return e&=255,"function"==typeof Uint8Array.prototype.indexOf?i?Uint8Array.prototype.indexOf.call(t,e,r):Uint8Array.prototype.lastIndexOf.call(t,e,r):m(t,[e],r,n,i);throw new TypeError("val must be string, number or Buffer")}function m(t,e,r,n,i){var a,o=1,s=t.length,l=e.length;if(void 0!==n&&("ucs2"===(n=String(n).toLowerCase())||"ucs-2"===n||"utf16le"===n||"utf-16le"===n)){if(t.length<2||e.length<2)return-1;o=2,s/=2,l/=2,r/=2}function c(t,e){return 1===o?t[e]:t.readUInt16BE(e*o)}if(i){var u=-1;for(a=r;a<s;a++)if(c(t,a)===c(e,-1===u?0:a-u)){if(-1===u&&(u=a),a-u+1===l)return u*o}else-1!==u&&(a-=a-u),u=-1}else for(r+l>s&&(r=s-l),a=r;a>=0;a--){for(var f=!0,h=0;h<l;h++)if(c(t,a+h)!==c(e,h)){f=!1;break}if(f)return a}return-1}function g(t,e,r,n){r=Number(r)||0;var i=t.length-r;n?(n=Number(n))>i&&(n=i):n=i;var a=e.length;n>a/2&&(n=a/2);for(var o=0;o<n;++o){var s=parseInt(e.substr(2*o,2),16);if(N(s))return o;t[r+o]=s}return o}function v(t,e,r,n){return F(D(e,t.length-r),t,r,n)}function y(t,e,r,n){return F(function(t){for(var e=[],r=0;r<t.length;++r)e.push(255&t.charCodeAt(r));return e}(e),t,r,n)}function x(t,e,r,n){return y(t,e,r,n)}function b(t,e,r,n){return F(R(e),t,r,n)}function _(t,e,r,n){return F(function(t,e){for(var r,n,i,a=[],o=0;o<t.length&&!((e-=2)<0);++o)r=t.charCodeAt(o),n=r>>8,i=r%256,a.push(i),a.push(n);return a}(e,t.length-r),t,r,n)}function w(t,r,n){return 0===r&&n===t.length?e.fromByteArray(t):e.fromByteArray(t.slice(r,n))}function T(t,e,r){r=Math.min(t.length,r);for(var n=[],i=e;i<r;){var a,o,s,l,c=t[i],u=null,f=c>239?4:c>223?3:c>191?2:1;if(i+f<=r)switch(f){case 1:c<128&&(u=c);break;case 2:128==(192&(a=t[i+1]))&&(l=(31&c)<<6|63&a)>127&&(u=l);break;case 3:a=t[i+1],o=t[i+2],128==(192&a)&&128==(192&o)&&(l=(15&c)<<12|(63&a)<<6|63&o)>2047&&(l<55296||l>57343)&&(u=l);break;case 4:a=t[i+1],o=t[i+2],s=t[i+3],128==(192&a)&&128==(192&o)&&128==(192&s)&&(l=(15&c)<<18|(63&a)<<12|(63&o)<<6|63&s)>65535&&l<1114112&&(u=l)}null===u?(u=65533,f=1):u>65535&&(u-=65536,n.push(u>>>10&1023|55296),u=56320|1023&u),n.push(u),i+=f}return function(t){var e=t.length;if(e<=4096)return String.fromCharCode.apply(String,t);var r="",n=0;for(;n<e;)r+=String.fromCharCode.apply(String,t.slice(n,n+=4096));return r}(n)}r.kMaxLength=2147483647,a.TYPED_ARRAY_SUPPORT=function(){try{var t=new Uint8Array(1);return t.__proto__={__proto__:Uint8Array.prototype,foo:function(){return 42}},42===t.foo()}catch(t){return!1}}(),a.TYPED_ARRAY_SUPPORT||"undefined"==typeof console||"function"!=typeof console.error||console.error("This browser lacks typed array (Uint8Array) support which is required by `buffer` v5.x. Use `buffer` v4.x if you require old browser support."),Object.defineProperty(a.prototype,"parent",{enumerable:!0,get:function(){if(a.isBuffer(this))return this.buffer}}),Object.defineProperty(a.prototype,"offset",{enumerable:!0,get:function(){if(a.isBuffer(this))return this.byteOffset}}),"undefined"!=typeof Symbol&&null!=Symbol.species&&a[Symbol.species]===a&&Object.defineProperty(a,Symbol.species,{value:null,configurable:!0,enumerable:!1,writable:!1}),a.poolSize=8192,a.from=function(t,e,r){return o(t,e,r)},a.prototype.__proto__=Uint8Array.prototype,a.__proto__=Uint8Array,a.alloc=function(t,e,r){return function(t,e,r){return s(t),t<=0?i(t):void 0!==e?"string"==typeof r?i(t).fill(e,r):i(t).fill(e):i(t)}(t,e,r)},a.allocUnsafe=function(t){return l(t)},a.allocUnsafeSlow=function(t){return l(t)},a.isBuffer=function(t){return null!=t&&!0===t._isBuffer&&t!==a.prototype},a.compare=function(t,e){if(B(t,Uint8Array)&&(t=a.from(t,t.offset,t.byteLength)),B(e,Uint8Array)&&(e=a.from(e,e.offset,e.byteLength)),!a.isBuffer(t)||!a.isBuffer(e))throw new TypeError('The "buf1", "buf2" arguments must be one of type Buffer or Uint8Array');if(t===e)return 0;for(var r=t.length,n=e.length,i=0,o=Math.min(r,n);i<o;++i)if(t[i]!==e[i]){r=t[i],n=e[i];break}return r<n?-1:n<r?1:0},a.isEncoding=function(t){switch(String(t).toLowerCase()){case"hex":case"utf8":case"utf-8":case"ascii":case"latin1":case"binary":case"base64":case"ucs2":case"ucs-2":case"utf16le":case"utf-16le":return!0;default:return!1}},a.concat=function(t,e){if(!Array.isArray(t))throw new TypeError('"list" argument must be an Array of Buffers');if(0===t.length)return a.alloc(0);var r;if(void 0===e)for(e=0,r=0;r<t.length;++r)e+=t[r].length;var n=a.allocUnsafe(e),i=0;for(r=0;r<t.length;++r){var o=t[r];if(B(o,Uint8Array)&&(o=a.from(o)),!a.isBuffer(o))throw new TypeError('"list" argument must be an Array of Buffers');o.copy(n,i),i+=o.length}return n},a.byteLength=f,a.prototype._isBuffer=!0,a.prototype.swap16=function(){var t=this.length;if(t%2!=0)throw new RangeError("Buffer size must be a multiple of 16-bits");for(var e=0;e<t;e+=2)p(this,e,e+1);return this},a.prototype.swap32=function(){var t=this.length;if(t%4!=0)throw new RangeError("Buffer size must be a multiple of 32-bits");for(var e=0;e<t;e+=4)p(this,e,e+3),p(this,e+1,e+2);return this},a.prototype.swap64=function(){var t=this.length;if(t%8!=0)throw new RangeError("Buffer size must be a multiple of 64-bits");for(var e=0;e<t;e+=8)p(this,e,e+7),p(this,e+1,e+6),p(this,e+2,e+5),p(this,e+3,e+4);return this},a.prototype.toString=function(){var t=this.length;return 0===t?"":0===arguments.length?T(this,0,t):h.apply(this,arguments)},a.prototype.toLocaleString=a.prototype.toString,a.prototype.equals=function(t){if(!a.isBuffer(t))throw new TypeError("Argument must be a Buffer");return this===t||0===a.compare(this,t)},a.prototype.inspect=function(){var t="",e=r.INSPECT_MAX_BYTES;return t=this.toString("hex",0,e).replace(/(.{2})/g,"$1 ").trim(),this.length>e&&(t+=" ... "),"<Buffer "+t+">"},a.prototype.compare=function(t,e,r,n,i){if(B(t,Uint8Array)&&(t=a.from(t,t.offset,t.byteLength)),!a.isBuffer(t))throw new TypeError('The "target" argument must be one of type Buffer or Uint8Array. Received type '+typeof t);if(void 0===e&&(e=0),void 0===r&&(r=t?t.length:0),void 0===n&&(n=0),void 0===i&&(i=this.length),e<0||r>t.length||n<0||i>this.length)throw new RangeError("out of range index");if(n>=i&&e>=r)return 0;if(n>=i)return-1;if(e>=r)return 1;if(this===t)return 0;for(var o=(i>>>=0)-(n>>>=0),s=(r>>>=0)-(e>>>=0),l=Math.min(o,s),c=this.slice(n,i),u=t.slice(e,r),f=0;f<l;++f)if(c[f]!==u[f]){o=c[f],s=u[f];break}return o<s?-1:s<o?1:0},a.prototype.includes=function(t,e,r){return-1!==this.indexOf(t,e,r)},a.prototype.indexOf=function(t,e,r){return d(this,t,e,r,!0)},a.prototype.lastIndexOf=function(t,e,r){return d(this,t,e,r,!1)},a.prototype.write=function(t,e,r,n){if(void 0===e)n="utf8",r=this.length,e=0;else if(void 0===r&&"string"==typeof e)n=e,r=this.length,e=0;else{if(!isFinite(e))throw new Error("Buffer.write(string, encoding, offset[, length]) is no longer supported");e>>>=0,isFinite(r)?(r>>>=0,void 0===n&&(n="utf8")):(n=r,r=void 0)}var i=this.length-e;if((void 0===r||r>i)&&(r=i),t.length>0&&(r<0||e<0)||e>this.length)throw new RangeError("Attempt to write outside buffer bounds");n||(n="utf8");for(var a=!1;;)switch(n){case"hex":return g(this,t,e,r);case"utf8":case"utf-8":return v(this,t,e,r);case"ascii":return y(this,t,e,r);case"latin1":case"binary":return x(this,t,e,r);case"base64":return b(this,t,e,r);case"ucs2":case"ucs-2":case"utf16le":case"utf-16le":return _(this,t,e,r);default:if(a)throw new TypeError("Unknown encoding: "+n);n=(""+n).toLowerCase(),a=!0}},a.prototype.toJSON=function(){return{type:"Buffer",data:Array.prototype.slice.call(this._arr||this,0)}};function k(t,e,r){var n="";r=Math.min(t.length,r);for(var i=e;i<r;++i)n+=String.fromCharCode(127&t[i]);return n}function A(t,e,r){var n="";r=Math.min(t.length,r);for(var i=e;i<r;++i)n+=String.fromCharCode(t[i]);return n}function M(t,e,r){var n=t.length;(!e||e<0)&&(e=0),(!r||r<0||r>n)&&(r=n);for(var i="",a=e;a<r;++a)i+=z(t[a]);return i}function S(t,e,r){for(var n=t.slice(e,r),i="",a=0;a<n.length;a+=2)i+=String.fromCharCode(n[a]+256*n[a+1]);return i}function E(t,e,r){if(t%1!=0||t<0)throw new RangeError("offset is not uint");if(t+e>r)throw new RangeError("Trying to access beyond buffer length")}function L(t,e,r,n,i,o){if(!a.isBuffer(t))throw new TypeError('"buffer" argument must be a Buffer instance');if(e>i||e<o)throw new RangeError('"value" argument is out of bounds');if(r+n>t.length)throw new RangeError("Index out of range")}function C(t,e,r,n,i,a){if(r+n>t.length)throw new RangeError("Index out of range");if(r<0)throw new RangeError("Index out of range")}function P(t,e,r,i,a){return e=+e,r>>>=0,a||C(t,0,r,4),n.write(t,e,r,i,23,4),r+4}function I(t,e,r,i,a){return e=+e,r>>>=0,a||C(t,0,r,8),n.write(t,e,r,i,52,8),r+8}a.prototype.slice=function(t,e){var r=this.length;(t=~~t)<0?(t+=r)<0&&(t=0):t>r&&(t=r),(e=void 0===e?r:~~e)<0?(e+=r)<0&&(e=0):e>r&&(e=r),e<t&&(e=t);var n=this.subarray(t,e);return n.__proto__=a.prototype,n},a.prototype.readUIntLE=function(t,e,r){t>>>=0,e>>>=0,r||E(t,e,this.length);for(var n=this[t],i=1,a=0;++a<e&&(i*=256);)n+=this[t+a]*i;return n},a.prototype.readUIntBE=function(t,e,r){t>>>=0,e>>>=0,r||E(t,e,this.length);for(var n=this[t+--e],i=1;e>0&&(i*=256);)n+=this[t+--e]*i;return n},a.prototype.readUInt8=function(t,e){return t>>>=0,e||E(t,1,this.length),this[t]},a.prototype.readUInt16LE=function(t,e){return t>>>=0,e||E(t,2,this.length),this[t]|this[t+1]<<8},a.prototype.readUInt16BE=function(t,e){return t>>>=0,e||E(t,2,this.length),this[t]<<8|this[t+1]},a.prototype.readUInt32LE=function(t,e){return t>>>=0,e||E(t,4,this.length),(this[t]|this[t+1]<<8|this[t+2]<<16)+16777216*this[t+3]},a.prototype.readUInt32BE=function(t,e){return t>>>=0,e||E(t,4,this.length),16777216*this[t]+(this[t+1]<<16|this[t+2]<<8|this[t+3])},a.prototype.readIntLE=function(t,e,r){t>>>=0,e>>>=0,r||E(t,e,this.length);for(var n=this[t],i=1,a=0;++a<e&&(i*=256);)n+=this[t+a]*i;return n>=(i*=128)&&(n-=Math.pow(2,8*e)),n},a.prototype.readIntBE=function(t,e,r){t>>>=0,e>>>=0,r||E(t,e,this.length);for(var n=e,i=1,a=this[t+--n];n>0&&(i*=256);)a+=this[t+--n]*i;return a>=(i*=128)&&(a-=Math.pow(2,8*e)),a},a.prototype.readInt8=function(t,e){return t>>>=0,e||E(t,1,this.length),128&this[t]?-1*(255-this[t]+1):this[t]},a.prototype.readInt16LE=function(t,e){t>>>=0,e||E(t,2,this.length);var r=this[t]|this[t+1]<<8;return 32768&r?4294901760|r:r},a.prototype.readInt16BE=function(t,e){t>>>=0,e||E(t,2,this.length);var r=this[t+1]|this[t]<<8;return 32768&r?4294901760|r:r},a.prototype.readInt32LE=function(t,e){return t>>>=0,e||E(t,4,this.length),this[t]|this[t+1]<<8|this[t+2]<<16|this[t+3]<<24},a.prototype.readInt32BE=function(t,e){return t>>>=0,e||E(t,4,this.length),this[t]<<24|this[t+1]<<16|this[t+2]<<8|this[t+3]},a.prototype.readFloatLE=function(t,e){return t>>>=0,e||E(t,4,this.length),n.read(this,t,!0,23,4)},a.prototype.readFloatBE=function(t,e){return t>>>=0,e||E(t,4,this.length),n.read(this,t,!1,23,4)},a.prototype.readDoubleLE=function(t,e){return t>>>=0,e||E(t,8,this.length),n.read(this,t,!0,52,8)},a.prototype.readDoubleBE=function(t,e){return t>>>=0,e||E(t,8,this.length),n.read(this,t,!1,52,8)},a.prototype.writeUIntLE=function(t,e,r,n){(t=+t,e>>>=0,r>>>=0,n)||L(this,t,e,r,Math.pow(2,8*r)-1,0);var i=1,a=0;for(this[e]=255&t;++a<r&&(i*=256);)this[e+a]=t/i&255;return e+r},a.prototype.writeUIntBE=function(t,e,r,n){(t=+t,e>>>=0,r>>>=0,n)||L(this,t,e,r,Math.pow(2,8*r)-1,0);var i=r-1,a=1;for(this[e+i]=255&t;--i>=0&&(a*=256);)this[e+i]=t/a&255;return e+r},a.prototype.writeUInt8=function(t,e,r){return t=+t,e>>>=0,r||L(this,t,e,1,255,0),this[e]=255&t,e+1},a.prototype.writeUInt16LE=function(t,e,r){return t=+t,e>>>=0,r||L(this,t,e,2,65535,0),this[e]=255&t,this[e+1]=t>>>8,e+2},a.prototype.writeUInt16BE=function(t,e,r){return t=+t,e>>>=0,r||L(this,t,e,2,65535,0),this[e]=t>>>8,this[e+1]=255&t,e+2},a.prototype.writeUInt32LE=function(t,e,r){return t=+t,e>>>=0,r||L(this,t,e,4,4294967295,0),this[e+3]=t>>>24,this[e+2]=t>>>16,this[e+1]=t>>>8,this[e]=255&t,e+4},a.prototype.writeUInt32BE=function(t,e,r){return t=+t,e>>>=0,r||L(this,t,e,4,4294967295,0),this[e]=t>>>24,this[e+1]=t>>>16,this[e+2]=t>>>8,this[e+3]=255&t,e+4},a.prototype.writeIntLE=function(t,e,r,n){if(t=+t,e>>>=0,!n){var i=Math.pow(2,8*r-1);L(this,t,e,r,i-1,-i)}var a=0,o=1,s=0;for(this[e]=255&t;++a<r&&(o*=256);)t<0&&0===s&&0!==this[e+a-1]&&(s=1),this[e+a]=(t/o>>0)-s&255;return e+r},a.prototype.writeIntBE=function(t,e,r,n){if(t=+t,e>>>=0,!n){var i=Math.pow(2,8*r-1);L(this,t,e,r,i-1,-i)}var a=r-1,o=1,s=0;for(this[e+a]=255&t;--a>=0&&(o*=256);)t<0&&0===s&&0!==this[e+a+1]&&(s=1),this[e+a]=(t/o>>0)-s&255;return e+r},a.prototype.writeInt8=function(t,e,r){return t=+t,e>>>=0,r||L(this,t,e,1,127,-128),t<0&&(t=255+t+1),this[e]=255&t,e+1},a.prototype.writeInt16LE=function(t,e,r){return t=+t,e>>>=0,r||L(this,t,e,2,32767,-32768),this[e]=255&t,this[e+1]=t>>>8,e+2},a.prototype.writeInt16BE=function(t,e,r){return t=+t,e>>>=0,r||L(this,t,e,2,32767,-32768),this[e]=t>>>8,this[e+1]=255&t,e+2},a.prototype.writeInt32LE=function(t,e,r){return t=+t,e>>>=0,r||L(this,t,e,4,2147483647,-2147483648),this[e]=255&t,this[e+1]=t>>>8,this[e+2]=t>>>16,this[e+3]=t>>>24,e+4},a.prototype.writeInt32BE=function(t,e,r){return t=+t,e>>>=0,r||L(this,t,e,4,2147483647,-2147483648),t<0&&(t=4294967295+t+1),this[e]=t>>>24,this[e+1]=t>>>16,this[e+2]=t>>>8,this[e+3]=255&t,e+4},a.prototype.writeFloatLE=function(t,e,r){return P(this,t,e,!0,r)},a.prototype.writeFloatBE=function(t,e,r){return P(this,t,e,!1,r)},a.prototype.writeDoubleLE=function(t,e,r){return I(this,t,e,!0,r)},a.prototype.writeDoubleBE=function(t,e,r){return I(this,t,e,!1,r)},a.prototype.copy=function(t,e,r,n){if(!a.isBuffer(t))throw new TypeError("argument should be a Buffer");if(r||(r=0),n||0===n||(n=this.length),e>=t.length&&(e=t.length),e||(e=0),n>0&&n<r&&(n=r),n===r)return 0;if(0===t.length||0===this.length)return 0;if(e<0)throw new RangeError("targetStart out of bounds");if(r<0||r>=this.length)throw new RangeError("Index out of range");if(n<0)throw new RangeError("sourceEnd out of bounds");n>this.length&&(n=this.length),t.length-e<n-r&&(n=t.length-e+r);var i=n-r;if(this===t&&"function"==typeof Uint8Array.prototype.copyWithin)this.copyWithin(e,r,n);else if(this===t&&r<e&&e<n)for(var o=i-1;o>=0;--o)t[o+e]=this[o+r];else Uint8Array.prototype.set.call(t,this.subarray(r,n),e);return i},a.prototype.fill=function(t,e,r,n){if("string"==typeof t){if("string"==typeof e?(n=e,e=0,r=this.length):"string"==typeof r&&(n=r,r=this.length),void 0!==n&&"string"!=typeof n)throw new TypeError("encoding must be a string");if("string"==typeof n&&!a.isEncoding(n))throw new TypeError("Unknown encoding: "+n);if(1===t.length){var i=t.charCodeAt(0);("utf8"===n&&i<128||"latin1"===n)&&(t=i)}}else"number"==typeof t&&(t&=255);if(e<0||this.length<e||this.length<r)throw new RangeError("Out of range index");if(r<=e)return this;var o;if(e>>>=0,r=void 0===r?this.length:r>>>0,t||(t=0),"number"==typeof t)for(o=e;o<r;++o)this[o]=t;else{var s=a.isBuffer(t)?t:a.from(t,n),l=s.length;if(0===l)throw new TypeError('The value "'+t+'" is invalid for argument "value"');for(o=0;o<r-e;++o)this[o+e]=s[o%l]}return this};var O=/[^+/0-9A-Za-z-_]/g;function z(t){return t<16?"0"+t.toString(16):t.toString(16)}function D(t,e){var r;e=e||1/0;for(var n=t.length,i=null,a=[],o=0;o<n;++o){if((r=t.charCodeAt(o))>55295&&r<57344){if(!i){if(r>56319){(e-=3)>-1&&a.push(239,191,189);continue}if(o+1===n){(e-=3)>-1&&a.push(239,191,189);continue}i=r;continue}if(r<56320){(e-=3)>-1&&a.push(239,191,189),i=r;continue}r=65536+(i-55296<<10|r-56320)}else i&&(e-=3)>-1&&a.push(239,191,189);if(i=null,r<128){if((e-=1)<0)break;a.push(r)}else if(r<2048){if((e-=2)<0)break;a.push(r>>6|192,63&r|128)}else if(r<65536){if((e-=3)<0)break;a.push(r>>12|224,r>>6&63|128,63&r|128)}else{if(!(r<1114112))throw new Error("Invalid code point");if((e-=4)<0)break;a.push(r>>18|240,r>>12&63|128,r>>6&63|128,63&r|128)}}return a}function R(t){return e.toByteArray(function(t){if((t=(t=t.split("=")[0]).trim().replace(O,"")).length<2)return"";for(;t.length%4!=0;)t+="=";return t}(t))}function F(t,e,r,n){for(var i=0;i<n&&!(i+r>=e.length||i>=t.length);++i)e[i+r]=t[i];return i}function B(t,e){return t instanceof e||null!=t&&null!=t.constructor&&null!=t.constructor.name&&t.constructor.name===e.name}function N(t){return t!=t}}).call(this)}).call(this,t("buffer").Buffer)},{"base64-js":75,buffer:80,ieee754:225}],81:[function(t,e,r){e.exports=function(t,e,r){return e<r?t<e?e:t>r?r:t:t<r?r:t>e?e:t}},{}],82:[function(t,e,r){"use strict";var n=t("clamp");function i(t,e){null==e&&(e=!0);var r=t[0],i=t[1],a=t[2],o=t[3];return null==o&&(o=e?1:255),e&&(r*=255,i*=255,a*=255,o*=255),16777216*(r=255&n(r,0,255))+((i=255&n(i,0,255))<<16)+((a=255&n(a,0,255))<<8)+(o=255&n(o,0,255))}e.exports=i,e.exports.to=i,e.exports.from=function(t,e){var r=(t=+t)>>>24,n=(16711680&t)>>>16,i=(65280&t)>>>8,a=255&t;return!1===e?[r,n,i,a]:[r/255,n/255,i/255,a/255]}},{clamp:81}],83:[function(t,e,r){"use strict";e.exports={aliceblue:[240,248,255],antiquewhite:[250,235,215],aqua:[0,255,255],aquamarine:[127,255,212],azure:[240,255,255],beige:[245,245,220],bisque:[255,228,196],black:[0,0,0],blanchedalmond:[255,235,205],blue:[0,0,255],blueviolet:[138,43,226],brown:[165,42,42],burlywood:[222,184,135],cadetblue:[95,158,160],chartreuse:[127,255,0],chocolate:[210,105,30],coral:[255,127,80],cornflowerblue:[100,149,237],cornsilk:[255,248,220],crimson:[220,20,60],cyan:[0,255,255],darkblue:[0,0,139],darkcyan:[0,139,139],darkgoldenrod:[184,134,11],darkgray:[169,169,169],darkgreen:[0,100,0],darkgrey:[169,169,169],darkkhaki:[189,183,107],darkmagenta:[139,0,139],darkolivegreen:[85,107,47],darkorange:[255,140,0],darkorchid:[153,50,204],darkred:[139,0,0],darksalmon:[233,150,122],darkseagreen:[143,188,143],darkslateblue:[72,61,139],darkslategray:[47,79,79],darkslategrey:[47,79,79],darkturquoise:[0,206,209],darkviolet:[148,0,211],deeppink:[255,20,147],deepskyblue:[0,191,255],dimgray:[105,105,105],dimgrey:[105,105,105],dodgerblue:[30,144,255],firebrick:[178,34,34],floralwhite:[255,250,240],forestgreen:[34,139,34],fuchsia:[255,0,255],gainsboro:[220,220,220],ghostwhite:[248,248,255],gold:[255,215,0],goldenrod:[218,165,32],gray:[128,128,128],green:[0,128,0],greenyellow:[173,255,47],grey:[128,128,128],honeydew:[240,255,240],hotpink:[255,105,180],indianred:[205,92,92],indigo:[75,0,130],ivory:[255,255,240],khaki:[240,230,140],lavender:[230,230,250],lavenderblush:[255,240,245],lawngreen:[124,252,0],lemonchiffon:[255,250,205],lightblue:[173,216,230],lightcoral:[240,128,128],lightcyan:[224,255,255],lightgoldenrodyellow:[250,250,210],lightgray:[211,211,211],lightgreen:[144,238,144],lightgrey:[211,211,211],lightpink:[255,182,193],lightsalmon:[255,160,122],lightseagreen:[32,178,170],lightskyblue:[135,206,250],lightslategray:[119,136,153],lightslategrey:[119,136,153],lightsteelblue:[176,196,222],lightyellow:[255,255,224],lime:[0,255,0],limegreen:[50,205,50],linen:[250,240,230],magenta:[255,0,255],maroon:[128,0,0],mediumaquamarine:[102,205,170],mediumblue:[0,0,205],mediumorchid:[186,85,211],mediumpurple:[147,112,219],mediumseagreen:[60,179,113],mediumslateblue:[123,104,238],mediumspringgreen:[0,250,154],mediumturquoise:[72,209,204],mediumvioletred:[199,21,133],midnightblue:[25,25,112],mintcream:[245,255,250],mistyrose:[255,228,225],moccasin:[255,228,181],navajowhite:[255,222,173],navy:[0,0,128],oldlace:[253,245,230],olive:[128,128,0],olivedrab:[107,142,35],orange:[255,165,0],orangered:[255,69,0],orchid:[218,112,214],palegoldenrod:[238,232,170],palegreen:[152,251,152],paleturquoise:[175,238,238],palevioletred:[219,112,147],papayawhip:[255,239,213],peachpuff:[255,218,185],peru:[205,133,63],pink:[255,192,203],plum:[221,160,221],powderblue:[176,224,230],purple:[128,0,128],rebeccapurple:[102,51,153],red:[255,0,0],rosybrown:[188,143,143],royalblue:[65,105,225],saddlebrown:[139,69,19],salmon:[250,128,114],sandybrown:[244,164,96],seagreen:[46,139,87],seashell:[255,245,238],sienna:[160,82,45],silver:[192,192,192],skyblue:[135,206,235],slateblue:[106,90,205],slategray:[112,128,144],slategrey:[112,128,144],snow:[255,250,250],springgreen:[0,255,127],steelblue:[70,130,180],tan:[210,180,140],teal:[0,128,128],thistle:[216,191,216],tomato:[255,99,71],turquoise:[64,224,208],violet:[238,130,238],wheat:[245,222,179],white:[255,255,255],whitesmoke:[245,245,245],yellow:[255,255,0],yellowgreen:[154,205,50]}},{}],84:[function(t,e,r){"use strict";var n=t("color-rgba"),i=t("clamp"),a=t("dtype");e.exports=function(t,e){"float"!==e&&e||(e="array"),"uint"===e&&(e="uint8"),"uint_clamped"===e&&(e="uint8_clamped");var r=new(a(e))(4),o="uint8"!==e&&"uint8_clamped"!==e;return t.length&&"string"!=typeof t||((t=n(t))[0]/=255,t[1]/=255,t[2]/=255),function(t){return t instanceof Uint8Array||t instanceof Uint8ClampedArray||!!(Array.isArray(t)&&(t[0]>1||0===t[0])&&(t[1]>1||0===t[1])&&(t[2]>1||0===t[2])&&(!t[3]||t[3]>1))}(t)?(r[0]=t[0],r[1]=t[1],r[2]=t[2],r[3]=null!=t[3]?t[3]:255,o&&(r[0]/=255,r[1]/=255,r[2]/=255,r[3]/=255),r):(o?(r[0]=t[0],r[1]=t[1],r[2]=t[2],r[3]=null!=t[3]?t[3]:1):(r[0]=i(Math.floor(255*t[0]),0,255),r[1]=i(Math.floor(255*t[1]),0,255),r[2]=i(Math.floor(255*t[2]),0,255),r[3]=null==t[3]?255:i(Math.floor(255*t[3]),0,255)),r)}},{clamp:81,"color-rgba":86,dtype:121}],85:[function(t,e,r){(function(r){(function(){"use strict";var n=t("color-name"),i=t("is-plain-obj"),a=t("defined");e.exports=function(t){var e,s,l=[],c=1;if("string"==typeof t)if(n[t])l=n[t].slice(),s="rgb";else if("transparent"===t)c=0,s="rgb",l=[0,0,0];else if(/^#[A-Fa-f0-9]+$/.test(t)){var u=(p=t.slice(1)).length;c=1,u<=4?(l=[parseInt(p[0]+p[0],16),parseInt(p[1]+p[1],16),parseInt(p[2]+p[2],16)],4===u&&(c=parseInt(p[3]+p[3],16)/255)):(l=[parseInt(p[0]+p[1],16),parseInt(p[2]+p[3],16),parseInt(p[4]+p[5],16)],8===u&&(c=parseInt(p[6]+p[7],16)/255)),l[0]||(l[0]=0),l[1]||(l[1]=0),l[2]||(l[2]=0),s="rgb"}else if(e=/^((?:rgb|hs[lvb]|hwb|cmyk?|xy[zy]|gray|lab|lchu?v?|[ly]uv|lms)a?)\s*\(([^\)]*)\)/.exec(t)){var f=e[1],h="rgb"===f,p=f.replace(/a$/,"");s=p;u="cmyk"===p?4:"gray"===p?1:3;l=e[2].trim().split(/\s*,\s*/).map((function(t,e){if(/%$/.test(t))return e===u?parseFloat(t)/100:"rgb"===p?255*parseFloat(t)/100:parseFloat(t);if("h"===p[e]){if(/deg$/.test(t))return parseFloat(t);if(void 0!==o[t])return o[t]}return parseFloat(t)})),f===p&&l.push(1),c=h||void 0===l[u]?1:l[u],l=l.slice(0,u)}else t.length>10&&/[0-9](?:\s|\/)/.test(t)&&(l=t.match(/([0-9]+)/g).map((function(t){return parseFloat(t)})),s=t.match(/([a-z])/gi).join("").toLowerCase());else if(isNaN(t))if(i(t)){var d=a(t.r,t.red,t.R,null);null!==d?(s="rgb",l=[d,a(t.g,t.green,t.G),a(t.b,t.blue,t.B)]):(s="hsl",l=[a(t.h,t.hue,t.H),a(t.s,t.saturation,t.S),a(t.l,t.lightness,t.L,t.b,t.brightness)]),c=a(t.a,t.alpha,t.opacity,1),null!=t.opacity&&(c/=100)}else(Array.isArray(t)||r.ArrayBuffer&&ArrayBuffer.isView&&ArrayBuffer.isView(t))&&(l=[t[0],t[1],t[2]],s="rgb",c=4===t.length?t[3]:1);else s="rgb",l=[t>>>16,(65280&t)>>>8,255&t];return{space:s,values:l,alpha:c}};var o={red:0,orange:60,yellow:120,green:180,blue:240,purple:300}}).call(this)}).call(this,"undefined"!=typeof global?global:"undefined"!=typeof self?self:"undefined"!=typeof window?window:{})},{"color-name":83,defined:118,"is-plain-obj":231}],86:[function(t,e,r){"use strict";var n=t("color-parse"),i=t("color-space/hsl"),a=t("clamp");e.exports=function(t){var e,r=n(t);return r.space?((e=Array(3))[0]=a(r.values[0],0,255),e[1]=a(r.values[1],0,255),e[2]=a(r.values[2],0,255),"h"===r.space[0]&&(e=i.rgb(e)),e.push(a(r.alpha,0,1)),e):[]}},{clamp:81,"color-parse":85,"color-space/hsl":87}],87:[function(t,e,r){"use strict";var n=t("./rgb");e.exports={name:"hsl",min:[0,0,0],max:[360,100,100],channel:["hue","saturation","lightness"],alias:["HSL"],rgb:function(t){var e,r,n,i,a,o=t[0]/360,s=t[1]/100,l=t[2]/100;if(0===s)return[a=255*l,a,a];e=2*l-(r=l<.5?l*(1+s):l+s-l*s),i=[0,0,0];for(var c=0;c<3;c++)(n=o+1/3*-(c-1))<0?n++:n>1&&n--,a=6*n<1?e+6*(r-e)*n:2*n<1?r:3*n<2?e+(r-e)*(2/3-n)*6:e,i[c]=255*a;return i}},n.hsl=function(t){var e,r,n=t[0]/255,i=t[1]/255,a=t[2]/255,o=Math.min(n,i,a),s=Math.max(n,i,a),l=s-o;return s===o?e=0:n===s?e=(i-a)/l:i===s?e=2+(a-n)/l:a===s&&(e=4+(n-i)/l),(e=Math.min(60*e,360))<0&&(e+=360),r=(o+s)/2,[e,100*(s===o?0:r<=.5?l/(s+o):l/(2-s-o)),100*r]}},{"./rgb":88}],88:[function(t,e,r){"use strict";e.exports={name:"rgb",min:[0,0,0],max:[255,255,255],channel:["red","green","blue"],alias:["RGB"]}},{}],89:[function(t,e,r){e.exports={AFG:"afghan",ALA:"\\b\\wland",ALB:"albania",DZA:"algeria",ASM:"^(?=.*americ).*samoa",AND:"andorra",AGO:"angola",AIA:"anguill?a",ATA:"antarctica",ATG:"antigua",ARG:"argentin",ARM:"armenia",ABW:"^(?!.*bonaire).*\\baruba",AUS:"australia",AUT:"^(?!.*hungary).*austria|\\baustri.*\\bemp",AZE:"azerbaijan",BHS:"bahamas",BHR:"bahrain",BGD:"bangladesh|^(?=.*east).*paki?stan",BRB:"barbados",BLR:"belarus|byelo",BEL:"^(?!.*luxem).*belgium",BLZ:"belize|^(?=.*british).*honduras",BEN:"benin|dahome",BMU:"bermuda",BTN:"bhutan",BOL:"bolivia",BES:"^(?=.*bonaire).*eustatius|^(?=.*carib).*netherlands|\\bbes.?islands",BIH:"herzegovina|bosnia",BWA:"botswana|bechuana",BVT:"bouvet",BRA:"brazil",IOT:"british.?indian.?ocean",BRN:"brunei",BGR:"bulgaria",BFA:"burkina|\\bfaso|upper.?volta",BDI:"burundi",CPV:"verde",KHM:"cambodia|kampuchea|khmer",CMR:"cameroon",CAN:"canada",CYM:"cayman",CAF:"\\bcentral.african.republic",TCD:"\\bchad",CHL:"\\bchile",CHN:"^(?!.*\\bmac)(?!.*\\bhong)(?!.*\\btai)(?!.*\\brep).*china|^(?=.*peo)(?=.*rep).*china",CXR:"christmas",CCK:"\\bcocos|keeling",COL:"colombia",COM:"comoro",COG:"^(?!.*\\bdem)(?!.*\\bd[\\.]?r)(?!.*kinshasa)(?!.*zaire)(?!.*belg)(?!.*l.opoldville)(?!.*free).*\\bcongo",COK:"\\bcook",CRI:"costa.?rica",CIV:"ivoire|ivory",HRV:"croatia",CUB:"\\bcuba",CUW:"^(?!.*bonaire).*\\bcura(c|\xe7)ao",CYP:"cyprus",CSK:"czechoslovakia",CZE:"^(?=.*rep).*czech|czechia|bohemia",COD:"\\bdem.*congo|congo.*\\bdem|congo.*\\bd[\\.]?r|\\bd[\\.]?r.*congo|belgian.?congo|congo.?free.?state|kinshasa|zaire|l.opoldville|drc|droc|rdc",DNK:"denmark",DJI:"djibouti",DMA:"dominica(?!n)",DOM:"dominican.rep",ECU:"ecuador",EGY:"egypt",SLV:"el.?salvador",GNQ:"guine.*eq|eq.*guine|^(?=.*span).*guinea",ERI:"eritrea",EST:"estonia",ETH:"ethiopia|abyssinia",FLK:"falkland|malvinas",FRO:"faroe|faeroe",FJI:"fiji",FIN:"finland",FRA:"^(?!.*\\bdep)(?!.*martinique).*france|french.?republic|\\bgaul",GUF:"^(?=.*french).*guiana",PYF:"french.?polynesia|tahiti",ATF:"french.?southern",GAB:"gabon",GMB:"gambia",GEO:"^(?!.*south).*georgia",DDR:"german.?democratic.?republic|democratic.?republic.*germany|east.germany",DEU:"^(?!.*east).*germany|^(?=.*\\bfed.*\\brep).*german",GHA:"ghana|gold.?coast",GIB:"gibraltar",GRC:"greece|hellenic|hellas",GRL:"greenland",GRD:"grenada",GLP:"guadeloupe",GUM:"\\bguam",GTM:"guatemala",GGY:"guernsey",GIN:"^(?!.*eq)(?!.*span)(?!.*bissau)(?!.*portu)(?!.*new).*guinea",GNB:"bissau|^(?=.*portu).*guinea",GUY:"guyana|british.?guiana",HTI:"haiti",HMD:"heard.*mcdonald",VAT:"holy.?see|vatican|papal.?st",HND:"^(?!.*brit).*honduras",HKG:"hong.?kong",HUN:"^(?!.*austr).*hungary",ISL:"iceland",IND:"india(?!.*ocea)",IDN:"indonesia",IRN:"\\biran|persia",IRQ:"\\biraq|mesopotamia",IRL:"(^ireland)|(^republic.*ireland)",IMN:"^(?=.*isle).*\\bman",ISR:"israel",ITA:"italy",JAM:"jamaica",JPN:"japan",JEY:"jersey",JOR:"jordan",KAZ:"kazak",KEN:"kenya|british.?east.?africa|east.?africa.?prot",KIR:"kiribati",PRK:"^(?=.*democrat|people|north|d.*p.*.r).*\\bkorea|dprk|korea.*(d.*p.*r)",KWT:"kuwait",KGZ:"kyrgyz|kirghiz",LAO:"\\blaos?\\b",LVA:"latvia",LBN:"lebanon",LSO:"lesotho|basuto",LBR:"liberia",LBY:"libya",LIE:"liechtenstein",LTU:"lithuania",LUX:"^(?!.*belg).*luxem",MAC:"maca(o|u)",MDG:"madagascar|malagasy",MWI:"malawi|nyasa",MYS:"malaysia",MDV:"maldive",MLI:"\\bmali\\b",MLT:"\\bmalta",MHL:"marshall",MTQ:"martinique",MRT:"mauritania",MUS:"mauritius",MYT:"\\bmayotte",MEX:"\\bmexic",FSM:"fed.*micronesia|micronesia.*fed",MCO:"monaco",MNG:"mongolia",MNE:"^(?!.*serbia).*montenegro",MSR:"montserrat",MAR:"morocco|\\bmaroc",MOZ:"mozambique",MMR:"myanmar|burma",NAM:"namibia",NRU:"nauru",NPL:"nepal",NLD:"^(?!.*\\bant)(?!.*\\bcarib).*netherlands",ANT:"^(?=.*\\bant).*(nether|dutch)",NCL:"new.?caledonia",NZL:"new.?zealand",NIC:"nicaragua",NER:"\\bniger(?!ia)",NGA:"nigeria",NIU:"niue",NFK:"norfolk",MNP:"mariana",NOR:"norway",OMN:"\\boman|trucial",PAK:"^(?!.*east).*paki?stan",PLW:"palau",PSE:"palestin|\\bgaza|west.?bank",PAN:"panama",PNG:"papua|new.?guinea",PRY:"paraguay",PER:"peru",PHL:"philippines",PCN:"pitcairn",POL:"poland",PRT:"portugal",PRI:"puerto.?rico",QAT:"qatar",KOR:"^(?!.*d.*p.*r)(?!.*democrat)(?!.*people)(?!.*north).*\\bkorea(?!.*d.*p.*r)",MDA:"moldov|b(a|e)ssarabia",REU:"r(e|\xe9)union",ROU:"r(o|u|ou)mania",RUS:"\\brussia|soviet.?union|u\\.?s\\.?s\\.?r|socialist.?republics",RWA:"rwanda",BLM:"barth(e|\xe9)lemy",SHN:"helena",KNA:"kitts|\\bnevis",LCA:"\\blucia",MAF:"^(?=.*collectivity).*martin|^(?=.*france).*martin(?!ique)|^(?=.*french).*martin(?!ique)",SPM:"miquelon",VCT:"vincent",WSM:"^(?!.*amer).*samoa",SMR:"san.?marino",STP:"\\bs(a|\xe3)o.?tom(e|\xe9)",SAU:"\\bsa\\w*.?arabia",SEN:"senegal",SRB:"^(?!.*monte).*serbia",SYC:"seychell",SLE:"sierra",SGP:"singapore",SXM:"^(?!.*martin)(?!.*saba).*maarten",SVK:"^(?!.*cze).*slovak",SVN:"slovenia",SLB:"solomon",SOM:"somali",ZAF:"south.africa|s\\\\..?africa",SGS:"south.?georgia|sandwich",SSD:"\\bs\\w*.?sudan",ESP:"spain",LKA:"sri.?lanka|ceylon",SDN:"^(?!.*\\bs(?!u)).*sudan",SUR:"surinam|dutch.?guiana",SJM:"svalbard",SWZ:"swaziland",SWE:"sweden",CHE:"switz|swiss",SYR:"syria",TWN:"taiwan|taipei|formosa|^(?!.*peo)(?=.*rep).*china",TJK:"tajik",THA:"thailand|\\bsiam",MKD:"macedonia|fyrom",TLS:"^(?=.*leste).*timor|^(?=.*east).*timor",TGO:"togo",TKL:"tokelau",TON:"tonga",TTO:"trinidad|tobago",TUN:"tunisia",TUR:"turkey",TKM:"turkmen",TCA:"turks",TUV:"tuvalu",UGA:"uganda",UKR:"ukrain",ARE:"emirates|^u\\.?a\\.?e\\.?$|united.?arab.?em",GBR:"united.?kingdom|britain|^u\\.?k\\.?$",TZA:"tanzania",USA:"united.?states\\b(?!.*islands)|\\bu\\.?s\\.?a\\.?\\b|^\\s*u\\.?s\\.?\\b(?!.*islands)",UMI:"minor.?outlying.?is",URY:"uruguay",UZB:"uzbek",VUT:"vanuatu|new.?hebrides",VEN:"venezuela",VNM:"^(?!.*republic).*viet.?nam|^(?=.*socialist).*viet.?nam",VGB:"^(?=.*\\bu\\.?\\s?k).*virgin|^(?=.*brit).*virgin|^(?=.*kingdom).*virgin",VIR:"^(?=.*\\bu\\.?\\s?s).*virgin|^(?=.*states).*virgin",WLF:"futuna|wallis",ESH:"western.sahara",YEM:"^(?!.*arab)(?!.*north)(?!.*sana)(?!.*peo)(?!.*dem)(?!.*south)(?!.*aden)(?!.*\\bp\\.?d\\.?r).*yemen",YMD:"^(?=.*peo).*yemen|^(?!.*rep)(?=.*dem).*yemen|^(?=.*south).*yemen|^(?=.*aden).*yemen|^(?=.*\\bp\\.?d\\.?r).*yemen",YUG:"yugoslavia",ZMB:"zambia|northern.?rhodesia",EAZ:"zanzibar",ZWE:"zimbabwe|^(?!.*northern).*rhodesia"}},{}],90:[function(t,e,r){e.exports=["xx-small","x-small","small","medium","large","x-large","xx-large","larger","smaller"]},{}],91:[function(t,e,r){e.exports=["normal","condensed","semi-condensed","extra-condensed","ultra-condensed","expanded","semi-expanded","extra-expanded","ultra-expanded"]},{}],92:[function(t,e,r){e.exports=["normal","italic","oblique"]},{}],93:[function(t,e,r){e.exports=["normal","bold","bolder","lighter","100","200","300","400","500","600","700","800","900"]},{}],94:[function(t,e,r){"use strict";e.exports={parse:t("./parse"),stringify:t("./stringify")}},{"./parse":96,"./stringify":97}],95:[function(t,e,r){"use strict";var n=t("css-font-size-keywords");e.exports={isSize:function(t){return/^[\d\.]/.test(t)||-1!==t.indexOf("/")||-1!==n.indexOf(t)}}},{"css-font-size-keywords":90}],96:[function(t,e,r){"use strict";var n=t("unquote"),i=t("css-global-keywords"),a=t("css-system-font-keywords"),o=t("css-font-weight-keywords"),s=t("css-font-style-keywords"),l=t("css-font-stretch-keywords"),c=t("string-split-by"),u=t("./lib/util").isSize;e.exports=h;var f=h.cache={};function h(t){if("string"!=typeof t)throw new Error("Font argument must be a string.");if(f[t])return f[t];if(""===t)throw new Error("Cannot parse an empty string.");if(-1!==a.indexOf(t))return f[t]={system:t};for(var e,r={style:"normal",variant:"normal",weight:"normal",stretch:"normal",lineHeight:"normal",size:"1rem",family:["serif"]},h=c(t,/\s+/);e=h.shift();){if(-1!==i.indexOf(e))return["style","variant","weight","stretch"].forEach((function(t){r[t]=e})),f[t]=r;if(-1===s.indexOf(e))if("normal"!==e&&"small-caps"!==e)if(-1===l.indexOf(e)){if(-1===o.indexOf(e)){if(u(e)){var d=c(e,"/");if(r.size=d[0],null!=d[1]?r.lineHeight=p(d[1]):"/"===h[0]&&(h.shift(),r.lineHeight=p(h.shift())),!h.length)throw new Error("Missing required font-family.");return r.family=c(h.join(" "),/\s*,\s*/).map(n),f[t]=r}throw new Error("Unknown or unsupported font token: "+e)}r.weight=e}else r.stretch=e;else r.variant=e;else r.style=e}throw new Error("Missing required font-size.")}function p(t){var e=parseFloat(t);return e.toString()===t?e:t}},{"./lib/util":95,"css-font-stretch-keywords":91,"css-font-style-keywords":92,"css-font-weight-keywords":93,"css-global-keywords":98,"css-system-font-keywords":99,"string-split-by":299,unquote:323}],97:[function(t,e,r){"use strict";var n=t("pick-by-alias"),i=t("./lib/util").isSize,a=m(t("css-global-keywords")),o=m(t("css-system-font-keywords")),s=m(t("css-font-weight-keywords")),l=m(t("css-font-style-keywords")),c=m(t("css-font-stretch-keywords")),u={normal:1,"small-caps":1},f={serif:1,"sans-serif":1,monospace:1,cursive:1,fantasy:1,"system-ui":1},h="1rem",p="serif";function d(t,e){if(t&&!e[t]&&!a[t])throw Error("Unknown keyword `"+t+"`");return t}function m(t){for(var e={},r=0;r<t.length;r++)e[t[r]]=1;return e}e.exports=function(t){if((t=n(t,{style:"style fontstyle fontStyle font-style slope distinction",variant:"variant font-variant fontVariant fontvariant var capitalization",weight:"weight w font-weight fontWeight fontweight",stretch:"stretch font-stretch fontStretch fontstretch width",size:"size s font-size fontSize fontsize height em emSize",lineHeight:"lh line-height lineHeight lineheight leading",family:"font family fontFamily font-family fontfamily type typeface face",system:"system reserved default global"})).system)return t.system&&d(t.system,o),t.system;if(d(t.style,l),d(t.variant,u),d(t.weight,s),d(t.stretch,c),null==t.size&&(t.size=h),"number"==typeof t.size&&(t.size+="px"),!i)throw Error("Bad size value `"+t.size+"`");t.family||(t.family=p),Array.isArray(t.family)&&(t.family.length||(t.family=[p]),t.family=t.family.map((function(t){return f[t]?t:'"'+t+'"'})).join(", "));var e=[];return e.push(t.style),t.variant!==t.style&&e.push(t.variant),t.weight!==t.variant&&t.weight!==t.style&&e.push(t.weight),t.stretch!==t.weight&&t.stretch!==t.variant&&t.stretch!==t.style&&e.push(t.stretch),e.push(t.size+(null==t.lineHeight||"normal"===t.lineHeight||t.lineHeight+""=="1"?"":"/"+t.lineHeight)),e.push(t.family),e.filter(Boolean).join(" ")}},{"./lib/util":95,"css-font-stretch-keywords":91,"css-font-style-keywords":92,"css-font-weight-keywords":93,"css-global-keywords":98,"css-system-font-keywords":99,"pick-by-alias":248}],98:[function(t,e,r){e.exports=["inherit","initial","unset"]},{}],99:[function(t,e,r){e.exports=["caption","icon","menu","message-box","small-caption","status-bar"]},{}],100:[function(t,e,r){"use strict";var n,i=t("type/value/is"),a=t("type/value/ensure"),o=t("type/plain-function/ensure"),s=t("es5-ext/object/copy"),l=t("es5-ext/object/normalize-options"),c=t("es5-ext/object/map"),u=Function.prototype.bind,f=Object.defineProperty,h=Object.prototype.hasOwnProperty;n=function(t,e,r){var n,i=a(e)&&o(e.value);return delete(n=s(e)).writable,delete n.value,n.get=function(){return!r.overwriteDefinition&&h.call(this,t)?i:(e.value=u.call(i,r.resolveContext?r.resolveContext(this):this),f(this,t,e),this[t])},n},e.exports=function(t){var e=l(arguments[1]);return i(e.resolveContext)&&o(e.resolveContext),c(t,(function(t,r){return n(r,t,e)}))}},{"es5-ext/object/copy":141,"es5-ext/object/map":149,"es5-ext/object/normalize-options":150,"type/plain-function/ensure":316,"type/value/ensure":320,"type/value/is":321}],101:[function(t,e,r){"use strict";var n=t("type/value/is"),i=t("type/plain-function/is"),a=t("es5-ext/object/assign"),o=t("es5-ext/object/normalize-options"),s=t("es5-ext/string/#/contains");(e.exports=function(t,e){var r,i,l,c,u;return arguments.length<2||"string"!=typeof t?(c=e,e=t,t=null):c=arguments[2],n(t)?(r=s.call(t,"c"),i=s.call(t,"e"),l=s.call(t,"w")):(r=l=!0,i=!1),u={value:e,configurable:r,enumerable:i,writable:l},c?a(o(c),u):u}).gs=function(t,e,r){var l,c,u,f;return"string"!=typeof t?(u=r,r=e,e=t,t=null):u=arguments[3],n(e)?i(e)?n(r)?i(r)||(u=r,r=void 0):r=void 0:(u=e,e=r=void 0):e=void 0,n(t)?(l=s.call(t,"c"),c=s.call(t,"e")):(l=!0,c=!1),f={get:e,set:r,configurable:l,enumerable:c},u?a(o(u),f):f}},{"es5-ext/object/assign":138,"es5-ext/object/normalize-options":150,"es5-ext/string/#/contains":157,"type/plain-function/is":317,"type/value/is":321}],102:[function(t,e,r){!function(t,n){n("object"==typeof r&&void 0!==e?r:t.d3=t.d3||{})}(this,(function(t){"use strict";function e(t,e){return t<e?-1:t>e?1:t>=e?0:NaN}function r(t){var r;return 1===t.length&&(r=t,t=function(t,n){return e(r(t),n)}),{left:function(e,r,n,i){for(null==n&&(n=0),null==i&&(i=e.length);n<i;){var a=n+i>>>1;t(e[a],r)<0?n=a+1:i=a}return n},right:function(e,r,n,i){for(null==n&&(n=0),null==i&&(i=e.length);n<i;){var a=n+i>>>1;t(e[a],r)>0?i=a:n=a+1}return n}}}var n=r(e),i=n.right,a=n.left;function o(t,e){return[t,e]}function s(t){return null===t?NaN:+t}function l(t,e){var r,n,i=t.length,a=0,o=-1,l=0,c=0;if(null==e)for(;++o<i;)isNaN(r=s(t[o]))||(c+=(n=r-l)*(r-(l+=n/++a)));else for(;++o<i;)isNaN(r=s(e(t[o],o,t)))||(c+=(n=r-l)*(r-(l+=n/++a)));if(a>1)return c/(a-1)}function c(t,e){var r=l(t,e);return r?Math.sqrt(r):r}function u(t,e){var r,n,i,a=t.length,o=-1;if(null==e){for(;++o<a;)if(null!=(r=t[o])&&r>=r)for(n=i=r;++o<a;)null!=(r=t[o])&&(n>r&&(n=r),i<r&&(i=r))}else for(;++o<a;)if(null!=(r=e(t[o],o,t))&&r>=r)for(n=i=r;++o<a;)null!=(r=e(t[o],o,t))&&(n>r&&(n=r),i<r&&(i=r));return[n,i]}var f=Array.prototype,h=f.slice,p=f.map;function d(t){return function(){return t}}function m(t){return t}function g(t,e,r){t=+t,e=+e,r=(i=arguments.length)<2?(e=t,t=0,1):i<3?1:+r;for(var n=-1,i=0|Math.max(0,Math.ceil((e-t)/r)),a=new Array(i);++n<i;)a[n]=t+n*r;return a}var v=Math.sqrt(50),y=Math.sqrt(10),x=Math.sqrt(2);function b(t,e,r){var n=(e-t)/Math.max(0,r),i=Math.floor(Math.log(n)/Math.LN10),a=n/Math.pow(10,i);return i>=0?(a>=v?10:a>=y?5:a>=x?2:1)*Math.pow(10,i):-Math.pow(10,-i)/(a>=v?10:a>=y?5:a>=x?2:1)}function _(t,e,r){var n=Math.abs(e-t)/Math.max(0,r),i=Math.pow(10,Math.floor(Math.log(n)/Math.LN10)),a=n/i;return a>=v?i*=10:a>=y?i*=5:a>=x&&(i*=2),e<t?-i:i}function w(t){return Math.ceil(Math.log(t.length)/Math.LN2)+1}function T(t,e,r){if(null==r&&(r=s),n=t.length){if((e=+e)<=0||n<2)return+r(t[0],0,t);if(e>=1)return+r(t[n-1],n-1,t);var n,i=(n-1)*e,a=Math.floor(i),o=+r(t[a],a,t);return o+(+r(t[a+1],a+1,t)-o)*(i-a)}}function k(t,e){var r,n,i=t.length,a=-1;if(null==e){for(;++a<i;)if(null!=(r=t[a])&&r>=r)for(n=r;++a<i;)null!=(r=t[a])&&n>r&&(n=r)}else for(;++a<i;)if(null!=(r=e(t[a],a,t))&&r>=r)for(n=r;++a<i;)null!=(r=e(t[a],a,t))&&n>r&&(n=r);return n}function A(t){if(!(i=t.length))return[];for(var e=-1,r=k(t,M),n=new Array(r);++e<r;)for(var i,a=-1,o=n[e]=new Array(i);++a<i;)o[a]=t[a][e];return n}function M(t){return t.length}t.bisect=i,t.bisectRight=i,t.bisectLeft=a,t.ascending=e,t.bisector=r,t.cross=function(t,e,r){var n,i,a,s,l=t.length,c=e.length,u=new Array(l*c);for(null==r&&(r=o),n=a=0;n<l;++n)for(s=t[n],i=0;i<c;++i,++a)u[a]=r(s,e[i]);return u},t.descending=function(t,e){return e<t?-1:e>t?1:e>=t?0:NaN},t.deviation=c,t.extent=u,t.histogram=function(){var t=m,e=u,r=w;function n(n){var a,o,s=n.length,l=new Array(s);for(a=0;a<s;++a)l[a]=t(n[a],a,n);var c=e(l),u=c[0],f=c[1],h=r(l,u,f);Array.isArray(h)||(h=_(u,f,h),h=g(Math.ceil(u/h)*h,f,h));for(var p=h.length;h[0]<=u;)h.shift(),--p;for(;h[p-1]>f;)h.pop(),--p;var d,m=new Array(p+1);for(a=0;a<=p;++a)(d=m[a]=[]).x0=a>0?h[a-1]:u,d.x1=a<p?h[a]:f;for(a=0;a<s;++a)u<=(o=l[a])&&o<=f&&m[i(h,o,0,p)].push(n[a]);return m}return n.value=function(e){return arguments.length?(t="function"==typeof e?e:d(e),n):t},n.domain=function(t){return arguments.length?(e="function"==typeof t?t:d([t[0],t[1]]),n):e},n.thresholds=function(t){return arguments.length?(r="function"==typeof t?t:Array.isArray(t)?d(h.call(t)):d(t),n):r},n},t.thresholdFreedmanDiaconis=function(t,r,n){return t=p.call(t,s).sort(e),Math.ceil((n-r)/(2*(T(t,.75)-T(t,.25))*Math.pow(t.length,-1/3)))},t.thresholdScott=function(t,e,r){return Math.ceil((r-e)/(3.5*c(t)*Math.pow(t.length,-1/3)))},t.thresholdSturges=w,t.max=function(t,e){var r,n,i=t.length,a=-1;if(null==e){for(;++a<i;)if(null!=(r=t[a])&&r>=r)for(n=r;++a<i;)null!=(r=t[a])&&r>n&&(n=r)}else for(;++a<i;)if(null!=(r=e(t[a],a,t))&&r>=r)for(n=r;++a<i;)null!=(r=e(t[a],a,t))&&r>n&&(n=r);return n},t.mean=function(t,e){var r,n=t.length,i=n,a=-1,o=0;if(null==e)for(;++a<n;)isNaN(r=s(t[a]))?--i:o+=r;else for(;++a<n;)isNaN(r=s(e(t[a],a,t)))?--i:o+=r;if(i)return o/i},t.median=function(t,r){var n,i=t.length,a=-1,o=[];if(null==r)for(;++a<i;)isNaN(n=s(t[a]))||o.push(n);else for(;++a<i;)isNaN(n=s(r(t[a],a,t)))||o.push(n);return T(o.sort(e),.5)},t.merge=function(t){for(var e,r,n,i=t.length,a=-1,o=0;++a<i;)o+=t[a].length;for(r=new Array(o);--i>=0;)for(e=(n=t[i]).length;--e>=0;)r[--o]=n[e];return r},t.min=k,t.pairs=function(t,e){null==e&&(e=o);for(var r=0,n=t.length-1,i=t[0],a=new Array(n<0?0:n);r<n;)a[r]=e(i,i=t[++r]);return a},t.permute=function(t,e){for(var r=e.length,n=new Array(r);r--;)n[r]=t[e[r]];return n},t.quantile=T,t.range=g,t.scan=function(t,r){if(n=t.length){var n,i,a=0,o=0,s=t[o];for(null==r&&(r=e);++a<n;)(r(i=t[a],s)<0||0!==r(s,s))&&(s=i,o=a);return 0===r(s,s)?o:void 0}},t.shuffle=function(t,e,r){for(var n,i,a=(null==r?t.length:r)-(e=null==e?0:+e);a;)i=Math.random()*a--|0,n=t[a+e],t[a+e]=t[i+e],t[i+e]=n;return t},t.sum=function(t,e){var r,n=t.length,i=-1,a=0;if(null==e)for(;++i<n;)(r=+t[i])&&(a+=r);else for(;++i<n;)(r=+e(t[i],i,t))&&(a+=r);return a},t.ticks=function(t,e,r){var n,i,a,o,s=-1;if(r=+r,(t=+t)===(e=+e)&&r>0)return[t];if((n=e<t)&&(i=t,t=e,e=i),0===(o=b(t,e,r))||!isFinite(o))return[];if(o>0)for(t=Math.ceil(t/o),e=Math.floor(e/o),a=new Array(i=Math.ceil(e-t+1));++s<i;)a[s]=(t+s)*o;else for(t=Math.floor(t*o),e=Math.ceil(e*o),a=new Array(i=Math.ceil(t-e+1));++s<i;)a[s]=(t-s)/o;return n&&a.reverse(),a},t.tickIncrement=b,t.tickStep=_,t.transpose=A,t.variance=l,t.zip=function(){return A(arguments)},Object.defineProperty(t,"__esModule",{value:!0})}))},{}],103:[function(t,e,r){!function(t,n){n("object"==typeof r&&void 0!==e?r:t.d3=t.d3||{})}(this,(function(t){"use strict";function e(){}function r(t,r){var n=new e;if(t instanceof e)t.each((function(t,e){n.set(e,t)}));else if(Array.isArray(t)){var i,a=-1,o=t.length;if(null==r)for(;++a<o;)n.set(a,t[a]);else for(;++a<o;)n.set(r(i=t[a],a,t),i)}else if(t)for(var s in t)n.set(s,t[s]);return n}function n(){return{}}function i(t,e,r){t[e]=r}function a(){return r()}function o(t,e,r){t.set(e,r)}function s(){}e.prototype=r.prototype={constructor:e,has:function(t){return"$"+t in this},get:function(t){return this["$"+t]},set:function(t,e){return this["$"+t]=e,this},remove:function(t){var e="$"+t;return e in this&&delete this[e]},clear:function(){for(var t in this)"$"===t[0]&&delete this[t]},keys:function(){var t=[];for(var e in this)"$"===e[0]&&t.push(e.slice(1));return t},values:function(){var t=[];for(var e in this)"$"===e[0]&&t.push(this[e]);return t},entries:function(){var t=[];for(var e in this)"$"===e[0]&&t.push({key:e.slice(1),value:this[e]});return t},size:function(){var t=0;for(var e in this)"$"===e[0]&&++t;return t},empty:function(){for(var t in this)if("$"===t[0])return!1;return!0},each:function(t){for(var e in this)"$"===e[0]&&t(this[e],e.slice(1),this)}};var l=r.prototype;function c(t,e){var r=new s;if(t instanceof s)t.each((function(t){r.add(t)}));else if(t){var n=-1,i=t.length;if(null==e)for(;++n<i;)r.add(t[n]);else for(;++n<i;)r.add(e(t[n],n,t))}return r}s.prototype=c.prototype={constructor:s,has:l.has,add:function(t){return this["$"+(t+="")]=t,this},remove:l.remove,clear:l.clear,values:l.keys,size:l.size,empty:l.empty,each:l.each},t.nest=function(){var t,e,s,l=[],c=[];function u(n,i,a,o){if(i>=l.length)return null!=t&&n.sort(t),null!=e?e(n):n;for(var s,c,f,h=-1,p=n.length,d=l[i++],m=r(),g=a();++h<p;)(f=m.get(s=d(c=n[h])+""))?f.push(c):m.set(s,[c]);return m.each((function(t,e){o(g,e,u(t,i,a,o))})),g}return s={object:function(t){return u(t,0,n,i)},map:function(t){return u(t,0,a,o)},entries:function(t){return function t(r,n){if(++n>l.length)return r;var i,a=c[n-1];return null!=e&&n>=l.length?i=r.entries():(i=[],r.each((function(e,r){i.push({key:r,values:t(e,n)})}))),null!=a?i.sort((function(t,e){return a(t.key,e.key)})):i}(u(t,0,a,o),0)},key:function(t){return l.push(t),s},sortKeys:function(t){return c[l.length-1]=t,s},sortValues:function(e){return t=e,s},rollup:function(t){return e=t,s}}},t.set=c,t.map=r,t.keys=function(t){var e=[];for(var r in t)e.push(r);return e},t.values=function(t){var e=[];for(var r in t)e.push(t[r]);return e},t.entries=function(t){var e=[];for(var r in t)e.push({key:r,value:t[r]});return e},Object.defineProperty(t,"__esModule",{value:!0})}))},{}],104:[function(t,e,r){!function(t,n){"object"==typeof r&&void 0!==e?n(r):n((t=t||self).d3=t.d3||{})}(this,(function(t){"use strict";function e(t,e,r){t.prototype=e.prototype=r,r.constructor=t}function r(t,e){var r=Object.create(t.prototype);for(var n in e)r[n]=e[n];return r}function n(){}var i="\\s*([+-]?\\d+)\\s*",a="\\s*([+-]?\\d*\\.?\\d+(?:[eE][+-]?\\d+)?)\\s*",o="\\s*([+-]?\\d*\\.?\\d+(?:[eE][+-]?\\d+)?)%\\s*",s=/^#([0-9a-f]{3,8})$/,l=new RegExp("^rgb\\("+[i,i,i]+"\\)$"),c=new RegExp("^rgb\\("+[o,o,o]+"\\)$"),u=new RegExp("^rgba\\("+[i,i,i,a]+"\\)$"),f=new RegExp("^rgba\\("+[o,o,o,a]+"\\)$"),h=new RegExp("^hsl\\("+[a,o,o]+"\\)$"),p=new RegExp("^hsla\\("+[a,o,o,a]+"\\)$"),d={aliceblue:15792383,antiquewhite:16444375,aqua:65535,aquamarine:8388564,azure:15794175,beige:16119260,bisque:16770244,black:0,blanchedalmond:16772045,blue:255,blueviolet:9055202,brown:10824234,burlywood:14596231,cadetblue:6266528,chartreuse:8388352,chocolate:13789470,coral:16744272,cornflowerblue:6591981,cornsilk:16775388,crimson:14423100,cyan:65535,darkblue:139,darkcyan:35723,darkgoldenrod:12092939,darkgray:11119017,darkgreen:25600,darkgrey:11119017,darkkhaki:12433259,darkmagenta:9109643,darkolivegreen:5597999,darkorange:16747520,darkorchid:10040012,darkred:9109504,darksalmon:15308410,darkseagreen:9419919,darkslateblue:4734347,darkslategray:3100495,darkslategrey:3100495,darkturquoise:52945,darkviolet:9699539,deeppink:16716947,deepskyblue:49151,dimgray:6908265,dimgrey:6908265,dodgerblue:2003199,firebrick:11674146,floralwhite:16775920,forestgreen:2263842,fuchsia:16711935,gainsboro:14474460,ghostwhite:16316671,gold:16766720,goldenrod:14329120,gray:8421504,green:32768,greenyellow:11403055,grey:8421504,honeydew:15794160,hotpink:16738740,indianred:13458524,indigo:4915330,ivory:16777200,khaki:15787660,lavender:15132410,lavenderblush:16773365,lawngreen:8190976,lemonchiffon:16775885,lightblue:11393254,lightcoral:15761536,lightcyan:14745599,lightgoldenrodyellow:16448210,lightgray:13882323,lightgreen:9498256,lightgrey:13882323,lightpink:16758465,lightsalmon:16752762,lightseagreen:2142890,lightskyblue:8900346,lightslategray:7833753,lightslategrey:7833753,lightsteelblue:11584734,lightyellow:16777184,lime:65280,limegreen:3329330,linen:16445670,magenta:16711935,maroon:8388608,mediumaquamarine:6737322,mediumblue:205,mediumorchid:12211667,mediumpurple:9662683,mediumseagreen:3978097,mediumslateblue:8087790,mediumspringgreen:64154,mediumturquoise:4772300,mediumvioletred:13047173,midnightblue:1644912,mintcream:16121850,mistyrose:16770273,moccasin:16770229,navajowhite:16768685,navy:128,oldlace:16643558,olive:8421376,olivedrab:7048739,orange:16753920,orangered:16729344,orchid:14315734,palegoldenrod:15657130,palegreen:10025880,paleturquoise:11529966,palevioletred:14381203,papayawhip:16773077,peachpuff:16767673,peru:13468991,pink:16761035,plum:14524637,powderblue:11591910,purple:8388736,rebeccapurple:6697881,red:16711680,rosybrown:12357519,royalblue:4286945,saddlebrown:9127187,salmon:16416882,sandybrown:16032864,seagreen:3050327,seashell:16774638,sienna:10506797,silver:12632256,skyblue:8900331,slateblue:6970061,slategray:7372944,slategrey:7372944,snow:16775930,springgreen:65407,steelblue:4620980,tan:13808780,teal:32896,thistle:14204888,tomato:16737095,turquoise:4251856,violet:15631086,wheat:16113331,white:16777215,whitesmoke:16119285,yellow:16776960,yellowgreen:10145074};function m(){return this.rgb().formatHex()}function g(){return this.rgb().formatRgb()}function v(t){var e,r;return t=(t+"").trim().toLowerCase(),(e=s.exec(t))?(r=e[1].length,e=parseInt(e[1],16),6===r?y(e):3===r?new w(e>>8&15|e>>4&240,e>>4&15|240&e,(15&e)<<4|15&e,1):8===r?x(e>>24&255,e>>16&255,e>>8&255,(255&e)/255):4===r?x(e>>12&15|e>>8&240,e>>8&15|e>>4&240,e>>4&15|240&e,((15&e)<<4|15&e)/255):null):(e=l.exec(t))?new w(e[1],e[2],e[3],1):(e=c.exec(t))?new w(255*e[1]/100,255*e[2]/100,255*e[3]/100,1):(e=u.exec(t))?x(e[1],e[2],e[3],e[4]):(e=f.exec(t))?x(255*e[1]/100,255*e[2]/100,255*e[3]/100,e[4]):(e=h.exec(t))?M(e[1],e[2]/100,e[3]/100,1):(e=p.exec(t))?M(e[1],e[2]/100,e[3]/100,e[4]):d.hasOwnProperty(t)?y(d[t]):"transparent"===t?new w(NaN,NaN,NaN,0):null}function y(t){return new w(t>>16&255,t>>8&255,255&t,1)}function x(t,e,r,n){return n<=0&&(t=e=r=NaN),new w(t,e,r,n)}function b(t){return t instanceof n||(t=v(t)),t?new w((t=t.rgb()).r,t.g,t.b,t.opacity):new w}function _(t,e,r,n){return 1===arguments.length?b(t):new w(t,e,r,null==n?1:n)}function w(t,e,r,n){this.r=+t,this.g=+e,this.b=+r,this.opacity=+n}function T(){return"#"+A(this.r)+A(this.g)+A(this.b)}function k(){var t=this.opacity;return(1===(t=isNaN(t)?1:Math.max(0,Math.min(1,t)))?"rgb(":"rgba(")+Math.max(0,Math.min(255,Math.round(this.r)||0))+", "+Math.max(0,Math.min(255,Math.round(this.g)||0))+", "+Math.max(0,Math.min(255,Math.round(this.b)||0))+(1===t?")":", "+t+")")}function A(t){return((t=Math.max(0,Math.min(255,Math.round(t)||0)))<16?"0":"")+t.toString(16)}function M(t,e,r,n){return n<=0?t=e=r=NaN:r<=0||r>=1?t=e=NaN:e<=0&&(t=NaN),new L(t,e,r,n)}function S(t){if(t instanceof L)return new L(t.h,t.s,t.l,t.opacity);if(t instanceof n||(t=v(t)),!t)return new L;if(t instanceof L)return t;var e=(t=t.rgb()).r/255,r=t.g/255,i=t.b/255,a=Math.min(e,r,i),o=Math.max(e,r,i),s=NaN,l=o-a,c=(o+a)/2;return l?(s=e===o?(r-i)/l+6*(r<i):r===o?(i-e)/l+2:(e-r)/l+4,l/=c<.5?o+a:2-o-a,s*=60):l=c>0&&c<1?0:s,new L(s,l,c,t.opacity)}function E(t,e,r,n){return 1===arguments.length?S(t):new L(t,e,r,null==n?1:n)}function L(t,e,r,n){this.h=+t,this.s=+e,this.l=+r,this.opacity=+n}function C(t,e,r){return 255*(t<60?e+(r-e)*t/60:t<180?r:t<240?e+(r-e)*(240-t)/60:e)}e(n,v,{copy:function(t){return Object.assign(new this.constructor,this,t)},displayable:function(){return this.rgb().displayable()},hex:m,formatHex:m,formatHsl:function(){return S(this).formatHsl()},formatRgb:g,toString:g}),e(w,_,r(n,{brighter:function(t){return t=null==t?1/.7:Math.pow(1/.7,t),new w(this.r*t,this.g*t,this.b*t,this.opacity)},darker:function(t){return t=null==t?.7:Math.pow(.7,t),new w(this.r*t,this.g*t,this.b*t,this.opacity)},rgb:function(){return this},displayable:function(){return-.5<=this.r&&this.r<255.5&&-.5<=this.g&&this.g<255.5&&-.5<=this.b&&this.b<255.5&&0<=this.opacity&&this.opacity<=1},hex:T,formatHex:T,formatRgb:k,toString:k})),e(L,E,r(n,{brighter:function(t){return t=null==t?1/.7:Math.pow(1/.7,t),new L(this.h,this.s,this.l*t,this.opacity)},darker:function(t){return t=null==t?.7:Math.pow(.7,t),new L(this.h,this.s,this.l*t,this.opacity)},rgb:function(){var t=this.h%360+360*(this.h<0),e=isNaN(t)||isNaN(this.s)?0:this.s,r=this.l,n=r+(r<.5?r:1-r)*e,i=2*r-n;return new w(C(t>=240?t-240:t+120,i,n),C(t,i,n),C(t<120?t+240:t-120,i,n),this.opacity)},displayable:function(){return(0<=this.s&&this.s<=1||isNaN(this.s))&&0<=this.l&&this.l<=1&&0<=this.opacity&&this.opacity<=1},formatHsl:function(){var t=this.opacity;return(1===(t=isNaN(t)?1:Math.max(0,Math.min(1,t)))?"hsl(":"hsla(")+(this.h||0)+", "+100*(this.s||0)+"%, "+100*(this.l||0)+"%"+(1===t?")":", "+t+")")}}));var P=Math.PI/180,I=180/Math.PI,O=6/29,z=3*O*O;function D(t){if(t instanceof F)return new F(t.l,t.a,t.b,t.opacity);if(t instanceof q)return G(t);t instanceof w||(t=b(t));var e,r,n=U(t.r),i=U(t.g),a=U(t.b),o=B((.2225045*n+.7168786*i+.0606169*a)/1);return n===i&&i===a?e=r=o:(e=B((.4360747*n+.3850649*i+.1430804*a)/.96422),r=B((.0139322*n+.0971045*i+.7141733*a)/.82521)),new F(116*o-16,500*(e-o),200*(o-r),t.opacity)}function R(t,e,r,n){return 1===arguments.length?D(t):new F(t,e,r,null==n?1:n)}function F(t,e,r,n){this.l=+t,this.a=+e,this.b=+r,this.opacity=+n}function B(t){return t>.008856451679035631?Math.pow(t,1/3):t/z+4/29}function N(t){return t>O?t*t*t:z*(t-4/29)}function j(t){return 255*(t<=.0031308?12.92*t:1.055*Math.pow(t,1/2.4)-.055)}function U(t){return(t/=255)<=.04045?t/12.92:Math.pow((t+.055)/1.055,2.4)}function V(t){if(t instanceof q)return new q(t.h,t.c,t.l,t.opacity);if(t instanceof F||(t=D(t)),0===t.a&&0===t.b)return new q(NaN,0<t.l&&t.l<100?0:NaN,t.l,t.opacity);var e=Math.atan2(t.b,t.a)*I;return new q(e<0?e+360:e,Math.sqrt(t.a*t.a+t.b*t.b),t.l,t.opacity)}function H(t,e,r,n){return 1===arguments.length?V(t):new q(t,e,r,null==n?1:n)}function q(t,e,r,n){this.h=+t,this.c=+e,this.l=+r,this.opacity=+n}function G(t){if(isNaN(t.h))return new F(t.l,0,0,t.opacity);var e=t.h*P;return new F(t.l,Math.cos(e)*t.c,Math.sin(e)*t.c,t.opacity)}e(F,R,r(n,{brighter:function(t){return new F(this.l+18*(null==t?1:t),this.a,this.b,this.opacity)},darker:function(t){return new F(this.l-18*(null==t?1:t),this.a,this.b,this.opacity)},rgb:function(){var t=(this.l+16)/116,e=isNaN(this.a)?t:t+this.a/500,r=isNaN(this.b)?t:t-this.b/200;return new w(j(3.1338561*(e=.96422*N(e))-1.6168667*(t=1*N(t))-.4906146*(r=.82521*N(r))),j(-.9787684*e+1.9161415*t+.033454*r),j(.0719453*e-.2289914*t+1.4052427*r),this.opacity)}})),e(q,H,r(n,{brighter:function(t){return new q(this.h,this.c,this.l+18*(null==t?1:t),this.opacity)},darker:function(t){return new q(this.h,this.c,this.l-18*(null==t?1:t),this.opacity)},rgb:function(){return G(this).rgb()}}));var Y=-.14861,W=1.78277,X=-.29227,Z=-.90649,J=1.97294,K=J*Z,Q=J*W,$=W*X-Z*Y;function tt(t){if(t instanceof rt)return new rt(t.h,t.s,t.l,t.opacity);t instanceof w||(t=b(t));var e=t.r/255,r=t.g/255,n=t.b/255,i=($*n+K*e-Q*r)/($+K-Q),a=n-i,o=(J*(r-i)-X*a)/Z,s=Math.sqrt(o*o+a*a)/(J*i*(1-i)),l=s?Math.atan2(o,a)*I-120:NaN;return new rt(l<0?l+360:l,s,i,t.opacity)}function et(t,e,r,n){return 1===arguments.length?tt(t):new rt(t,e,r,null==n?1:n)}function rt(t,e,r,n){this.h=+t,this.s=+e,this.l=+r,this.opacity=+n}e(rt,et,r(n,{brighter:function(t){return t=null==t?1/.7:Math.pow(1/.7,t),new rt(this.h,this.s,this.l*t,this.opacity)},darker:function(t){return t=null==t?.7:Math.pow(.7,t),new rt(this.h,this.s,this.l*t,this.opacity)},rgb:function(){var t=isNaN(this.h)?0:(this.h+120)*P,e=+this.l,r=isNaN(this.s)?0:this.s*e*(1-e),n=Math.cos(t),i=Math.sin(t);return new w(255*(e+r*(Y*n+W*i)),255*(e+r*(X*n+Z*i)),255*(e+r*(J*n)),this.opacity)}})),t.color=v,t.cubehelix=et,t.gray=function(t,e){return new F(t,0,0,null==e?1:e)},t.hcl=H,t.hsl=E,t.lab=R,t.lch=function(t,e,r,n){return 1===arguments.length?V(t):new q(r,e,t,null==n?1:n)},t.rgb=_,Object.defineProperty(t,"__esModule",{value:!0})}))},{}],105:[function(t,e,r){!function(t,n){"object"==typeof r&&void 0!==e?n(r):n((t=t||self).d3=t.d3||{})}(this,(function(t){"use strict";var e={value:function(){}};function r(){for(var t,e=0,r=arguments.length,i={};e<r;++e){if(!(t=arguments[e]+"")||t in i||/[\s.]/.test(t))throw new Error("illegal type: "+t);i[t]=[]}return new n(i)}function n(t){this._=t}function i(t,e){return t.trim().split(/^|\s+/).map((function(t){var r="",n=t.indexOf(".");if(n>=0&&(r=t.slice(n+1),t=t.slice(0,n)),t&&!e.hasOwnProperty(t))throw new Error("unknown type: "+t);return{type:t,name:r}}))}function a(t,e){for(var r,n=0,i=t.length;n<i;++n)if((r=t[n]).name===e)return r.value}function o(t,r,n){for(var i=0,a=t.length;i<a;++i)if(t[i].name===r){t[i]=e,t=t.slice(0,i).concat(t.slice(i+1));break}return null!=n&&t.push({name:r,value:n}),t}n.prototype=r.prototype={constructor:n,on:function(t,e){var r,n=this._,s=i(t+"",n),l=-1,c=s.length;if(!(arguments.length<2)){if(null!=e&&"function"!=typeof e)throw new Error("invalid callback: "+e);for(;++l<c;)if(r=(t=s[l]).type)n[r]=o(n[r],t.name,e);else if(null==e)for(r in n)n[r]=o(n[r],t.name,null);return this}for(;++l<c;)if((r=(t=s[l]).type)&&(r=a(n[r],t.name)))return r},copy:function(){var t={},e=this._;for(var r in e)t[r]=e[r].slice();return new n(t)},call:function(t,e){if((r=arguments.length-2)>0)for(var r,n,i=new Array(r),a=0;a<r;++a)i[a]=arguments[a+2];if(!this._.hasOwnProperty(t))throw new Error("unknown type: "+t);for(a=0,r=(n=this._[t]).length;a<r;++a)n[a].value.apply(e,i)},apply:function(t,e,r){if(!this._.hasOwnProperty(t))throw new Error("unknown type: "+t);for(var n=this._[t],i=0,a=n.length;i<a;++i)n[i].value.apply(e,r)}},t.dispatch=r,Object.defineProperty(t,"__esModule",{value:!0})}))},{}],106:[function(t,e,r){!function(n,i){"object"==typeof r&&void 0!==e?i(r,t("d3-quadtree"),t("d3-collection"),t("d3-dispatch"),t("d3-timer")):i(n.d3=n.d3||{},n.d3,n.d3,n.d3,n.d3)}(this,(function(t,e,r,n,i){"use strict";function a(t){return function(){return t}}function o(){return 1e-6*(Math.random()-.5)}function s(t){return t.x+t.vx}function l(t){return t.y+t.vy}function c(t){return t.index}function u(t,e){var r=t.get(e);if(!r)throw new Error("missing: "+e);return r}function f(t){return t.x}function h(t){return t.y}var p=Math.PI*(3-Math.sqrt(5));t.forceCenter=function(t,e){var r;function n(){var n,i,a=r.length,o=0,s=0;for(n=0;n<a;++n)o+=(i=r[n]).x,s+=i.y;for(o=o/a-t,s=s/a-e,n=0;n<a;++n)(i=r[n]).x-=o,i.y-=s}return null==t&&(t=0),null==e&&(e=0),n.initialize=function(t){r=t},n.x=function(e){return arguments.length?(t=+e,n):t},n.y=function(t){return arguments.length?(e=+t,n):e},n},t.forceCollide=function(t){var r,n,i=1,c=1;function u(){for(var t,a,u,h,p,d,m,g=r.length,v=0;v<c;++v)for(a=e.quadtree(r,s,l).visitAfter(f),t=0;t<g;++t)u=r[t],d=n[u.index],m=d*d,h=u.x+u.vx,p=u.y+u.vy,a.visit(y);function y(t,e,r,n,a){var s=t.data,l=t.r,c=d+l;if(!s)return e>h+c||n<h-c||r>p+c||a<p-c;if(s.index>u.index){var f=h-s.x-s.vx,g=p-s.y-s.vy,v=f*f+g*g;v<c*c&&(0===f&&(v+=(f=o())*f),0===g&&(v+=(g=o())*g),v=(c-(v=Math.sqrt(v)))/v*i,u.vx+=(f*=v)*(c=(l*=l)/(m+l)),u.vy+=(g*=v)*c,s.vx-=f*(c=1-c),s.vy-=g*c)}}}function f(t){if(t.data)return t.r=n[t.data.index];for(var e=t.r=0;e<4;++e)t[e]&&t[e].r>t.r&&(t.r=t[e].r)}function h(){if(r){var e,i,a=r.length;for(n=new Array(a),e=0;e<a;++e)i=r[e],n[i.index]=+t(i,e,r)}}return"function"!=typeof t&&(t=a(null==t?1:+t)),u.initialize=function(t){r=t,h()},u.iterations=function(t){return arguments.length?(c=+t,u):c},u.strength=function(t){return arguments.length?(i=+t,u):i},u.radius=function(e){return arguments.length?(t="function"==typeof e?e:a(+e),h(),u):t},u},t.forceLink=function(t){var e,n,i,s,l,f=c,h=function(t){return 1/Math.min(s[t.source.index],s[t.target.index])},p=a(30),d=1;function m(r){for(var i=0,a=t.length;i<d;++i)for(var s,c,u,f,h,p,m,g=0;g<a;++g)c=(s=t[g]).source,f=(u=s.target).x+u.vx-c.x-c.vx||o(),h=u.y+u.vy-c.y-c.vy||o(),f*=p=((p=Math.sqrt(f*f+h*h))-n[g])/p*r*e[g],h*=p,u.vx-=f*(m=l[g]),u.vy-=h*m,c.vx+=f*(m=1-m),c.vy+=h*m}function g(){if(i){var a,o,c=i.length,h=t.length,p=r.map(i,f);for(a=0,s=new Array(c);a<h;++a)(o=t[a]).index=a,"object"!=typeof o.source&&(o.source=u(p,o.source)),"object"!=typeof o.target&&(o.target=u(p,o.target)),s[o.source.index]=(s[o.source.index]||0)+1,s[o.target.index]=(s[o.target.index]||0)+1;for(a=0,l=new Array(h);a<h;++a)o=t[a],l[a]=s[o.source.index]/(s[o.source.index]+s[o.target.index]);e=new Array(h),v(),n=new Array(h),y()}}function v(){if(i)for(var r=0,n=t.length;r<n;++r)e[r]=+h(t[r],r,t)}function y(){if(i)for(var e=0,r=t.length;e<r;++e)n[e]=+p(t[e],e,t)}return null==t&&(t=[]),m.initialize=function(t){i=t,g()},m.links=function(e){return arguments.length?(t=e,g(),m):t},m.id=function(t){return arguments.length?(f=t,m):f},m.iterations=function(t){return arguments.length?(d=+t,m):d},m.strength=function(t){return arguments.length?(h="function"==typeof t?t:a(+t),v(),m):h},m.distance=function(t){return arguments.length?(p="function"==typeof t?t:a(+t),y(),m):p},m},t.forceManyBody=function(){var t,r,n,i,s=a(-30),l=1,c=1/0,u=.81;function p(i){var a,o=t.length,s=e.quadtree(t,f,h).visitAfter(m);for(n=i,a=0;a<o;++a)r=t[a],s.visit(g)}function d(){if(t){var e,r,n=t.length;for(i=new Array(n),e=0;e<n;++e)r=t[e],i[r.index]=+s(r,e,t)}}function m(t){var e,r,n,a,o,s=0,l=0;if(t.length){for(n=a=o=0;o<4;++o)(e=t[o])&&(r=Math.abs(e.value))&&(s+=e.value,l+=r,n+=r*e.x,a+=r*e.y);t.x=n/l,t.y=a/l}else{(e=t).x=e.data.x,e.y=e.data.y;do{s+=i[e.data.index]}while(e=e.next)}t.value=s}function g(t,e,a,s){if(!t.value)return!0;var f=t.x-r.x,h=t.y-r.y,p=s-e,d=f*f+h*h;if(p*p/u<d)return d<c&&(0===f&&(d+=(f=o())*f),0===h&&(d+=(h=o())*h),d<l&&(d=Math.sqrt(l*d)),r.vx+=f*t.value*n/d,r.vy+=h*t.value*n/d),!0;if(!(t.length||d>=c)){(t.data!==r||t.next)&&(0===f&&(d+=(f=o())*f),0===h&&(d+=(h=o())*h),d<l&&(d=Math.sqrt(l*d)));do{t.data!==r&&(p=i[t.data.index]*n/d,r.vx+=f*p,r.vy+=h*p)}while(t=t.next)}}return p.initialize=function(e){t=e,d()},p.strength=function(t){return arguments.length?(s="function"==typeof t?t:a(+t),d(),p):s},p.distanceMin=function(t){return arguments.length?(l=t*t,p):Math.sqrt(l)},p.distanceMax=function(t){return arguments.length?(c=t*t,p):Math.sqrt(c)},p.theta=function(t){return arguments.length?(u=t*t,p):Math.sqrt(u)},p},t.forceRadial=function(t,e,r){var n,i,o,s=a(.1);function l(t){for(var a=0,s=n.length;a<s;++a){var l=n[a],c=l.x-e||1e-6,u=l.y-r||1e-6,f=Math.sqrt(c*c+u*u),h=(o[a]-f)*i[a]*t/f;l.vx+=c*h,l.vy+=u*h}}function c(){if(n){var e,r=n.length;for(i=new Array(r),o=new Array(r),e=0;e<r;++e)o[e]=+t(n[e],e,n),i[e]=isNaN(o[e])?0:+s(n[e],e,n)}}return"function"!=typeof t&&(t=a(+t)),null==e&&(e=0),null==r&&(r=0),l.initialize=function(t){n=t,c()},l.strength=function(t){return arguments.length?(s="function"==typeof t?t:a(+t),c(),l):s},l.radius=function(e){return arguments.length?(t="function"==typeof e?e:a(+e),c(),l):t},l.x=function(t){return arguments.length?(e=+t,l):e},l.y=function(t){return arguments.length?(r=+t,l):r},l},t.forceSimulation=function(t){var e,a=1,o=.001,s=1-Math.pow(o,1/300),l=0,c=.6,u=r.map(),f=i.timer(d),h=n.dispatch("tick","end");function d(){m(),h.call("tick",e),a<o&&(f.stop(),h.call("end",e))}function m(r){var n,i,o=t.length;void 0===r&&(r=1);for(var f=0;f<r;++f)for(a+=(l-a)*s,u.each((function(t){t(a)})),n=0;n<o;++n)null==(i=t[n]).fx?i.x+=i.vx*=c:(i.x=i.fx,i.vx=0),null==i.fy?i.y+=i.vy*=c:(i.y=i.fy,i.vy=0);return e}function g(){for(var e,r=0,n=t.length;r<n;++r){if((e=t[r]).index=r,null!=e.fx&&(e.x=e.fx),null!=e.fy&&(e.y=e.fy),isNaN(e.x)||isNaN(e.y)){var i=10*Math.sqrt(r),a=r*p;e.x=i*Math.cos(a),e.y=i*Math.sin(a)}(isNaN(e.vx)||isNaN(e.vy))&&(e.vx=e.vy=0)}}function v(e){return e.initialize&&e.initialize(t),e}return null==t&&(t=[]),g(),e={tick:m,restart:function(){return f.restart(d),e},stop:function(){return f.stop(),e},nodes:function(r){return arguments.length?(t=r,g(),u.each(v),e):t},alpha:function(t){return arguments.length?(a=+t,e):a},alphaMin:function(t){return arguments.length?(o=+t,e):o},alphaDecay:function(t){return arguments.length?(s=+t,e):+s},alphaTarget:function(t){return arguments.length?(l=+t,e):l},velocityDecay:function(t){return arguments.length?(c=1-t,e):1-c},force:function(t,r){return arguments.length>1?(null==r?u.remove(t):u.set(t,v(r)),e):u.get(t)},find:function(e,r,n){var i,a,o,s,l,c=0,u=t.length;for(null==n?n=1/0:n*=n,c=0;c<u;++c)(o=(i=e-(s=t[c]).x)*i+(a=r-s.y)*a)<n&&(l=s,n=o);return l},on:function(t,r){return arguments.length>1?(h.on(t,r),e):h.on(t)}}},t.forceX=function(t){var e,r,n,i=a(.1);function o(t){for(var i,a=0,o=e.length;a<o;++a)(i=e[a]).vx+=(n[a]-i.x)*r[a]*t}function s(){if(e){var a,o=e.length;for(r=new Array(o),n=new Array(o),a=0;a<o;++a)r[a]=isNaN(n[a]=+t(e[a],a,e))?0:+i(e[a],a,e)}}return"function"!=typeof t&&(t=a(null==t?0:+t)),o.initialize=function(t){e=t,s()},o.strength=function(t){return arguments.length?(i="function"==typeof t?t:a(+t),s(),o):i},o.x=function(e){return arguments.length?(t="function"==typeof e?e:a(+e),s(),o):t},o},t.forceY=function(t){var e,r,n,i=a(.1);function o(t){for(var i,a=0,o=e.length;a<o;++a)(i=e[a]).vy+=(n[a]-i.y)*r[a]*t}function s(){if(e){var a,o=e.length;for(r=new Array(o),n=new Array(o),a=0;a<o;++a)r[a]=isNaN(n[a]=+t(e[a],a,e))?0:+i(e[a],a,e)}}return"function"!=typeof t&&(t=a(null==t?0:+t)),o.initialize=function(t){e=t,s()},o.strength=function(t){return arguments.length?(i="function"==typeof t?t:a(+t),s(),o):i},o.y=function(e){return arguments.length?(t="function"==typeof e?e:a(+e),s(),o):t},o},Object.defineProperty(t,"__esModule",{value:!0})}))},{"d3-collection":103,"d3-dispatch":105,"d3-quadtree":113,"d3-timer":117}],107:[function(t,e,r){!function(t,n){"object"==typeof r&&void 0!==e?n(r):n((t="undefined"!=typeof globalThis?globalThis:t||self).d3=t.d3||{})}(this,(function(t){"use strict";function e(t,e){if((r=(t=e?t.toExponential(e-1):t.toExponential()).indexOf("e"))<0)return null;var r,n=t.slice(0,r);return[n.length>1?n[0]+n.slice(2):n,+t.slice(r+1)]}function r(t){return(t=e(Math.abs(t)))?t[1]:NaN}var n,i=/^(?:(.)?([<>=^]))?([+\-( ])?([$#])?(0)?(\d+)?(,)?(\.\d+)?(~)?([a-z%])?$/i;function a(t){if(!(e=i.exec(t)))throw new Error("invalid format: "+t);var e;return new o({fill:e[1],align:e[2],sign:e[3],symbol:e[4],zero:e[5],width:e[6],comma:e[7],precision:e[8]&&e[8].slice(1),trim:e[9],type:e[10]})}function o(t){this.fill=void 0===t.fill?" ":t.fill+"",this.align=void 0===t.align?">":t.align+"",this.sign=void 0===t.sign?"-":t.sign+"",this.symbol=void 0===t.symbol?"":t.symbol+"",this.zero=!!t.zero,this.width=void 0===t.width?void 0:+t.width,this.comma=!!t.comma,this.precision=void 0===t.precision?void 0:+t.precision,this.trim=!!t.trim,this.type=void 0===t.type?"":t.type+""}function s(t,r){var n=e(t,r);if(!n)return t+"";var i=n[0],a=n[1];return a<0?"0."+new Array(-a).join("0")+i:i.length>a+1?i.slice(0,a+1)+"."+i.slice(a+1):i+new Array(a-i.length+2).join("0")}a.prototype=o.prototype,o.prototype.toString=function(){return this.fill+this.align+this.sign+this.symbol+(this.zero?"0":"")+(void 0===this.width?"":Math.max(1,0|this.width))+(this.comma?",":"")+(void 0===this.precision?"":"."+Math.max(0,0|this.precision))+(this.trim?"~":"")+this.type};var l={"%":function(t,e){return(100*t).toFixed(e)},b:function(t){return Math.round(t).toString(2)},c:function(t){return t+""},d:function(t){return Math.abs(t=Math.round(t))>=1e21?t.toLocaleString("en").replace(/,/g,""):t.toString(10)},e:function(t,e){return t.toExponential(e)},f:function(t,e){return t.toFixed(e)},g:function(t,e){return t.toPrecision(e)},o:function(t){return Math.round(t).toString(8)},p:function(t,e){return s(100*t,e)},r:s,s:function(t,r){var i=e(t,r);if(!i)return t+"";var a=i[0],o=i[1],s=o-(n=3*Math.max(-8,Math.min(8,Math.floor(o/3))))+1,l=a.length;return s===l?a:s>l?a+new Array(s-l+1).join("0"):s>0?a.slice(0,s)+"."+a.slice(s):"0."+new Array(1-s).join("0")+e(t,Math.max(0,r+s-1))[0]},X:function(t){return Math.round(t).toString(16).toUpperCase()},x:function(t){return Math.round(t).toString(16)}};function c(t){return t}var u,f=Array.prototype.map,h=["y","z","a","f","p","n","\xb5","m","","k","M","G","T","P","E","Z","Y"];function p(t){var e,i,o=void 0===t.grouping||void 0===t.thousands?c:(e=f.call(t.grouping,Number),i=t.thousands+"",function(t,r){for(var n=t.length,a=[],o=0,s=e[0],l=0;n>0&&s>0&&(l+s+1>r&&(s=Math.max(1,r-l)),a.push(t.substring(n-=s,n+s)),!((l+=s+1)>r));)s=e[o=(o+1)%e.length];return a.reverse().join(i)}),s=void 0===t.currency?"":t.currency[0]+"",u=void 0===t.currency?"":t.currency[1]+"",p=void 0===t.decimal?".":t.decimal+"",d=void 0===t.numerals?c:function(t){return function(e){return e.replace(/[0-9]/g,(function(e){return t[+e]}))}}(f.call(t.numerals,String)),m=void 0===t.percent?"%":t.percent+"",g=void 0===t.minus?"-":t.minus+"",v=void 0===t.nan?"NaN":t.nan+"";function y(t){var e=(t=a(t)).fill,r=t.align,i=t.sign,c=t.symbol,f=t.zero,y=t.width,x=t.comma,b=t.precision,_=t.trim,w=t.type;"n"===w?(x=!0,w="g"):l[w]||(void 0===b&&(b=12),_=!0,w="g"),(f||"0"===e&&"="===r)&&(f=!0,e="0",r="=");var T="$"===c?s:"#"===c&&/[boxX]/.test(w)?"0"+w.toLowerCase():"",k="$"===c?u:/[%p]/.test(w)?m:"",A=l[w],M=/[defgprs%]/.test(w);function S(t){var a,s,l,c=T,u=k;if("c"===w)u=A(t)+u,t="";else{var m=(t=+t)<0||1/t<0;if(t=isNaN(t)?v:A(Math.abs(t),b),_&&(t=function(t){t:for(var e,r=t.length,n=1,i=-1;n<r;++n)switch(t[n]){case".":i=e=n;break;case"0":0===i&&(i=n),e=n;break;default:if(!+t[n])break t;i>0&&(i=0)}return i>0?t.slice(0,i)+t.slice(e+1):t}(t)),m&&0==+t&&"+"!==i&&(m=!1),c=(m?"("===i?i:g:"-"===i||"("===i?"":i)+c,u=("s"===w?h[8+n/3]:"")+u+(m&&"("===i?")":""),M)for(a=-1,s=t.length;++a<s;)if(48>(l=t.charCodeAt(a))||l>57){u=(46===l?p+t.slice(a+1):t.slice(a))+u,t=t.slice(0,a);break}}x&&!f&&(t=o(t,1/0));var S=c.length+t.length+u.length,E=S<y?new Array(y-S+1).join(e):"";switch(x&&f&&(t=o(E+t,E.length?y-u.length:1/0),E=""),r){case"<":t=c+t+u+E;break;case"=":t=c+E+t+u;break;case"^":t=E.slice(0,S=E.length>>1)+c+t+u+E.slice(S);break;default:t=E+c+t+u}return d(t)}return b=void 0===b?6:/[gprs]/.test(w)?Math.max(1,Math.min(21,b)):Math.max(0,Math.min(20,b)),S.toString=function(){return t+""},S}return{format:y,formatPrefix:function(t,e){var n=y(((t=a(t)).type="f",t)),i=3*Math.max(-8,Math.min(8,Math.floor(r(e)/3))),o=Math.pow(10,-i),s=h[8+i/3];return function(t){return n(o*t)+s}}}}function d(e){return u=p(e),t.format=u.format,t.formatPrefix=u.formatPrefix,u}d({decimal:".",thousands:",",grouping:[3],currency:["$",""],minus:"-"}),t.FormatSpecifier=o,t.formatDefaultLocale=d,t.formatLocale=p,t.formatSpecifier=a,t.precisionFixed=function(t){return Math.max(0,-r(Math.abs(t)))},t.precisionPrefix=function(t,e){return Math.max(0,3*Math.max(-8,Math.min(8,Math.floor(r(e)/3)))-r(Math.abs(t)))},t.precisionRound=function(t,e){return t=Math.abs(t),e=Math.abs(e)-t,Math.max(0,r(e)-r(t))+1},Object.defineProperty(t,"__esModule",{value:!0})}))},{}],108:[function(t,e,r){!function(n,i){"object"==typeof r&&void 0!==e?i(r,t("d3-geo"),t("d3-array")):i(n.d3=n.d3||{},n.d3,n.d3)}(this,(function(t,e,r){"use strict";var n=Math.abs,i=Math.atan,a=Math.atan2,o=Math.cos,s=Math.exp,l=Math.floor,c=Math.log,u=Math.max,f=Math.min,h=Math.pow,p=Math.round,d=Math.sign||function(t){return t>0?1:t<0?-1:0},m=Math.sin,g=Math.tan,v=1e-6,y=Math.PI,x=y/2,b=y/4,_=Math.SQRT1_2,w=L(2),T=L(y),k=2*y,A=180/y,M=y/180;function S(t){return t>1?x:t<-1?-x:Math.asin(t)}function E(t){return t>1?0:t<-1?y:Math.acos(t)}function L(t){return t>0?Math.sqrt(t):0}function C(t){return(s(t)-s(-t))/2}function P(t){return(s(t)+s(-t))/2}function I(t){var e=g(t/2),r=2*c(o(t/2))/(e*e);function i(t,e){var n=o(t),i=o(e),a=m(e),s=i*n,l=-((1-s?c((1+s)/2)/(1-s):-.5)+r/(1+s));return[l*i*m(t),l*a]}return i.invert=function(e,i){var s,l=L(e*e+i*i),u=-t/2,f=50;if(!l)return[0,0];do{var h=u/2,p=o(h),d=m(h),g=d/p,y=-c(n(p));u-=s=(2/g*y-r*g-l)/(-y/(d*d)+1-r/(2*p*p))*(p<0?.7:1)}while(n(s)>v&&--f>0);var x=m(u);return[a(e*x,l*o(u)),S(i*x/l)]},i}function O(t,e){var r=o(e),n=function(t){return t?t/Math.sin(t):1}(E(r*o(t/=2)));return[2*r*m(t)*n,m(e)*n]}function z(t){var e=m(t),r=o(t),i=t>=0?1:-1,s=g(i*t),l=(1+e-r)/2;function c(t,n){var c=o(n),u=o(t/=2);return[(1+c)*m(t),(i*n>-a(u,s)-.001?0:10*-i)+l+m(n)*r-(1+c)*e*u]}return c.invert=function(t,c){var u=0,f=0,h=50;do{var p=o(u),d=m(u),g=o(f),y=m(f),x=1+g,b=x*d-t,_=l+y*r-x*e*p-c,w=x*p/2,T=-d*y,k=e*x*d/2,A=r*g+e*p*y,M=T*k-A*w,S=(_*T-b*A)/M/2,E=(b*k-_*w)/M;n(E)>2&&(E/=2),u-=S,f-=E}while((n(S)>v||n(E)>v)&&--h>0);return i*f>-a(o(u),s)-.001?[2*u,f]:null},c}function D(t,e){var r=g(e/2),n=L(1-r*r),i=1+n*o(t/=2),a=m(t)*n/i,s=r/i,l=a*a,c=s*s;return[4/3*a*(3+l-3*c),4/3*s*(3+3*l-c)]}O.invert=function(t,e){if(!(t*t+4*e*e>y*y+v)){var r=t,i=e,a=25;do{var s,l=m(r),c=m(r/2),u=o(r/2),f=m(i),h=o(i),p=m(2*i),d=f*f,g=h*h,x=c*c,b=1-g*u*u,_=b?E(h*u)*L(s=1/b):s=0,w=2*_*h*c-t,T=_*f-e,k=s*(g*x+_*h*u*d),A=s*(.5*l*p-2*_*f*c),M=.25*s*(p*c-_*f*g*l),S=s*(d*u+_*x*h),C=A*M-S*k;if(!C)break;var P=(T*A-w*S)/C,I=(w*M-T*k)/C;r-=P,i-=I}while((n(P)>v||n(I)>v)&&--a>0);return[r,i]}},D.invert=function(t,e){if(e*=3/8,!(t*=3/8)&&n(e)>1)return null;var r=1+t*t+e*e,i=L((r-L(r*r-4*e*e))/2),s=S(i)/3,l=i?function(t){return c(t+L(t*t-1))}(n(e/i))/3:function(t){return c(t+L(t*t+1))}(n(t))/3,u=o(s),f=P(l),h=f*f-u*u;return[2*d(t)*a(C(l)*u,.25-h),2*d(e)*a(f*m(s),.25+h)]};var R=L(8),F=c(1+w);function B(t,e){var r=n(e);return r<b?[t,c(g(b+e/2))]:[t*o(r)*(2*w-1/m(r)),d(e)*(2*w*(r-b)-c(g(r/2)))]}function N(t){var r=2*y/t;function s(t,i){var s=e.geoAzimuthalEquidistantRaw(t,i);if(n(t)>x){var l=a(s[1],s[0]),c=L(s[0]*s[0]+s[1]*s[1]),u=r*p((l-x)/r)+x,f=a(m(l-=u),2-o(l));l=u+S(y/c*m(f))-f,s[0]=c*o(l),s[1]=c*m(l)}return s}return s.invert=function(t,n){var s=L(t*t+n*n);if(s>x){var l=a(n,t),c=r*p((l-x)/r)+x,u=l>c?-1:1,f=s*o(c-l),h=1/g(u*E((f-y)/L(y*(y-2*f)+s*s)));l=c+2*i((h+u*L(h*h-3))/3),t=s*o(l),n=s*m(l)}return e.geoAzimuthalEquidistantRaw.invert(t,n)},s}function j(t,r){if(arguments.length<2&&(r=t),1===r)return e.geoAzimuthalEqualAreaRaw;if(r===1/0)return U;function n(n,i){var a=e.geoAzimuthalEqualAreaRaw(n/r,i);return a[0]*=t,a}return n.invert=function(n,i){var a=e.geoAzimuthalEqualAreaRaw.invert(n/t,i);return a[0]*=r,a},n}function U(t,e){return[t*o(e)/o(e/=2),2*m(e)]}function V(t,e,r){var i,a,o,s=100;r=void 0===r?0:+r,e=+e;do{(a=t(r))===(o=t(r+v))&&(o=a+v),r-=i=-1*v*(a-e)/(a-o)}while(s-- >0&&n(i)>v);return s<0?NaN:r}function H(t,e,r){return void 0===e&&(e=40),void 0===r&&(r=1e-12),function(i,a,o,s){var l,c,u;o=void 0===o?0:+o,s=void 0===s?0:+s;for(var f=0;f<e;f++){var h=t(o,s),p=h[0]-i,d=h[1]-a;if(n(p)<r&&n(d)<r)break;var m=p*p+d*d;if(m>l)o-=c/=2,s-=u/=2;else{l=m;var g=(o>0?-1:1)*r,v=(s>0?-1:1)*r,y=t(o+g,s),x=t(o,s+v),b=(y[0]-h[0])/g,_=(y[1]-h[1])/g,w=(x[0]-h[0])/v,T=(x[1]-h[1])/v,k=T*b-_*w,A=(n(k)<.5?.5:1)/k;if(o+=c=(d*w-p*T)*A,s+=u=(p*_-d*b)*A,n(c)<r&&n(u)<r)break}}return[o,s]}}function q(){var t=j(1.68,2);function e(e,r){if(e+r<-1.4){var n=(e-r+1.6)*(e+r+1.4)/8;e+=n,r-=.8*n*m(r+y/2)}var i=t(e,r),a=(1-o(e*r))/12;return i[1]<0&&(i[0]*=1+a),i[1]>0&&(i[1]*=1+a/1.5*i[0]*i[0]),i}return e.invert=H(e),e}function G(t,e){var r,i=t*m(e),a=30;do{e-=r=(e+m(e)-i)/(1+o(e))}while(n(r)>v&&--a>0);return e/2}function Y(t,e,r){function n(n,i){return[t*n*o(i=G(r,i)),e*m(i)]}return n.invert=function(n,i){return i=S(i/e),[n/(t*o(i)),S((2*i+m(2*i))/r)]},n}B.invert=function(t,e){if((a=n(e))<F)return[t,2*i(s(e))-x];var r,a,l=b,u=25;do{var f=o(l/2),h=g(l/2);l-=r=(R*(l-b)-c(h)-a)/(R-f*f/(2*h))}while(n(r)>1e-12&&--u>0);return[t/(o(l)*(R-1/m(l))),d(e)*l]},U.invert=function(t,e){var r=2*S(e/2);return[t*o(r/2)/o(r),r]};var W=Y(w/x,w,y);var X=2.00276,Z=1.11072;function J(t,e){var r=G(y,e);return[X*t/(1/o(e)+Z/o(r)),(e+w*m(r))/X]}function K(t){var r=0,n=e.geoProjectionMutator(t),i=n(r);return i.parallel=function(t){return arguments.length?n(r=t*M):r*A},i}function Q(t,e){return[t*o(e),e]}function $(t){if(!t)return Q;var e=1/g(t);function r(r,n){var i=e+t-n,a=i?r*o(n)/i:i;return[i*m(a),e-i*o(a)]}return r.invert=function(r,n){var i=L(r*r+(n=e-n)*n),s=e+t-i;return[i/o(s)*a(r,n),s]},r}function tt(t){function e(e,r){var n=x-r,i=n?e*t*m(n)/n:n;return[n*m(i)/t,x-n*o(i)]}return e.invert=function(e,r){var n=e*t,i=x-r,o=L(n*n+i*i),s=a(n,i);return[(o?o/m(o):1)*s/t,x-o]},e}J.invert=function(t,e){var r,i,a=X*e,s=e<0?-b:b,l=25;do{i=a-w*m(s),s-=r=(m(2*s)+2*s-y*m(i))/(2*o(2*s)+2+y*o(i)*w*o(s))}while(n(r)>v&&--l>0);return i=a-w*m(s),[t*(1/o(i)+Z/o(s))/X,i]},Q.invert=function(t,e){return[t/o(e),e]};var et=Y(1,4/y,y);function rt(t,e,r,i,s,l){var c,u=o(l);if(n(t)>1||n(l)>1)c=E(r*s+e*i*u);else{var f=m(t/2),h=m(l/2);c=2*S(L(f*f+e*i*h*h))}return n(c)>v?[c,a(i*m(l),e*s-r*i*u)]:[0,0]}function nt(t,e,r){return E((t*t+e*e-r*r)/(2*t*e))}function it(t){return t-2*y*l((t+y)/(2*y))}function at(t,e,r){for(var n,i=[[t[0],t[1],m(t[1]),o(t[1])],[e[0],e[1],m(e[1]),o(e[1])],[r[0],r[1],m(r[1]),o(r[1])]],a=i[2],s=0;s<3;++s,a=n)n=i[s],a.v=rt(n[1]-a[1],a[3],a[2],n[3],n[2],n[0]-a[0]),a.point=[0,0];var l=nt(i[0].v[0],i[2].v[0],i[1].v[0]),c=nt(i[0].v[0],i[1].v[0],i[2].v[0]),u=y-l;i[2].point[1]=0,i[0].point[0]=-(i[1].point[0]=i[0].v[0]/2);var f=[i[2].point[0]=i[0].point[0]+i[2].v[0]*o(l),2*(i[0].point[1]=i[1].point[1]=i[2].v[0]*m(l))];return function(t,e){var r,n=m(e),a=o(e),s=new Array(3);for(r=0;r<3;++r){var l=i[r];if(s[r]=rt(e-l[1],l[3],l[2],a,n,t-l[0]),!s[r][0])return l.point;s[r][1]=it(s[r][1]-l.v[1])}var h=f.slice();for(r=0;r<3;++r){var p=2==r?0:r+1,d=nt(i[r].v[0],s[r][0],s[p][0]);s[r][1]<0&&(d=-d),r?1==r?(d=c-d,h[0]-=s[r][0]*o(d),h[1]-=s[r][0]*m(d)):(d=u-d,h[0]+=s[r][0]*o(d),h[1]+=s[r][0]*m(d)):(h[0]+=s[r][0]*o(d),h[1]-=s[r][0]*m(d))}return h[0]/=3,h[1]/=3,h}}function ot(t){return t[0]*=M,t[1]*=M,t}function st(t,r,n){var i=e.geoCentroid({type:"MultiPoint",coordinates:[t,r,n]}),a=[-i[0],-i[1]],o=e.geoRotation(a),s=at(ot(o(t)),ot(o(r)),ot(o(n)));s.invert=H(s);var l=e.geoProjection(s).rotate(a),c=l.center;return delete l.rotate,l.center=function(t){return arguments.length?c(o(t)):o.invert(c())},l.clipAngle(90)}function lt(t,e){var r=L(1-m(e));return[2/T*t*r,T*(1-r)]}function ct(t){var e=g(t);function r(t,r){return[t,(t?t/m(t):1)*(m(r)*o(t)-e*o(r))]}return r.invert=e?function(t,r){t&&(r*=m(t)/t);var n=o(t);return[t,2*a(L(n*n+e*e-r*r)-n,e-r)]}:function(t,e){return[t,S(t?e*g(t)/t:e)]},r}lt.invert=function(t,e){var r=(r=e/T-1)*r;return[r>0?t*L(y/r)/2:0,S(1-r)]};var ut=L(3);function ft(t,e){return[ut*t*(2*o(2*e/3)-1)/T,ut*T*m(e/3)]}function ht(t){var e=o(t);function r(t,r){return[t*e,m(r)/e]}return r.invert=function(t,r){return[t/e,S(r*e)]},r}function pt(t){var e=o(t);function r(t,r){return[t*e,(1+e)*g(r/2)]}return r.invert=function(t,r){return[t/e,2*i(r/(1+e))]},r}function dt(t,e){var r=L(8/(3*y));return[r*t*(1-n(e)/y),r*e]}function mt(t,e){var r=L(4-3*m(n(e)));return[2/L(6*y)*t*r,d(e)*L(2*y/3)*(2-r)]}function gt(t,e){var r=L(y*(4+y));return[2/r*t*(1+L(1-4*e*e/(y*y))),4/r*e]}function vt(t,e){var r=(2+x)*m(e);e/=2;for(var i=0,a=1/0;i<10&&n(a)>v;i++){var s=o(e);e-=a=(e+m(e)*(s+2)-r)/(2*s*(1+s))}return[2/L(y*(4+y))*t*(1+o(e)),2*L(y/(4+y))*m(e)]}function yt(t,e){return[t*(1+o(e))/L(2+y),2*e/L(2+y)]}function xt(t,e){for(var r=(1+x)*m(e),i=0,a=1/0;i<10&&n(a)>v;i++)e-=a=(e+m(e)-r)/(1+o(e));return r=L(2+y),[t*(1+o(e))/r,2*e/r]}ft.invert=function(t,e){var r=3*S(e/(ut*T));return[T*t/(ut*(2*o(2*r/3)-1)),r]},dt.invert=function(t,e){var r=L(8/(3*y)),i=e/r;return[t/(r*(1-n(i)/y)),i]},mt.invert=function(t,e){var r=2-n(e)/L(2*y/3);return[t*L(6*y)/(2*r),d(e)*S((4-r*r)/3)]},gt.invert=function(t,e){var r=L(y*(4+y))/2;return[t*r/(1+L(1-e*e*(4+y)/(4*y))),e*r/2]},vt.invert=function(t,e){var r=e*L((4+y)/y)/2,n=S(r),i=o(n);return[t/(2/L(y*(4+y))*(1+i)),S((n+r*(i+2))/(2+x))]},yt.invert=function(t,e){var r=L(2+y),n=e*r/2;return[r*t/(1+o(n)),n]},xt.invert=function(t,e){var r=1+x,n=L(r/2);return[2*t*n/(1+o(e*=n)),S((e+m(e))/r)]};var bt=3+2*w;function _t(t,e){var r=m(t/=2),n=o(t),a=L(o(e)),s=o(e/=2),l=m(e)/(s+w*n*a),u=L(2/(1+l*l)),f=L((w*s+(n+r)*a)/(w*s+(n-r)*a));return[bt*(u*(f-1/f)-2*c(f)),bt*(u*l*(f+1/f)-2*i(l))]}_t.invert=function(t,e){if(!(r=D.invert(t/1.2,1.065*e)))return null;var r,a=r[0],s=r[1],l=20;t/=bt,e/=bt;do{var h=a/2,p=s/2,d=m(h),g=o(h),y=m(p),b=o(p),T=o(s),k=L(T),A=y/(b+w*g*k),M=A*A,S=L(2/(1+M)),E=(w*b+(g+d)*k)/(w*b+(g-d)*k),C=L(E),P=C-1/C,I=C+1/C,O=S*P-2*c(C)-t,z=S*A*I-2*i(A)-e,R=y&&_*k*d*M/y,F=(w*g*b+k)/(2*(b+w*g*k)*(b+w*g*k)*k),B=-.5*A*S*S*S,N=B*R,j=B*F,U=(U=2*b+w*k*(g-d))*U*C,V=(w*g*b*k+T)/U,H=-w*d*y/(k*U),q=P*N-2*V/C+S*(V+V/E),G=P*j-2*H/C+S*(H+H/E),Y=A*I*N-2*R/(1+M)+S*I*R+S*A*(V-V/E),W=A*I*j-2*F/(1+M)+S*I*F+S*A*(H-H/E),X=G*Y-W*q;if(!X)break;var Z=(z*G-O*W)/X,J=(O*Y-z*q)/X;a-=Z,s=u(-x,f(x,s-J))}while((n(Z)>v||n(J)>v)&&--l>0);return n(n(s)-x)<v?[0,s]:l&&[a,s]};var wt=o(35*M);function Tt(t,e){var r=g(e/2);return[t*wt*L(1-r*r),(1+wt)*r]}function kt(t,e){var r=e/2,n=o(r);return[2*t/T*o(e)*n*n,T*g(r)]}function At(t){var e=1-t,r=i(y,0)[0]-i(-y,0)[0],n=L(2*(i(0,x)[1]-i(0,-x)[1])/r);function i(r,n){var i=o(n),a=m(n);return[i/(e+t*i)*r,e*n+t*a]}function a(t,e){var r=i(t,e);return[r[0]*n,r[1]/n]}function s(t){return a(0,t)[1]}return a.invert=function(r,i){var a=V(s,i);return[r/n*(t+e/o(a)),a]},a}function Mt(t){return[t[0]/2,S(g(t[1]/2*M))*A]}function St(t){return[2*t[0],2*i(m(t[1]*M))*A]}function Et(t,r){var i=2*y/r,s=t*t;function l(r,l){var c=e.geoAzimuthalEquidistantRaw(r,l),u=c[0],f=c[1],h=u*u+f*f;if(h>s){var d=L(h),g=a(f,u),b=i*p(g/i),_=g-b,w=t*o(_),T=(t*m(_)-_*m(w))/(x-w),k=Lt(_,T),A=(y-t)/Ct(k,w,y);u=d;var M,S=50;do{u-=M=(t+Ct(k,w,u)*A-d)/(k(u)*A)}while(n(M)>v&&--S>0);f=_*m(u),u<x&&(f-=T*(u-x));var E=m(b),C=o(b);c[0]=u*C-f*E,c[1]=u*E+f*C}return c}return l.invert=function(r,l){var c=r*r+l*l;if(c>s){var u=L(c),f=a(l,r),h=i*p(f/i),d=f-h;r=u*o(d),l=u*m(d);for(var g=r-x,v=m(r),b=l/v,_=r<x?1/0:0,w=10;;){var T=t*m(b),k=t*o(b),A=m(k),M=x-k,S=(T-b*A)/M,E=Lt(b,S);if(n(_)<1e-12||!--w)break;b-=_=(b*v-S*g-l)/(v-2*g*(M*(k+b*T*o(k)-A)-T*(T-b*A))/(M*M))}r=(u=t+Ct(E,k,r)*(y-t)/Ct(E,k,y))*o(f=h+b),l=u*m(f)}return e.geoAzimuthalEquidistantRaw.invert(r,l)},l}function Lt(t,e){return function(r){var n=t*o(r);return r<x&&(n-=e),L(1+n*n)}}function Ct(t,e,r){for(var n=(r-e)/50,i=t(e)+t(r),a=1,o=e;a<50;++a)i+=2*t(o+=n);return.5*i*n}function Pt(t,e,r,i,a,s,l,c){function u(n,u){if(!u)return[t*n/y,0];var f=u*u,h=t+f*(e+f*(r+f*i)),p=u*(a-1+f*(s-c+f*l)),d=(h*h+p*p)/(2*p),g=n*S(h/d)/y;return[d*m(g),u*(1+f*c)+d*(1-o(g))]}return arguments.length<8&&(c=0),u.invert=function(u,f){var h,p,d=y*u/t,g=f,x=50;do{var b=g*g,_=t+b*(e+b*(r+b*i)),w=g*(a-1+b*(s-c+b*l)),T=_*_+w*w,k=2*w,A=T/k,M=A*A,E=S(_/A)/y,C=d*E,P=_*_,I=(2*e+b*(4*r+6*b*i))*g,O=a+b*(3*s+5*b*l),z=(2*(_*I+w*(O-1))*k-T*(2*(O-1)))/(k*k),D=o(C),R=m(C),F=A*D,B=A*R,N=d/y*(1/L(1-P/M))*(I*A-_*z)/M,j=B-u,U=g*(1+b*c)+A-F-f,V=z*R+F*N,H=F*E,q=1+z-(z*D-B*N),G=B*E,Y=V*G-q*H;if(!Y)break;d-=h=(U*V-j*q)/Y,g-=p=(j*G-U*H)/Y}while((n(h)>v||n(p)>v)&&--x>0);return[d,g]},u}Tt.invert=function(t,e){var r=e/(1+wt);return[t&&t/(wt*L(1-r*r)),2*i(r)]},kt.invert=function(t,e){var r=i(e/T),n=o(r),a=2*r;return[t*T/2/(o(a)*n*n),a]};var It=Pt(2.8284,-1.6988,.75432,-.18071,1.76003,-.38914,.042555);var Ot=Pt(2.583819,-.835827,.170354,-.038094,1.543313,-.411435,.082742);var zt=Pt(5/6*y,-.62636,-.0344,0,1.3493,-.05524,0,.045);function Dt(t,e){var r=t*t,n=e*e;return[t*(1-.162388*n)*(.87-952426e-9*r*r),e*(1+n/12)]}Dt.invert=function(t,e){var r,i=t,a=e,o=50;do{var s=a*a;a-=r=(a*(1+s/12)-e)/(1+s/4)}while(n(r)>v&&--o>0);o=50,t/=1-.162388*s;do{var l=(l=i*i)*l;i-=r=(i*(.87-952426e-9*l)-t)/(.87-.00476213*l)}while(n(r)>v&&--o>0);return[i,a]};var Rt=Pt(2.6516,-.76534,.19123,-.047094,1.36289,-.13965,.031762);function Ft(t){var e=t(x,0)[0]-t(-x,0)[0];function r(r,n){var i=r>0?-.5:.5,a=t(r+i*y,n);return a[0]-=i*e,a}return t.invert&&(r.invert=function(r,n){var i=r>0?-.5:.5,a=t.invert(r+i*e,n),o=a[0]-i*y;return o<-y?o+=2*y:o>y&&(o-=2*y),a[0]=o,a}),r}function Bt(t,e){var r=d(t),i=d(e),s=o(e),l=o(t)*s,c=m(t)*s,u=m(i*e);t=n(a(c,u)),e=S(l),n(t-x)>v&&(t%=x);var f=function(t,e){if(e===x)return[0,0];var r,i,a=m(e),s=a*a,l=s*s,c=1+l,u=1+3*l,f=1-l,h=S(1/L(c)),p=f+s*c*h,d=(1-a)/p,g=L(d),b=d*c,_=L(b),w=g*f;if(0===t)return[0,-(w+s*_)];var T,k=o(e),A=1/k,M=2*a*k,E=(-p*k-(-3*s+h*u)*M*(1-a))/(p*p),C=-A*M,P=-A*(s*c*E+d*u*M),I=-2*A*(f*(.5*E/g)-2*s*g*M),O=4*t/y;if(t>.222*y||e<y/4&&t>.175*y){if(r=(w+s*L(b*(1+l)-w*w))/(1+l),t>y/4)return[r,r];var z=r,D=.5*r;r=.5*(D+z),i=50;do{var R=L(b-r*r),F=r*(I+C*R)+P*S(r/_)-O;if(!F)break;F<0?D=r:z=r,r=.5*(D+z)}while(n(z-D)>v&&--i>0)}else{r=v,i=25;do{var B=r*r,N=L(b-B),j=I+C*N,U=r*j+P*S(r/_)-O,V=j+(P-C*B)/N;r-=T=N?U/V:0}while(n(T)>v&&--i>0)}return[r,-w-s*L(b-r*r)]}(t>y/4?x-t:t,e);return t>y/4&&(u=f[0],f[0]=-f[1],f[1]=-u),f[0]*=r,f[1]*=-i,f}function Nt(t,e){var r,a,l,c,u,f;if(e<v)return[(c=m(t))-(r=e*(t-c*(a=o(t)))/4)*a,a+r*c,1-e*c*c/2,t-r];if(e>=1-v)return r=(1-e)/4,l=1/(a=P(t)),[(c=((f=s(2*(f=t)))-1)/(f+1))+r*((u=a*C(t))-t)/(a*a),l-r*c*l*(u-t),l+r*c*l*(u+t),2*i(s(t))-x+r*(u-t)/a];var h=[1,0,0,0,0,0,0,0,0],p=[L(e),0,0,0,0,0,0,0,0],d=0;for(a=L(1-e),u=1;n(p[d]/h[d])>v&&d<8;)r=h[d++],p[d]=(r-a)/2,h[d]=(r+a)/2,a=L(r*a),u*=2;l=u*h[d]*t;do{l=(S(c=p[d]*m(a=l)/h[d])+l)/2}while(--d);return[m(l),c=o(l),c/o(l-a),l]}function jt(t,e){if(!e)return t;if(1===e)return c(g(t/2+b));for(var r=1,a=L(1-e),o=L(e),s=0;n(o)>v;s++){if(t%y){var l=i(a*g(t)/r);l<0&&(l+=y),t+=l+~~(t/y)*y}else t+=t;o=(r+a)/2,a=L(r*a),o=((r=o)-a)/2}return t/(h(2,s)*r)}function Ut(t,e){var r=(w-1)/(w+1),l=L(1-r*r),u=jt(x,l*l),f=c(g(y/4+n(e)/2)),h=s(-1*f)/L(r),p=function(t,e){var r=t*t,n=e+1,i=1-r-e*e;return[.5*((t>=0?x:-x)-a(i,2*t)),-.25*c(i*i+4*r)+.5*c(n*n+r)]}(h*o(-1*t),h*m(-1*t)),v=function(t,e,r){var a=n(t),o=C(n(e));if(a){var s=1/m(a),l=1/(g(a)*g(a)),c=-(l+r*(o*o*s*s)-1+r),u=(-c+L(c*c-4*((r-1)*l)))/2;return[jt(i(1/L(u)),r)*d(t),jt(i(L((u/l-1)/r)),1-r)*d(e)]}return[0,jt(i(o),1-r)*d(e)]}(p[0],p[1],l*l);return[-v[1],(e>=0?1:-1)*(.5*u-v[0])]}function Vt(t){var e=m(t),r=o(t),i=Ht(t);function s(t,a){var s=i(t,a);t=s[0],a=s[1];var l=m(a),c=o(a),u=o(t),f=E(e*l+r*c*u),h=m(f),p=n(h)>v?f/h:1;return[p*r*m(t),(n(t)>x?p:-p)*(e*c-r*l*u)]}return i.invert=Ht(-t),s.invert=function(t,r){var n=L(t*t+r*r),s=-m(n),l=o(n),c=n*l,u=-r*s,f=n*e,h=L(c*c+u*u-f*f),p=a(c*f+u*h,u*f-c*h),d=(n>x?-1:1)*a(t*s,n*o(p)*l+r*m(p)*s);return i.invert(d,p)},s}function Ht(t){var e=m(t),r=o(t);return function(t,n){var i=o(n),s=o(t)*i,l=m(t)*i,c=m(n);return[a(l,s*r-c*e),S(c*r+s*e)]}}Bt.invert=function(t,e){n(t)>1&&(t=2*d(t)-t),n(e)>1&&(e=2*d(e)-e);var r=d(t),i=d(e),s=-r*t,l=-i*e,c=l/s<1,u=function(t,e){var r=0,i=1,a=.5,s=50;for(;;){var l=a*a,c=L(a),u=S(1/L(1+l)),f=1-l+a*(1+l)*u,h=(1-c)/f,p=L(h),d=h*(1+l),m=p*(1-l),g=L(d-t*t),v=e+m+a*g;if(n(i-r)<1e-12||0==--s||0===v)break;v>0?r=a:i=a,a=.5*(r+i)}if(!s)return null;var x=S(c),b=o(x),_=1/b,w=2*c*b,T=(-f*b-(-3*a+u*(1+3*l))*w*(1-c))/(f*f);return[y/4*(t*(-2*_*(.5*T/p*(1-l)-2*a*p*w)+-_*w*g)+-_*(a*(1+l)*T+h*(1+3*l)*w)*S(t/L(d))),x]}(c?l:s,c?s:l),f=u[0],h=u[1],p=o(h);return c&&(f=-x-f),[r*(a(m(f)*p,-m(h))+y),i*S(o(f)*p)]},Ut.invert=function(t,e){var r,n,o,l,u,f,h=(w-1)/(w+1),p=L(1-h*h),d=jt(x,p*p),m=(n=-t,o=p*p,(r=.5*d-e)?(l=Nt(r,o),n?(f=(u=Nt(n,1-o))[1]*u[1]+o*l[0]*l[0]*u[0]*u[0],[[l[0]*u[2]/f,l[1]*l[2]*u[0]*u[1]/f],[l[1]*u[1]/f,-l[0]*l[2]*u[0]*u[2]/f],[l[2]*u[1]*u[2]/f,-o*l[0]*l[1]*u[0]/f]]):[[l[0],0],[l[1],0],[l[2],0]]):[[0,(u=Nt(n,1-o))[0]/u[1]],[1/u[1],0],[u[2]/u[1],0]]),g=function(t,e){var r=e[0]*e[0]+e[1]*e[1];return[(t[0]*e[0]+t[1]*e[1])/r,(t[1]*e[0]-t[0]*e[1])/r]}(m[0],m[1]);return[a(g[1],g[0])/-1,2*i(s(-.5*c(h*g[0]*g[0]+h*g[1]*g[1])))-x]};var qt=S(1-1/3)*A,Gt=ht(0);function Yt(t){var e=qt*M,r=lt(y,e)[0]-lt(-y,e)[0],i=Gt(0,e)[1],a=lt(0,e)[1],o=T-a,s=k/t,c=4/k,h=i+o*o*4/k;function p(p,d){var m,g=n(d);if(g>e){var v=f(t-1,u(0,l((p+y)/s)));(m=lt(p+=y*(t-1)/t-v*s,g))[0]=m[0]*k/r-k*(t-1)/(2*t)+v*k/t,m[1]=i+4*(m[1]-a)*o/k,d<0&&(m[1]=-m[1])}else m=Gt(p,d);return m[0]*=c,m[1]/=h,m}return p.invert=function(e,p){e/=c;var d=n(p*=h);if(d>i){var m=f(t-1,u(0,l((e+y)/s)));e=(e+y*(t-1)/t-m*s)*r/k;var g=lt.invert(e,.25*(d-i)*k/o+a);return g[0]-=y*(t-1)/t-m*s,p<0&&(g[1]=-g[1]),g}return Gt.invert(e,p)},p}function Wt(t,e){return[t,1&e?90-v:qt]}function Xt(t,e){return[t,1&e?-90+v:-qt]}function Zt(t){return[t[0]*(1-v),t[1]]}function Jt(t){var e,r=1+t,i=S(m(1/r)),s=2*L(y/(e=y+4*i*r)),l=.5*s*(r+L(t*(2+t))),c=t*t,u=r*r;function f(f,h){var p,d,g=1-m(h);if(g&&g<2){var v,b=x-h,_=25;do{var w=m(b),T=o(b),k=i+a(w,r-T),A=1+u-2*r*T;b-=v=(b-c*i-r*w+A*k-.5*g*e)/(2*r*w*k)}while(n(v)>1e-12&&--_>0);p=s*L(A),d=f*k/y}else p=s*(t+g),d=f*i/y;return[p*m(d),l-p*o(d)]}return f.invert=function(t,n){var o=t*t+(n-=l)*n,f=(1+u-o/(s*s))/(2*r),h=E(f),p=m(h),d=i+a(p,r-f);return[S(t/L(o))*y/d,S(1-2*(h-c*i-r*p+(1+u-2*r*f)*d)/e)]},f}function Kt(t,e){return e>-.7109889596207567?((t=W(t,e))[1]+=.0528035274542,t):Q(t,e)}function Qt(t,e){return n(e)>.7109889596207567?((t=W(t,e))[1]-=e>0?.0528035274542:-.0528035274542,t):Q(t,e)}function $t(t,e,r,n){var i=L(4*y/(2*r+(1+t-e/2)*m(2*r)+(t+e)/2*m(4*r)+e/2*m(6*r))),a=L(n*m(r)*L((1+t*o(2*r)+e*o(4*r))/(1+t+e))),s=r*c(1);function l(r){return L(1+t*o(2*r)+e*o(4*r))}function c(n){var i=n*r;return(2*i+(1+t-e/2)*m(2*i)+(t+e)/2*m(4*i)+e/2*m(6*i))/r}function u(t){return l(t)*m(t)}var f=function(t,e){var n=r*V(c,s*m(e)/r,e/y);isNaN(n)&&(n=r*d(e));var u=i*l(n);return[u*a*t/y*o(n),u/a*m(n)]};return f.invert=function(t,e){var n=V(u,e*a/i);return[t*y/(o(n)*i*a*l(n)),S(r*c(n/r)/s)]},0===r&&(i=L(n/y),(f=function(t,e){return[t*i,m(e)/i]}).invert=function(t,e){return[t/i,S(e*i)]}),f}function te(t,e,r,n,i){void 0===n&&(n=1e-8),void 0===i&&(i=20);var a=t(e),o=t(.5*(e+r)),s=t(r);return function t(e,r,n,i,a,o,s,l,c,u,f){if(f.nanEncountered)return NaN;var h,p,d,m,g,v,y,x,b,_;if(p=e(r+.25*(h=n-r)),d=e(n-.25*h),isNaN(p))f.nanEncountered=!0;else{if(!isNaN(d))return _=((v=(m=h*(i+4*p+a)/12)+(g=h*(a+4*d+o)/12))-s)/15,u>c?(f.maxDepthCount++,v+_):Math.abs(_)<l?v+_:(x=t(e,r,y=r+.5*h,i,p,a,m,.5*l,c,u+1,f),isNaN(x)?(f.nanEncountered=!0,NaN):(b=t(e,y,n,a,d,o,g,.5*l,c,u+1,f),isNaN(b)?(f.nanEncountered=!0,NaN):x+b));f.nanEncountered=!0}}(t,e,r,a,o,s,(a+4*o+s)*(r-e)/6,n,i,1,{maxDepthCount:0,nanEncountered:!1})}function ee(t,e,r){function i(r){return t+(1-t)*h(1-h(r,e),1/e)}function a(t){return te(i,0,t,1e-4)}for(var o=1/a(1),s=1e3,l=(1+1e-8)*o,c=[],u=0;u<=s;u++)c.push(a(u/s)*l);function f(t){var e=0,r=s,n=500;do{c[n]>t?r=n:e=n,n=e+r>>1}while(n>e);var i=c[n+1]-c[n];return i&&(i=(t-c[n+1])/i),(n+1+i)/s}var p=2*f(1)/y*o/r,g=function(t,e){var r=f(n(m(e))),a=i(r)*t;return r/=p,[a,e>=0?r:-r]};return g.invert=function(t,e){var r;return n(e*=p)<1&&(r=d(e)*S(a(n(e))*o)),[t/i(n(e)),r]},g}function re(t,e){return n(t[0]-e[0])<v&&n(t[1]-e[1])<v}function ne(t,e){for(var r,n,i,a=-1,o=t.length,s=t[0],l=[];++a<o;){n=((r=t[a])[0]-s[0])/e,i=(r[1]-s[1])/e;for(var c=0;c<e;++c)l.push([s[0]+c*n,s[1]+c*i]);s=r}return l.push(r),l}function ie(t){var e,n,i,a,o,s,l,c=[],u=t[0].length;for(l=0;l<u;++l)n=(e=t[0][l])[0][0],i=e[0][1],a=e[1][1],o=e[2][0],s=e[2][1],c.push(ne([[n+v,i+v],[n+v,a-v],[o-v,a-v],[o-v,s+v]],30));for(l=t[1].length-1;l>=0;--l)n=(e=t[1][l])[0][0],i=e[0][1],a=e[1][1],o=e[2][0],s=e[2][1],c.push(ne([[o-v,s-v],[o-v,a+v],[n+v,a+v],[n+v,i-v]],30));return{type:"Polygon",coordinates:[r.merge(c)]}}function ae(t,r,n){var i,a;function o(e,n){for(var i=n<0?-1:1,a=r[+(n<0)],o=0,s=a.length-1;o<s&&e>a[o][2][0];++o);var l=t(e-a[o][1][0],n);return l[0]+=t(a[o][1][0],i*n>i*a[o][0][1]?a[o][0][1]:n)[0],l}n?o.invert=n(o):t.invert&&(o.invert=function(e,n){for(var i=a[+(n<0)],s=r[+(n<0)],l=0,c=i.length;l<c;++l){var u=i[l];if(u[0][0]<=e&&e<u[1][0]&&u[0][1]<=n&&n<u[1][1]){var f=t.invert(e-t(s[l][1][0],0)[0],n);return f[0]+=s[l][1][0],re(o(f[0],f[1]),[e,n])?f:null}}});var s=e.geoProjection(o),l=s.stream;return s.stream=function(t){var r=s.rotate(),n=l(t),a=(s.rotate([0,0]),l(t));return s.rotate(r),n.sphere=function(){e.geoStream(i,a)},n},s.lobes=function(e){return arguments.length?(i=ie(e),r=e.map((function(t){return t.map((function(t){return[[t[0][0]*M,t[0][1]*M],[t[1][0]*M,t[1][1]*M],[t[2][0]*M,t[2][1]*M]]}))})),a=r.map((function(e){return e.map((function(e){var r,n=t(e[0][0],e[0][1])[0],i=t(e[2][0],e[2][1])[0],a=t(e[1][0],e[0][1])[1],o=t(e[1][0],e[1][1])[1];return a>o&&(r=a,a=o,o=r),[[n,a],[i,o]]}))})),s):r.map((function(t){return t.map((function(t){return[[t[0][0]*A,t[0][1]*A],[t[1][0]*A,t[1][1]*A],[t[2][0]*A,t[2][1]*A]]}))}))},null!=r&&s.lobes(r),s}Kt.invert=function(t,e){return e>-.7109889596207567?W.invert(t,e-.0528035274542):Q.invert(t,e)},Qt.invert=function(t,e){return n(e)>.7109889596207567?W.invert(t,e+(e>0?.0528035274542:-.0528035274542)):Q.invert(t,e)};var oe=[[[[-180,0],[-100,90],[-40,0]],[[-40,0],[30,90],[180,0]]],[[[-180,0],[-160,-90],[-100,0]],[[-100,0],[-60,-90],[-20,0]],[[-20,0],[20,-90],[80,0]],[[80,0],[140,-90],[180,0]]]];var se=[[[[-180,0],[-100,90],[-40,0]],[[-40,0],[30,90],[180,0]]],[[[-180,0],[-160,-90],[-100,0]],[[-100,0],[-60,-90],[-20,0]],[[-20,0],[20,-90],[80,0]],[[80,0],[140,-90],[180,0]]]];var le=[[[[-180,0],[-100,90],[-40,0]],[[-40,0],[30,90],[180,0]]],[[[-180,0],[-160,-90],[-100,0]],[[-100,0],[-60,-90],[-20,0]],[[-20,0],[20,-90],[80,0]],[[80,0],[140,-90],[180,0]]]];var ce=[[[[-180,0],[-90,90],[0,0]],[[0,0],[90,90],[180,0]]],[[[-180,0],[-90,-90],[0,0]],[[0,0],[90,-90],[180,0]]]];var ue=[[[[-180,35],[-30,90],[0,35]],[[0,35],[30,90],[180,35]]],[[[-180,-10],[-102,-90],[-65,-10]],[[-65,-10],[5,-90],[77,-10]],[[77,-10],[103,-90],[180,-10]]]];var fe=[[[[-180,0],[-110,90],[-40,0]],[[-40,0],[0,90],[40,0]],[[40,0],[110,90],[180,0]]],[[[-180,0],[-110,-90],[-40,0]],[[-40,0],[0,-90],[40,0]],[[40,0],[110,-90],[180,0]]]];function he(t,e){return[3/k*t*L(y*y/3-e*e),e]}function pe(t){function e(e,r){if(n(n(r)-x)<v)return[0,r<0?-2:2];var i=m(r),a=h((1+i)/(1-i),t/2),s=.5*(a+1/a)+o(e*=t);return[2*m(e)/s,(a-1/a)/s]}return e.invert=function(e,r){var i=n(r);if(n(i-2)<v)return e?null:[0,d(r)*x];if(i>2)return null;var o=(e/=2)*e,s=(r/=2)*r,l=2*r/(1+o+s);return l=h((1+l)/(1-l),1/t),[a(2*e,1-o-s)/t,S((l-1)/(l+1))]},e}he.invert=function(t,e){return[k/3*t/L(y*y/3-e*e),e]};var de=y/w;function me(t,e){return[t*(1+L(o(e)))/2,e/(o(e/2)*o(t/6))]}function ge(t,e){var r=t*t,n=e*e;return[t*(.975534+n*(-.0143059*r-.119161+-.0547009*n)),e*(1.00384+r*(.0802894+-.02855*n+199025e-9*r)+n*(.0998909+-.0491032*n))]}function ve(t,e){return[m(t)/o(e),g(e)*o(t)]}function ye(t){var e=o(t),r=g(b+t/2);function i(i,a){var o=a-t,s=n(o)<v?i*e:n(s=b+a/2)<v||n(n(s)-x)<v?0:i*o/c(g(s)/r);return[s,o]}return i.invert=function(i,a){var o,s=a+t;return[n(a)<v?i/e:n(o=b+s/2)<v||n(n(o)-x)<v?0:i*c(g(o)/r)/a,s]},i}function xe(t,e){return[t,1.25*c(g(b+.4*e))]}function be(t){var e=t.length-1;function r(r,n){for(var i,a=o(n),s=2/(1+a*o(r)),l=s*a*m(r),c=s*m(n),u=e,f=t[u],h=f[0],p=f[1];--u>=0;)h=(f=t[u])[0]+l*(i=h)-c*p,p=f[1]+l*p+c*i;return[h=l*(i=h)-c*p,p=l*p+c*i]}return r.invert=function(r,s){var l=20,c=r,u=s;do{for(var f,h=e,p=t[h],d=p[0],g=p[1],v=0,y=0;--h>=0;)v=d+c*(f=v)-u*y,y=g+c*y+u*f,d=(p=t[h])[0]+c*(f=d)-u*g,g=p[1]+c*g+u*f;var x,b,_=(v=d+c*(f=v)-u*y)*v+(y=g+c*y+u*f)*y;c-=x=((d=c*(f=d)-u*g-r)*v+(g=c*g+u*f-s)*y)/_,u-=b=(g*v-d*y)/_}while(n(x)+n(b)>1e-12&&--l>0);if(l){var w=L(c*c+u*u),T=2*i(.5*w),k=m(T);return[a(c*k,w*o(T)),w?S(u*k/w):0]}},r}me.invert=function(t,e){var r=n(t),i=n(e),a=v,s=x;i<de?s*=i/de:a+=6*E(de/i);for(var l=0;l<25;l++){var c=m(s),u=L(o(s)),f=m(s/2),h=o(s/2),p=m(a/6),d=o(a/6),g=.5*a*(1+u)-r,y=s/(h*d)-i,b=u?-.25*a*c/u:0,_=.5*(1+u),w=(1+.5*s*f/h)/(h*d),T=s/h*(p/6)/(d*d),k=b*T-w*_,A=(g*T-y*_)/k,M=(y*b-g*w)/k;if(s-=A,a-=M,n(A)<v&&n(M)<v)break}return[t<0?-a:a,e<0?-s:s]},ge.invert=function(t,e){var r=d(t)*y,i=e/2,a=50;do{var o=r*r,s=i*i,l=r*i,c=r*(.975534+s*(-.0143059*o-.119161+-.0547009*s))-t,u=i*(1.00384+o*(.0802894+-.02855*s+199025e-9*o)+s*(.0998909+-.0491032*s))-e,f=.975534-s*(.119161+3*o*.0143059+.0547009*s),h=-l*(.238322+.2188036*s+.0286118*o),p=l*(.1605788+7961e-7*o+-.0571*s),m=1.00384+o*(.0802894+199025e-9*o)+s*(3*(.0998909-.02855*o)-.245516*s),g=h*p-m*f,x=(u*h-c*m)/g,b=(c*p-u*f)/g;r-=x,i-=b}while((n(x)>v||n(b)>v)&&--a>0);return a&&[r,i]},ve.invert=function(t,e){var r=t*t,n=e*e+1,i=r+n,a=t?_*L((i-L(i*i-4*r))/r):1/L(n);return[S(t*a),d(e)*E(a)]},xe.invert=function(t,e){return[t,2.5*i(s(.8*e))-.625*y]};var _e=[[.9972523,0],[.0052513,-.0041175],[.0074606,.0048125],[-.0153783,-.1968253],[.0636871,-.1408027],[.3660976,-.2937382]],we=[[.98879,0],[0,0],[-.050909,0],[0,0],[.075528,0]],Te=[[.984299,0],[.0211642,.0037608],[-.1036018,-.0575102],[-.0329095,-.0320119],[.0499471,.1223335],[.026046,.0899805],[7388e-7,-.1435792],[.0075848,-.1334108],[-.0216473,.0776645],[-.0225161,.0853673]],ke=[[.9245,0],[0,0],[.01943,0]],Ae=[[.721316,0],[0,0],[-.00881625,-.00617325]];function Me(t,r){var n=e.geoProjection(be(t)).rotate(r).clipAngle(90),i=e.geoRotation(r),a=n.center;return delete n.rotate,n.center=function(t){return arguments.length?a(i(t)):i.invert(a())},n}var Se=L(6),Ee=L(7);function Le(t,e){var r=S(7*m(e)/(3*Se));return[Se*t*(2*o(2*r/3)-1)/Ee,9*m(r/3)/Ee]}function Ce(t,e){for(var r,i=(1+_)*m(e),a=e,s=0;s<25&&(a-=r=(m(a/2)+m(a)-i)/(.5*o(a/2)+o(a)),!(n(r)<v));s++);return[t*(1+2*o(a)/o(a/2))/(3*w),2*L(3)*m(a/2)/L(2+w)]}function Pe(t,e){for(var r,i=L(6/(4+y)),a=(1+y/4)*m(e),s=e/2,l=0;l<25&&(s-=r=(s/2+m(s)-a)/(.5+o(s)),!(n(r)<v));l++);return[i*(.5+o(s))*t/1.5,i*s]}function Ie(t,e){var r=e*e,n=r*r,i=r*n;return[t*(.84719-.13063*r+i*i*(.05494*r-.04515-.02326*n+.00331*i)),e*(1.01183+n*n*(.01926*r-.02625-.00396*n))]}function Oe(t,e){return[t*(1+o(e))/2,2*(e-g(e/2))]}Le.invert=function(t,e){var r=3*S(e*Ee/9);return[t*Ee/(Se*(2*o(2*r/3)-1)),S(3*m(r)*Se/7)]},Ce.invert=function(t,e){var r=e*L(2+w)/(2*L(3)),n=2*S(r);return[3*w*t/(1+2*o(n)/o(n/2)),S((r+m(n))/(1+_))]},Pe.invert=function(t,e){var r=L(6/(4+y)),i=e/r;return n(n(i)-x)<v&&(i=i<0?-x:x),[1.5*t/(r*(.5+o(i))),S((i/2+m(i))/(1+y/4))]},Ie.invert=function(t,e){var r,i,a,o,s=e,l=25;do{s-=r=(s*(1.01183+(a=(i=s*s)*i)*a*(.01926*i-.02625-.00396*a))-e)/(1.01183+a*a*(.21186*i-.23625+-.05148*a))}while(n(r)>1e-12&&--l>0);return[t/(.84719-.13063*(i=s*s)+(o=i*(a=i*i))*o*(.05494*i-.04515-.02326*a+.00331*o)),s]},Oe.invert=function(t,e){for(var r=e/2,i=0,a=1/0;i<10&&n(a)>v;++i){var s=o(e/2);e-=a=(e-g(e/2)-r)/(1-.5/(s*s))}return[2*t/(1+o(e)),e]};var ze=[[[[-180,0],[-90,90],[0,0]],[[0,0],[90,90],[180,0]]],[[[-180,0],[-90,-90],[0,0]],[[0,0],[90,-90],[180,0]]]];function De(t,e){var r=m(e),i=o(e),a=d(t);if(0===t||n(e)===x)return[0,e];if(0===e)return[t,0];if(n(t)===x)return[t*i,x*r];var s=y/(2*t)-2*t/y,l=2*e/y,c=(1-l*l)/(r-l),u=s*s,f=c*c,h=1+u/f,p=1+f/u,g=(s*r/c-s/2)/h,v=(f*r/u+c/2)/p,b=v*v-(f*r*r/u+c*r-1)/p;return[x*(g+L(g*g+i*i/h)*a),x*(v+L(b<0?0:b)*d(-e*s)*a)]}De.invert=function(t,e){var r=(t/=x)*t,n=r+(e/=x)*e,i=y*y;return[t?(n-1+L((1-n)*(1-n)+4*r))/(2*t)*x:0,V((function(t){return n*(y*m(t)-2*t)*y+4*t*t*(e-m(t))+2*y*t-i*e}),0)]};function Re(t,e){var r=e*e;return[t,e*(1.0148+r*r*(.23185+r*(.02406*r-.14499)))]}function Fe(t,e){if(n(e)<v)return[t,0];var r=g(e),i=t*m(e);return[m(i)/r,e+(1-o(i))/r]}function Be(t,e){var r=je(t[1],t[0]),n=je(e[1],e[0]),i=function(t,e){return a(t[0]*e[1]-t[1]*e[0],t[0]*e[0]+t[1]*e[1])}(r,n),s=Ue(r)/Ue(n);return Ne([1,0,t[0][0],0,1,t[0][1]],Ne([s,0,0,0,s,0],Ne([o(i),m(i),0,-m(i),o(i),0],[1,0,-e[0][0],0,1,-e[0][1]])))}function Ne(t,e){return[t[0]*e[0]+t[1]*e[3],t[0]*e[1]+t[1]*e[4],t[0]*e[2]+t[1]*e[5]+t[2],t[3]*e[0]+t[4]*e[3],t[3]*e[1]+t[4]*e[4],t[3]*e[2]+t[4]*e[5]+t[5]]}function je(t,e){return[t[0]-e[0],t[1]-e[1]]}function Ue(t){return L(t[0]*t[0]+t[1]*t[1])}function Ve(t,r,i){function a(t,e){var n,i=r(t,e),a=i.project([t*A,e*A]);return(n=i.transform)?[n[0]*a[0]+n[1]*a[1]+n[2],-(n[3]*a[0]+n[4]*a[1]+n[5])]:(a[1]=-a[1],a)}!function t(e,r){if(e.edges=function(t){for(var e=t.length,r=[],n=t[e-1],i=0;i<e;++i)r.push([n,n=t[i]]);return r}(e.face),r.face){var n=e.shared=function(t,e){for(var r,n,i=t.length,a=null,o=0;o<i;++o){r=t[o];for(var s=e.length;--s>=0;)if(n=e[s],r[0]===n[0]&&r[1]===n[1]){if(a)return[a,r];a=r}}}(e.face,r.face),i=Be(n.map(r.project),n.map(e.project));e.transform=r.transform?Ne(r.transform,i):i;for(var a=r.edges,o=0,s=a.length;o<s;++o)He(n[0],a[o][1])&&He(n[1],a[o][0])&&(a[o]=e),He(n[0],a[o][0])&&He(n[1],a[o][1])&&(a[o]=e);for(a=e.edges,o=0,s=a.length;o<s;++o)He(n[0],a[o][0])&&He(n[1],a[o][1])&&(a[o]=r),He(n[0],a[o][1])&&He(n[1],a[o][0])&&(a[o]=r)}else e.transform=r.transform;e.children&&e.children.forEach((function(r){t(r,e)}));return e}(t,{transform:null}),qe(t)&&(a.invert=function(e,n){var i=function t(e,n){var i=e.project.invert,a=e.transform,o=n;a&&(a=function(t){var e=1/(t[0]*t[4]-t[1]*t[3]);return[e*t[4],-e*t[1],e*(t[1]*t[5]-t[2]*t[4]),-e*t[3],e*t[0],e*(t[2]*t[3]-t[0]*t[5])]}(a),o=[a[0]*o[0]+a[1]*o[1]+a[2],a[3]*o[0]+a[4]*o[1]+a[5]]);if(i&&e===function(t){return r(t[0]*M,t[1]*M)}(s=i(o)))return s;for(var s,l=e.children,c=0,u=l&&l.length;c<u;++c)if(s=t(l[c],n))return s}(t,[e,-n]);return i&&(i[0]*=M,i[1]*=M,i)});var o=e.geoProjection(a),s=o.stream;return o.stream=function(r){var i=o.rotate(),a=s(r),l=(o.rotate([0,0]),s(r));return o.rotate(i),a.sphere=function(){l.polygonStart(),l.lineStart(),function t(r,i,a){var o,s,l=i.edges,c=l.length,u={type:"MultiPoint",coordinates:i.face},f=i.face.filter((function(t){return 90!==n(t[1])})),h=e.geoBounds({type:"MultiPoint",coordinates:f}),p=!1,d=-1,m=h[1][0]-h[0][0],g=180===m||360===m?[(h[0][0]+h[1][0])/2,(h[0][1]+h[1][1])/2]:e.geoCentroid(u);if(a)for(;++d<c&&l[d]!==a;);++d;for(var y=0;y<c;++y)s=l[(y+d)%c],Array.isArray(s)?(p||(r.point((o=e.geoInterpolate(s[0],g)(v))[0],o[1]),p=!0),r.point((o=e.geoInterpolate(s[1],g)(v))[0],o[1])):(p=!1,s!==a&&t(r,s,i))}(l,t),l.lineEnd(),l.polygonEnd()},a},o.angle(null==i?-30:i*A)}function He(t,e){return t&&e&&t[0]===e[0]&&t[1]===e[1]}function qe(t){return t.project.invert||t.children&&t.children.some(qe)}Re.invert=function(t,e){e>1.790857183?e=1.790857183:e<-1.790857183&&(e=-1.790857183);var r,i=e;do{var a=i*i;i-=r=(i*(1.0148+a*a*(.23185+a*(.02406*a-.14499)))-e)/(1.0148+a*a*(5*.23185+a*(.21654*a-1.01493)))}while(n(r)>v);return[t,i]},Fe.invert=function(t,e){if(n(e)<v)return[t,0];var r,i=t*t+e*e,a=.5*e,s=10;do{var l=g(a),c=1/o(a),u=i-2*e*a+a*a;a-=r=(l*u+2*(a-e))/(2+u*c*c+2*(a-e)*l)}while(n(r)>v&&--s>0);return l=g(a),[(n(e)<n(a+1/l)?S(t*l):d(e)*d(t)*(E(n(t*l))+x))/m(a),a]};var Ge=[[0,90],[-90,0],[0,0],[90,0],[180,0],[0,-90]],Ye=[[0,2,1],[0,3,2],[5,1,2],[5,2,3],[0,1,4],[0,4,3],[5,4,1],[5,3,4]].map((function(t){return t.map((function(t){return Ge[t]}))}));var We=2/L(3);function Xe(t,e){var r=lt(t,e);return[r[0]*We,r[1]]}function Ze(t,e){for(var r=0,n=t.length,i=0;r<n;++r)i+=t[r]*e[r];return i}function Je(t){return[a(t[1],t[0])*A,S(u(-1,f(1,t[2])))*A]}function Ke(t){var e=t[0]*M,r=t[1]*M,n=o(r);return[n*o(e),n*m(e),m(r)]}function Qe(){}function $e(t,e){return{type:"FeatureCollection",features:t.features.map((function(t){return tr(t,e)}))}}function tr(t,e){return{type:"Feature",id:t.id,properties:t.properties,geometry:er(t.geometry,e)}}function er(t,r){if(!t)return null;if("GeometryCollection"===t.type)return function(t,e){return{type:"GeometryCollection",geometries:t.geometries.map((function(t){return er(t,e)}))}}(t,r);var n;switch(t.type){case"Point":case"MultiPoint":n=ir;break;case"LineString":case"MultiLineString":n=ar;break;case"Polygon":case"MultiPolygon":case"Sphere":n=or;break;default:return null}return e.geoStream(t,r(n)),n.result()}Xe.invert=function(t,e){return lt.invert(t/We,e)};var rr=[],nr=[],ir={point:function(t,e){rr.push([t,e])},result:function(){var t=rr.length?rr.length<2?{type:"Point",coordinates:rr[0]}:{type:"MultiPoint",coordinates:rr}:null;return rr=[],t}},ar={lineStart:Qe,point:function(t,e){rr.push([t,e])},lineEnd:function(){rr.length&&(nr.push(rr),rr=[])},result:function(){var t=nr.length?nr.length<2?{type:"LineString",coordinates:nr[0]}:{type:"MultiLineString",coordinates:nr}:null;return nr=[],t}},or={polygonStart:Qe,lineStart:Qe,point:function(t,e){rr.push([t,e])},lineEnd:function(){var t=rr.length;if(t){do{rr.push(rr[0].slice())}while(++t<4);nr.push(rr),rr=[]}},polygonEnd:Qe,result:function(){if(!nr.length)return null;var t=[],e=[];return nr.forEach((function(r){!function(t){if((e=t.length)<4)return!1;for(var e,r=0,n=t[e-1][1]*t[0][0]-t[e-1][0]*t[0][1];++r<e;)n+=t[r-1][1]*t[r][0]-t[r-1][0]*t[r][1];return n<=0}(r)?e.push(r):t.push([r])})),e.forEach((function(e){var r=e[0];t.some((function(t){if(function(t,e){for(var r=e[0],n=e[1],i=!1,a=0,o=t.length,s=o-1;a<o;s=a++){var l=t[a],c=l[0],u=l[1],f=t[s],h=f[0],p=f[1];u>n^p>n&&r<(h-c)*(n-u)/(p-u)+c&&(i=!i)}return i}(t[0],r))return t.push(e),!0}))||t.push([e])})),nr=[],t.length?t.length>1?{type:"MultiPolygon",coordinates:t}:{type:"Polygon",coordinates:t[0]}:null}};function sr(t){var r=t(x,0)[0]-t(-x,0)[0];function i(e,i){var a=n(e)<x,o=t(a?e:e>0?e-y:e+y,i),s=(o[0]-o[1])*_,l=(o[0]+o[1])*_;if(a)return[s,l];var c=r*_,u=s>0^l>0?-1:1;return[u*s-d(l)*c,u*l-d(s)*c]}return t.invert&&(i.invert=function(e,i){var a=(e+i)*_,o=(i-e)*_,s=n(a)<.5*r&&n(o)<.5*r;if(!s){var l=r*_,c=a>0^o>0?-1:1,u=-c*e+(o>0?1:-1)*l,f=-c*i+(a>0?1:-1)*l;a=(-u-f)*_,o=(u-f)*_}var h=t.invert(a,o);return s||(h[0]+=a>0?y:-y),h}),e.geoProjection(i).rotate([-90,-90,45]).clipAngle(179.999)}function lr(){return sr(Ut).scale(111.48)}function cr(t){var e=m(t);function r(r,n){var a=e?g(r*e/2)/e:r/2;if(!n)return[2*a,-t];var s=2*i(a*m(n)),l=1/g(n);return[m(s)*l,n+(1-o(s))*l-t]}return r.invert=function(r,a){if(n(a+=t)<v)return[e?2*i(e*r/2)/e:r,0];var s,l=r*r+a*a,c=0,u=10;do{var f=g(c),h=1/o(c),p=l-2*a*c+c*c;c-=s=(f*p+2*(c-a))/(2+p*h*h+2*(c-a)*f)}while(n(s)>v&&--u>0);var d=r*(f=g(c)),x=g(n(a)<n(c+1/f)?.5*S(d):.5*E(d)+y/4)/m(c);return[e?2*i(e*x)/e:2*x,c]},r}var ur=[[.9986,-.062],[1,0],[.9986,.062],[.9954,.124],[.99,.186],[.9822,.248],[.973,.31],[.96,.372],[.9427,.434],[.9216,.4958],[.8962,.5571],[.8679,.6176],[.835,.6769],[.7986,.7346],[.7597,.7903],[.7186,.8435],[.6732,.8936],[.6213,.9394],[.5722,.9761],[.5322,1]];function fr(t,e){var r,i=f(18,36*n(e)/y),a=l(i),o=i-a,s=(r=ur[a])[0],c=r[1],u=(r=ur[++a])[0],h=r[1],p=(r=ur[f(19,++a)])[0],d=r[1];return[t*(u+o*(p-s)/2+o*o*(p-2*u+s)/2),(e>0?x:-x)*(h+o*(d-c)/2+o*o*(d-2*h+c)/2)]}function hr(t,e){var r=function(t){function e(e,r){var n=o(r),i=(t-1)/(t-n*o(e));return[i*n*m(e),i*m(r)]}return e.invert=function(e,r){var n=e*e+r*r,i=L(n),o=(t-L(1-n*(t+1)/(t-1)))/((t-1)/i+i/(t-1));return[a(e*o,i*L(1-o*o)),i?S(r*o/i):0]},e}(t);if(!e)return r;var n=o(e),i=m(e);function s(e,a){var o=r(e,a),s=o[1],l=s*i/(t-1)+n;return[o[0]*n/l,s/l]}return s.invert=function(e,a){var o=(t-1)/(t-1-a*i);return r.invert(o*e,o*a*n)},s}ur.forEach((function(t){t[1]*=1.0144})),fr.invert=function(t,e){var r=e/x,i=90*r,a=f(18,n(i/5)),o=u(0,l(a));do{var s=ur[o][1],c=ur[o+1][1],h=ur[f(19,o+2)][1],p=h-s,d=h-2*c+s,m=2*(n(r)-c)/p,g=d/p,v=m*(1-g*m*(1-2*g*m));if(v>=0||1===o){i=(e>=0?5:-5)*(v+a);var y,b=50;do{v=(a=f(18,n(i)/5))-(o=l(a)),s=ur[o][1],c=ur[o+1][1],h=ur[f(19,o+2)][1],i-=(y=(e>=0?x:-x)*(c+v*(h-s)/2+v*v*(h-2*c+s)/2)-e)*A}while(n(y)>1e-12&&--b>0);break}}while(--o>=0);var _=ur[o][0],w=ur[o+1][0],T=ur[f(19,o+2)][0];return[t/(w+v*(T-_)/2+v*v*(T-2*w+_)/2),i*M]};var pr=-179.9999,dr=179.9999,mr=-89.9999;function gr(t){return t.length>0}function vr(t){return-90===t||90===t?[0,t]:[-180,(e=t,Math.floor(1e4*e)/1e4)];var e}function yr(t){var e=t[0],r=t[1],n=!1;return e<=pr?(e=-180,n=!0):e>=dr&&(e=180,n=!0),r<=mr?(r=-90,n=!0):r>=89.9999&&(r=90,n=!0),n?[e,r]:t}function xr(t){return t.map(yr)}function br(t,e,r){for(var n=0,i=t.length;n<i;++n){var a=t[n].slice();r.push({index:-1,polygon:e,ring:a});for(var o=0,s=a.length;o<s;++o){var l=a[o],c=l[0],u=l[1];if(c<=pr||c>=dr||u<=mr||u>=89.9999){a[o]=yr(l);for(var f=o+1;f<s;++f){var h=a[f],p=h[0],d=h[1];if(p>pr&&p<dr&&d>mr&&d<89.9999)break}if(f===o+1)continue;if(o){var m={index:-1,polygon:e,ring:a.slice(0,o+1)};m.ring[m.ring.length-1]=vr(u),r[r.length-1]=m}else r.pop();if(f>=s)break;r.push({index:-1,polygon:e,ring:a=a.slice(f-1)}),a[0]=vr(a[0][1]),o=-1,s=a.length}}}}function _r(t){var e,r,n,i,a,o,s=t.length,l={},c={};for(e=0;e<s;++e)n=(r=t[e]).ring[0],a=r.ring[r.ring.length-1],n[0]!==a[0]||n[1]!==a[1]?(r.index=e,l[n]=c[a]=r):(r.polygon.push(r.ring),t[e]=null);for(e=0;e<s;++e)if(r=t[e]){if(n=r.ring[0],a=r.ring[r.ring.length-1],i=c[n],o=l[a],delete l[n],delete c[a],n[0]===a[0]&&n[1]===a[1]){r.polygon.push(r.ring);continue}i?(delete c[n],delete l[i.ring[0]],i.ring.pop(),t[i.index]=null,r={index:-1,polygon:i.polygon,ring:i.ring.concat(r.ring)},i===o?r.polygon.push(r.ring):(r.index=s++,t.push(l[r.ring[0]]=c[r.ring[r.ring.length-1]]=r))):o?(delete l[a],delete c[o.ring[o.ring.length-1]],r.ring.pop(),r={index:s++,polygon:o.polygon,ring:r.ring.concat(o.ring)},t[o.index]=null,t.push(l[r.ring[0]]=c[r.ring[r.ring.length-1]]=r)):(r.ring.push(r.ring[0]),r.polygon.push(r.ring))}}function wr(t){var e={type:"Feature",geometry:Tr(t.geometry)};return null!=t.id&&(e.id=t.id),null!=t.bbox&&(e.bbox=t.bbox),null!=t.properties&&(e.properties=t.properties),e}function Tr(t){if(null==t)return t;var e,r,n,i;switch(t.type){case"GeometryCollection":e={type:"GeometryCollection",geometries:t.geometries.map(Tr)};break;case"Point":e={type:"Point",coordinates:yr(t.coordinates)};break;case"MultiPoint":case"LineString":e={type:t.type,coordinates:xr(t.coordinates)};break;case"MultiLineString":e={type:"MultiLineString",coordinates:t.coordinates.map(xr)};break;case"Polygon":var a=[];br(t.coordinates,a,r=[]),_r(r),e={type:"Polygon",coordinates:a};break;case"MultiPolygon":r=[],n=-1,i=t.coordinates.length;for(var o=new Array(i);++n<i;)br(t.coordinates[n],o[n]=[],r);_r(r),e={type:"MultiPolygon",coordinates:o.filter(gr)};break;default:return t}return null!=t.bbox&&(e.bbox=t.bbox),e}function kr(t,e){var r=g(e/2),n=m(b*r);return[t*(.74482-.34588*n*n),1.70711*r]}function Ar(t,r,n){var i=e.geoInterpolate(r,n),a=i(.5),o=e.geoRotation([-a[0],-a[1]])(r),s=i.distance/2,l=-S(m(o[1]*M)/m(s)),c=[-a[0],-a[1],-(o[0]>0?y-l:l)*A],u=e.geoProjection(t(s)).rotate(c),f=e.geoRotation(c),h=u.center;return delete u.rotate,u.center=function(t){return arguments.length?h(f(t)):f.invert(h())},u.clipAngle(90)}function Mr(t){var r=o(t);function n(t,n){var i=e.geoGnomonicRaw(t,n);return i[0]*=r,i}return n.invert=function(t,n){return e.geoGnomonicRaw.invert(t/r,n)},n}function Sr(t,e){return Ar(Mr,t,e)}function Er(t){if(!(t*=2))return e.geoAzimuthalEquidistantRaw;var r=-t/2,n=-r,i=t*t,s=g(n),l=.5/m(n);function c(e,a){var s=E(o(a)*o(e-r)),l=E(o(a)*o(e-n));return[((s*=s)-(l*=l))/(2*t),(a<0?-1:1)*L(4*i*l-(i-s+l)*(i-s+l))/(2*t)]}return c.invert=function(t,e){var i,c,u=e*e,f=o(L(u+(i=t+r)*i)),h=o(L(u+(i=t+n)*i));return[a(c=f-h,i=(f+h)*s),(e<0?-1:1)*E(L(i*i+c*c)*l)]},c}function Lr(t,e){return Ar(Er,t,e)}function Cr(t,e){if(n(e)<v)return[t,0];var r=n(e/x),i=S(r);if(n(t)<v||n(n(e)-x)<v)return[0,d(e)*y*g(i/2)];var a=o(i),s=n(y/t-t/y)/2,l=s*s,c=a/(r+a-1),u=c*(2/r-1),f=u*u,h=f+l,p=c-f,m=l+c;return[d(t)*y*(s*p+L(l*p*p-h*(c*c-f)))/h,d(e)*y*(u*m-s*L((l+1)*h-m*m))/h]}function Pr(t,e){if(n(e)<v)return[t,0];var r=n(e/x),i=S(r);if(n(t)<v||n(n(e)-x)<v)return[0,d(e)*y*g(i/2)];var a=o(i),s=n(y/t-t/y)/2,l=s*s,c=a*(L(1+l)-s*a)/(1+l*r*r);return[d(t)*y*c,d(e)*y*L(1-c*(2*s+c))]}function Ir(t,e){if(n(e)<v)return[t,0];var r=e/x,i=S(r);if(n(t)<v||n(n(e)-x)<v)return[0,y*g(i/2)];var a=(y/t-t/y)/2,s=r/(1+o(i));return[y*(d(t)*L(a*a+1-s*s)-a),y*s]}function Or(t,e){if(!e)return[t,0];var r=n(e);if(!t||r===x)return[0,e];var i=r/x,a=i*i,o=(8*i-a*(a+2)-5)/(2*a*(i-1)),s=o*o,l=i*o,c=a+s+2*l,u=i+3*o,f=t/x,h=f+1/f,p=d(n(t)-x)*L(h*h-4),m=p*p,g=(p*(c+s-1)+2*L(c*(a+s*m-1)+(1-a)*(a*(u*u+4*s)+12*l*s+4*s*s)))/(4*c+m);return[d(t)*x*g,d(e)*x*L(1+p*n(g)-g*g)]}function zr(t,e,r,n){var i=y/3;t=u(t,v),e=u(e,v),t=f(t,x),e=f(e,y-v),r=u(r,0),r=f(r,100-v);var s=(n=u(n,v))/100,l=E((r/100+1)*o(i))/i,c=m(t)/m(l*x),h=e/y,p=L(s*m(t/2)/m(e/2));return function(t,e,r,n,i){function s(a,s){var l=r*m(n*s),c=L(1-l*l),u=L(2/(1+c*o(a*=i)));return[t*c*u*m(a),e*l*u]}return s.invert=function(o,s){var l=o/t,c=s/e,u=L(l*l+c*c),f=2*S(u/2);return[a(o*g(f),t*u)/i,u&&S(s*m(f)/(e*r*u))/n]},s}(p/L(h*c*l),1/(p*L(h*c*l)),c,l,h)}function Dr(){var t=65*M,r=60*M,n=20,i=200,a=e.geoProjectionMutator(zr),o=a(t,r,n,i);return o.poleline=function(e){return arguments.length?a(t=+e*M,r,n,i):t*A},o.parallels=function(e){return arguments.length?a(t,r=+e*M,n,i):r*A},o.inflation=function(e){return arguments.length?a(t,r,n=+e,i):n},o.ratio=function(e){return arguments.length?a(t,r,n,i=+e):i},o.scale(163.775)}kr.invert=function(t,e){var r=e/1.70711,n=m(b*r);return[t/(.74482-.34588*n*n),2*i(r)]},Cr.invert=function(t,e){if(n(e)<v)return[t,0];if(n(t)<v)return[0,x*m(2*i(e/y))];var r=(t/=y)*t,a=(e/=y)*e,s=r+a,l=s*s,c=-n(e)*(1+s),u=c-2*a+r,f=-2*c+1+2*a+l,h=a/f+(2*u*u*u/(f*f*f)-9*c*u/(f*f))/27,p=(c-u*u/(3*f))/f,g=2*L(-p/3),b=E(3*h/(p*g))/3;return[y*(s-1+L(1+2*(r-a)+l))/(2*t),d(e)*y*(-g*o(b+y/3)-u/(3*f))]},Pr.invert=function(t,e){if(!t)return[0,x*m(2*i(e/y))];var r=n(t/y),o=(1-r*r-(e/=y)*e)/(2*r),s=L(o*o+1);return[d(t)*y*(s-o),d(e)*x*m(2*a(L((1-2*o*r)*(o+s)-r),L(s+o+r)))]},Ir.invert=function(t,e){if(!e)return[t,0];var r=e/y,n=(y*y*(1-r*r)-t*t)/(2*y*t);return[t?y*(d(t)*L(n*n+1)-n):0,x*m(2*i(r))]},Or.invert=function(t,e){var r;if(!t||!e)return[t,e];e/=y;var i=d(t)*t/x,a=(i*i-1+4*e*e)/n(i),o=a*a,s=2*e,l=50;do{var c=s*s,u=(8*s-c*(c+2)-5)/(2*c*(s-1)),f=(3*s-c*s-10)/(2*c*s),h=u*u,p=s*u,m=s+u,g=m*m,b=s+3*u,_=-2*m*(4*p*h+(1-4*c+3*c*c)*(1+f)+h*(14*c-6-o+(8*c-8-2*o)*f)+p*(12*c-8+(10*c-10-o)*f)),w=L(g*(c+h*o-1)+(1-c)*(c*(b*b+4*h)+h*(12*p+4*h)));s-=r=(a*(g+h-1)+2*w-i*(4*g+o))/(a*(2*u*f+2*m*(1+f))+_/w-8*m*(a*(-1+h+g)+2*w)*(1+f)/(o+4*g))}while(r>v&&--l>0);return[d(t)*(L(a*a+4)+a)*y/4,x*s]};var Rr=4*y+3*L(3),Fr=2*L(2*y*L(3)/Rr),Br=Y(Fr*L(3)/y,Fr,Rr/6);function Nr(t,e){return[t*L(1-3*e*e/(y*y)),e]}function jr(t,e){var r=o(e),n=o(t)*r,i=1-n,s=o(t=a(m(t)*r,-m(e))),l=m(t);return[l*(r=L(1-n*n))-s*i,-s*r-l*i]}function Ur(t,e){var r=O(t,e);return[(r[0]+t/x)/2,(r[1]+e)/2]}Nr.invert=function(t,e){return[t/L(1-3*e*e/(y*y)),e]},jr.invert=function(t,e){var r=(t*t+e*e)/-2,n=L(-r*(2+r)),i=e*r+t*n,o=t*r-e*n,s=L(o*o+i*i);return[a(n*i,s*(1+r)),s?-S(n*o/s):0]},Ur.invert=function(t,e){var r=t,i=e,a=25;do{var s,l=o(i),c=m(i),u=m(2*i),f=c*c,h=l*l,p=m(r),d=o(r/2),g=m(r/2),y=g*g,b=1-h*d*d,_=b?E(l*d)*L(s=1/b):s=0,w=.5*(2*_*l*g+r/x)-t,T=.5*(_*c+i)-e,k=.5*s*(h*y+_*l*d*f)+.5/x,A=s*(p*u/4-_*c*g),M=.125*s*(u*g-_*c*h*p),S=.5*s*(f*d+_*y*l)+.5,C=A*M-S*k,P=(T*A-w*S)/C,I=(w*M-T*k)/C;r-=P,i-=I}while((n(P)>v||n(I)>v)&&--a>0);return[r,i]},t.geoNaturalEarth=e.geoNaturalEarth1,t.geoNaturalEarthRaw=e.geoNaturalEarth1Raw,t.geoAiry=function(){var t=x,r=e.geoProjectionMutator(I),n=r(t);return n.radius=function(e){return arguments.length?r(t=e*M):t*A},n.scale(179.976).clipAngle(147)},t.geoAiryRaw=I,t.geoAitoff=function(){return e.geoProjection(O).scale(152.63)},t.geoAitoffRaw=O,t.geoArmadillo=function(){var t=20*M,r=t>=0?1:-1,n=g(r*t),i=e.geoProjectionMutator(z),s=i(t),l=s.stream;return s.parallel=function(e){return arguments.length?(n=g((r=(t=e*M)>=0?1:-1)*t),i(t)):t*A},s.stream=function(e){var i=s.rotate(),c=l(e),u=(s.rotate([0,0]),l(e)),f=s.precision();return s.rotate(i),c.sphere=function(){u.polygonStart(),u.lineStart();for(var e=-180*r;r*e<180;e+=90*r)u.point(e,90*r);if(t)for(;r*(e-=3*r*f)>=-180;)u.point(e,r*-a(o(e*M/2),n)*A);u.lineEnd(),u.polygonEnd()},c},s.scale(218.695).center([0,28.0974])},t.geoArmadilloRaw=z,t.geoAugust=function(){return e.geoProjection(D).scale(66.1603)},t.geoAugustRaw=D,t.geoBaker=function(){return e.geoProjection(B).scale(112.314)},t.geoBakerRaw=B,t.geoBerghaus=function(){var t=5,r=e.geoProjectionMutator(N),n=r(t),i=n.stream,s=-o(.01*M),l=m(.01*M);return n.lobes=function(e){return arguments.length?r(t=+e):t},n.stream=function(e){var r=n.rotate(),c=i(e),u=(n.rotate([0,0]),i(e));return n.rotate(r),c.sphere=function(){u.polygonStart(),u.lineStart();for(var e=0,r=360/t,n=2*y/t,i=90-180/t,c=x;e<t;++e,i-=r,c-=n)u.point(a(l*o(c),s)*A,S(l*m(c))*A),i<-90?(u.point(-90,-180-i-.01),u.point(-90,-180-i+.01)):(u.point(90,i+.01),u.point(90,i-.01));u.lineEnd(),u.polygonEnd()},c},n.scale(87.8076).center([0,17.1875]).clipAngle(179.999)},t.geoBerghausRaw=N,t.geoBertin1953=function(){return e.geoProjection(q()).rotate([-16.5,-42]).scale(176.57).center([7.93,.09])},t.geoBertin1953Raw=q,t.geoBoggs=function(){return e.geoProjection(J).scale(160.857)},t.geoBoggsRaw=J,t.geoBonne=function(){return K($).scale(123.082).center([0,26.1441]).parallel(45)},t.geoBonneRaw=$,t.geoBottomley=function(){var t=.5,r=e.geoProjectionMutator(tt),n=r(t);return n.fraction=function(e){return arguments.length?r(t=+e):t},n.scale(158.837)},t.geoBottomleyRaw=tt,t.geoBromley=function(){return e.geoProjection(et).scale(152.63)},t.geoBromleyRaw=et,t.geoChamberlin=st,t.geoChamberlinRaw=at,t.geoChamberlinAfrica=function(){return st([0,22],[45,22],[22.5,-22]).scale(380).center([22.5,2])},t.geoCollignon=function(){return e.geoProjection(lt).scale(95.6464).center([0,30])},t.geoCollignonRaw=lt,t.geoCraig=function(){return K(ct).scale(249.828).clipAngle(90)},t.geoCraigRaw=ct,t.geoCraster=function(){return e.geoProjection(ft).scale(156.19)},t.geoCrasterRaw=ft,t.geoCylindricalEqualArea=function(){return K(ht).parallel(38.58).scale(195.044)},t.geoCylindricalEqualAreaRaw=ht,t.geoCylindricalStereographic=function(){return K(pt).scale(124.75)},t.geoCylindricalStereographicRaw=pt,t.geoEckert1=function(){return e.geoProjection(dt).scale(165.664)},t.geoEckert1Raw=dt,t.geoEckert2=function(){return e.geoProjection(mt).scale(165.664)},t.geoEckert2Raw=mt,t.geoEckert3=function(){return e.geoProjection(gt).scale(180.739)},t.geoEckert3Raw=gt,t.geoEckert4=function(){return e.geoProjection(vt).scale(180.739)},t.geoEckert4Raw=vt,t.geoEckert5=function(){return e.geoProjection(yt).scale(173.044)},t.geoEckert5Raw=yt,t.geoEckert6=function(){return e.geoProjection(xt).scale(173.044)},t.geoEckert6Raw=xt,t.geoEisenlohr=function(){return e.geoProjection(_t).scale(62.5271)},t.geoEisenlohrRaw=_t,t.geoFahey=function(){return e.geoProjection(Tt).scale(137.152)},t.geoFaheyRaw=Tt,t.geoFoucaut=function(){return e.geoProjection(kt).scale(135.264)},t.geoFoucautRaw=kt,t.geoFoucautSinusoidal=function(){var t=.5,r=e.geoProjectionMutator(At),n=r(t);return n.alpha=function(e){return arguments.length?r(t=+e):t},n.scale(168.725)},t.geoFoucautSinusoidalRaw=At,t.geoGilbert=function(t){null==t&&(t=e.geoOrthographic);var r=t(),n=e.geoEquirectangular().scale(A).precision(0).clipAngle(null).translate([0,0]);function i(t){return r(Mt(t))}function a(t){i[t]=function(){return arguments.length?(r[t].apply(r,arguments),i):r[t]()}}return r.invert&&(i.invert=function(t){return St(r.invert(t))}),i.stream=function(t){var e=r.stream(t),i=n.stream({point:function(t,r){e.point(t/2,S(g(-r/2*M))*A)},lineStart:function(){e.lineStart()},lineEnd:function(){e.lineEnd()},polygonStart:function(){e.polygonStart()},polygonEnd:function(){e.polygonEnd()}});return i.sphere=e.sphere,i},i.rotate=function(t){return arguments.length?(n.rotate(t),i):n.rotate()},i.center=function(t){return arguments.length?(r.center(Mt(t)),i):St(r.center())},a("angle"),a("clipAngle"),a("clipExtent"),a("fitExtent"),a("fitHeight"),a("fitSize"),a("fitWidth"),a("scale"),a("translate"),a("precision"),i.scale(249.5)},t.geoGingery=function(){var t=6,r=30*M,n=o(r),i=m(r),s=e.geoProjectionMutator(Et),l=s(r,t),c=l.stream,u=-o(.01*M),f=m(.01*M);return l.radius=function(e){return arguments.length?(n=o(r=e*M),i=m(r),s(r,t)):r*A},l.lobes=function(e){return arguments.length?s(r,t=+e):t},l.stream=function(e){var r=l.rotate(),s=c(e),h=(l.rotate([0,0]),c(e));return l.rotate(r),s.sphere=function(){h.polygonStart(),h.lineStart();for(var e=0,r=2*y/t,s=0;e<t;++e,s-=r)h.point(a(f*o(s),u)*A,S(f*m(s))*A),h.point(a(i*o(s-r/2),n)*A,S(i*m(s-r/2))*A);h.lineEnd(),h.polygonEnd()},s},l.rotate([90,-40]).scale(91.7095).clipAngle(179.999)},t.geoGingeryRaw=Et,t.geoGinzburg4=function(){return e.geoProjection(It).scale(149.995)},t.geoGinzburg4Raw=It,t.geoGinzburg5=function(){return e.geoProjection(Ot).scale(153.93)},t.geoGinzburg5Raw=Ot,t.geoGinzburg6=function(){return e.geoProjection(zt).scale(130.945)},t.geoGinzburg6Raw=zt,t.geoGinzburg8=function(){return e.geoProjection(Dt).scale(131.747)},t.geoGinzburg8Raw=Dt,t.geoGinzburg9=function(){return e.geoProjection(Rt).scale(131.087)},t.geoGinzburg9Raw=Rt,t.geoGringorten=function(){return e.geoProjection(Ft(Bt)).scale(239.75)},t.geoGringortenRaw=Bt,t.geoGuyou=function(){return e.geoProjection(Ft(Ut)).scale(151.496)},t.geoGuyouRaw=Ut,t.geoHammer=function(){var t=2,r=e.geoProjectionMutator(j),n=r(t);return n.coefficient=function(e){return arguments.length?r(t=+e):t},n.scale(169.529)},t.geoHammerRaw=j,t.geoHammerRetroazimuthal=function(){var t=0,r=e.geoProjectionMutator(Vt),n=r(t),i=n.rotate,a=n.stream,o=e.geoCircle();return n.parallel=function(e){if(!arguments.length)return t*A;var i=n.rotate();return r(t=e*M).rotate(i)},n.rotate=function(e){return arguments.length?(i.call(n,[e[0],e[1]-t*A]),o.center([-e[0],-e[1]]),n):((e=i.call(n))[1]+=t*A,e)},n.stream=function(t){return(t=a(t)).sphere=function(){t.polygonStart();var e,r=o.radius(89.99)().coordinates[0],n=r.length-1,i=-1;for(t.lineStart();++i<n;)t.point((e=r[i])[0],e[1]);for(t.lineEnd(),n=(r=o.radius(90.01)().coordinates[0]).length-1,t.lineStart();--i>=0;)t.point((e=r[i])[0],e[1]);t.lineEnd(),t.polygonEnd()},t},n.scale(79.4187).parallel(45).clipAngle(179.999)},t.geoHammerRetroazimuthalRaw=Vt,t.geoHealpix=function(){var t=4,n=e.geoProjectionMutator(Yt),i=n(t),a=i.stream;return i.lobes=function(e){return arguments.length?n(t=+e):t},i.stream=function(n){var o=i.rotate(),s=a(n),l=(i.rotate([0,0]),a(n));return i.rotate(o),s.sphere=function(){var n,i;e.geoStream((n=180/t,i=[].concat(r.range(-180,180+n/2,n).map(Wt),r.range(180,-180-n/2,-n).map(Xt)),{type:"Polygon",coordinates:[180===n?i.map(Zt):i]}),l)},s},i.scale(239.75)},t.geoHealpixRaw=Yt,t.geoHill=function(){var t=1,r=e.geoProjectionMutator(Jt),n=r(t);return n.ratio=function(e){return arguments.length?r(t=+e):t},n.scale(167.774).center([0,18.67])},t.geoHillRaw=Jt,t.geoHomolosine=function(){return e.geoProjection(Qt).scale(152.63)},t.geoHomolosineRaw=Qt,t.geoHufnagel=function(){var t=1,r=0,n=45*M,i=2,a=e.geoProjectionMutator($t),o=a(t,r,n,i);return o.a=function(e){return arguments.length?a(t=+e,r,n,i):t},o.b=function(e){return arguments.length?a(t,r=+e,n,i):r},o.psiMax=function(e){return arguments.length?a(t,r,n=+e*M,i):n*A},o.ratio=function(e){return arguments.length?a(t,r,n,i=+e):i},o.scale(180.739)},t.geoHufnagelRaw=$t,t.geoHyperelliptical=function(){var t=0,r=2.5,n=1.183136,i=e.geoProjectionMutator(ee),a=i(t,r,n);return a.alpha=function(e){return arguments.length?i(t=+e,r,n):t},a.k=function(e){return arguments.length?i(t,r=+e,n):r},a.gamma=function(e){return arguments.length?i(t,r,n=+e):n},a.scale(152.63)},t.geoHyperellipticalRaw=ee,t.geoInterrupt=ae,t.geoInterruptedBoggs=function(){return ae(J,oe).scale(160.857)},t.geoInterruptedHomolosine=function(){return ae(Qt,se).scale(152.63)},t.geoInterruptedMollweide=function(){return ae(W,le).scale(169.529)},t.geoInterruptedMollweideHemispheres=function(){return ae(W,ce).scale(169.529).rotate([20,0])},t.geoInterruptedSinuMollweide=function(){return ae(Kt,ue,H).rotate([-20,-55]).scale(164.263).center([0,-5.4036])},t.geoInterruptedSinusoidal=function(){return ae(Q,fe).scale(152.63).rotate([-20,0])},t.geoKavrayskiy7=function(){return e.geoProjection(he).scale(158.837)},t.geoKavrayskiy7Raw=he,t.geoLagrange=function(){var t=.5,r=e.geoProjectionMutator(pe),n=r(t);return n.spacing=function(e){return arguments.length?r(t=+e):t},n.scale(124.75)},t.geoLagrangeRaw=pe,t.geoLarrivee=function(){return e.geoProjection(me).scale(97.2672)},t.geoLarriveeRaw=me,t.geoLaskowski=function(){return e.geoProjection(ge).scale(139.98)},t.geoLaskowskiRaw=ge,t.geoLittrow=function(){return e.geoProjection(ve).scale(144.049).clipAngle(89.999)},t.geoLittrowRaw=ve,t.geoLoximuthal=function(){return K(ye).parallel(40).scale(158.837)},t.geoLoximuthalRaw=ye,t.geoMiller=function(){return e.geoProjection(xe).scale(108.318)},t.geoMillerRaw=xe,t.geoModifiedStereographic=Me,t.geoModifiedStereographicRaw=be,t.geoModifiedStereographicAlaska=function(){return Me(_e,[152,-64]).scale(1400).center([-160.908,62.4864]).clipAngle(30).angle(7.8)},t.geoModifiedStereographicGs48=function(){return Me(we,[95,-38]).scale(1e3).clipAngle(55).center([-96.5563,38.8675])},t.geoModifiedStereographicGs50=function(){return Me(Te,[120,-45]).scale(359.513).clipAngle(55).center([-117.474,53.0628])},t.geoModifiedStereographicMiller=function(){return Me(ke,[-20,-18]).scale(209.091).center([20,16.7214]).clipAngle(82)},t.geoModifiedStereographicLee=function(){return Me(Ae,[165,10]).scale(250).clipAngle(130).center([-165,-10])},t.geoMollweide=function(){return e.geoProjection(W).scale(169.529)},t.geoMollweideRaw=W,t.geoMtFlatPolarParabolic=function(){return e.geoProjection(Le).scale(164.859)},t.geoMtFlatPolarParabolicRaw=Le,t.geoMtFlatPolarQuartic=function(){return e.geoProjection(Ce).scale(188.209)},t.geoMtFlatPolarQuarticRaw=Ce,t.geoMtFlatPolarSinusoidal=function(){return e.geoProjection(Pe).scale(166.518)},t.geoMtFlatPolarSinusoidalRaw=Pe,t.geoNaturalEarth2=function(){return e.geoProjection(Ie).scale(175.295)},t.geoNaturalEarth2Raw=Ie,t.geoNellHammer=function(){return e.geoProjection(Oe).scale(152.63)},t.geoNellHammerRaw=Oe,t.geoInterruptedQuarticAuthalic=function(){return ae(j(1/0),ze).rotate([20,0]).scale(152.63)},t.geoNicolosi=function(){return e.geoProjection(De).scale(127.267)},t.geoNicolosiRaw=De,t.geoPatterson=function(){return e.geoProjection(Re).scale(139.319)},t.geoPattersonRaw=Re,t.geoPolyconic=function(){return e.geoProjection(Fe).scale(103.74)},t.geoPolyconicRaw=Fe,t.geoPolyhedral=Ve,t.geoPolyhedralButterfly=function(t){t=t||function(t){var r=e.geoCentroid({type:"MultiPoint",coordinates:t});return e.geoGnomonic().scale(1).translate([0,0]).rotate([-r[0],-r[1]])};var r=Ye.map((function(e){return{face:e,project:t(e)}}));return[-1,0,0,1,0,1,4,5].forEach((function(t,e){var n=r[t];n&&(n.children||(n.children=[])).push(r[e])})),Ve(r[0],(function(t,e){return r[t<-y/2?e<0?6:4:t<0?e<0?2:0:t<y/2?e<0?3:1:e<0?7:5]})).angle(-30).scale(101.858).center([0,45])},t.geoPolyhedralCollignon=function(t){t=t||function(t){var r=e.geoCentroid({type:"MultiPoint",coordinates:t});return e.geoProjection(Xe).translate([0,0]).scale(1).rotate(r[1]>0?[-r[0],0]:[180-r[0],180])};var r=Ye.map((function(e){return{face:e,project:t(e)}}));return[-1,0,0,1,0,1,4,5].forEach((function(t,e){var n=r[t];n&&(n.children||(n.children=[])).push(r[e])})),Ve(r[0],(function(t,e){return r[t<-y/2?e<0?6:4:t<0?e<0?2:0:t<y/2?e<0?3:1:e<0?7:5]})).angle(-30).scale(121.906).center([0,48.5904])},t.geoPolyhedralWaterman=function(t){t=t||function(t){var r=6===t.length?e.geoCentroid({type:"MultiPoint",coordinates:t}):t[0];return e.geoGnomonic().scale(1).translate([0,0]).rotate([-r[0],-r[1]])};var r=Ye.map((function(t){for(var e,r=t.map(Ke),n=r.length,i=r[n-1],a=[],o=0;o<n;++o)e=r[o],a.push(Je([.9486832980505138*i[0]+.31622776601683794*e[0],.9486832980505138*i[1]+.31622776601683794*e[1],.9486832980505138*i[2]+.31622776601683794*e[2]]),Je([.9486832980505138*e[0]+.31622776601683794*i[0],.9486832980505138*e[1]+.31622776601683794*i[1],.9486832980505138*e[2]+.31622776601683794*i[2]])),i=e;return a})),n=[],i=[-1,0,0,1,0,1,4,5];r.forEach((function(t,e){for(var a,o,s=Ye[e],l=s.length,c=n[e]=[],u=0;u<l;++u)r.push([s[u],t[(2*u+2)%(2*l)],t[(2*u+1)%(2*l)]]),i.push(e),c.push((a=Ke(t[(2*u+2)%(2*l)]),o=Ke(t[(2*u+1)%(2*l)]),[a[1]*o[2]-a[2]*o[1],a[2]*o[0]-a[0]*o[2],a[0]*o[1]-a[1]*o[0]]))}));var a=r.map((function(e){return{project:t(e),face:e}}));return i.forEach((function(t,e){var r=a[t];r&&(r.children||(r.children=[])).push(a[e])})),Ve(a[0],(function(t,e){var r=o(e),i=[r*o(t),r*m(t),m(e)],s=t<-y/2?e<0?6:4:t<0?e<0?2:0:t<y/2?e<0?3:1:e<0?7:5,l=n[s];return a[Ze(l[0],i)<0?8+3*s:Ze(l[1],i)<0?8+3*s+1:Ze(l[2],i)<0?8+3*s+2:s]})).angle(-30).scale(110.625).center([0,45])},t.geoProject=function(t,e){var r,n=e.stream;if(!n)throw new Error("invalid projection");switch(t&&t.type){case"Feature":r=tr;break;case"FeatureCollection":r=$e;break;default:r=er}return r(t,n)},t.geoGringortenQuincuncial=function(){return sr(Bt).scale(176.423)},t.geoPeirceQuincuncial=lr,t.geoPierceQuincuncial=lr,t.geoQuantize=function(t,e){if(!(0<=(e=+e)&&e<=20))throw new Error("invalid digits");function r(t){var r=t.length,n=2,i=new Array(r);for(i[0]=+t[0].toFixed(e),i[1]=+t[1].toFixed(e);n<r;)i[n]=t[n],++n;return i}function n(t){return t.map(r)}function i(t){for(var e=r(t[0]),n=[e],i=1;i<t.length;i++){var a=r(t[i]);(a.length>2||a[0]!=e[0]||a[1]!=e[1])&&(n.push(a),e=a)}return 1===n.length&&t.length>1&&n.push(r(t[t.length-1])),n}function a(t){return t.map(i)}function o(t){if(null==t)return t;var e;switch(t.type){case"GeometryCollection":e={type:"GeometryCollection",geometries:t.geometries.map(o)};break;case"Point":e={type:"Point",coordinates:r(t.coordinates)};break;case"MultiPoint":e={type:t.type,coordinates:n(t.coordinates)};break;case"LineString":e={type:t.type,coordinates:i(t.coordinates)};break;case"MultiLineString":case"Polygon":e={type:t.type,coordinates:a(t.coordinates)};break;case"MultiPolygon":e={type:"MultiPolygon",coordinates:t.coordinates.map(a)};break;default:return t}return null!=t.bbox&&(e.bbox=t.bbox),e}function s(t){var e={type:"Feature",properties:t.properties,geometry:o(t.geometry)};return null!=t.id&&(e.id=t.id),null!=t.bbox&&(e.bbox=t.bbox),e}if(null!=t)switch(t.type){case"Feature":return s(t);case"FeatureCollection":var l={type:"FeatureCollection",features:t.features.map(s)};return null!=t.bbox&&(l.bbox=t.bbox),l;default:return o(t)}return t},t.geoQuincuncial=sr,t.geoRectangularPolyconic=function(){return K(cr).scale(131.215)},t.geoRectangularPolyconicRaw=cr,t.geoRobinson=function(){return e.geoProjection(fr).scale(152.63)},t.geoRobinsonRaw=fr,t.geoSatellite=function(){var t=2,r=0,n=e.geoProjectionMutator(hr),i=n(t,r);return i.distance=function(e){return arguments.length?n(t=+e,r):t},i.tilt=function(e){return arguments.length?n(t,r=e*M):r*A},i.scale(432.147).clipAngle(E(1/t)*A-1e-6)},t.geoSatelliteRaw=hr,t.geoSinuMollweide=function(){return e.geoProjection(Kt).rotate([-20,-55]).scale(164.263).center([0,-5.4036])},t.geoSinuMollweideRaw=Kt,t.geoSinusoidal=function(){return e.geoProjection(Q).scale(152.63)},t.geoSinusoidalRaw=Q,t.geoStitch=function(t){if(null==t)return t;switch(t.type){case"Feature":return wr(t);case"FeatureCollection":var e={type:"FeatureCollection",features:t.features.map(wr)};return null!=t.bbox&&(e.bbox=t.bbox),e;default:return Tr(t)}},t.geoTimes=function(){return e.geoProjection(kr).scale(146.153)},t.geoTimesRaw=kr,t.geoTwoPointAzimuthal=Sr,t.geoTwoPointAzimuthalRaw=Mr,t.geoTwoPointAzimuthalUsa=function(){return Sr([-158,21.5],[-77,39]).clipAngle(60).scale(400)},t.geoTwoPointEquidistant=Lr,t.geoTwoPointEquidistantRaw=Er,t.geoTwoPointEquidistantUsa=function(){return Lr([-158,21.5],[-77,39]).clipAngle(130).scale(122.571)},t.geoVanDerGrinten=function(){return e.geoProjection(Cr).scale(79.4183)},t.geoVanDerGrintenRaw=Cr,t.geoVanDerGrinten2=function(){return e.geoProjection(Pr).scale(79.4183)},t.geoVanDerGrinten2Raw=Pr,t.geoVanDerGrinten3=function(){return e.geoProjection(Ir).scale(79.4183)},t.geoVanDerGrinten3Raw=Ir,t.geoVanDerGrinten4=function(){return e.geoProjection(Or).scale(127.16)},t.geoVanDerGrinten4Raw=Or,t.geoWagner=Dr,t.geoWagner7=function(){return Dr().poleline(65).parallels(60).inflation(0).ratio(200).scale(172.633)},t.geoWagnerRaw=zr,t.geoWagner4=function(){return e.geoProjection(Br).scale(176.84)},t.geoWagner4Raw=Br,t.geoWagner6=function(){return e.geoProjection(Nr).scale(152.63)},t.geoWagner6Raw=Nr,t.geoWiechel=function(){return e.geoProjection(jr).rotate([0,-90,45]).scale(124.75).clipAngle(179.999)},t.geoWiechelRaw=jr,t.geoWinkel3=function(){return e.geoProjection(Ur).scale(158.837)},t.geoWinkel3Raw=Ur,Object.defineProperty(t,"__esModule",{value:!0})}))},{"d3-array":102,"d3-geo":109}],109:[function(t,e,r){!function(n,i){"object"==typeof r&&void 0!==e?i(r,t("d3-array")):i((n=n||self).d3=n.d3||{},n.d3)}(this,(function(t,e){"use strict";function r(){return new n}function n(){this.reset()}n.prototype={constructor:n,reset:function(){this.s=this.t=0},add:function(t){a(i,t,this.t),a(this,i.s,this.s),this.s?this.t+=i.t:this.s=i.t},valueOf:function(){return this.s}};var i=new n;function a(t,e,r){var n=t.s=e+r,i=n-e,a=n-i;t.t=e-a+(r-i)}var o=1e-6,s=Math.PI,l=s/2,c=s/4,u=2*s,f=180/s,h=s/180,p=Math.abs,d=Math.atan,m=Math.atan2,g=Math.cos,v=Math.ceil,y=Math.exp,x=Math.log,b=Math.pow,_=Math.sin,w=Math.sign||function(t){return t>0?1:t<0?-1:0},T=Math.sqrt,k=Math.tan;function A(t){return t>1?0:t<-1?s:Math.acos(t)}function M(t){return t>1?l:t<-1?-l:Math.asin(t)}function S(t){return(t=_(t/2))*t}function E(){}function L(t,e){t&&P.hasOwnProperty(t.type)&&P[t.type](t,e)}var C={Feature:function(t,e){L(t.geometry,e)},FeatureCollection:function(t,e){for(var r=t.features,n=-1,i=r.length;++n<i;)L(r[n].geometry,e)}},P={Sphere:function(t,e){e.sphere()},Point:function(t,e){t=t.coordinates,e.point(t[0],t[1],t[2])},MultiPoint:function(t,e){for(var r=t.coordinates,n=-1,i=r.length;++n<i;)t=r[n],e.point(t[0],t[1],t[2])},LineString:function(t,e){I(t.coordinates,e,0)},MultiLineString:function(t,e){for(var r=t.coordinates,n=-1,i=r.length;++n<i;)I(r[n],e,0)},Polygon:function(t,e){O(t.coordinates,e)},MultiPolygon:function(t,e){for(var r=t.coordinates,n=-1,i=r.length;++n<i;)O(r[n],e)},GeometryCollection:function(t,e){for(var r=t.geometries,n=-1,i=r.length;++n<i;)L(r[n],e)}};function I(t,e,r){var n,i=-1,a=t.length-r;for(e.lineStart();++i<a;)n=t[i],e.point(n[0],n[1],n[2]);e.lineEnd()}function O(t,e){var r=-1,n=t.length;for(e.polygonStart();++r<n;)I(t[r],e,1);e.polygonEnd()}function z(t,e){t&&C.hasOwnProperty(t.type)?C[t.type](t,e):L(t,e)}var D,R,F,B,N,j=r(),U=r(),V={point:E,lineStart:E,lineEnd:E,polygonStart:function(){j.reset(),V.lineStart=H,V.lineEnd=q},polygonEnd:function(){var t=+j;U.add(t<0?u+t:t),this.lineStart=this.lineEnd=this.point=E},sphere:function(){U.add(u)}};function H(){V.point=G}function q(){Y(D,R)}function G(t,e){V.point=Y,D=t,R=e,F=t*=h,B=g(e=(e*=h)/2+c),N=_(e)}function Y(t,e){var r=(t*=h)-F,n=r>=0?1:-1,i=n*r,a=g(e=(e*=h)/2+c),o=_(e),s=N*o,l=B*a+s*g(i),u=s*n*_(i);j.add(m(u,l)),F=t,B=a,N=o}function W(t){return[m(t[1],t[0]),M(t[2])]}function X(t){var e=t[0],r=t[1],n=g(r);return[n*g(e),n*_(e),_(r)]}function Z(t,e){return t[0]*e[0]+t[1]*e[1]+t[2]*e[2]}function J(t,e){return[t[1]*e[2]-t[2]*e[1],t[2]*e[0]-t[0]*e[2],t[0]*e[1]-t[1]*e[0]]}function K(t,e){t[0]+=e[0],t[1]+=e[1],t[2]+=e[2]}function Q(t,e){return[t[0]*e,t[1]*e,t[2]*e]}function $(t){var e=T(t[0]*t[0]+t[1]*t[1]+t[2]*t[2]);t[0]/=e,t[1]/=e,t[2]/=e}var tt,et,rt,nt,it,at,ot,st,lt,ct,ut,ft,ht,pt,dt,mt,gt,vt,yt,xt,bt,_t,wt,Tt,kt,At,Mt=r(),St={point:Et,lineStart:Ct,lineEnd:Pt,polygonStart:function(){St.point=It,St.lineStart=Ot,St.lineEnd=zt,Mt.reset(),V.polygonStart()},polygonEnd:function(){V.polygonEnd(),St.point=Et,St.lineStart=Ct,St.lineEnd=Pt,j<0?(tt=-(rt=180),et=-(nt=90)):Mt>o?nt=90:Mt<-o&&(et=-90),ct[0]=tt,ct[1]=rt},sphere:function(){tt=-(rt=180),et=-(nt=90)}};function Et(t,e){lt.push(ct=[tt=t,rt=t]),e<et&&(et=e),e>nt&&(nt=e)}function Lt(t,e){var r=X([t*h,e*h]);if(st){var n=J(st,r),i=J([n[1],-n[0],0],n);$(i),i=W(i);var a,o=t-it,s=o>0?1:-1,l=i[0]*f*s,c=p(o)>180;c^(s*it<l&&l<s*t)?(a=i[1]*f)>nt&&(nt=a):c^(s*it<(l=(l+360)%360-180)&&l<s*t)?(a=-i[1]*f)<et&&(et=a):(e<et&&(et=e),e>nt&&(nt=e)),c?t<it?Dt(tt,t)>Dt(tt,rt)&&(rt=t):Dt(t,rt)>Dt(tt,rt)&&(tt=t):rt>=tt?(t<tt&&(tt=t),t>rt&&(rt=t)):t>it?Dt(tt,t)>Dt(tt,rt)&&(rt=t):Dt(t,rt)>Dt(tt,rt)&&(tt=t)}else lt.push(ct=[tt=t,rt=t]);e<et&&(et=e),e>nt&&(nt=e),st=r,it=t}function Ct(){St.point=Lt}function Pt(){ct[0]=tt,ct[1]=rt,St.point=Et,st=null}function It(t,e){if(st){var r=t-it;Mt.add(p(r)>180?r+(r>0?360:-360):r)}else at=t,ot=e;V.point(t,e),Lt(t,e)}function Ot(){V.lineStart()}function zt(){It(at,ot),V.lineEnd(),p(Mt)>o&&(tt=-(rt=180)),ct[0]=tt,ct[1]=rt,st=null}function Dt(t,e){return(e-=t)<0?e+360:e}function Rt(t,e){return t[0]-e[0]}function Ft(t,e){return t[0]<=t[1]?t[0]<=e&&e<=t[1]:e<t[0]||t[1]<e}var Bt={sphere:E,point:Nt,lineStart:Ut,lineEnd:qt,polygonStart:function(){Bt.lineStart=Gt,Bt.lineEnd=Yt},polygonEnd:function(){Bt.lineStart=Ut,Bt.lineEnd=qt}};function Nt(t,e){t*=h;var r=g(e*=h);jt(r*g(t),r*_(t),_(e))}function jt(t,e,r){++ut,ht+=(t-ht)/ut,pt+=(e-pt)/ut,dt+=(r-dt)/ut}function Ut(){Bt.point=Vt}function Vt(t,e){t*=h;var r=g(e*=h);Tt=r*g(t),kt=r*_(t),At=_(e),Bt.point=Ht,jt(Tt,kt,At)}function Ht(t,e){t*=h;var r=g(e*=h),n=r*g(t),i=r*_(t),a=_(e),o=m(T((o=kt*a-At*i)*o+(o=At*n-Tt*a)*o+(o=Tt*i-kt*n)*o),Tt*n+kt*i+At*a);ft+=o,mt+=o*(Tt+(Tt=n)),gt+=o*(kt+(kt=i)),vt+=o*(At+(At=a)),jt(Tt,kt,At)}function qt(){Bt.point=Nt}function Gt(){Bt.point=Wt}function Yt(){Xt(_t,wt),Bt.point=Nt}function Wt(t,e){_t=t,wt=e,t*=h,e*=h,Bt.point=Xt;var r=g(e);Tt=r*g(t),kt=r*_(t),At=_(e),jt(Tt,kt,At)}function Xt(t,e){t*=h;var r=g(e*=h),n=r*g(t),i=r*_(t),a=_(e),o=kt*a-At*i,s=At*n-Tt*a,l=Tt*i-kt*n,c=T(o*o+s*s+l*l),u=M(c),f=c&&-u/c;yt+=f*o,xt+=f*s,bt+=f*l,ft+=u,mt+=u*(Tt+(Tt=n)),gt+=u*(kt+(kt=i)),vt+=u*(At+(At=a)),jt(Tt,kt,At)}function Zt(t){return function(){return t}}function Jt(t,e){function r(r,n){return r=t(r,n),e(r[0],r[1])}return t.invert&&e.invert&&(r.invert=function(r,n){return(r=e.invert(r,n))&&t.invert(r[0],r[1])}),r}function Kt(t,e){return[p(t)>s?t+Math.round(-t/u)*u:t,e]}function Qt(t,e,r){return(t%=u)?e||r?Jt(te(t),ee(e,r)):te(t):e||r?ee(e,r):Kt}function $t(t){return function(e,r){return[(e+=t)>s?e-u:e<-s?e+u:e,r]}}function te(t){var e=$t(t);return e.invert=$t(-t),e}function ee(t,e){var r=g(t),n=_(t),i=g(e),a=_(e);function o(t,e){var o=g(e),s=g(t)*o,l=_(t)*o,c=_(e),u=c*r+s*n;return[m(l*i-u*a,s*r-c*n),M(u*i+l*a)]}return o.invert=function(t,e){var o=g(e),s=g(t)*o,l=_(t)*o,c=_(e),u=c*i-l*a;return[m(l*i+c*a,s*r+u*n),M(u*r-s*n)]},o}function re(t){function e(e){return(e=t(e[0]*h,e[1]*h))[0]*=f,e[1]*=f,e}return t=Qt(t[0]*h,t[1]*h,t.length>2?t[2]*h:0),e.invert=function(e){return(e=t.invert(e[0]*h,e[1]*h))[0]*=f,e[1]*=f,e},e}function ne(t,e,r,n,i,a){if(r){var o=g(e),s=_(e),l=n*r;null==i?(i=e+n*u,a=e-l/2):(i=ie(o,i),a=ie(o,a),(n>0?i<a:i>a)&&(i+=n*u));for(var c,f=i;n>0?f>a:f<a;f-=l)c=W([o,-s*g(f),-s*_(f)]),t.point(c[0],c[1])}}function ie(t,e){(e=X(e))[0]-=t,$(e);var r=A(-e[1]);return((-e[2]<0?-r:r)+u-o)%u}function ae(){var t,e=[];return{point:function(e,r,n){t.push([e,r,n])},lineStart:function(){e.push(t=[])},lineEnd:E,rejoin:function(){e.length>1&&e.push(e.pop().concat(e.shift()))},result:function(){var r=e;return e=[],t=null,r}}}function oe(t,e){return p(t[0]-e[0])<o&&p(t[1]-e[1])<o}function se(t,e,r,n){this.x=t,this.z=e,this.o=r,this.e=n,this.v=!1,this.n=this.p=null}function le(t,e,r,n,i){var a,s,l=[],c=[];if(t.forEach((function(t){if(!((e=t.length-1)<=0)){var e,r,n=t[0],s=t[e];if(oe(n,s)){if(!n[2]&&!s[2]){for(i.lineStart(),a=0;a<e;++a)i.point((n=t[a])[0],n[1]);return void i.lineEnd()}s[0]+=2*o}l.push(r=new se(n,t,null,!0)),c.push(r.o=new se(n,null,r,!1)),l.push(r=new se(s,t,null,!1)),c.push(r.o=new se(s,null,r,!0))}})),l.length){for(c.sort(e),ce(l),ce(c),a=0,s=c.length;a<s;++a)c[a].e=r=!r;for(var u,f,h=l[0];;){for(var p=h,d=!0;p.v;)if((p=p.n)===h)return;u=p.z,i.lineStart();do{if(p.v=p.o.v=!0,p.e){if(d)for(a=0,s=u.length;a<s;++a)i.point((f=u[a])[0],f[1]);else n(p.x,p.n.x,1,i);p=p.n}else{if(d)for(u=p.p.z,a=u.length-1;a>=0;--a)i.point((f=u[a])[0],f[1]);else n(p.x,p.p.x,-1,i);p=p.p}u=(p=p.o).z,d=!d}while(!p.v);i.lineEnd()}}}function ce(t){if(e=t.length){for(var e,r,n=0,i=t[0];++n<e;)i.n=r=t[n],r.p=i,i=r;i.n=r=t[0],r.p=i}}Kt.invert=Kt;var ue=r();function fe(t){return p(t[0])<=s?t[0]:w(t[0])*((p(t[0])+s)%u-s)}function he(t,e){var r=fe(e),n=e[1],i=_(n),a=[_(r),-g(r),0],f=0,h=0;ue.reset(),1===i?n=l+o:-1===i&&(n=-l-o);for(var p=0,d=t.length;p<d;++p)if(y=(v=t[p]).length)for(var v,y,x=v[y-1],b=fe(x),w=x[1]/2+c,T=_(w),k=g(w),A=0;A<y;++A,b=E,T=C,k=P,x=S){var S=v[A],E=fe(S),L=S[1]/2+c,C=_(L),P=g(L),I=E-b,O=I>=0?1:-1,z=O*I,D=z>s,R=T*C;if(ue.add(m(R*O*_(z),k*P+R*g(z))),f+=D?I+O*u:I,D^b>=r^E>=r){var F=J(X(x),X(S));$(F);var B=J(a,F);$(B);var N=(D^I>=0?-1:1)*M(B[2]);(n>N||n===N&&(F[0]||F[1]))&&(h+=D^I>=0?1:-1)}}return(f<-o||f<o&&ue<-o)^1&h}function pe(t,r,n,i){return function(a){var o,s,l,c=r(a),u=ae(),f=r(u),h=!1,p={point:d,lineStart:g,lineEnd:v,polygonStart:function(){p.point=y,p.lineStart=x,p.lineEnd=b,s=[],o=[]},polygonEnd:function(){p.point=d,p.lineStart=g,p.lineEnd=v,s=e.merge(s);var t=he(o,i);s.length?(h||(a.polygonStart(),h=!0),le(s,me,t,n,a)):t&&(h||(a.polygonStart(),h=!0),a.lineStart(),n(null,null,1,a),a.lineEnd()),h&&(a.polygonEnd(),h=!1),s=o=null},sphere:function(){a.polygonStart(),a.lineStart(),n(null,null,1,a),a.lineEnd(),a.polygonEnd()}};function d(e,r){t(e,r)&&a.point(e,r)}function m(t,e){c.point(t,e)}function g(){p.point=m,c.lineStart()}function v(){p.point=d,c.lineEnd()}function y(t,e){l.push([t,e]),f.point(t,e)}function x(){f.lineStart(),l=[]}function b(){y(l[0][0],l[0][1]),f.lineEnd();var t,e,r,n,i=f.clean(),c=u.result(),p=c.length;if(l.pop(),o.push(l),l=null,p)if(1&i){if((e=(r=c[0]).length-1)>0){for(h||(a.polygonStart(),h=!0),a.lineStart(),t=0;t<e;++t)a.point((n=r[t])[0],n[1]);a.lineEnd()}}else p>1&&2&i&&c.push(c.pop().concat(c.shift())),s.push(c.filter(de))}return p}}function de(t){return t.length>1}function me(t,e){return((t=t.x)[0]<0?t[1]-l-o:l-t[1])-((e=e.x)[0]<0?e[1]-l-o:l-e[1])}var ge=pe((function(){return!0}),(function(t){var e,r=NaN,n=NaN,i=NaN;return{lineStart:function(){t.lineStart(),e=1},point:function(a,c){var u=a>0?s:-s,f=p(a-r);p(f-s)<o?(t.point(r,n=(n+c)/2>0?l:-l),t.point(i,n),t.lineEnd(),t.lineStart(),t.point(u,n),t.point(a,n),e=0):i!==u&&f>=s&&(p(r-i)<o&&(r-=i*o),p(a-u)<o&&(a-=u*o),n=function(t,e,r,n){var i,a,s=_(t-r);return p(s)>o?d((_(e)*(a=g(n))*_(r)-_(n)*(i=g(e))*_(t))/(i*a*s)):(e+n)/2}(r,n,a,c),t.point(i,n),t.lineEnd(),t.lineStart(),t.point(u,n),e=0),t.point(r=a,n=c),i=u},lineEnd:function(){t.lineEnd(),r=n=NaN},clean:function(){return 2-e}}}),(function(t,e,r,n){var i;if(null==t)i=r*l,n.point(-s,i),n.point(0,i),n.point(s,i),n.point(s,0),n.point(s,-i),n.point(0,-i),n.point(-s,-i),n.point(-s,0),n.point(-s,i);else if(p(t[0]-e[0])>o){var a=t[0]<e[0]?s:-s;i=r*a/2,n.point(-a,i),n.point(0,i),n.point(a,i)}else n.point(e[0],e[1])}),[-s,-l]);function ve(t){var e=g(t),r=6*h,n=e>0,i=p(e)>o;function a(t,r){return g(t)*g(r)>e}function l(t,r,n){var i=[1,0,0],a=J(X(t),X(r)),l=Z(a,a),c=a[0],u=l-c*c;if(!u)return!n&&t;var f=e*l/u,h=-e*c/u,d=J(i,a),m=Q(i,f);K(m,Q(a,h));var g=d,v=Z(m,g),y=Z(g,g),x=v*v-y*(Z(m,m)-1);if(!(x<0)){var b=T(x),_=Q(g,(-v-b)/y);if(K(_,m),_=W(_),!n)return _;var w,k=t[0],A=r[0],M=t[1],S=r[1];A<k&&(w=k,k=A,A=w);var E=A-k,L=p(E-s)<o;if(!L&&S<M&&(w=M,M=S,S=w),L||E<o?L?M+S>0^_[1]<(p(_[0]-k)<o?M:S):M<=_[1]&&_[1]<=S:E>s^(k<=_[0]&&_[0]<=A)){var C=Q(g,(-v+b)/y);return K(C,m),[_,W(C)]}}}function c(e,r){var i=n?t:s-t,a=0;return e<-i?a|=1:e>i&&(a|=2),r<-i?a|=4:r>i&&(a|=8),a}return pe(a,(function(t){var e,r,o,u,f;return{lineStart:function(){u=o=!1,f=1},point:function(h,p){var d,m=[h,p],g=a(h,p),v=n?g?0:c(h,p):g?c(h+(h<0?s:-s),p):0;if(!e&&(u=o=g)&&t.lineStart(),g!==o&&(!(d=l(e,m))||oe(e,d)||oe(m,d))&&(m[2]=1),g!==o)f=0,g?(t.lineStart(),d=l(m,e),t.point(d[0],d[1])):(d=l(e,m),t.point(d[0],d[1],2),t.lineEnd()),e=d;else if(i&&e&&n^g){var y;v&r||!(y=l(m,e,!0))||(f=0,n?(t.lineStart(),t.point(y[0][0],y[0][1]),t.point(y[1][0],y[1][1]),t.lineEnd()):(t.point(y[1][0],y[1][1]),t.lineEnd(),t.lineStart(),t.point(y[0][0],y[0][1],3)))}!g||e&&oe(e,m)||t.point(m[0],m[1]),e=m,o=g,r=v},lineEnd:function(){o&&t.lineEnd(),e=null},clean:function(){return f|(u&&o)<<1}}}),(function(e,n,i,a){ne(a,t,r,i,e,n)}),n?[0,-t]:[-s,t-s])}function ye(t,r,n,i){function a(e,a){return t<=e&&e<=n&&r<=a&&a<=i}function s(e,a,o,s){var c=0,f=0;if(null==e||(c=l(e,o))!==(f=l(a,o))||u(e,a)<0^o>0)do{s.point(0===c||3===c?t:n,c>1?i:r)}while((c=(c+o+4)%4)!==f);else s.point(a[0],a[1])}function l(e,i){return p(e[0]-t)<o?i>0?0:3:p(e[0]-n)<o?i>0?2:1:p(e[1]-r)<o?i>0?1:0:i>0?3:2}function c(t,e){return u(t.x,e.x)}function u(t,e){var r=l(t,1),n=l(e,1);return r!==n?r-n:0===r?e[1]-t[1]:1===r?t[0]-e[0]:2===r?t[1]-e[1]:e[0]-t[0]}return function(o){var l,u,f,h,p,d,m,g,v,y,x,b=o,_=ae(),w={point:T,lineStart:function(){w.point=k,u&&u.push(f=[]);y=!0,v=!1,m=g=NaN},lineEnd:function(){l&&(k(h,p),d&&v&&_.rejoin(),l.push(_.result()));w.point=T,v&&b.lineEnd()},polygonStart:function(){b=_,l=[],u=[],x=!0},polygonEnd:function(){var r=function(){for(var e=0,r=0,n=u.length;r<n;++r)for(var a,o,s=u[r],l=1,c=s.length,f=s[0],h=f[0],p=f[1];l<c;++l)a=h,o=p,f=s[l],h=f[0],p=f[1],o<=i?p>i&&(h-a)*(i-o)>(p-o)*(t-a)&&++e:p<=i&&(h-a)*(i-o)<(p-o)*(t-a)&&--e;return e}(),n=x&&r,a=(l=e.merge(l)).length;(n||a)&&(o.polygonStart(),n&&(o.lineStart(),s(null,null,1,o),o.lineEnd()),a&&le(l,c,r,s,o),o.polygonEnd());b=o,l=u=f=null}};function T(t,e){a(t,e)&&b.point(t,e)}function k(e,o){var s=a(e,o);if(u&&f.push([e,o]),y)h=e,p=o,d=s,y=!1,s&&(b.lineStart(),b.point(e,o));else if(s&&v)b.point(e,o);else{var l=[m=Math.max(-1e9,Math.min(1e9,m)),g=Math.max(-1e9,Math.min(1e9,g))],c=[e=Math.max(-1e9,Math.min(1e9,e)),o=Math.max(-1e9,Math.min(1e9,o))];!function(t,e,r,n,i,a){var o,s=t[0],l=t[1],c=0,u=1,f=e[0]-s,h=e[1]-l;if(o=r-s,f||!(o>0)){if(o/=f,f<0){if(o<c)return;o<u&&(u=o)}else if(f>0){if(o>u)return;o>c&&(c=o)}if(o=i-s,f||!(o<0)){if(o/=f,f<0){if(o>u)return;o>c&&(c=o)}else if(f>0){if(o<c)return;o<u&&(u=o)}if(o=n-l,h||!(o>0)){if(o/=h,h<0){if(o<c)return;o<u&&(u=o)}else if(h>0){if(o>u)return;o>c&&(c=o)}if(o=a-l,h||!(o<0)){if(o/=h,h<0){if(o>u)return;o>c&&(c=o)}else if(h>0){if(o<c)return;o<u&&(u=o)}return c>0&&(t[0]=s+c*f,t[1]=l+c*h),u<1&&(e[0]=s+u*f,e[1]=l+u*h),!0}}}}}(l,c,t,r,n,i)?s&&(b.lineStart(),b.point(e,o),x=!1):(v||(b.lineStart(),b.point(l[0],l[1])),b.point(c[0],c[1]),s||b.lineEnd(),x=!1)}m=e,g=o,v=s}return w}}var xe,be,_e,we=r(),Te={sphere:E,point:E,lineStart:function(){Te.point=Ae,Te.lineEnd=ke},lineEnd:E,polygonStart:E,polygonEnd:E};function ke(){Te.point=Te.lineEnd=E}function Ae(t,e){xe=t*=h,be=_(e*=h),_e=g(e),Te.point=Me}function Me(t,e){t*=h;var r=_(e*=h),n=g(e),i=p(t-xe),a=g(i),o=n*_(i),s=_e*r-be*n*a,l=be*r+_e*n*a;we.add(m(T(o*o+s*s),l)),xe=t,be=r,_e=n}function Se(t){return we.reset(),z(t,Te),+we}var Ee=[null,null],Le={type:"LineString",coordinates:Ee};function Ce(t,e){return Ee[0]=t,Ee[1]=e,Se(Le)}var Pe={Feature:function(t,e){return Oe(t.geometry,e)},FeatureCollection:function(t,e){for(var r=t.features,n=-1,i=r.length;++n<i;)if(Oe(r[n].geometry,e))return!0;return!1}},Ie={Sphere:function(){return!0},Point:function(t,e){return ze(t.coordinates,e)},MultiPoint:function(t,e){for(var r=t.coordinates,n=-1,i=r.length;++n<i;)if(ze(r[n],e))return!0;return!1},LineString:function(t,e){return De(t.coordinates,e)},MultiLineString:function(t,e){for(var r=t.coordinates,n=-1,i=r.length;++n<i;)if(De(r[n],e))return!0;return!1},Polygon:function(t,e){return Re(t.coordinates,e)},MultiPolygon:function(t,e){for(var r=t.coordinates,n=-1,i=r.length;++n<i;)if(Re(r[n],e))return!0;return!1},GeometryCollection:function(t,e){for(var r=t.geometries,n=-1,i=r.length;++n<i;)if(Oe(r[n],e))return!0;return!1}};function Oe(t,e){return!(!t||!Ie.hasOwnProperty(t.type))&&Ie[t.type](t,e)}function ze(t,e){return 0===Ce(t,e)}function De(t,e){for(var r,n,i,a=0,o=t.length;a<o;a++){if(0===(n=Ce(t[a],e)))return!0;if(a>0&&(i=Ce(t[a],t[a-1]))>0&&r<=i&&n<=i&&(r+n-i)*(1-Math.pow((r-n)/i,2))<1e-12*i)return!0;r=n}return!1}function Re(t,e){return!!he(t.map(Fe),Be(e))}function Fe(t){return(t=t.map(Be)).pop(),t}function Be(t){return[t[0]*h,t[1]*h]}function Ne(t,r,n){var i=e.range(t,r-o,n).concat(r);return function(t){return i.map((function(e){return[t,e]}))}}function je(t,r,n){var i=e.range(t,r-o,n).concat(r);return function(t){return i.map((function(e){return[e,t]}))}}function Ue(){var t,r,n,i,a,s,l,c,u,f,h,d,m=10,g=m,y=90,x=360,b=2.5;function _(){return{type:"MultiLineString",coordinates:w()}}function w(){return e.range(v(i/y)*y,n,y).map(h).concat(e.range(v(c/x)*x,l,x).map(d)).concat(e.range(v(r/m)*m,t,m).filter((function(t){return p(t%y)>o})).map(u)).concat(e.range(v(s/g)*g,a,g).filter((function(t){return p(t%x)>o})).map(f))}return _.lines=function(){return w().map((function(t){return{type:"LineString",coordinates:t}}))},_.outline=function(){return{type:"Polygon",coordinates:[h(i).concat(d(l).slice(1),h(n).reverse().slice(1),d(c).reverse().slice(1))]}},_.extent=function(t){return arguments.length?_.extentMajor(t).extentMinor(t):_.extentMinor()},_.extentMajor=function(t){return arguments.length?(i=+t[0][0],n=+t[1][0],c=+t[0][1],l=+t[1][1],i>n&&(t=i,i=n,n=t),c>l&&(t=c,c=l,l=t),_.precision(b)):[[i,c],[n,l]]},_.extentMinor=function(e){return arguments.length?(r=+e[0][0],t=+e[1][0],s=+e[0][1],a=+e[1][1],r>t&&(e=r,r=t,t=e),s>a&&(e=s,s=a,a=e),_.precision(b)):[[r,s],[t,a]]},_.step=function(t){return arguments.length?_.stepMajor(t).stepMinor(t):_.stepMinor()},_.stepMajor=function(t){return arguments.length?(y=+t[0],x=+t[1],_):[y,x]},_.stepMinor=function(t){return arguments.length?(m=+t[0],g=+t[1],_):[m,g]},_.precision=function(e){return arguments.length?(b=+e,u=Ne(s,a,90),f=je(r,t,b),h=Ne(c,l,90),d=je(i,n,b),_):b},_.extentMajor([[-180,-90+o],[180,90-o]]).extentMinor([[-180,-80-o],[180,80+o]])}function Ve(t){return t}var He,qe,Ge,Ye,We=r(),Xe=r(),Ze={point:E,lineStart:E,lineEnd:E,polygonStart:function(){Ze.lineStart=Je,Ze.lineEnd=$e},polygonEnd:function(){Ze.lineStart=Ze.lineEnd=Ze.point=E,We.add(p(Xe)),Xe.reset()},result:function(){var t=We/2;return We.reset(),t}};function Je(){Ze.point=Ke}function Ke(t,e){Ze.point=Qe,He=Ge=t,qe=Ye=e}function Qe(t,e){Xe.add(Ye*t-Ge*e),Ge=t,Ye=e}function $e(){Qe(He,qe)}var tr=1/0,er=tr,rr=-tr,nr=rr,ir={point:function(t,e){t<tr&&(tr=t);t>rr&&(rr=t);e<er&&(er=e);e>nr&&(nr=e)},lineStart:E,lineEnd:E,polygonStart:E,polygonEnd:E,result:function(){var t=[[tr,er],[rr,nr]];return rr=nr=-(er=tr=1/0),t}};var ar,or,sr,lr,cr=0,ur=0,fr=0,hr=0,pr=0,dr=0,mr=0,gr=0,vr=0,yr={point:xr,lineStart:br,lineEnd:Tr,polygonStart:function(){yr.lineStart=kr,yr.lineEnd=Ar},polygonEnd:function(){yr.point=xr,yr.lineStart=br,yr.lineEnd=Tr},result:function(){var t=vr?[mr/vr,gr/vr]:dr?[hr/dr,pr/dr]:fr?[cr/fr,ur/fr]:[NaN,NaN];return cr=ur=fr=hr=pr=dr=mr=gr=vr=0,t}};function xr(t,e){cr+=t,ur+=e,++fr}function br(){yr.point=_r}function _r(t,e){yr.point=wr,xr(sr=t,lr=e)}function wr(t,e){var r=t-sr,n=e-lr,i=T(r*r+n*n);hr+=i*(sr+t)/2,pr+=i*(lr+e)/2,dr+=i,xr(sr=t,lr=e)}function Tr(){yr.point=xr}function kr(){yr.point=Mr}function Ar(){Sr(ar,or)}function Mr(t,e){yr.point=Sr,xr(ar=sr=t,or=lr=e)}function Sr(t,e){var r=t-sr,n=e-lr,i=T(r*r+n*n);hr+=i*(sr+t)/2,pr+=i*(lr+e)/2,dr+=i,mr+=(i=lr*t-sr*e)*(sr+t),gr+=i*(lr+e),vr+=3*i,xr(sr=t,lr=e)}function Er(t){this._context=t}Er.prototype={_radius:4.5,pointRadius:function(t){return this._radius=t,this},polygonStart:function(){this._line=0},polygonEnd:function(){this._line=NaN},lineStart:function(){this._point=0},lineEnd:function(){0===this._line&&this._context.closePath(),this._point=NaN},point:function(t,e){switch(this._point){case 0:this._context.moveTo(t,e),this._point=1;break;case 1:this._context.lineTo(t,e);break;default:this._context.moveTo(t+this._radius,e),this._context.arc(t,e,this._radius,0,u)}},result:E};var Lr,Cr,Pr,Ir,Or,zr=r(),Dr={point:E,lineStart:function(){Dr.point=Rr},lineEnd:function(){Lr&&Fr(Cr,Pr),Dr.point=E},polygonStart:function(){Lr=!0},polygonEnd:function(){Lr=null},result:function(){var t=+zr;return zr.reset(),t}};function Rr(t,e){Dr.point=Fr,Cr=Ir=t,Pr=Or=e}function Fr(t,e){Ir-=t,Or-=e,zr.add(T(Ir*Ir+Or*Or)),Ir=t,Or=e}function Br(){this._string=[]}function Nr(t){return"m0,"+t+"a"+t+","+t+" 0 1,1 0,"+-2*t+"a"+t+","+t+" 0 1,1 0,"+2*t+"z"}function jr(t){return function(e){var r=new Ur;for(var n in t)r[n]=t[n];return r.stream=e,r}}function Ur(){}function Vr(t,e,r){var n=t.clipExtent&&t.clipExtent();return t.scale(150).translate([0,0]),null!=n&&t.clipExtent(null),z(r,t.stream(ir)),e(ir.result()),null!=n&&t.clipExtent(n),t}function Hr(t,e,r){return Vr(t,(function(r){var n=e[1][0]-e[0][0],i=e[1][1]-e[0][1],a=Math.min(n/(r[1][0]-r[0][0]),i/(r[1][1]-r[0][1])),o=+e[0][0]+(n-a*(r[1][0]+r[0][0]))/2,s=+e[0][1]+(i-a*(r[1][1]+r[0][1]))/2;t.scale(150*a).translate([o,s])}),r)}function qr(t,e,r){return Hr(t,[[0,0],e],r)}function Gr(t,e,r){return Vr(t,(function(r){var n=+e,i=n/(r[1][0]-r[0][0]),a=(n-i*(r[1][0]+r[0][0]))/2,o=-i*r[0][1];t.scale(150*i).translate([a,o])}),r)}function Yr(t,e,r){return Vr(t,(function(r){var n=+e,i=n/(r[1][1]-r[0][1]),a=-i*r[0][0],o=(n-i*(r[1][1]+r[0][1]))/2;t.scale(150*i).translate([a,o])}),r)}Br.prototype={_radius:4.5,_circle:Nr(4.5),pointRadius:function(t){return(t=+t)!==this._radius&&(this._radius=t,this._circle=null),this},polygonStart:function(){this._line=0},polygonEnd:function(){this._line=NaN},lineStart:function(){this._point=0},lineEnd:function(){0===this._line&&this._string.push("Z"),this._point=NaN},point:function(t,e){switch(this._point){case 0:this._string.push("M",t,",",e),this._point=1;break;case 1:this._string.push("L",t,",",e);break;default:null==this._circle&&(this._circle=Nr(this._radius)),this._string.push("M",t,",",e,this._circle)}},result:function(){if(this._string.length){var t=this._string.join("");return this._string=[],t}return null}},Ur.prototype={constructor:Ur,point:function(t,e){this.stream.point(t,e)},sphere:function(){this.stream.sphere()},lineStart:function(){this.stream.lineStart()},lineEnd:function(){this.stream.lineEnd()},polygonStart:function(){this.stream.polygonStart()},polygonEnd:function(){this.stream.polygonEnd()}};var Wr=g(30*h);function Xr(t,e){return+e?function(t,e){function r(n,i,a,s,l,c,u,f,h,d,g,v,y,x){var b=u-n,_=f-i,w=b*b+_*_;if(w>4*e&&y--){var k=s+d,A=l+g,S=c+v,E=T(k*k+A*A+S*S),L=M(S/=E),C=p(p(S)-1)<o||p(a-h)<o?(a+h)/2:m(A,k),P=t(C,L),I=P[0],O=P[1],z=I-n,D=O-i,R=_*z-b*D;(R*R/w>e||p((b*z+_*D)/w-.5)>.3||s*d+l*g+c*v<Wr)&&(r(n,i,a,s,l,c,I,O,C,k/=E,A/=E,S,y,x),x.point(I,O),r(I,O,C,k,A,S,u,f,h,d,g,v,y,x))}}return function(e){var n,i,a,o,s,l,c,u,f,h,p,d,m={point:g,lineStart:v,lineEnd:x,polygonStart:function(){e.polygonStart(),m.lineStart=b},polygonEnd:function(){e.polygonEnd(),m.lineStart=v}};function g(r,n){r=t(r,n),e.point(r[0],r[1])}function v(){u=NaN,m.point=y,e.lineStart()}function y(n,i){var a=X([n,i]),o=t(n,i);r(u,f,c,h,p,d,u=o[0],f=o[1],c=n,h=a[0],p=a[1],d=a[2],16,e),e.point(u,f)}function x(){m.point=g,e.lineEnd()}function b(){v(),m.point=_,m.lineEnd=w}function _(t,e){y(n=t,e),i=u,a=f,o=h,s=p,l=d,m.point=y}function w(){r(u,f,c,h,p,d,i,a,n,o,s,l,16,e),m.lineEnd=x,x()}return m}}(t,e):function(t){return jr({point:function(e,r){e=t(e,r),this.stream.point(e[0],e[1])}})}(t)}var Zr=jr({point:function(t,e){this.stream.point(t*h,e*h)}});function Jr(t,e,r,n,i){function a(a,o){return[e+t*(a*=n),r-t*(o*=i)]}return a.invert=function(a,o){return[(a-e)/t*n,(r-o)/t*i]},a}function Kr(t,e,r,n,i,a){var o=g(a),s=_(a),l=o*t,c=s*t,u=o/t,f=s/t,h=(s*r-o*e)/t,p=(s*e+o*r)/t;function d(t,a){return[l*(t*=n)-c*(a*=i)+e,r-c*t-l*a]}return d.invert=function(t,e){return[n*(u*t-f*e+h),i*(p-f*t-u*e)]},d}function Qr(t){return $r((function(){return t}))()}function $r(t){var e,r,n,i,a,o,s,l,c,u,p=150,d=480,m=250,g=0,v=0,y=0,x=0,b=0,_=0,w=1,k=1,A=null,M=ge,S=null,E=Ve,L=.5;function C(t){return l(t[0]*h,t[1]*h)}function P(t){return(t=l.invert(t[0],t[1]))&&[t[0]*f,t[1]*f]}function I(){var t=Kr(p,0,0,w,k,_).apply(null,e(g,v)),n=(_?Kr:Jr)(p,d-t[0],m-t[1],w,k,_);return r=Qt(y,x,b),s=Jt(e,n),l=Jt(r,s),o=Xr(s,L),O()}function O(){return c=u=null,C}return C.stream=function(t){return c&&u===t?c:c=Zr(function(t){return jr({point:function(e,r){var n=t(e,r);return this.stream.point(n[0],n[1])}})}(r)(M(o(E(u=t)))))},C.preclip=function(t){return arguments.length?(M=t,A=void 0,O()):M},C.postclip=function(t){return arguments.length?(E=t,S=n=i=a=null,O()):E},C.clipAngle=function(t){return arguments.length?(M=+t?ve(A=t*h):(A=null,ge),O()):A*f},C.clipExtent=function(t){return arguments.length?(E=null==t?(S=n=i=a=null,Ve):ye(S=+t[0][0],n=+t[0][1],i=+t[1][0],a=+t[1][1]),O()):null==S?null:[[S,n],[i,a]]},C.scale=function(t){return arguments.length?(p=+t,I()):p},C.translate=function(t){return arguments.length?(d=+t[0],m=+t[1],I()):[d,m]},C.center=function(t){return arguments.length?(g=t[0]%360*h,v=t[1]%360*h,I()):[g*f,v*f]},C.rotate=function(t){return arguments.length?(y=t[0]%360*h,x=t[1]%360*h,b=t.length>2?t[2]%360*h:0,I()):[y*f,x*f,b*f]},C.angle=function(t){return arguments.length?(_=t%360*h,I()):_*f},C.reflectX=function(t){return arguments.length?(w=t?-1:1,I()):w<0},C.reflectY=function(t){return arguments.length?(k=t?-1:1,I()):k<0},C.precision=function(t){return arguments.length?(o=Xr(s,L=t*t),O()):T(L)},C.fitExtent=function(t,e){return Hr(C,t,e)},C.fitSize=function(t,e){return qr(C,t,e)},C.fitWidth=function(t,e){return Gr(C,t,e)},C.fitHeight=function(t,e){return Yr(C,t,e)},function(){return e=t.apply(this,arguments),C.invert=e.invert&&P,I()}}function tn(t){var e=0,r=s/3,n=$r(t),i=n(e,r);return i.parallels=function(t){return arguments.length?n(e=t[0]*h,r=t[1]*h):[e*f,r*f]},i}function en(t,e){var r=_(t),n=(r+_(e))/2;if(p(n)<o)return function(t){var e=g(t);function r(t,r){return[t*e,_(r)/e]}return r.invert=function(t,r){return[t/e,M(r*e)]},r}(t);var i=1+r*(2*n-r),a=T(i)/n;function l(t,e){var r=T(i-2*n*_(e))/n;return[r*_(t*=n),a-r*g(t)]}return l.invert=function(t,e){var r=a-e,o=m(t,p(r))*w(r);return r*n<0&&(o-=s*w(t)*w(r)),[o/n,M((i-(t*t+r*r)*n*n)/(2*n))]},l}function rn(){return tn(en).scale(155.424).center([0,33.6442])}function nn(){return rn().parallels([29.5,45.5]).scale(1070).translate([480,250]).rotate([96,0]).center([-.6,38.7])}function an(t){return function(e,r){var n=g(e),i=g(r),a=t(n*i);return[a*i*_(e),a*_(r)]}}function on(t){return function(e,r){var n=T(e*e+r*r),i=t(n),a=_(i),o=g(i);return[m(e*a,n*o),M(n&&r*a/n)]}}var sn=an((function(t){return T(2/(1+t))}));sn.invert=on((function(t){return 2*M(t/2)}));var ln=an((function(t){return(t=A(t))&&t/_(t)}));function cn(t,e){return[t,x(k((l+e)/2))]}function un(t){var e,r,n,i=Qr(t),a=i.center,o=i.scale,l=i.translate,c=i.clipExtent,u=null;function f(){var a=s*o(),l=i(re(i.rotate()).invert([0,0]));return c(null==u?[[l[0]-a,l[1]-a],[l[0]+a,l[1]+a]]:t===cn?[[Math.max(l[0]-a,u),e],[Math.min(l[0]+a,r),n]]:[[u,Math.max(l[1]-a,e)],[r,Math.min(l[1]+a,n)]])}return i.scale=function(t){return arguments.length?(o(t),f()):o()},i.translate=function(t){return arguments.length?(l(t),f()):l()},i.center=function(t){return arguments.length?(a(t),f()):a()},i.clipExtent=function(t){return arguments.length?(null==t?u=e=r=n=null:(u=+t[0][0],e=+t[0][1],r=+t[1][0],n=+t[1][1]),f()):null==u?null:[[u,e],[r,n]]},f()}function fn(t){return k((l+t)/2)}function hn(t,e){var r=g(t),n=t===e?_(t):x(r/g(e))/x(fn(e)/fn(t)),i=r*b(fn(t),n)/n;if(!n)return cn;function a(t,e){i>0?e<-l+o&&(e=-l+o):e>l-o&&(e=l-o);var r=i/b(fn(e),n);return[r*_(n*t),i-r*g(n*t)]}return a.invert=function(t,e){var r=i-e,a=w(n)*T(t*t+r*r),o=m(t,p(r))*w(r);return r*n<0&&(o-=s*w(t)*w(r)),[o/n,2*d(b(i/a,1/n))-l]},a}function pn(t,e){return[t,e]}function dn(t,e){var r=g(t),n=t===e?_(t):(r-g(e))/(e-t),i=r/n+t;if(p(n)<o)return pn;function a(t,e){var r=i-e,a=n*t;return[r*_(a),i-r*g(a)]}return a.invert=function(t,e){var r=i-e,a=m(t,p(r))*w(r);return r*n<0&&(a-=s*w(t)*w(r)),[a/n,i-w(n)*T(t*t+r*r)]},a}ln.invert=on((function(t){return t})),cn.invert=function(t,e){return[t,2*d(y(e))-l]},pn.invert=pn;var mn=1.340264,gn=-.081106,vn=893e-6,yn=.003796,xn=T(3)/2;function bn(t,e){var r=M(xn*_(e)),n=r*r,i=n*n*n;return[t*g(r)/(xn*(mn+3*gn*n+i*(7*vn+9*yn*n))),r*(mn+gn*n+i*(vn+yn*n))]}function _n(t,e){var r=g(e),n=g(t)*r;return[r*_(t)/n,_(e)/n]}function wn(t,e){var r=e*e,n=r*r;return[t*(.8707-.131979*r+n*(n*(.003971*r-.001529*n)-.013791)),e*(1.007226+r*(.015085+n*(.028874*r-.044475-.005916*n)))]}function Tn(t,e){return[g(e)*_(t),_(e)]}function kn(t,e){var r=g(e),n=1+g(t)*r;return[r*_(t)/n,_(e)/n]}function An(t,e){return[x(k((l+e)/2)),-t]}bn.invert=function(t,e){for(var r,n=e,i=n*n,a=i*i*i,o=0;o<12&&(a=(i=(n-=r=(n*(mn+gn*i+a*(vn+yn*i))-e)/(mn+3*gn*i+a*(7*vn+9*yn*i)))*n)*i*i,!(p(r)<1e-12));++o);return[xn*t*(mn+3*gn*i+a*(7*vn+9*yn*i))/g(n),M(_(n)/xn)]},_n.invert=on(d),wn.invert=function(t,e){var r,n=e,i=25;do{var a=n*n,s=a*a;n-=r=(n*(1.007226+a*(.015085+s*(.028874*a-.044475-.005916*s)))-e)/(1.007226+a*(.045255+s*(.259866*a-.311325-.005916*11*s)))}while(p(r)>o&&--i>0);return[t/(.8707+(a=n*n)*(a*(a*a*a*(.003971-.001529*a)-.013791)-.131979)),n]},Tn.invert=on(M),kn.invert=on((function(t){return 2*d(t)})),An.invert=function(t,e){return[-e,2*d(y(t))-l]},t.geoAlbers=nn,t.geoAlbersUsa=function(){var t,e,r,n,i,a,s=nn(),l=rn().rotate([154,0]).center([-2,58.5]).parallels([55,65]),c=rn().rotate([157,0]).center([-3,19.9]).parallels([8,18]),u={point:function(t,e){a=[t,e]}};function f(t){var e=t[0],o=t[1];return a=null,r.point(e,o),a||(n.point(e,o),a)||(i.point(e,o),a)}function h(){return t=e=null,f}return f.invert=function(t){var e=s.scale(),r=s.translate(),n=(t[0]-r[0])/e,i=(t[1]-r[1])/e;return(i>=.12&&i<.234&&n>=-.425&&n<-.214?l:i>=.166&&i<.234&&n>=-.214&&n<-.115?c:s).invert(t)},f.stream=function(r){return t&&e===r?t:(n=[s.stream(e=r),l.stream(r),c.stream(r)],i=n.length,t={point:function(t,e){for(var r=-1;++r<i;)n[r].point(t,e)},sphere:function(){for(var t=-1;++t<i;)n[t].sphere()},lineStart:function(){for(var t=-1;++t<i;)n[t].lineStart()},lineEnd:function(){for(var t=-1;++t<i;)n[t].lineEnd()},polygonStart:function(){for(var t=-1;++t<i;)n[t].polygonStart()},polygonEnd:function(){for(var t=-1;++t<i;)n[t].polygonEnd()}});var n,i},f.precision=function(t){return arguments.length?(s.precision(t),l.precision(t),c.precision(t),h()):s.precision()},f.scale=function(t){return arguments.length?(s.scale(t),l.scale(.35*t),c.scale(t),f.translate(s.translate())):s.scale()},f.translate=function(t){if(!arguments.length)return s.translate();var e=s.scale(),a=+t[0],f=+t[1];return r=s.translate(t).clipExtent([[a-.455*e,f-.238*e],[a+.455*e,f+.238*e]]).stream(u),n=l.translate([a-.307*e,f+.201*e]).clipExtent([[a-.425*e+o,f+.12*e+o],[a-.214*e-o,f+.234*e-o]]).stream(u),i=c.translate([a-.205*e,f+.212*e]).clipExtent([[a-.214*e+o,f+.166*e+o],[a-.115*e-o,f+.234*e-o]]).stream(u),h()},f.fitExtent=function(t,e){return Hr(f,t,e)},f.fitSize=function(t,e){return qr(f,t,e)},f.fitWidth=function(t,e){return Gr(f,t,e)},f.fitHeight=function(t,e){return Yr(f,t,e)},f.scale(1070)},t.geoArea=function(t){return U.reset(),z(t,V),2*U},t.geoAzimuthalEqualArea=function(){return Qr(sn).scale(124.75).clipAngle(179.999)},t.geoAzimuthalEqualAreaRaw=sn,t.geoAzimuthalEquidistant=function(){return Qr(ln).scale(79.4188).clipAngle(179.999)},t.geoAzimuthalEquidistantRaw=ln,t.geoBounds=function(t){var e,r,n,i,a,o,s;if(nt=rt=-(tt=et=1/0),lt=[],z(t,St),r=lt.length){for(lt.sort(Rt),e=1,a=[n=lt[0]];e<r;++e)Ft(n,(i=lt[e])[0])||Ft(n,i[1])?(Dt(n[0],i[1])>Dt(n[0],n[1])&&(n[1]=i[1]),Dt(i[0],n[1])>Dt(n[0],n[1])&&(n[0]=i[0])):a.push(n=i);for(o=-1/0,e=0,n=a[r=a.length-1];e<=r;n=i,++e)i=a[e],(s=Dt(n[1],i[0]))>o&&(o=s,tt=i[0],rt=n[1])}return lt=ct=null,tt===1/0||et===1/0?[[NaN,NaN],[NaN,NaN]]:[[tt,et],[rt,nt]]},t.geoCentroid=function(t){ut=ft=ht=pt=dt=mt=gt=vt=yt=xt=bt=0,z(t,Bt);var e=yt,r=xt,n=bt,i=e*e+r*r+n*n;return i<1e-12&&(e=mt,r=gt,n=vt,ft<o&&(e=ht,r=pt,n=dt),(i=e*e+r*r+n*n)<1e-12)?[NaN,NaN]:[m(r,e)*f,M(n/T(i))*f]},t.geoCircle=function(){var t,e,r=Zt([0,0]),n=Zt(90),i=Zt(6),a={point:function(r,n){t.push(r=e(r,n)),r[0]*=f,r[1]*=f}};function o(){var o=r.apply(this,arguments),s=n.apply(this,arguments)*h,l=i.apply(this,arguments)*h;return t=[],e=Qt(-o[0]*h,-o[1]*h,0).invert,ne(a,s,l,1),o={type:"Polygon",coordinates:[t]},t=e=null,o}return o.center=function(t){return arguments.length?(r="function"==typeof t?t:Zt([+t[0],+t[1]]),o):r},o.radius=function(t){return arguments.length?(n="function"==typeof t?t:Zt(+t),o):n},o.precision=function(t){return arguments.length?(i="function"==typeof t?t:Zt(+t),o):i},o},t.geoClipAntimeridian=ge,t.geoClipCircle=ve,t.geoClipExtent=function(){var t,e,r,n=0,i=0,a=960,o=500;return r={stream:function(r){return t&&e===r?t:t=ye(n,i,a,o)(e=r)},extent:function(s){return arguments.length?(n=+s[0][0],i=+s[0][1],a=+s[1][0],o=+s[1][1],t=e=null,r):[[n,i],[a,o]]}}},t.geoClipRectangle=ye,t.geoConicConformal=function(){return tn(hn).scale(109.5).parallels([30,30])},t.geoConicConformalRaw=hn,t.geoConicEqualArea=rn,t.geoConicEqualAreaRaw=en,t.geoConicEquidistant=function(){return tn(dn).scale(131.154).center([0,13.9389])},t.geoConicEquidistantRaw=dn,t.geoContains=function(t,e){return(t&&Pe.hasOwnProperty(t.type)?Pe[t.type]:Oe)(t,e)},t.geoDistance=Ce,t.geoEqualEarth=function(){return Qr(bn).scale(177.158)},t.geoEqualEarthRaw=bn,t.geoEquirectangular=function(){return Qr(pn).scale(152.63)},t.geoEquirectangularRaw=pn,t.geoGnomonic=function(){return Qr(_n).scale(144.049).clipAngle(60)},t.geoGnomonicRaw=_n,t.geoGraticule=Ue,t.geoGraticule10=function(){return Ue()()},t.geoIdentity=function(){var t,e,r,n,i,a,o,s=1,l=0,c=0,u=1,p=1,d=0,m=null,v=1,y=1,x=jr({point:function(t,e){var r=T([t,e]);this.stream.point(r[0],r[1])}}),b=Ve;function w(){return v=s*u,y=s*p,a=o=null,T}function T(r){var n=r[0]*v,i=r[1]*y;if(d){var a=i*t-n*e;n=n*t+i*e,i=a}return[n+l,i+c]}return T.invert=function(r){var n=r[0]-l,i=r[1]-c;if(d){var a=i*t+n*e;n=n*t-i*e,i=a}return[n/v,i/y]},T.stream=function(t){return a&&o===t?a:a=x(b(o=t))},T.postclip=function(t){return arguments.length?(b=t,m=r=n=i=null,w()):b},T.clipExtent=function(t){return arguments.length?(b=null==t?(m=r=n=i=null,Ve):ye(m=+t[0][0],r=+t[0][1],n=+t[1][0],i=+t[1][1]),w()):null==m?null:[[m,r],[n,i]]},T.scale=function(t){return arguments.length?(s=+t,w()):s},T.translate=function(t){return arguments.length?(l=+t[0],c=+t[1],w()):[l,c]},T.angle=function(r){return arguments.length?(e=_(d=r%360*h),t=g(d),w()):d*f},T.reflectX=function(t){return arguments.length?(u=t?-1:1,w()):u<0},T.reflectY=function(t){return arguments.length?(p=t?-1:1,w()):p<0},T.fitExtent=function(t,e){return Hr(T,t,e)},T.fitSize=function(t,e){return qr(T,t,e)},T.fitWidth=function(t,e){return Gr(T,t,e)},T.fitHeight=function(t,e){return Yr(T,t,e)},T},t.geoInterpolate=function(t,e){var r=t[0]*h,n=t[1]*h,i=e[0]*h,a=e[1]*h,o=g(n),s=_(n),l=g(a),c=_(a),u=o*g(r),p=o*_(r),d=l*g(i),v=l*_(i),y=2*M(T(S(a-n)+o*l*S(i-r))),x=_(y),b=y?function(t){var e=_(t*=y)/x,r=_(y-t)/x,n=r*u+e*d,i=r*p+e*v,a=r*s+e*c;return[m(i,n)*f,m(a,T(n*n+i*i))*f]}:function(){return[r*f,n*f]};return b.distance=y,b},t.geoLength=Se,t.geoMercator=function(){return un(cn).scale(961/u)},t.geoMercatorRaw=cn,t.geoNaturalEarth1=function(){return Qr(wn).scale(175.295)},t.geoNaturalEarth1Raw=wn,t.geoOrthographic=function(){return Qr(Tn).scale(249.5).clipAngle(90+o)},t.geoOrthographicRaw=Tn,t.geoPath=function(t,e){var r,n,i=4.5;function a(t){return t&&("function"==typeof i&&n.pointRadius(+i.apply(this,arguments)),z(t,r(n))),n.result()}return a.area=function(t){return z(t,r(Ze)),Ze.result()},a.measure=function(t){return z(t,r(Dr)),Dr.result()},a.bounds=function(t){return z(t,r(ir)),ir.result()},a.centroid=function(t){return z(t,r(yr)),yr.result()},a.projection=function(e){return arguments.length?(r=null==e?(t=null,Ve):(t=e).stream,a):t},a.context=function(t){return arguments.length?(n=null==t?(e=null,new Br):new Er(e=t),"function"!=typeof i&&n.pointRadius(i),a):e},a.pointRadius=function(t){return arguments.length?(i="function"==typeof t?t:(n.pointRadius(+t),+t),a):i},a.projection(t).context(e)},t.geoProjection=Qr,t.geoProjectionMutator=$r,t.geoRotation=re,t.geoStereographic=function(){return Qr(kn).scale(250).clipAngle(142)},t.geoStereographicRaw=kn,t.geoStream=z,t.geoTransform=function(t){return{stream:jr(t)}},t.geoTransverseMercator=function(){var t=un(An),e=t.center,r=t.rotate;return t.center=function(t){return arguments.length?e([-t[1],t[0]]):[(t=e())[1],-t[0]]},t.rotate=function(t){return arguments.length?r([t[0],t[1],t.length>2?t[2]+90:90]):[(t=r())[0],t[1],t[2]-90]},r([0,0,90]).scale(159.155)},t.geoTransverseMercatorRaw=An,Object.defineProperty(t,"__esModule",{value:!0})}))},{"d3-array":102}],110:[function(t,e,r){!function(t,n){"object"==typeof r&&void 0!==e?n(r):n((t=t||self).d3=t.d3||{})}(this,(function(t){"use strict";function e(t,e){return t.parent===e.parent?1:2}function r(t,e){return t+e.x}function n(t,e){return Math.max(t,e.y)}function i(t){var e=0,r=t.children,n=r&&r.length;if(n)for(;--n>=0;)e+=r[n].value;else e=1;t.value=e}function a(t,e){var r,n,i,a,s,u=new c(t),f=+t.value&&(u.value=t.value),h=[u];for(null==e&&(e=o);r=h.pop();)if(f&&(r.value=+r.data.value),(i=e(r.data))&&(s=i.length))for(r.children=new Array(s),a=s-1;a>=0;--a)h.push(n=r.children[a]=new c(i[a])),n.parent=r,n.depth=r.depth+1;return u.eachBefore(l)}function o(t){return t.children}function s(t){t.data=t.data.data}function l(t){var e=0;do{t.height=e}while((t=t.parent)&&t.height<++e)}function c(t){this.data=t,this.depth=this.height=0,this.parent=null}c.prototype=a.prototype={constructor:c,count:function(){return this.eachAfter(i)},each:function(t){var e,r,n,i,a=this,o=[a];do{for(e=o.reverse(),o=[];a=e.pop();)if(t(a),r=a.children)for(n=0,i=r.length;n<i;++n)o.push(r[n])}while(o.length);return this},eachAfter:function(t){for(var e,r,n,i=this,a=[i],o=[];i=a.pop();)if(o.push(i),e=i.children)for(r=0,n=e.length;r<n;++r)a.push(e[r]);for(;i=o.pop();)t(i);return this},eachBefore:function(t){for(var e,r,n=this,i=[n];n=i.pop();)if(t(n),e=n.children)for(r=e.length-1;r>=0;--r)i.push(e[r]);return this},sum:function(t){return this.eachAfter((function(e){for(var r=+t(e.data)||0,n=e.children,i=n&&n.length;--i>=0;)r+=n[i].value;e.value=r}))},sort:function(t){return this.eachBefore((function(e){e.children&&e.children.sort(t)}))},path:function(t){for(var e=this,r=function(t,e){if(t===e)return t;var r=t.ancestors(),n=e.ancestors(),i=null;t=r.pop(),e=n.pop();for(;t===e;)i=t,t=r.pop(),e=n.pop();return i}(e,t),n=[e];e!==r;)e=e.parent,n.push(e);for(var i=n.length;t!==r;)n.splice(i,0,t),t=t.parent;return n},ancestors:function(){for(var t=this,e=[t];t=t.parent;)e.push(t);return e},descendants:function(){var t=[];return this.each((function(e){t.push(e)})),t},leaves:function(){var t=[];return this.eachBefore((function(e){e.children||t.push(e)})),t},links:function(){var t=this,e=[];return t.each((function(r){r!==t&&e.push({source:r.parent,target:r})})),e},copy:function(){return a(this).eachBefore(s)}};var u=Array.prototype.slice;function f(t){for(var e,r,n=0,i=(t=function(t){for(var e,r,n=t.length;n;)r=Math.random()*n--|0,e=t[n],t[n]=t[r],t[r]=e;return t}(u.call(t))).length,a=[];n<i;)e=t[n],r&&d(r,e)?++n:(r=g(a=h(a,e)),n=0);return r}function h(t,e){var r,n;if(m(e,t))return[e];for(r=0;r<t.length;++r)if(p(e,t[r])&&m(v(t[r],e),t))return[t[r],e];for(r=0;r<t.length-1;++r)for(n=r+1;n<t.length;++n)if(p(v(t[r],t[n]),e)&&p(v(t[r],e),t[n])&&p(v(t[n],e),t[r])&&m(y(t[r],t[n],e),t))return[t[r],t[n],e];throw new Error}function p(t,e){var r=t.r-e.r,n=e.x-t.x,i=e.y-t.y;return r<0||r*r<n*n+i*i}function d(t,e){var r=t.r-e.r+1e-6,n=e.x-t.x,i=e.y-t.y;return r>0&&r*r>n*n+i*i}function m(t,e){for(var r=0;r<e.length;++r)if(!d(t,e[r]))return!1;return!0}function g(t){switch(t.length){case 1:return{x:(e=t[0]).x,y:e.y,r:e.r};case 2:return v(t[0],t[1]);case 3:return y(t[0],t[1],t[2])}var e}function v(t,e){var r=t.x,n=t.y,i=t.r,a=e.x,o=e.y,s=e.r,l=a-r,c=o-n,u=s-i,f=Math.sqrt(l*l+c*c);return{x:(r+a+l/f*u)/2,y:(n+o+c/f*u)/2,r:(f+i+s)/2}}function y(t,e,r){var n=t.x,i=t.y,a=t.r,o=e.x,s=e.y,l=e.r,c=r.x,u=r.y,f=r.r,h=n-o,p=n-c,d=i-s,m=i-u,g=l-a,v=f-a,y=n*n+i*i-a*a,x=y-o*o-s*s+l*l,b=y-c*c-u*u+f*f,_=p*d-h*m,w=(d*b-m*x)/(2*_)-n,T=(m*g-d*v)/_,k=(p*x-h*b)/(2*_)-i,A=(h*v-p*g)/_,M=T*T+A*A-1,S=2*(a+w*T+k*A),E=w*w+k*k-a*a,L=-(M?(S+Math.sqrt(S*S-4*M*E))/(2*M):E/S);return{x:n+w+T*L,y:i+k+A*L,r:L}}function x(t,e,r){var n,i,a,o,s=t.x-e.x,l=t.y-e.y,c=s*s+l*l;c?(i=e.r+r.r,i*=i,o=t.r+r.r,i>(o*=o)?(n=(c+o-i)/(2*c),a=Math.sqrt(Math.max(0,o/c-n*n)),r.x=t.x-n*s-a*l,r.y=t.y-n*l+a*s):(n=(c+i-o)/(2*c),a=Math.sqrt(Math.max(0,i/c-n*n)),r.x=e.x+n*s-a*l,r.y=e.y+n*l+a*s)):(r.x=e.x+r.r,r.y=e.y)}function b(t,e){var r=t.r+e.r-1e-6,n=e.x-t.x,i=e.y-t.y;return r>0&&r*r>n*n+i*i}function _(t){var e=t._,r=t.next._,n=e.r+r.r,i=(e.x*r.r+r.x*e.r)/n,a=(e.y*r.r+r.y*e.r)/n;return i*i+a*a}function w(t){this._=t,this.next=null,this.previous=null}function T(t){if(!(i=t.length))return 0;var e,r,n,i,a,o,s,l,c,u,h;if((e=t[0]).x=0,e.y=0,!(i>1))return e.r;if(r=t[1],e.x=-r.r,r.x=e.r,r.y=0,!(i>2))return e.r+r.r;x(r,e,n=t[2]),e=new w(e),r=new w(r),n=new w(n),e.next=n.previous=r,r.next=e.previous=n,n.next=r.previous=e;t:for(s=3;s<i;++s){x(e._,r._,n=t[s]),n=new w(n),l=r.next,c=e.previous,u=r._.r,h=e._.r;do{if(u<=h){if(b(l._,n._)){r=l,e.next=r,r.previous=e,--s;continue t}u+=l._.r,l=l.next}else{if(b(c._,n._)){(e=c).next=r,r.previous=e,--s;continue t}h+=c._.r,c=c.previous}}while(l!==c.next);for(n.previous=e,n.next=r,e.next=r.previous=r=n,a=_(e);(n=n.next)!==r;)(o=_(n))<a&&(e=n,a=o);r=e.next}for(e=[r._],n=r;(n=n.next)!==r;)e.push(n._);for(n=f(e),s=0;s<i;++s)(e=t[s]).x-=n.x,e.y-=n.y;return n.r}function k(t){return null==t?null:A(t)}function A(t){if("function"!=typeof t)throw new Error;return t}function M(){return 0}function S(t){return function(){return t}}function E(t){return Math.sqrt(t.value)}function L(t){return function(e){e.children||(e.r=Math.max(0,+t(e)||0))}}function C(t,e){return function(r){if(n=r.children){var n,i,a,o=n.length,s=t(r)*e||0;if(s)for(i=0;i<o;++i)n[i].r+=s;if(a=T(n),s)for(i=0;i<o;++i)n[i].r-=s;r.r=a+s}}}function P(t){return function(e){var r=e.parent;e.r*=t,r&&(e.x=r.x+t*e.x,e.y=r.y+t*e.y)}}function I(t){t.x0=Math.round(t.x0),t.y0=Math.round(t.y0),t.x1=Math.round(t.x1),t.y1=Math.round(t.y1)}function O(t,e,r,n,i){for(var a,o=t.children,s=-1,l=o.length,c=t.value&&(n-e)/t.value;++s<l;)(a=o[s]).y0=r,a.y1=i,a.x0=e,a.x1=e+=a.value*c}var z={depth:-1},D={};function R(t){return t.id}function F(t){return t.parentId}function B(t,e){return t.parent===e.parent?1:2}function N(t){var e=t.children;return e?e[0]:t.t}function j(t){var e=t.children;return e?e[e.length-1]:t.t}function U(t,e,r){var n=r/(e.i-t.i);e.c-=n,e.s+=r,t.c+=n,e.z+=r,e.m+=r}function V(t,e,r){return t.a.parent===e.parent?t.a:r}function H(t,e){this._=t,this.parent=null,this.children=null,this.A=null,this.a=this,this.z=0,this.m=0,this.c=0,this.s=0,this.t=null,this.i=e}function q(t,e,r,n,i){for(var a,o=t.children,s=-1,l=o.length,c=t.value&&(i-r)/t.value;++s<l;)(a=o[s]).x0=e,a.x1=n,a.y0=r,a.y1=r+=a.value*c}H.prototype=Object.create(c.prototype);var G=(1+Math.sqrt(5))/2;function Y(t,e,r,n,i,a){for(var o,s,l,c,u,f,h,p,d,m,g,v=[],y=e.children,x=0,b=0,_=y.length,w=e.value;x<_;){l=i-r,c=a-n;do{u=y[b++].value}while(!u&&b<_);for(f=h=u,g=u*u*(m=Math.max(c/l,l/c)/(w*t)),d=Math.max(h/g,g/f);b<_;++b){if(u+=s=y[b].value,s<f&&(f=s),s>h&&(h=s),g=u*u*m,(p=Math.max(h/g,g/f))>d){u-=s;break}d=p}v.push(o={value:u,dice:l<c,children:y.slice(x,b)}),o.dice?O(o,r,n,i,w?n+=c*u/w:a):q(o,r,n,w?r+=l*u/w:i,a),w-=u,x=b}return v}var W=function t(e){function r(t,r,n,i,a){Y(e,t,r,n,i,a)}return r.ratio=function(e){return t((e=+e)>1?e:1)},r}(G);var X=function t(e){function r(t,r,n,i,a){if((o=t._squarify)&&o.ratio===e)for(var o,s,l,c,u,f=-1,h=o.length,p=t.value;++f<h;){for(l=(s=o[f]).children,c=s.value=0,u=l.length;c<u;++c)s.value+=l[c].value;s.dice?O(s,r,n,i,n+=(a-n)*s.value/p):q(s,r,n,r+=(i-r)*s.value/p,a),p-=s.value}else t._squarify=o=Y(e,t,r,n,i,a),o.ratio=e}return r.ratio=function(e){return t((e=+e)>1?e:1)},r}(G);t.cluster=function(){var t=e,i=1,a=1,o=!1;function s(e){var s,l=0;e.eachAfter((function(e){var i=e.children;i?(e.x=function(t){return t.reduce(r,0)/t.length}(i),e.y=function(t){return 1+t.reduce(n,0)}(i)):(e.x=s?l+=t(e,s):0,e.y=0,s=e)}));var c=function(t){for(var e;e=t.children;)t=e[0];return t}(e),u=function(t){for(var e;e=t.children;)t=e[e.length-1];return t}(e),f=c.x-t(c,u)/2,h=u.x+t(u,c)/2;return e.eachAfter(o?function(t){t.x=(t.x-e.x)*i,t.y=(e.y-t.y)*a}:function(t){t.x=(t.x-f)/(h-f)*i,t.y=(1-(e.y?t.y/e.y:1))*a})}return s.separation=function(e){return arguments.length?(t=e,s):t},s.size=function(t){return arguments.length?(o=!1,i=+t[0],a=+t[1],s):o?null:[i,a]},s.nodeSize=function(t){return arguments.length?(o=!0,i=+t[0],a=+t[1],s):o?[i,a]:null},s},t.hierarchy=a,t.pack=function(){var t=null,e=1,r=1,n=M;function i(i){return i.x=e/2,i.y=r/2,t?i.eachBefore(L(t)).eachAfter(C(n,.5)).eachBefore(P(1)):i.eachBefore(L(E)).eachAfter(C(M,1)).eachAfter(C(n,i.r/Math.min(e,r))).eachBefore(P(Math.min(e,r)/(2*i.r))),i}return i.radius=function(e){return arguments.length?(t=k(e),i):t},i.size=function(t){return arguments.length?(e=+t[0],r=+t[1],i):[e,r]},i.padding=function(t){return arguments.length?(n="function"==typeof t?t:S(+t),i):n},i},t.packEnclose=f,t.packSiblings=function(t){return T(t),t},t.partition=function(){var t=1,e=1,r=0,n=!1;function i(i){var a=i.height+1;return i.x0=i.y0=r,i.x1=t,i.y1=e/a,i.eachBefore(function(t,e){return function(n){n.children&&O(n,n.x0,t*(n.depth+1)/e,n.x1,t*(n.depth+2)/e);var i=n.x0,a=n.y0,o=n.x1-r,s=n.y1-r;o<i&&(i=o=(i+o)/2),s<a&&(a=s=(a+s)/2),n.x0=i,n.y0=a,n.x1=o,n.y1=s}}(e,a)),n&&i.eachBefore(I),i}return i.round=function(t){return arguments.length?(n=!!t,i):n},i.size=function(r){return arguments.length?(t=+r[0],e=+r[1],i):[t,e]},i.padding=function(t){return arguments.length?(r=+t,i):r},i},t.stratify=function(){var t=R,e=F;function r(r){var n,i,a,o,s,u,f,h=r.length,p=new Array(h),d={};for(i=0;i<h;++i)n=r[i],s=p[i]=new c(n),null!=(u=t(n,i,r))&&(u+="")&&(d[f="$"+(s.id=u)]=f in d?D:s);for(i=0;i<h;++i)if(s=p[i],null!=(u=e(r[i],i,r))&&(u+="")){if(!(o=d["$"+u]))throw new Error("missing: "+u);if(o===D)throw new Error("ambiguous: "+u);o.children?o.children.push(s):o.children=[s],s.parent=o}else{if(a)throw new Error("multiple roots");a=s}if(!a)throw new Error("no root");if(a.parent=z,a.eachBefore((function(t){t.depth=t.parent.depth+1,--h})).eachBefore(l),a.parent=null,h>0)throw new Error("cycle");return a}return r.id=function(e){return arguments.length?(t=A(e),r):t},r.parentId=function(t){return arguments.length?(e=A(t),r):e},r},t.tree=function(){var t=B,e=1,r=1,n=null;function i(i){var l=function(t){for(var e,r,n,i,a,o=new H(t,0),s=[o];e=s.pop();)if(n=e._.children)for(e.children=new Array(a=n.length),i=a-1;i>=0;--i)s.push(r=e.children[i]=new H(n[i],i)),r.parent=e;return(o.parent=new H(null,0)).children=[o],o}(i);if(l.eachAfter(a),l.parent.m=-l.z,l.eachBefore(o),n)i.eachBefore(s);else{var c=i,u=i,f=i;i.eachBefore((function(t){t.x<c.x&&(c=t),t.x>u.x&&(u=t),t.depth>f.depth&&(f=t)}));var h=c===u?1:t(c,u)/2,p=h-c.x,d=e/(u.x+h+p),m=r/(f.depth||1);i.eachBefore((function(t){t.x=(t.x+p)*d,t.y=t.depth*m}))}return i}function a(e){var r=e.children,n=e.parent.children,i=e.i?n[e.i-1]:null;if(r){!function(t){for(var e,r=0,n=0,i=t.children,a=i.length;--a>=0;)(e=i[a]).z+=r,e.m+=r,r+=e.s+(n+=e.c)}(e);var a=(r[0].z+r[r.length-1].z)/2;i?(e.z=i.z+t(e._,i._),e.m=e.z-a):e.z=a}else i&&(e.z=i.z+t(e._,i._));e.parent.A=function(e,r,n){if(r){for(var i,a=e,o=e,s=r,l=a.parent.children[0],c=a.m,u=o.m,f=s.m,h=l.m;s=j(s),a=N(a),s&&a;)l=N(l),(o=j(o)).a=e,(i=s.z+f-a.z-c+t(s._,a._))>0&&(U(V(s,e,n),e,i),c+=i,u+=i),f+=s.m,c+=a.m,h+=l.m,u+=o.m;s&&!j(o)&&(o.t=s,o.m+=f-u),a&&!N(l)&&(l.t=a,l.m+=c-h,n=e)}return n}(e,i,e.parent.A||n[0])}function o(t){t._.x=t.z+t.parent.m,t.m+=t.parent.m}function s(t){t.x*=e,t.y=t.depth*r}return i.separation=function(e){return arguments.length?(t=e,i):t},i.size=function(t){return arguments.length?(n=!1,e=+t[0],r=+t[1],i):n?null:[e,r]},i.nodeSize=function(t){return arguments.length?(n=!0,e=+t[0],r=+t[1],i):n?[e,r]:null},i},t.treemap=function(){var t=W,e=!1,r=1,n=1,i=[0],a=M,o=M,s=M,l=M,c=M;function u(t){return t.x0=t.y0=0,t.x1=r,t.y1=n,t.eachBefore(f),i=[0],e&&t.eachBefore(I),t}function f(e){var r=i[e.depth],n=e.x0+r,u=e.y0+r,f=e.x1-r,h=e.y1-r;f<n&&(n=f=(n+f)/2),h<u&&(u=h=(u+h)/2),e.x0=n,e.y0=u,e.x1=f,e.y1=h,e.children&&(r=i[e.depth+1]=a(e)/2,n+=c(e)-r,u+=o(e)-r,(f-=s(e)-r)<n&&(n=f=(n+f)/2),(h-=l(e)-r)<u&&(u=h=(u+h)/2),t(e,n,u,f,h))}return u.round=function(t){return arguments.length?(e=!!t,u):e},u.size=function(t){return arguments.length?(r=+t[0],n=+t[1],u):[r,n]},u.tile=function(e){return arguments.length?(t=A(e),u):t},u.padding=function(t){return arguments.length?u.paddingInner(t).paddingOuter(t):u.paddingInner()},u.paddingInner=function(t){return arguments.length?(a="function"==typeof t?t:S(+t),u):a},u.paddingOuter=function(t){return arguments.length?u.paddingTop(t).paddingRight(t).paddingBottom(t).paddingLeft(t):u.paddingTop()},u.paddingTop=function(t){return arguments.length?(o="function"==typeof t?t:S(+t),u):o},u.paddingRight=function(t){return arguments.length?(s="function"==typeof t?t:S(+t),u):s},u.paddingBottom=function(t){return arguments.length?(l="function"==typeof t?t:S(+t),u):l},u.paddingLeft=function(t){return arguments.length?(c="function"==typeof t?t:S(+t),u):c},u},t.treemapBinary=function(t,e,r,n,i){var a,o,s=t.children,l=s.length,c=new Array(l+1);for(c[0]=o=a=0;a<l;++a)c[a+1]=o+=s[a].value;!function t(e,r,n,i,a,o,l){if(e>=r-1){var u=s[e];return u.x0=i,u.y0=a,u.x1=o,void(u.y1=l)}var f=c[e],h=n/2+f,p=e+1,d=r-1;for(;p<d;){var m=p+d>>>1;c[m]<h?p=m+1:d=m}h-c[p-1]<c[p]-h&&e+1<p&&--p;var g=c[p]-f,v=n-g;if(o-i>l-a){var y=(i*v+o*g)/n;t(e,p,g,i,a,y,l),t(p,r,v,y,a,o,l)}else{var x=(a*v+l*g)/n;t(e,p,g,i,a,o,x),t(p,r,v,i,x,o,l)}}(0,l,t.value,e,r,n,i)},t.treemapDice=O,t.treemapResquarify=X,t.treemapSlice=q,t.treemapSliceDice=function(t,e,r,n,i){(1&t.depth?q:O)(t,e,r,n,i)},t.treemapSquarify=W,Object.defineProperty(t,"__esModule",{value:!0})}))},{}],111:[function(t,e,r){!function(n,i){"object"==typeof r&&void 0!==e?i(r,t("d3-color")):i((n=n||self).d3=n.d3||{},n.d3)}(this,(function(t,e){"use strict";function r(t,e,r,n,i){var a=t*t,o=a*t;return((1-3*t+3*a-o)*e+(4-6*a+3*o)*r+(1+3*t+3*a-3*o)*n+o*i)/6}function n(t){var e=t.length-1;return function(n){var i=n<=0?n=0:n>=1?(n=1,e-1):Math.floor(n*e),a=t[i],o=t[i+1],s=i>0?t[i-1]:2*a-o,l=i<e-1?t[i+2]:2*o-a;return r((n-i/e)*e,s,a,o,l)}}function i(t){var e=t.length;return function(n){var i=Math.floor(((n%=1)<0?++n:n)*e),a=t[(i+e-1)%e],o=t[i%e],s=t[(i+1)%e],l=t[(i+2)%e];return r((n-i/e)*e,a,o,s,l)}}function a(t){return function(){return t}}function o(t,e){return function(r){return t+r*e}}function s(t,e){var r=e-t;return r?o(t,r>180||r<-180?r-360*Math.round(r/360):r):a(isNaN(t)?e:t)}function l(t){return 1==(t=+t)?c:function(e,r){return r-e?function(t,e,r){return t=Math.pow(t,r),e=Math.pow(e,r)-t,r=1/r,function(n){return Math.pow(t+n*e,r)}}(e,r,t):a(isNaN(e)?r:e)}}function c(t,e){var r=e-t;return r?o(t,r):a(isNaN(t)?e:t)}var u=function t(r){var n=l(r);function i(t,r){var i=n((t=e.rgb(t)).r,(r=e.rgb(r)).r),a=n(t.g,r.g),o=n(t.b,r.b),s=c(t.opacity,r.opacity);return function(e){return t.r=i(e),t.g=a(e),t.b=o(e),t.opacity=s(e),t+""}}return i.gamma=t,i}(1);function f(t){return function(r){var n,i,a=r.length,o=new Array(a),s=new Array(a),l=new Array(a);for(n=0;n<a;++n)i=e.rgb(r[n]),o[n]=i.r||0,s[n]=i.g||0,l[n]=i.b||0;return o=t(o),s=t(s),l=t(l),i.opacity=1,function(t){return i.r=o(t),i.g=s(t),i.b=l(t),i+""}}}var h=f(n),p=f(i);function d(t,e){e||(e=[]);var r,n=t?Math.min(e.length,t.length):0,i=e.slice();return function(a){for(r=0;r<n;++r)i[r]=t[r]*(1-a)+e[r]*a;return i}}function m(t){return ArrayBuffer.isView(t)&&!(t instanceof DataView)}function g(t,e){var r,n=e?e.length:0,i=t?Math.min(n,t.length):0,a=new Array(i),o=new Array(n);for(r=0;r<i;++r)a[r]=T(t[r],e[r]);for(;r<n;++r)o[r]=e[r];return function(t){for(r=0;r<i;++r)o[r]=a[r](t);return o}}function v(t,e){var r=new Date;return t=+t,e=+e,function(n){return r.setTime(t*(1-n)+e*n),r}}function y(t,e){return t=+t,e=+e,function(r){return t*(1-r)+e*r}}function x(t,e){var r,n={},i={};for(r in null!==t&&"object"==typeof t||(t={}),null!==e&&"object"==typeof e||(e={}),e)r in t?n[r]=T(t[r],e[r]):i[r]=e[r];return function(t){for(r in n)i[r]=n[r](t);return i}}var b=/[-+]?(?:\d+\.?\d*|\.?\d+)(?:[eE][-+]?\d+)?/g,_=new RegExp(b.source,"g");function w(t,e){var r,n,i,a=b.lastIndex=_.lastIndex=0,o=-1,s=[],l=[];for(t+="",e+="";(r=b.exec(t))&&(n=_.exec(e));)(i=n.index)>a&&(i=e.slice(a,i),s[o]?s[o]+=i:s[++o]=i),(r=r[0])===(n=n[0])?s[o]?s[o]+=n:s[++o]=n:(s[++o]=null,l.push({i:o,x:y(r,n)})),a=_.lastIndex;return a<e.length&&(i=e.slice(a),s[o]?s[o]+=i:s[++o]=i),s.length<2?l[0]?function(t){return function(e){return t(e)+""}}(l[0].x):function(t){return function(){return t}}(e):(e=l.length,function(t){for(var r,n=0;n<e;++n)s[(r=l[n]).i]=r.x(t);return s.join("")})}function T(t,r){var n,i=typeof r;return null==r||"boolean"===i?a(r):("number"===i?y:"string"===i?(n=e.color(r))?(r=n,u):w:r instanceof e.color?u:r instanceof Date?v:m(r)?d:Array.isArray(r)?g:"function"!=typeof r.valueOf&&"function"!=typeof r.toString||isNaN(r)?x:y)(t,r)}var k,A,M,S,E=180/Math.PI,L={translateX:0,translateY:0,rotate:0,skewX:0,scaleX:1,scaleY:1};function C(t,e,r,n,i,a){var o,s,l;return(o=Math.sqrt(t*t+e*e))&&(t/=o,e/=o),(l=t*r+e*n)&&(r-=t*l,n-=e*l),(s=Math.sqrt(r*r+n*n))&&(r/=s,n/=s,l/=s),t*n<e*r&&(t=-t,e=-e,l=-l,o=-o),{translateX:i,translateY:a,rotate:Math.atan2(e,t)*E,skewX:Math.atan(l)*E,scaleX:o,scaleY:s}}function P(t,e,r,n){function i(t){return t.length?t.pop()+" ":""}return function(a,o){var s=[],l=[];return a=t(a),o=t(o),function(t,n,i,a,o,s){if(t!==i||n!==a){var l=o.push("translate(",null,e,null,r);s.push({i:l-4,x:y(t,i)},{i:l-2,x:y(n,a)})}else(i||a)&&o.push("translate("+i+e+a+r)}(a.translateX,a.translateY,o.translateX,o.translateY,s,l),function(t,e,r,a){t!==e?(t-e>180?e+=360:e-t>180&&(t+=360),a.push({i:r.push(i(r)+"rotate(",null,n)-2,x:y(t,e)})):e&&r.push(i(r)+"rotate("+e+n)}(a.rotate,o.rotate,s,l),function(t,e,r,a){t!==e?a.push({i:r.push(i(r)+"skewX(",null,n)-2,x:y(t,e)}):e&&r.push(i(r)+"skewX("+e+n)}(a.skewX,o.skewX,s,l),function(t,e,r,n,a,o){if(t!==r||e!==n){var s=a.push(i(a)+"scale(",null,",",null,")");o.push({i:s-4,x:y(t,r)},{i:s-2,x:y(e,n)})}else 1===r&&1===n||a.push(i(a)+"scale("+r+","+n+")")}(a.scaleX,a.scaleY,o.scaleX,o.scaleY,s,l),a=o=null,function(t){for(var e,r=-1,n=l.length;++r<n;)s[(e=l[r]).i]=e.x(t);return s.join("")}}}var I=P((function(t){return"none"===t?L:(k||(k=document.createElement("DIV"),A=document.documentElement,M=document.defaultView),k.style.transform=t,t=M.getComputedStyle(A.appendChild(k),null).getPropertyValue("transform"),A.removeChild(k),C(+(t=t.slice(7,-1).split(","))[0],+t[1],+t[2],+t[3],+t[4],+t[5]))}),"px, ","px)","deg)"),O=P((function(t){return null==t?L:(S||(S=document.createElementNS("http://www.w3.org/2000/svg","g")),S.setAttribute("transform",t),(t=S.transform.baseVal.consolidate())?C((t=t.matrix).a,t.b,t.c,t.d,t.e,t.f):L)}),", ",")",")"),z=Math.SQRT2;function D(t){return((t=Math.exp(t))+1/t)/2}function R(t){return function(r,n){var i=t((r=e.hsl(r)).h,(n=e.hsl(n)).h),a=c(r.s,n.s),o=c(r.l,n.l),s=c(r.opacity,n.opacity);return function(t){return r.h=i(t),r.s=a(t),r.l=o(t),r.opacity=s(t),r+""}}}var F=R(s),B=R(c);function N(t){return function(r,n){var i=t((r=e.hcl(r)).h,(n=e.hcl(n)).h),a=c(r.c,n.c),o=c(r.l,n.l),s=c(r.opacity,n.opacity);return function(t){return r.h=i(t),r.c=a(t),r.l=o(t),r.opacity=s(t),r+""}}}var j=N(s),U=N(c);function V(t){return function r(n){function i(r,i){var a=t((r=e.cubehelix(r)).h,(i=e.cubehelix(i)).h),o=c(r.s,i.s),s=c(r.l,i.l),l=c(r.opacity,i.opacity);return function(t){return r.h=a(t),r.s=o(t),r.l=s(Math.pow(t,n)),r.opacity=l(t),r+""}}return n=+n,i.gamma=r,i}(1)}var H=V(s),q=V(c);t.interpolate=T,t.interpolateArray=function(t,e){return(m(e)?d:g)(t,e)},t.interpolateBasis=n,t.interpolateBasisClosed=i,t.interpolateCubehelix=H,t.interpolateCubehelixLong=q,t.interpolateDate=v,t.interpolateDiscrete=function(t){var e=t.length;return function(r){return t[Math.max(0,Math.min(e-1,Math.floor(r*e)))]}},t.interpolateHcl=j,t.interpolateHclLong=U,t.interpolateHsl=F,t.interpolateHslLong=B,t.interpolateHue=function(t,e){var r=s(+t,+e);return function(t){var e=r(t);return e-360*Math.floor(e/360)}},t.interpolateLab=function(t,r){var n=c((t=e.lab(t)).l,(r=e.lab(r)).l),i=c(t.a,r.a),a=c(t.b,r.b),o=c(t.opacity,r.opacity);return function(e){return t.l=n(e),t.a=i(e),t.b=a(e),t.opacity=o(e),t+""}},t.interpolateNumber=y,t.interpolateNumberArray=d,t.interpolateObject=x,t.interpolateRgb=u,t.interpolateRgbBasis=h,t.interpolateRgbBasisClosed=p,t.interpolateRound=function(t,e){return t=+t,e=+e,function(r){return Math.round(t*(1-r)+e*r)}},t.interpolateString=w,t.interpolateTransformCss=I,t.interpolateTransformSvg=O,t.interpolateZoom=function(t,e){var r,n,i=t[0],a=t[1],o=t[2],s=e[0],l=e[1],c=e[2],u=s-i,f=l-a,h=u*u+f*f;if(h<1e-12)n=Math.log(c/o)/z,r=function(t){return[i+t*u,a+t*f,o*Math.exp(z*t*n)]};else{var p=Math.sqrt(h),d=(c*c-o*o+4*h)/(2*o*2*p),m=(c*c-o*o-4*h)/(2*c*2*p),g=Math.log(Math.sqrt(d*d+1)-d),v=Math.log(Math.sqrt(m*m+1)-m);n=(v-g)/z,r=function(t){var e,r=t*n,s=D(g),l=o/(2*p)*(s*(e=z*r+g,((e=Math.exp(2*e))-1)/(e+1))-function(t){return((t=Math.exp(t))-1/t)/2}(g));return[i+l*u,a+l*f,o*s/D(z*r+g)]}}return r.duration=1e3*n,r},t.piecewise=function(t,e){for(var r=0,n=e.length-1,i=e[0],a=new Array(n<0?0:n);r<n;)a[r]=t(i,i=e[++r]);return function(t){var e=Math.max(0,Math.min(n-1,Math.floor(t*=n)));return a[e](t-e)}},t.quantize=function(t,e){for(var r=new Array(e),n=0;n<e;++n)r[n]=t(n/(e-1));return r},Object.defineProperty(t,"__esModule",{value:!0})}))},{"d3-color":104}],112:[function(t,e,r){!function(t,n){"object"==typeof r&&void 0!==e?n(r):n((t=t||self).d3=t.d3||{})}(this,(function(t){"use strict";var e=Math.PI,r=2*e,n=r-1e-6;function i(){this._x0=this._y0=this._x1=this._y1=null,this._=""}function a(){return new i}i.prototype=a.prototype={constructor:i,moveTo:function(t,e){this._+="M"+(this._x0=this._x1=+t)+","+(this._y0=this._y1=+e)},closePath:function(){null!==this._x1&&(this._x1=this._x0,this._y1=this._y0,this._+="Z")},lineTo:function(t,e){this._+="L"+(this._x1=+t)+","+(this._y1=+e)},quadraticCurveTo:function(t,e,r,n){this._+="Q"+ +t+","+ +e+","+(this._x1=+r)+","+(this._y1=+n)},bezierCurveTo:function(t,e,r,n,i,a){this._+="C"+ +t+","+ +e+","+ +r+","+ +n+","+(this._x1=+i)+","+(this._y1=+a)},arcTo:function(t,r,n,i,a){t=+t,r=+r,n=+n,i=+i,a=+a;var o=this._x1,s=this._y1,l=n-t,c=i-r,u=o-t,f=s-r,h=u*u+f*f;if(a<0)throw new Error("negative radius: "+a);if(null===this._x1)this._+="M"+(this._x1=t)+","+(this._y1=r);else if(h>1e-6)if(Math.abs(f*l-c*u)>1e-6&&a){var p=n-o,d=i-s,m=l*l+c*c,g=p*p+d*d,v=Math.sqrt(m),y=Math.sqrt(h),x=a*Math.tan((e-Math.acos((m+h-g)/(2*v*y)))/2),b=x/y,_=x/v;Math.abs(b-1)>1e-6&&(this._+="L"+(t+b*u)+","+(r+b*f)),this._+="A"+a+","+a+",0,0,"+ +(f*p>u*d)+","+(this._x1=t+_*l)+","+(this._y1=r+_*c)}else this._+="L"+(this._x1=t)+","+(this._y1=r);else;},arc:function(t,i,a,o,s,l){t=+t,i=+i,l=!!l;var c=(a=+a)*Math.cos(o),u=a*Math.sin(o),f=t+c,h=i+u,p=1^l,d=l?o-s:s-o;if(a<0)throw new Error("negative radius: "+a);null===this._x1?this._+="M"+f+","+h:(Math.abs(this._x1-f)>1e-6||Math.abs(this._y1-h)>1e-6)&&(this._+="L"+f+","+h),a&&(d<0&&(d=d%r+r),d>n?this._+="A"+a+","+a+",0,1,"+p+","+(t-c)+","+(i-u)+"A"+a+","+a+",0,1,"+p+","+(this._x1=f)+","+(this._y1=h):d>1e-6&&(this._+="A"+a+","+a+",0,"+ +(d>=e)+","+p+","+(this._x1=t+a*Math.cos(s))+","+(this._y1=i+a*Math.sin(s))))},rect:function(t,e,r,n){this._+="M"+(this._x0=this._x1=+t)+","+(this._y0=this._y1=+e)+"h"+ +r+"v"+ +n+"h"+-r+"Z"},toString:function(){return this._}},t.path=a,Object.defineProperty(t,"__esModule",{value:!0})}))},{}],113:[function(t,e,r){!function(t,n){"object"==typeof r&&void 0!==e?n(r):n((t=t||self).d3=t.d3||{})}(this,(function(t){"use strict";function e(t,e,r,n){if(isNaN(e)||isNaN(r))return t;var i,a,o,s,l,c,u,f,h,p=t._root,d={data:n},m=t._x0,g=t._y0,v=t._x1,y=t._y1;if(!p)return t._root=d,t;for(;p.length;)if((c=e>=(a=(m+v)/2))?m=a:v=a,(u=r>=(o=(g+y)/2))?g=o:y=o,i=p,!(p=p[f=u<<1|c]))return i[f]=d,t;if(s=+t._x.call(null,p.data),l=+t._y.call(null,p.data),e===s&&r===l)return d.next=p,i?i[f]=d:t._root=d,t;do{i=i?i[f]=new Array(4):t._root=new Array(4),(c=e>=(a=(m+v)/2))?m=a:v=a,(u=r>=(o=(g+y)/2))?g=o:y=o}while((f=u<<1|c)==(h=(l>=o)<<1|s>=a));return i[h]=p,i[f]=d,t}function r(t,e,r,n,i){this.node=t,this.x0=e,this.y0=r,this.x1=n,this.y1=i}function n(t){return t[0]}function i(t){return t[1]}function a(t,e,r){var a=new o(null==e?n:e,null==r?i:r,NaN,NaN,NaN,NaN);return null==t?a:a.addAll(t)}function o(t,e,r,n,i,a){this._x=t,this._y=e,this._x0=r,this._y0=n,this._x1=i,this._y1=a,this._root=void 0}function s(t){for(var e={data:t.data},r=e;t=t.next;)r=r.next={data:t.data};return e}var l=a.prototype=o.prototype;l.copy=function(){var t,e,r=new o(this._x,this._y,this._x0,this._y0,this._x1,this._y1),n=this._root;if(!n)return r;if(!n.length)return r._root=s(n),r;for(t=[{source:n,target:r._root=new Array(4)}];n=t.pop();)for(var i=0;i<4;++i)(e=n.source[i])&&(e.length?t.push({source:e,target:n.target[i]=new Array(4)}):n.target[i]=s(e));return r},l.add=function(t){var r=+this._x.call(null,t),n=+this._y.call(null,t);return e(this.cover(r,n),r,n,t)},l.addAll=function(t){var r,n,i,a,o=t.length,s=new Array(o),l=new Array(o),c=1/0,u=1/0,f=-1/0,h=-1/0;for(n=0;n<o;++n)isNaN(i=+this._x.call(null,r=t[n]))||isNaN(a=+this._y.call(null,r))||(s[n]=i,l[n]=a,i<c&&(c=i),i>f&&(f=i),a<u&&(u=a),a>h&&(h=a));if(c>f||u>h)return this;for(this.cover(c,u).cover(f,h),n=0;n<o;++n)e(this,s[n],l[n],t[n]);return this},l.cover=function(t,e){if(isNaN(t=+t)||isNaN(e=+e))return this;var r=this._x0,n=this._y0,i=this._x1,a=this._y1;if(isNaN(r))i=(r=Math.floor(t))+1,a=(n=Math.floor(e))+1;else{for(var o,s,l=i-r,c=this._root;r>t||t>=i||n>e||e>=a;)switch(s=(e<n)<<1|t<r,(o=new Array(4))[s]=c,c=o,l*=2,s){case 0:i=r+l,a=n+l;break;case 1:r=i-l,a=n+l;break;case 2:i=r+l,n=a-l;break;case 3:r=i-l,n=a-l}this._root&&this._root.length&&(this._root=c)}return this._x0=r,this._y0=n,this._x1=i,this._y1=a,this},l.data=function(){var t=[];return this.visit((function(e){if(!e.length)do{t.push(e.data)}while(e=e.next)})),t},l.extent=function(t){return arguments.length?this.cover(+t[0][0],+t[0][1]).cover(+t[1][0],+t[1][1]):isNaN(this._x0)?void 0:[[this._x0,this._y0],[this._x1,this._y1]]},l.find=function(t,e,n){var i,a,o,s,l,c,u,f=this._x0,h=this._y0,p=this._x1,d=this._y1,m=[],g=this._root;for(g&&m.push(new r(g,f,h,p,d)),null==n?n=1/0:(f=t-n,h=e-n,p=t+n,d=e+n,n*=n);c=m.pop();)if(!(!(g=c.node)||(a=c.x0)>p||(o=c.y0)>d||(s=c.x1)<f||(l=c.y1)<h))if(g.length){var v=(a+s)/2,y=(o+l)/2;m.push(new r(g[3],v,y,s,l),new r(g[2],a,y,v,l),new r(g[1],v,o,s,y),new r(g[0],a,o,v,y)),(u=(e>=y)<<1|t>=v)&&(c=m[m.length-1],m[m.length-1]=m[m.length-1-u],m[m.length-1-u]=c)}else{var x=t-+this._x.call(null,g.data),b=e-+this._y.call(null,g.data),_=x*x+b*b;if(_<n){var w=Math.sqrt(n=_);f=t-w,h=e-w,p=t+w,d=e+w,i=g.data}}return i},l.remove=function(t){if(isNaN(a=+this._x.call(null,t))||isNaN(o=+this._y.call(null,t)))return this;var e,r,n,i,a,o,s,l,c,u,f,h,p=this._root,d=this._x0,m=this._y0,g=this._x1,v=this._y1;if(!p)return this;if(p.length)for(;;){if((c=a>=(s=(d+g)/2))?d=s:g=s,(u=o>=(l=(m+v)/2))?m=l:v=l,e=p,!(p=p[f=u<<1|c]))return this;if(!p.length)break;(e[f+1&3]||e[f+2&3]||e[f+3&3])&&(r=e,h=f)}for(;p.data!==t;)if(n=p,!(p=p.next))return this;return(i=p.next)&&delete p.next,n?(i?n.next=i:delete n.next,this):e?(i?e[f]=i:delete e[f],(p=e[0]||e[1]||e[2]||e[3])&&p===(e[3]||e[2]||e[1]||e[0])&&!p.length&&(r?r[h]=p:this._root=p),this):(this._root=i,this)},l.removeAll=function(t){for(var e=0,r=t.length;e<r;++e)this.remove(t[e]);return this},l.root=function(){return this._root},l.size=function(){var t=0;return this.visit((function(e){if(!e.length)do{++t}while(e=e.next)})),t},l.visit=function(t){var e,n,i,a,o,s,l=[],c=this._root;for(c&&l.push(new r(c,this._x0,this._y0,this._x1,this._y1));e=l.pop();)if(!t(c=e.node,i=e.x0,a=e.y0,o=e.x1,s=e.y1)&&c.length){var u=(i+o)/2,f=(a+s)/2;(n=c[3])&&l.push(new r(n,u,f,o,s)),(n=c[2])&&l.push(new r(n,i,f,u,s)),(n=c[1])&&l.push(new r(n,u,a,o,f)),(n=c[0])&&l.push(new r(n,i,a,u,f))}return this},l.visitAfter=function(t){var e,n=[],i=[];for(this._root&&n.push(new r(this._root,this._x0,this._y0,this._x1,this._y1));e=n.pop();){var a=e.node;if(a.length){var o,s=e.x0,l=e.y0,c=e.x1,u=e.y1,f=(s+c)/2,h=(l+u)/2;(o=a[0])&&n.push(new r(o,s,l,f,h)),(o=a[1])&&n.push(new r(o,f,l,c,h)),(o=a[2])&&n.push(new r(o,s,h,f,u)),(o=a[3])&&n.push(new r(o,f,h,c,u))}i.push(e)}for(;e=i.pop();)t(e.node,e.x0,e.y0,e.x1,e.y1);return this},l.x=function(t){return arguments.length?(this._x=t,this):this._x},l.y=function(t){return arguments.length?(this._y=t,this):this._y},t.quadtree=a,Object.defineProperty(t,"__esModule",{value:!0})}))},{}],114:[function(t,e,r){!function(n,i){"object"==typeof r&&void 0!==e?i(r,t("d3-path")):i((n=n||self).d3=n.d3||{},n.d3)}(this,(function(t,e){"use strict";function r(t){return function(){return t}}var n=Math.abs,i=Math.atan2,a=Math.cos,o=Math.max,s=Math.min,l=Math.sin,c=Math.sqrt,u=Math.PI,f=u/2,h=2*u;function p(t){return t>1?0:t<-1?u:Math.acos(t)}function d(t){return t>=1?f:t<=-1?-f:Math.asin(t)}function m(t){return t.innerRadius}function g(t){return t.outerRadius}function v(t){return t.startAngle}function y(t){return t.endAngle}function x(t){return t&&t.padAngle}function b(t,e,r,n,i,a,o,s){var l=r-t,c=n-e,u=o-i,f=s-a,h=f*l-u*c;if(!(h*h<1e-12))return[t+(h=(u*(e-a)-f*(t-i))/h)*l,e+h*c]}function _(t,e,r,n,i,a,s){var l=t-r,u=e-n,f=(s?a:-a)/c(l*l+u*u),h=f*u,p=-f*l,d=t+h,m=e+p,g=r+h,v=n+p,y=(d+g)/2,x=(m+v)/2,b=g-d,_=v-m,w=b*b+_*_,T=i-a,k=d*v-g*m,A=(_<0?-1:1)*c(o(0,T*T*w-k*k)),M=(k*_-b*A)/w,S=(-k*b-_*A)/w,E=(k*_+b*A)/w,L=(-k*b+_*A)/w,C=M-y,P=S-x,I=E-y,O=L-x;return C*C+P*P>I*I+O*O&&(M=E,S=L),{cx:M,cy:S,x01:-h,y01:-p,x11:M*(i/T-1),y11:S*(i/T-1)}}function w(t){this._context=t}function T(t){return new w(t)}function k(t){return t[0]}function A(t){return t[1]}function M(){var t=k,n=A,i=r(!0),a=null,o=T,s=null;function l(r){var l,c,u,f=r.length,h=!1;for(null==a&&(s=o(u=e.path())),l=0;l<=f;++l)!(l<f&&i(c=r[l],l,r))===h&&((h=!h)?s.lineStart():s.lineEnd()),h&&s.point(+t(c,l,r),+n(c,l,r));if(u)return s=null,u+""||null}return l.x=function(e){return arguments.length?(t="function"==typeof e?e:r(+e),l):t},l.y=function(t){return arguments.length?(n="function"==typeof t?t:r(+t),l):n},l.defined=function(t){return arguments.length?(i="function"==typeof t?t:r(!!t),l):i},l.curve=function(t){return arguments.length?(o=t,null!=a&&(s=o(a)),l):o},l.context=function(t){return arguments.length?(null==t?a=s=null:s=o(a=t),l):a},l}function S(){var t=k,n=null,i=r(0),a=A,o=r(!0),s=null,l=T,c=null;function u(r){var u,f,h,p,d,m=r.length,g=!1,v=new Array(m),y=new Array(m);for(null==s&&(c=l(d=e.path())),u=0;u<=m;++u){if(!(u<m&&o(p=r[u],u,r))===g)if(g=!g)f=u,c.areaStart(),c.lineStart();else{for(c.lineEnd(),c.lineStart(),h=u-1;h>=f;--h)c.point(v[h],y[h]);c.lineEnd(),c.areaEnd()}g&&(v[u]=+t(p,u,r),y[u]=+i(p,u,r),c.point(n?+n(p,u,r):v[u],a?+a(p,u,r):y[u]))}if(d)return c=null,d+""||null}function f(){return M().defined(o).curve(l).context(s)}return u.x=function(e){return arguments.length?(t="function"==typeof e?e:r(+e),n=null,u):t},u.x0=function(e){return arguments.length?(t="function"==typeof e?e:r(+e),u):t},u.x1=function(t){return arguments.length?(n=null==t?null:"function"==typeof t?t:r(+t),u):n},u.y=function(t){return arguments.length?(i="function"==typeof t?t:r(+t),a=null,u):i},u.y0=function(t){return arguments.length?(i="function"==typeof t?t:r(+t),u):i},u.y1=function(t){return arguments.length?(a=null==t?null:"function"==typeof t?t:r(+t),u):a},u.lineX0=u.lineY0=function(){return f().x(t).y(i)},u.lineY1=function(){return f().x(t).y(a)},u.lineX1=function(){return f().x(n).y(i)},u.defined=function(t){return arguments.length?(o="function"==typeof t?t:r(!!t),u):o},u.curve=function(t){return arguments.length?(l=t,null!=s&&(c=l(s)),u):l},u.context=function(t){return arguments.length?(null==t?s=c=null:c=l(s=t),u):s},u}function E(t,e){return e<t?-1:e>t?1:e>=t?0:NaN}function L(t){return t}w.prototype={areaStart:function(){this._line=0},areaEnd:function(){this._line=NaN},lineStart:function(){this._point=0},lineEnd:function(){(this._line||0!==this._line&&1===this._point)&&this._context.closePath(),this._line=1-this._line},point:function(t,e){switch(t=+t,e=+e,this._point){case 0:this._point=1,this._line?this._context.lineTo(t,e):this._context.moveTo(t,e);break;case 1:this._point=2;default:this._context.lineTo(t,e)}}};var C=I(T);function P(t){this._curve=t}function I(t){function e(e){return new P(t(e))}return e._curve=t,e}function O(t){var e=t.curve;return t.angle=t.x,delete t.x,t.radius=t.y,delete t.y,t.curve=function(t){return arguments.length?e(I(t)):e()._curve},t}function z(){return O(M().curve(C))}function D(){var t=S().curve(C),e=t.curve,r=t.lineX0,n=t.lineX1,i=t.lineY0,a=t.lineY1;return t.angle=t.x,delete t.x,t.startAngle=t.x0,delete t.x0,t.endAngle=t.x1,delete t.x1,t.radius=t.y,delete t.y,t.innerRadius=t.y0,delete t.y0,t.outerRadius=t.y1,delete t.y1,t.lineStartAngle=function(){return O(r())},delete t.lineX0,t.lineEndAngle=function(){return O(n())},delete t.lineX1,t.lineInnerRadius=function(){return O(i())},delete t.lineY0,t.lineOuterRadius=function(){return O(a())},delete t.lineY1,t.curve=function(t){return arguments.length?e(I(t)):e()._curve},t}function R(t,e){return[(e=+e)*Math.cos(t-=Math.PI/2),e*Math.sin(t)]}P.prototype={areaStart:function(){this._curve.areaStart()},areaEnd:function(){this._curve.areaEnd()},lineStart:function(){this._curve.lineStart()},lineEnd:function(){this._curve.lineEnd()},point:function(t,e){this._curve.point(e*Math.sin(t),e*-Math.cos(t))}};var F=Array.prototype.slice;function B(t){return t.source}function N(t){return t.target}function j(t){var n=B,i=N,a=k,o=A,s=null;function l(){var r,l=F.call(arguments),c=n.apply(this,l),u=i.apply(this,l);if(s||(s=r=e.path()),t(s,+a.apply(this,(l[0]=c,l)),+o.apply(this,l),+a.apply(this,(l[0]=u,l)),+o.apply(this,l)),r)return s=null,r+""||null}return l.source=function(t){return arguments.length?(n=t,l):n},l.target=function(t){return arguments.length?(i=t,l):i},l.x=function(t){return arguments.length?(a="function"==typeof t?t:r(+t),l):a},l.y=function(t){return arguments.length?(o="function"==typeof t?t:r(+t),l):o},l.context=function(t){return arguments.length?(s=null==t?null:t,l):s},l}function U(t,e,r,n,i){t.moveTo(e,r),t.bezierCurveTo(e=(e+n)/2,r,e,i,n,i)}function V(t,e,r,n,i){t.moveTo(e,r),t.bezierCurveTo(e,r=(r+i)/2,n,r,n,i)}function H(t,e,r,n,i){var a=R(e,r),o=R(e,r=(r+i)/2),s=R(n,r),l=R(n,i);t.moveTo(a[0],a[1]),t.bezierCurveTo(o[0],o[1],s[0],s[1],l[0],l[1])}var q={draw:function(t,e){var r=Math.sqrt(e/u);t.moveTo(r,0),t.arc(0,0,r,0,h)}},G={draw:function(t,e){var r=Math.sqrt(e/5)/2;t.moveTo(-3*r,-r),t.lineTo(-r,-r),t.lineTo(-r,-3*r),t.lineTo(r,-3*r),t.lineTo(r,-r),t.lineTo(3*r,-r),t.lineTo(3*r,r),t.lineTo(r,r),t.lineTo(r,3*r),t.lineTo(-r,3*r),t.lineTo(-r,r),t.lineTo(-3*r,r),t.closePath()}},Y=Math.sqrt(1/3),W=2*Y,X={draw:function(t,e){var r=Math.sqrt(e/W),n=r*Y;t.moveTo(0,-r),t.lineTo(n,0),t.lineTo(0,r),t.lineTo(-n,0),t.closePath()}},Z=Math.sin(u/10)/Math.sin(7*u/10),J=Math.sin(h/10)*Z,K=-Math.cos(h/10)*Z,Q={draw:function(t,e){var r=Math.sqrt(.8908130915292852*e),n=J*r,i=K*r;t.moveTo(0,-r),t.lineTo(n,i);for(var a=1;a<5;++a){var o=h*a/5,s=Math.cos(o),l=Math.sin(o);t.lineTo(l*r,-s*r),t.lineTo(s*n-l*i,l*n+s*i)}t.closePath()}},$={draw:function(t,e){var r=Math.sqrt(e),n=-r/2;t.rect(n,n,r,r)}},tt=Math.sqrt(3),et={draw:function(t,e){var r=-Math.sqrt(e/(3*tt));t.moveTo(0,2*r),t.lineTo(-tt*r,-r),t.lineTo(tt*r,-r),t.closePath()}},rt=-.5,nt=Math.sqrt(3)/2,it=1/Math.sqrt(12),at=3*(it/2+1),ot={draw:function(t,e){var r=Math.sqrt(e/at),n=r/2,i=r*it,a=n,o=r*it+r,s=-a,l=o;t.moveTo(n,i),t.lineTo(a,o),t.lineTo(s,l),t.lineTo(rt*n-nt*i,nt*n+rt*i),t.lineTo(rt*a-nt*o,nt*a+rt*o),t.lineTo(rt*s-nt*l,nt*s+rt*l),t.lineTo(rt*n+nt*i,rt*i-nt*n),t.lineTo(rt*a+nt*o,rt*o-nt*a),t.lineTo(rt*s+nt*l,rt*l-nt*s),t.closePath()}},st=[q,G,X,$,Q,et,ot];function lt(){}function ct(t,e,r){t._context.bezierCurveTo((2*t._x0+t._x1)/3,(2*t._y0+t._y1)/3,(t._x0+2*t._x1)/3,(t._y0+2*t._y1)/3,(t._x0+4*t._x1+e)/6,(t._y0+4*t._y1+r)/6)}function ut(t){this._context=t}function ft(t){this._context=t}function ht(t){this._context=t}function pt(t,e){this._basis=new ut(t),this._beta=e}ut.prototype={areaStart:function(){this._line=0},areaEnd:function(){this._line=NaN},lineStart:function(){this._x0=this._x1=this._y0=this._y1=NaN,this._point=0},lineEnd:function(){switch(this._point){case 3:ct(this,this._x1,this._y1);case 2:this._context.lineTo(this._x1,this._y1)}(this._line||0!==this._line&&1===this._point)&&this._context.closePath(),this._line=1-this._line},point:function(t,e){switch(t=+t,e=+e,this._point){case 0:this._point=1,this._line?this._context.lineTo(t,e):this._context.moveTo(t,e);break;case 1:this._point=2;break;case 2:this._point=3,this._context.lineTo((5*this._x0+this._x1)/6,(5*this._y0+this._y1)/6);default:ct(this,t,e)}this._x0=this._x1,this._x1=t,this._y0=this._y1,this._y1=e}},ft.prototype={areaStart:lt,areaEnd:lt,lineStart:function(){this._x0=this._x1=this._x2=this._x3=this._x4=this._y0=this._y1=this._y2=this._y3=this._y4=NaN,this._point=0},lineEnd:function(){switch(this._point){case 1:this._context.moveTo(this._x2,this._y2),this._context.closePath();break;case 2:this._context.moveTo((this._x2+2*this._x3)/3,(this._y2+2*this._y3)/3),this._context.lineTo((this._x3+2*this._x2)/3,(this._y3+2*this._y2)/3),this._context.closePath();break;case 3:this.point(this._x2,this._y2),this.point(this._x3,this._y3),this.point(this._x4,this._y4)}},point:function(t,e){switch(t=+t,e=+e,this._point){case 0:this._point=1,this._x2=t,this._y2=e;break;case 1:this._point=2,this._x3=t,this._y3=e;break;case 2:this._point=3,this._x4=t,this._y4=e,this._context.moveTo((this._x0+4*this._x1+t)/6,(this._y0+4*this._y1+e)/6);break;default:ct(this,t,e)}this._x0=this._x1,this._x1=t,this._y0=this._y1,this._y1=e}},ht.prototype={areaStart:function(){this._line=0},areaEnd:function(){this._line=NaN},lineStart:function(){this._x0=this._x1=this._y0=this._y1=NaN,this._point=0},lineEnd:function(){(this._line||0!==this._line&&3===this._point)&&this._context.closePath(),this._line=1-this._line},point:function(t,e){switch(t=+t,e=+e,this._point){case 0:this._point=1;break;case 1:this._point=2;break;case 2:this._point=3;var r=(this._x0+4*this._x1+t)/6,n=(this._y0+4*this._y1+e)/6;this._line?this._context.lineTo(r,n):this._context.moveTo(r,n);break;case 3:this._point=4;default:ct(this,t,e)}this._x0=this._x1,this._x1=t,this._y0=this._y1,this._y1=e}},pt.prototype={lineStart:function(){this._x=[],this._y=[],this._basis.lineStart()},lineEnd:function(){var t=this._x,e=this._y,r=t.length-1;if(r>0)for(var n,i=t[0],a=e[0],o=t[r]-i,s=e[r]-a,l=-1;++l<=r;)n=l/r,this._basis.point(this._beta*t[l]+(1-this._beta)*(i+n*o),this._beta*e[l]+(1-this._beta)*(a+n*s));this._x=this._y=null,this._basis.lineEnd()},point:function(t,e){this._x.push(+t),this._y.push(+e)}};var dt=function t(e){function r(t){return 1===e?new ut(t):new pt(t,e)}return r.beta=function(e){return t(+e)},r}(.85);function mt(t,e,r){t._context.bezierCurveTo(t._x1+t._k*(t._x2-t._x0),t._y1+t._k*(t._y2-t._y0),t._x2+t._k*(t._x1-e),t._y2+t._k*(t._y1-r),t._x2,t._y2)}function gt(t,e){this._context=t,this._k=(1-e)/6}gt.prototype={areaStart:function(){this._line=0},areaEnd:function(){this._line=NaN},lineStart:function(){this._x0=this._x1=this._x2=this._y0=this._y1=this._y2=NaN,this._point=0},lineEnd:function(){switch(this._point){case 2:this._context.lineTo(this._x2,this._y2);break;case 3:mt(this,this._x1,this._y1)}(this._line||0!==this._line&&1===this._point)&&this._context.closePath(),this._line=1-this._line},point:function(t,e){switch(t=+t,e=+e,this._point){case 0:this._point=1,this._line?this._context.lineTo(t,e):this._context.moveTo(t,e);break;case 1:this._point=2,this._x1=t,this._y1=e;break;case 2:this._point=3;default:mt(this,t,e)}this._x0=this._x1,this._x1=this._x2,this._x2=t,this._y0=this._y1,this._y1=this._y2,this._y2=e}};var vt=function t(e){function r(t){return new gt(t,e)}return r.tension=function(e){return t(+e)},r}(0);function yt(t,e){this._context=t,this._k=(1-e)/6}yt.prototype={areaStart:lt,areaEnd:lt,lineStart:function(){this._x0=this._x1=this._x2=this._x3=this._x4=this._x5=this._y0=this._y1=this._y2=this._y3=this._y4=this._y5=NaN,this._point=0},lineEnd:function(){switch(this._point){case 1:this._context.moveTo(this._x3,this._y3),this._context.closePath();break;case 2:this._context.lineTo(this._x3,this._y3),this._context.closePath();break;case 3:this.point(this._x3,this._y3),this.point(this._x4,this._y4),this.point(this._x5,this._y5)}},point:function(t,e){switch(t=+t,e=+e,this._point){case 0:this._point=1,this._x3=t,this._y3=e;break;case 1:this._point=2,this._context.moveTo(this._x4=t,this._y4=e);break;case 2:this._point=3,this._x5=t,this._y5=e;break;default:mt(this,t,e)}this._x0=this._x1,this._x1=this._x2,this._x2=t,this._y0=this._y1,this._y1=this._y2,this._y2=e}};var xt=function t(e){function r(t){return new yt(t,e)}return r.tension=function(e){return t(+e)},r}(0);function bt(t,e){this._context=t,this._k=(1-e)/6}bt.prototype={areaStart:function(){this._line=0},areaEnd:function(){this._line=NaN},lineStart:function(){this._x0=this._x1=this._x2=this._y0=this._y1=this._y2=NaN,this._point=0},lineEnd:function(){(this._line||0!==this._line&&3===this._point)&&this._context.closePath(),this._line=1-this._line},point:function(t,e){switch(t=+t,e=+e,this._point){case 0:this._point=1;break;case 1:this._point=2;break;case 2:this._point=3,this._line?this._context.lineTo(this._x2,this._y2):this._context.moveTo(this._x2,this._y2);break;case 3:this._point=4;default:mt(this,t,e)}this._x0=this._x1,this._x1=this._x2,this._x2=t,this._y0=this._y1,this._y1=this._y2,this._y2=e}};var _t=function t(e){function r(t){return new bt(t,e)}return r.tension=function(e){return t(+e)},r}(0);function wt(t,e,r){var n=t._x1,i=t._y1,a=t._x2,o=t._y2;if(t._l01_a>1e-12){var s=2*t._l01_2a+3*t._l01_a*t._l12_a+t._l12_2a,l=3*t._l01_a*(t._l01_a+t._l12_a);n=(n*s-t._x0*t._l12_2a+t._x2*t._l01_2a)/l,i=(i*s-t._y0*t._l12_2a+t._y2*t._l01_2a)/l}if(t._l23_a>1e-12){var c=2*t._l23_2a+3*t._l23_a*t._l12_a+t._l12_2a,u=3*t._l23_a*(t._l23_a+t._l12_a);a=(a*c+t._x1*t._l23_2a-e*t._l12_2a)/u,o=(o*c+t._y1*t._l23_2a-r*t._l12_2a)/u}t._context.bezierCurveTo(n,i,a,o,t._x2,t._y2)}function Tt(t,e){this._context=t,this._alpha=e}Tt.prototype={areaStart:function(){this._line=0},areaEnd:function(){this._line=NaN},lineStart:function(){this._x0=this._x1=this._x2=this._y0=this._y1=this._y2=NaN,this._l01_a=this._l12_a=this._l23_a=this._l01_2a=this._l12_2a=this._l23_2a=this._point=0},lineEnd:function(){switch(this._point){case 2:this._context.lineTo(this._x2,this._y2);break;case 3:this.point(this._x2,this._y2)}(this._line||0!==this._line&&1===this._point)&&this._context.closePath(),this._line=1-this._line},point:function(t,e){if(t=+t,e=+e,this._point){var r=this._x2-t,n=this._y2-e;this._l23_a=Math.sqrt(this._l23_2a=Math.pow(r*r+n*n,this._alpha))}switch(this._point){case 0:this._point=1,this._line?this._context.lineTo(t,e):this._context.moveTo(t,e);break;case 1:this._point=2;break;case 2:this._point=3;default:wt(this,t,e)}this._l01_a=this._l12_a,this._l12_a=this._l23_a,this._l01_2a=this._l12_2a,this._l12_2a=this._l23_2a,this._x0=this._x1,this._x1=this._x2,this._x2=t,this._y0=this._y1,this._y1=this._y2,this._y2=e}};var kt=function t(e){function r(t){return e?new Tt(t,e):new gt(t,0)}return r.alpha=function(e){return t(+e)},r}(.5);function At(t,e){this._context=t,this._alpha=e}At.prototype={areaStart:lt,areaEnd:lt,lineStart:function(){this._x0=this._x1=this._x2=this._x3=this._x4=this._x5=this._y0=this._y1=this._y2=this._y3=this._y4=this._y5=NaN,this._l01_a=this._l12_a=this._l23_a=this._l01_2a=this._l12_2a=this._l23_2a=this._point=0},lineEnd:function(){switch(this._point){case 1:this._context.moveTo(this._x3,this._y3),this._context.closePath();break;case 2:this._context.lineTo(this._x3,this._y3),this._context.closePath();break;case 3:this.point(this._x3,this._y3),this.point(this._x4,this._y4),this.point(this._x5,this._y5)}},point:function(t,e){if(t=+t,e=+e,this._point){var r=this._x2-t,n=this._y2-e;this._l23_a=Math.sqrt(this._l23_2a=Math.pow(r*r+n*n,this._alpha))}switch(this._point){case 0:this._point=1,this._x3=t,this._y3=e;break;case 1:this._point=2,this._context.moveTo(this._x4=t,this._y4=e);break;case 2:this._point=3,this._x5=t,this._y5=e;break;default:wt(this,t,e)}this._l01_a=this._l12_a,this._l12_a=this._l23_a,this._l01_2a=this._l12_2a,this._l12_2a=this._l23_2a,this._x0=this._x1,this._x1=this._x2,this._x2=t,this._y0=this._y1,this._y1=this._y2,this._y2=e}};var Mt=function t(e){function r(t){return e?new At(t,e):new yt(t,0)}return r.alpha=function(e){return t(+e)},r}(.5);function St(t,e){this._context=t,this._alpha=e}St.prototype={areaStart:function(){this._line=0},areaEnd:function(){this._line=NaN},lineStart:function(){this._x0=this._x1=this._x2=this._y0=this._y1=this._y2=NaN,this._l01_a=this._l12_a=this._l23_a=this._l01_2a=this._l12_2a=this._l23_2a=this._point=0},lineEnd:function(){(this._line||0!==this._line&&3===this._point)&&this._context.closePath(),this._line=1-this._line},point:function(t,e){if(t=+t,e=+e,this._point){var r=this._x2-t,n=this._y2-e;this._l23_a=Math.sqrt(this._l23_2a=Math.pow(r*r+n*n,this._alpha))}switch(this._point){case 0:this._point=1;break;case 1:this._point=2;break;case 2:this._point=3,this._line?this._context.lineTo(this._x2,this._y2):this._context.moveTo(this._x2,this._y2);break;case 3:this._point=4;default:wt(this,t,e)}this._l01_a=this._l12_a,this._l12_a=this._l23_a,this._l01_2a=this._l12_2a,this._l12_2a=this._l23_2a,this._x0=this._x1,this._x1=this._x2,this._x2=t,this._y0=this._y1,this._y1=this._y2,this._y2=e}};var Et=function t(e){function r(t){return e?new St(t,e):new bt(t,0)}return r.alpha=function(e){return t(+e)},r}(.5);function Lt(t){this._context=t}function Ct(t){return t<0?-1:1}function Pt(t,e,r){var n=t._x1-t._x0,i=e-t._x1,a=(t._y1-t._y0)/(n||i<0&&-0),o=(r-t._y1)/(i||n<0&&-0),s=(a*i+o*n)/(n+i);return(Ct(a)+Ct(o))*Math.min(Math.abs(a),Math.abs(o),.5*Math.abs(s))||0}function It(t,e){var r=t._x1-t._x0;return r?(3*(t._y1-t._y0)/r-e)/2:e}function Ot(t,e,r){var n=t._x0,i=t._y0,a=t._x1,o=t._y1,s=(a-n)/3;t._context.bezierCurveTo(n+s,i+s*e,a-s,o-s*r,a,o)}function zt(t){this._context=t}function Dt(t){this._context=new Rt(t)}function Rt(t){this._context=t}function Ft(t){this._context=t}function Bt(t){var e,r,n=t.length-1,i=new Array(n),a=new Array(n),o=new Array(n);for(i[0]=0,a[0]=2,o[0]=t[0]+2*t[1],e=1;e<n-1;++e)i[e]=1,a[e]=4,o[e]=4*t[e]+2*t[e+1];for(i[n-1]=2,a[n-1]=7,o[n-1]=8*t[n-1]+t[n],e=1;e<n;++e)r=i[e]/a[e-1],a[e]-=r,o[e]-=r*o[e-1];for(i[n-1]=o[n-1]/a[n-1],e=n-2;e>=0;--e)i[e]=(o[e]-i[e+1])/a[e];for(a[n-1]=(t[n]+i[n-1])/2,e=0;e<n-1;++e)a[e]=2*t[e+1]-i[e+1];return[i,a]}function Nt(t,e){this._context=t,this._t=e}function jt(t,e){if((i=t.length)>1)for(var r,n,i,a=1,o=t[e[0]],s=o.length;a<i;++a)for(n=o,o=t[e[a]],r=0;r<s;++r)o[r][1]+=o[r][0]=isNaN(n[r][1])?n[r][0]:n[r][1]}function Ut(t){for(var e=t.length,r=new Array(e);--e>=0;)r[e]=e;return r}function Vt(t,e){return t[e]}function Ht(t){var e=t.map(qt);return Ut(t).sort((function(t,r){return e[t]-e[r]}))}function qt(t){for(var e,r=-1,n=0,i=t.length,a=-1/0;++r<i;)(e=+t[r][1])>a&&(a=e,n=r);return n}function Gt(t){var e=t.map(Yt);return Ut(t).sort((function(t,r){return e[t]-e[r]}))}function Yt(t){for(var e,r=0,n=-1,i=t.length;++n<i;)(e=+t[n][1])&&(r+=e);return r}Lt.prototype={areaStart:lt,areaEnd:lt,lineStart:function(){this._point=0},lineEnd:function(){this._point&&this._context.closePath()},point:function(t,e){t=+t,e=+e,this._point?this._context.lineTo(t,e):(this._point=1,this._context.moveTo(t,e))}},zt.prototype={areaStart:function(){this._line=0},areaEnd:function(){this._line=NaN},lineStart:function(){this._x0=this._x1=this._y0=this._y1=this._t0=NaN,this._point=0},lineEnd:function(){switch(this._point){case 2:this._context.lineTo(this._x1,this._y1);break;case 3:Ot(this,this._t0,It(this,this._t0))}(this._line||0!==this._line&&1===this._point)&&this._context.closePath(),this._line=1-this._line},point:function(t,e){var r=NaN;if(e=+e,(t=+t)!==this._x1||e!==this._y1){switch(this._point){case 0:this._point=1,this._line?this._context.lineTo(t,e):this._context.moveTo(t,e);break;case 1:this._point=2;break;case 2:this._point=3,Ot(this,It(this,r=Pt(this,t,e)),r);break;default:Ot(this,this._t0,r=Pt(this,t,e))}this._x0=this._x1,this._x1=t,this._y0=this._y1,this._y1=e,this._t0=r}}},(Dt.prototype=Object.create(zt.prototype)).point=function(t,e){zt.prototype.point.call(this,e,t)},Rt.prototype={moveTo:function(t,e){this._context.moveTo(e,t)},closePath:function(){this._context.closePath()},lineTo:function(t,e){this._context.lineTo(e,t)},bezierCurveTo:function(t,e,r,n,i,a){this._context.bezierCurveTo(e,t,n,r,a,i)}},Ft.prototype={areaStart:function(){this._line=0},areaEnd:function(){this._line=NaN},lineStart:function(){this._x=[],this._y=[]},lineEnd:function(){var t=this._x,e=this._y,r=t.length;if(r)if(this._line?this._context.lineTo(t[0],e[0]):this._context.moveTo(t[0],e[0]),2===r)this._context.lineTo(t[1],e[1]);else for(var n=Bt(t),i=Bt(e),a=0,o=1;o<r;++a,++o)this._context.bezierCurveTo(n[0][a],i[0][a],n[1][a],i[1][a],t[o],e[o]);(this._line||0!==this._line&&1===r)&&this._context.closePath(),this._line=1-this._line,this._x=this._y=null},point:function(t,e){this._x.push(+t),this._y.push(+e)}},Nt.prototype={areaStart:function(){this._line=0},areaEnd:function(){this._line=NaN},lineStart:function(){this._x=this._y=NaN,this._point=0},lineEnd:function(){0<this._t&&this._t<1&&2===this._point&&this._context.lineTo(this._x,this._y),(this._line||0!==this._line&&1===this._point)&&this._context.closePath(),this._line>=0&&(this._t=1-this._t,this._line=1-this._line)},point:function(t,e){switch(t=+t,e=+e,this._point){case 0:this._point=1,this._line?this._context.lineTo(t,e):this._context.moveTo(t,e);break;case 1:this._point=2;default:if(this._t<=0)this._context.lineTo(this._x,e),this._context.lineTo(t,e);else{var r=this._x*(1-this._t)+t*this._t;this._context.lineTo(r,this._y),this._context.lineTo(r,e)}}this._x=t,this._y=e}},t.arc=function(){var t=m,o=g,w=r(0),T=null,k=v,A=y,M=x,S=null;function E(){var r,m,g=+t.apply(this,arguments),v=+o
```
